# The evolutionary dynamics of genetic mutational load throughout tomato domestication history

**DOI:** 10.1101/2021.11.08.467620

**Authors:** Hamid Razifard, Sofia Visa, Denise Tieman, Esther van der Knaap, Ana L. Caicedo

## Abstract

Understanding the evolution of deleterious mutations through domestication has fascinated evolutionary biologists and breeders alike. Some domesticated organisms have been reported to accumulate deleterious mutations, i.e. radical amino acid changes, through their domestication history (“cost of domestication”). However, more recent evidence paints a more complex picture of this phenomenon in different domesticated organisms. In this study, we used genomic sequences of 253 tomato accessions to investigate the evolution of deleterious mutations and genomic structural variants (SVs) through tomato domestication history. Specifically, we used phylogeny-based methods to identify deleterious mutations in the cultivated tomato as well as its closely related semi-wild and wild populations. We also explored a potential correlation between deleterious mutations and SVs. To create a functional link between deleterious alleles and phenotypes of interest for tomato breeding, we also conducted Genome-wide Association Studies (GWAS) on several fruit volatiles.

Our results implicate a downward trend, throughout tomato domestication history, in diversity of most alleles, regardless of their functional impact. This suggests that demographic factors, such as bottleneck events and inbreeding, have reduced overall genetic diversity, leading to lower deleterious load and SVs as well as loss of some beneficial alleles during tomato domestication. We also detected an increase in proportions of nonsynonymous and deleterious alleles (relative to synonymous and neutral nonsynonymous alleles, respectively) during the initial stage of tomato domestication in Ecuador, although the final stage of tomato domestication in Mexico did not seem to involve such an increase. However, deleterious alleles in cultivated tomato seem to be more frequent than expected by neutral theory of molecular evolution. Additionally, for all tomato populations, we found a higher proportion of deleterious mutations in genomic regions impacted by SVs.

Our analyses also revealed frequent deleterious alleles in several well-studied tomato genes, probably involved in response to biotic and abiotic stress as well as fruit development and flavor regulation. Also, through genome-wide association studies (GWAS), we discovered deleterious alleles associated with two volatiles: isobutyl acetate, which is important for tomato fruit flavor, and methyl salicylate, involved in disease resistance and regulating flowering time. To provide a practical guide for breeding experiments, we created TomDel, a public searchable database of 21,162 deleterious alleles identified in this study (https://github.com/hrazif/TomDel-0.1)

## Introduction

Understanding the circumstances that allow proliferation of deleterious mutations in genomes is a fascinating evolutionary question and one of great relevance for breeding applications. In recent decades, several studies have reported accumulation of deleterious mutations during domestication of both plants (e.g. Lu et al. 2006; Liu et al. 2017; Zhou et al. 2017) and animals (e.g. Schubert et al. 2014; Marsden et al. 2016), often referred to as “cost of domestication” (Moyers et al. 2017). A recent review of studies on deleterious mutations in two cultivated plants, i.e. soybean and rice, as well as five domesticated animals, i.e. dog, pig, rabbit, chicken, and silkworm (Makino et al. 2018) concluded that, except in pigs, the studied organisms showed elevated proportions of nonsynonymous mutations, when compared with their wild relatives. The exception observed in pigs was attributed to absence of strong bottleneck during pig domestication as well as possible gene flow with its wild ancestors. Similarly, a study on sunflower (Renaut and Rieseberg 2015) revealed a higher proportion of deleterious alleles in the domesticated sunflower, although the absolute number of deleterious mutations was similar to that of its wild ancestor. In tomato, a prior study has implied accumulation of deleterious mutations during tomato domestication, based on a transcriptomic comparison between cultivated and wild tomato species (Koenig et al. 2013).

A comprehensive understanding of evolution of deleterious mutations through domestication requires population-wide genomic surveys based on comparisons between domesticated organisms and their closely related wild and semi-domesticated populations. Adopting such approach, more recent studies have and revealed complexities in the evolutionary patterns of deleterious mutations in a small number of domesticated organisms. For example, in a study by Lozano et al. (2021), the authors estimated mutational load (the sum of derived deleterious alleles) in sorghum and maize. That study revealed that, contrary to maize, domesticated sorghum had a lower number of sites with deleterious alleles than its wild relative. Therefore, the fate of deleterious mutations might differ even between closely related species. Similar to sorghum, domesticated soybean was recently reported to harbor fewer deleterious alleles than its wild relatives (Kim et al. 2021), although prior studies had implied the opposite pattern (Lam et al. 2010). A difficulty in comparing prior studies has been lack of consensus on definitions and methods to evaluate deleterious alleles, which has also been a focus on recent studies.

The mutational load, defined as reduction in fitness due to harmful mutations (Kimura 1983), is largely shaped by the cumulative count of deleterious mutations occurring in a population (see Moyers et al. 2017 and references therein). The mutational load of a population is shaped by a unique combination of demographic and genomic factors. One of the major demographic processes impacting deleterious mutations through domestication is reduction in effective population size due to bottleneck events, which leads to more genetic drift and less efficient selection (Gaut et al. 2018), thus predicted to allow deleterious alleles to persist. This is more evident in populations that have undergone a recent bottleneck event. However, sustained bottleneck events can lead to removal of deleterious mutations, along with neutral and beneficial alleles, by lowering overall genetic diversity in the population (Bortoluzzi et al. 2020; Grossen et al. 2020). Additionally, a transition to an inbreeding mating system, which is a common phenomenon in domesticated organisms, can more efficiently remove recessive deleterious mutations from the population by exposing them to selection through an increase in homozygosity (Byers and Waller 1999).

Domesticated organisms often experience rapid population growth as humans expand their use in new geographic regions; the action of drift on the expanding edges of the population can lead to accumulation of new deleterious mutations via “gene surfing” (Klopfstein et al. 2006; Travis et al. 2007). Asexual reproduction, e.g. clonal propagation in plants, can also lead to accumulation of deleterious mutations, due to lack of crossing-over in asexually reproducing organisms. This phenomenon is often referred to as “Muller’s ratchet” (Muller 1932) and has been reported in plants, e.g. cassava (Ramu et al. 2017), animals, e.g. Bdelloid rotifers (Barraclough et al. 2007), and bacteria (Elena and Lenski 1997).

As a genomic factor influencing the evolution of deleterious mutations, genomic structural variants (SVs) can disrupt genes, which might negatively affect the fitness of an organism (reviewed by Yuan et al. (2021). With the rapid advances in genomic research in recent years, several studies have revealed valuable insights on the prevalence of SVs in different plant crops, e.g. *Brassica napus* L. (Chawla et al. 2021), rice (Fuentes et al. 2019), grapevine (Zhou et al. 2019), soybean (McHale et al. 2012; Anderson et al. 2014), maize (Mahmoud et al. 2020); Yang et al. 2019), and tomato (Wang et al. 2020; Alonge et al. 2020; Domínguez et al. 2020). However, the relationship between SVs and deleterious mutations has rarely been studied. Notably, a study by (Hämälä et al. 2021) on the chocolate tree (*Theobroma cacao*) revealed higher genetic load in genomic regions impacted by SVs.

Moreover, the degree of genetic linkage (infrequent crossing-over between genomic regions with smaller physical distance or due to other variations in recombination rate) is also an important factor in the prevalence of deleterious mutations in domesticated organisms. Due to linkage, artificial selection on favorable traits in the organisms under domestication can lead to accumulation of deleterious mutations within or near genomic regions experiencing selection; this phenomenon is referred to as “genetic hitchhiking” (Barton 1995).

Considering the complex interactions of demographic and genomic factors operating on a population, a robust analysis of mutational load in any domesticated organism must be placed within the unique context of the domestication history of that organism. Currently available methods for detecting deleterious mutations (Moyers et al. 2017) are based on comparing protein sequences from an organism of interest and the orthologous protein sequences from closely related species (Ng and Henikoff 2003; Choi et al. 2012; Kono et al. 2016). The underlying assumption of these methods is that amino acid changes in positions that are highly conserved phylogenetically are considered to be deleterious.

To date, population-level insight on deleterious mutations, especially in the intermediate stages of domestication between wild and domesticated populations, is lacking for many crops, including the common cultivated tomato. Insights on the evolution of deleterious mutations through tomato domestication could have immense economical value considering tomato’s status as the world’s most valuable vegetable crop (FAO 2013). The common cultivated tomato has a complex domestication history, with several intermediate populations collected in Latin America (Blanca et al. 2015; Razifard et al. 2020). These populations have experienced different demographic fates; thus a thorough analysis of deleterious load in tomato entails special attention to population genomic factors affecting deleterious load in the intermediate stages as well as the functions of genes impacted by deleterious mutations.

In this study, we investigate the evolution of deleterious mutations through tomato domestication history, with special attention to the evolution of deleterious mutations in the context of SVs. Specifically, we used population genomic tools to a) create novel insights on mutational load in the cultivated tomato and its closely related populations b) track the trajectories of deleterious alleles within wild, semi-domesticated, and domesticated tomato populations, c) illuminate potential correlation between SVs and deleterious mutations d) create a functional link between deleterious mutations in the cultivated tomato and phenotypes of interest for breeding purposes.

## Results

### Population history and statistics

We used a dataset of 15,011,193 single nucleotide polymorphisms (SNPs) obtained from genome-wide sequencing of a diverse set of tomato accessions (295 accessions), as described in Razifard et al. (2020), to investigate the dynamics of deleterious mutations through tomato domestication history. These accessions included the closest fully wild relative of the cultivated tomato, i.e. *Solanum pimpinellifolium* L. (SP), from South America; the intermediate group between SP and the cultivated tomato, i.e. *S. lycoperiscum* L. var. *cerasiforme* (SLC), from South America and Mesoamerica; landraces of the domesticated tomato, i.e. *S. lycopersicum* L. var. *lycopersicum* (SLL), from the Americas as well as its modern varieties grown worldwide; and one accession of *Solanum pennellii* L., as an outgroup.

A maximum-likelihood population tree was built using a TreeMix analysis (fig. 1a) of the populations identified in Razifard et al. (2020) with two exceptions; 1-for simplicity and to ensure sufficient sample size, all SP populations were considered as one group; 2-SLL was split into two groups: “SLL Americas”, a group mostly limited to the Americas, and “SLL modern”, a group of modern tomato accessions. This tree and the geographic distribution of the accessions from each group (fig. 1b) recapitulate our current understanding of tomato domestication history in South America and Mesoamerica, which has been discussed in detail in Razifard et al. (2020). Briefly, the SLC are a highly diverse group both phenotypically and geographically, with SLC populations extending from South America to Central America and Mexico. Three SLC populations from South America (SLC ECU, SLC PER, and SLC San Martin) are considered semi-domesticated because they tend to display domesticated-like phenotypes. Of these SLC groups, SLC from Ecuador was likely the first intermediate group of tomatoes to diverge from wild SP. Two “weedy” SLC populations, one mostly distributed in Mexico (SLC MEX) and another with an extensive distribution ranging from Mexico to Central America and northern South America (SLC MEX-CA-NSA), are the northernmost extensions of SLC and tend to display some wild-like phenotypes (e.g. fruit size) similar to SP (Fig. 1a; Razifard et al. 2020). Finally, the closest relative of the cultivated tomato (SLL) is SLC MEX.

**Fig. 1.**
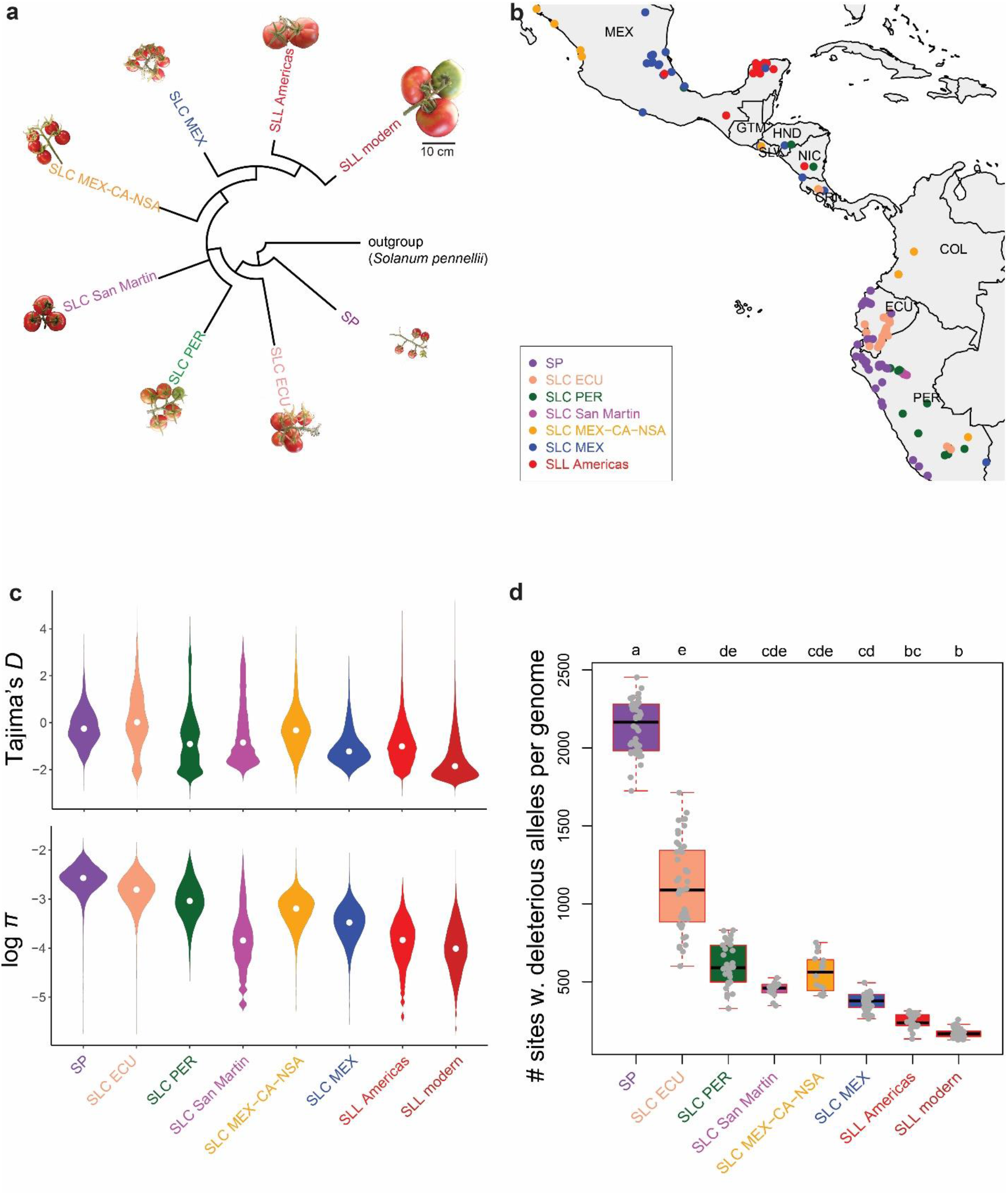
Tomato population history and statistics. (**a**) A maximum-likelihood phylogeny reconstructed using TreeMix. All nodes in the population tree received bootstrap support of 100%. (**b**) Distribution map of all tomato accessions included in this study, except SLL Modern, which have a worldwide distribution. (**c**) Evidence of selection (*Tajima’s D*) and nucleotide diversity (log *π*) calculated for each accession within all tomato populations included in this study. All pairwise comparisons between populations are statistically significant for *Tajima’s D* and *π* (p-value < 2 × 10^−16^ based on Dunn’s test). (**d**) Number of sites with derived alleles identified as deleterious by SIFT and PROVEAN in each of the tomato accessions within different populations. Results of pairwise statistical comparisons (Dunn’s test) have been presented with lowercase letters; populations with a shared letter did not have a significant difference in their distribution of the number of sites with deleterious alleles.

To gain insights on the overall frequency of alleles as well as genetic diversity in the populations studied here, we estimated nucleotide diversity (using *π*) as well as the deviation from a neutral allele frequency spectrum (using *Tajima’s D*) (Fig. 1c). Comparing median *π* between different populations reveals a general downward trend during the transition from SP to SLC and from SLC to SLL, implying bottleneck events through tomato domestication history. Also, an abundance of rare alleles, implied by negative median *Tajima’s D*, in SLL has been described previously (Razifard et al. 2020) However, median *Tajima’s D* in the populations surveyed here imply that abundance of rare alleles in SLL is primarily due to variants in “SLL modern”, i.e. tomato accessions collected from outside the centers of tomato domestication in South America and Mesoamerica.

### Assessing mutational load of different tomato populations

To assess the occurrence of deleterious mutations (i.e. the “mutational load”) in each tomato population, we explored the functional impact of each SNP using SIFT (Kumar et al. 2009) and Protein Variation Effect Analyzer (PROVEAN; Choi et al. 2012). To avoid reference bias, all variants in all tomato populations were polarized using a *Solanum pennellli* L. accession (outgroup) and only derived alleles were evaluated for their functional impact. Similar to other tools used for assessing variant impacts, SIFT and PROVEAN operate based on the theory that mutations occurring at evolutionary conserved coding regions of the genome are likely to be deleterious. However, these programs use different mathematical models for calculating a score for each variant. SIFT computes a score based on the frequency of observed amino acids in the protein coded by a certain gene, across the gene phylogeny, as well as the distribution of unobserved amino acids based on a Dirichlet mixture model. PROVEAN provides a score for each variant based on similarity of each protein sequence to a functional homolog and the change in similarity caused by incorporating each amino acid change to that protein. According to the PROVEAN model, amino acid changes that make a protein sequence less similar to a functional homolog are expected to be deleterious.

A total of 134,446 SNPs were identified as containing nonsynonymous alleles by SIFT, of which 53,222 were identified as sites containing potentially deleterious alleles (“deleterious” or “deleterious, low confidence”). The SIFT list of nonsynonymous sites was provided as input to PROVEAN, which identified 36,583 sites as potentially deleterious. In the final list of deleterious sites, we included 21,162 sites identified as potentially deleterious by both SIFT and PROVEAN, and we assessed the number of deleterious alleles carried by each tomato genotype. Similarly to nucleotide diversity (Fig. 1c), we observed a general reduction in the total number of derived deleterious alleles per genome throughout tomato domestication history (Fig. 1d; supplementary fig. 1a). The number of derived deleterious mutations per genome has been significantly reduced in the transition from SP to SLC and from SLC to SLL (Suppl. Fig. 1a), but a closer look at individual tomato populations presents a more nuanced view.

The number of derived deleterious alleles seems to have been significantly reduced during the divergence between SP and SLC ECU. Another significant reduction, compared to SLC ECU, seems to have occurred during the northward expansion of SLC to Mexico, represented by SLC MEX. Interestingly, the domestication of SLL from SLC MEX, represented by SLL Americas, does not seem to have entailed a significant change in the number of deleterious alleles per genome. However modern cultivated tomatoes seem to harbor a significantly lower number of deleterious mutations than SLC MEX, with most genomes surveyed containing just a few hundred deleterious nucleotide substitutions.

The number of derived alleles per genome from other categories (e.g. non-coding, synonymous, and non-synonymous) show a similar pattern of decrease in members of populations through tomato domestication history (supplementary fig. 1). In fact, the patterns of decrease all mirror each other and that of nucleotide diversity, suggesting that the drastic changes in genetic variation may be obscuring any patterns specific to deleterious mutations. We thus focused on the proportions of alleles from different mutational categories within each population, to detect any differences in the rates at which mutational loads have accumulated. Specifically we looked at the ratio of the number of sites with derived nonsynonymous alleles to the number of sites with synonymous alleles (*nonsyn./syn.*) per genome, as well as the ratio of the number of sites with derived deleterious alleles to the number of sites with neutral nonsynonymous alleles (*del./neutr. nonsyn.*; heretofore referred to as “relative mutational load”). These ratio estimates are expected to be more robust to the effects of decreased diversity, due to bottlenecks and inbreeding, than the absolute number of deleterious mutations.

Per-accession *nonsyn./syn.* (Fig. 2a) varied widely between groups, but seems to have significantly increased during the origin of SLC, represented by SLC ECU. In contrast, there are no significant differences in *nonsyn./syn.* between SLC ECU and the other two South American SLC populations (SLC PER and SLC San Martin). The northward spreads of SLC, represented by SLC MEX-CA-NSA and SLC MEX, seem to have involved significant increases in *nonsyn./syn.* ratio, compared to SLC ECU. Conversely, a significant reduction in *nonsyn./syn.* ratio in modern cultivated tomato (“SLL modern”) is observed compared to SLC MEX, although the variance in large. However, the divergence of SLL Americas from SLC MEX does not seem to have entailed a significant change in *nonsyn./syn.* ratio.

**Fig. 2.**
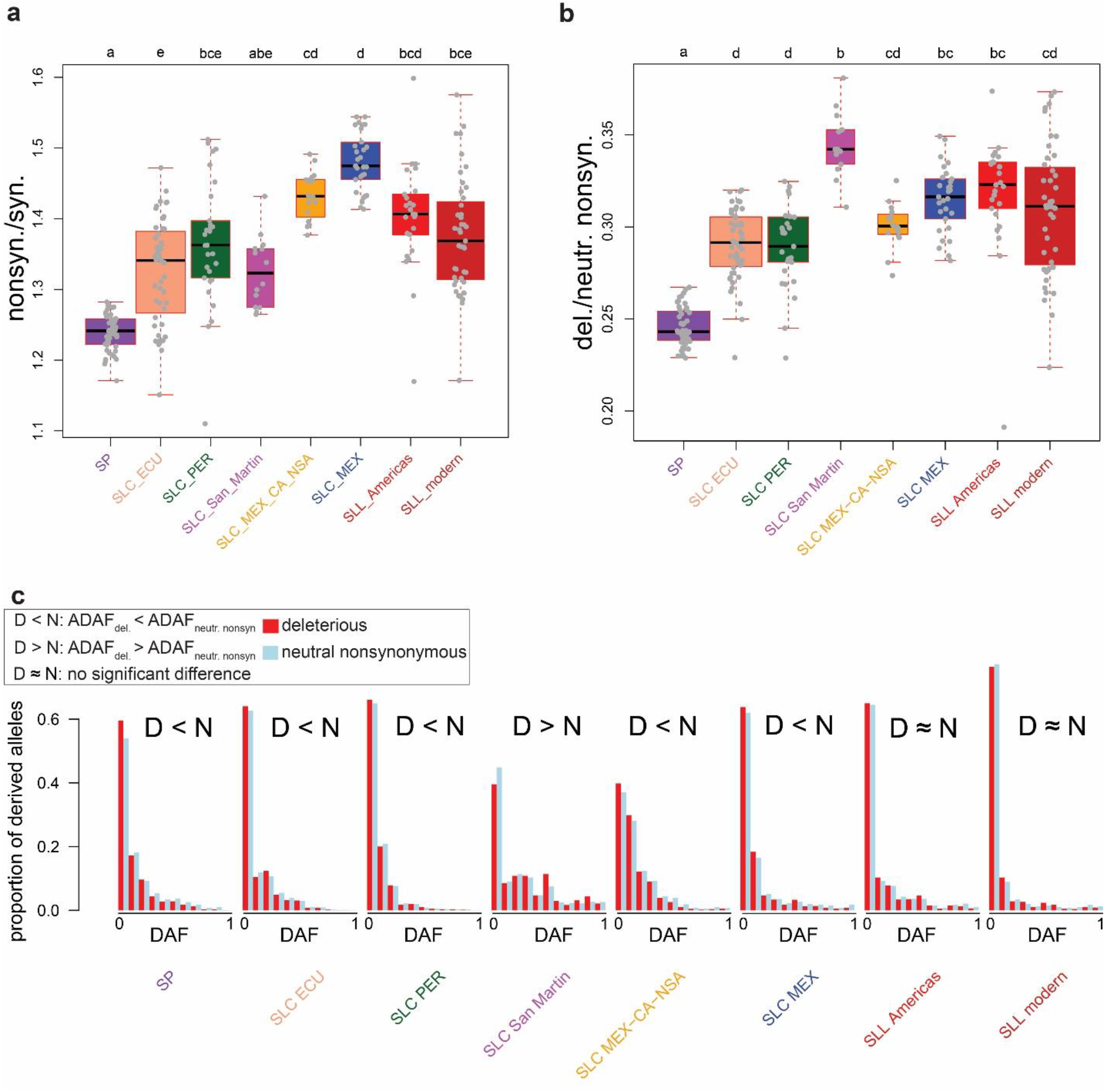
Prevalence of alleles with potentially high impact on fitness, compared with neutral alleles. (a) *nonsyn./syn.*, the ratio of the number of sites with nonsynonymous alleles to the number of sites with synonymous alleles in each accession categorized within different tomato populations. (b) del./neutr., the ratio of the number of sites with deleterious alleles to the number of sites with neutral nonsynonymous alleles (del/neutr) in each accession within different tomato populations. In both (a) and (b) results of a pairwise Dunn’s test are shown with lowercase letters (see Fig. 1 caption). (c) Site frequency spectra for derived deleterious (red) and derived neutral nonsynonymous alleles (blue) in each tomato population. For reach population, derived allele frequency (DAF) is shown in the range of 0 to 1, with bins of 0.1. Results of statistical tests of average derived allele frequency of deleterious alleles (ADAF_del_) versus average derived allele frequency of neutral nonsynonymous alleles (ADAF_neutr. nonsyn._) are shown with D < N (ADAF_del_ is smaller than ADAF_neutr. nonsyn._), D > N (ADAF_del_ is greater than ADAF_neutr. nonsyn._ l), and D ≈ N (ADAF_del_ is not significantly different from ADAF_neutr. nonsyn._).

While nonsynonymous substitutions have a higher probability of being deleterious than synonymous changes, the true mutational load in genomes is given by the proportion of nonsynonymous mutations deemed to be deleterious. We thus explored the patterns of per-accession relative mutational load among and between all the populations included in this study (Fig. 2b).Two major differences also appear when comparing the patterns of relative mutational load and *nonsyn./syn.* in tomato population (Figs. 2a–b). First, the northward spread of SLC (SLC MEX-CA-NSA) does not seem to have involved a significant change in relative mutational load compared to SLC ECU, although its *nonsyn./syn.* ratio is significantly higher than in SLC ECU. This incongruence might suggest that the first northward spread of SLC might have entailed release from artificial selection pressure, probably leading to positive selection on nonsynonymous substitutions that might have been beneficial for the survival of this population in wild-like habitats. Second, compared to SLC ECU, SLC San Martin seems to harbor a significantly higher relative mutational load (Fig. 2b), although its average *nonsyn./syn.* is not significantly different from SLC ECU. Finally, there seems to be no significant difference in relative mutational load between the two SLL populations and their closely related populations, SLC MEX-CA-NSA and SLC MEX (Fig. 2b), despite the reduction in *nonsyn./syn.* in modern SLL in comparison with SLC MEX (Fig. 2a). This implies that the origin of the cultivated tomato in Mexico did not entail an increase in relative mutational load. However, we observed a very high variance in proportions of nonsynonymous and deleterious alleles among modern SLL accessions.

### The frequency spectra of deleterious mutations in tomato populations

We further investigated the frequencies of derived neutral nonsynonymous and derived deleterious alleles in each tomato population (Fig. 2c). In theory, deleterious alleles are expected to occur at low frequency in populations under drift-selection equilibrium due to the action of background selection which removes deleterious variants more readily than neutral variants. However, deleterious alleles that “hitchhike” with positively selected variants can reach higher frequency in populations, and the reduced efficiency of selection in populations with small effective population size can also lead to higher-than-expected deleterious allele frequencies.

In most tomato populations studied here, average derived allele frequency of deleterious alleles (ADAF_del_) is smaller than average derived allele frequency of neutral nonsynonymous alleles (ADAF_neutr. nonsyn_) (Fig. 2c). Three populations deviated from this pattern: first, SLC San Martin has a higher ADAF_del_ than ADAF_neutr. nonsyn._ Second, the two SLL populations show no significant difference in ADAF_del_ vs. ADAF_neutr. nonsyn._, indicating that despite low relative mutational loads, the deleterious alleles that exist in these populations are relatively common across accessions.

For each accession within a certain population, we also calculated the proportion of sites with derived deleterious alleles that are private to the population to which that accession belongs i.e. #Priv_del_./#All_del_.. Similarly, we calculated #Priv_neutr. nonsyn._/#All _neutr. nonsyn_. for each individual (supplementary fig. 2). In all populations, except SP, a higher proportion of deleterious alleles than neutral nonsynonymous alleles were found to be private. This is consistent with the expectation that deleterious alleles are more readily removed by selection and thus are less likely to be shared with other populations. This does not hold true for SP, in which deleterious alleles are as likely as neutral alleles to be shared with other populations. This observation may be attributed to higher levels of heterozygosity in SP, which may mask the effects of some deleterious alleles to selection.

Despite a high proportion of deleterious alleles being private in SLC and SLL, we found that the no significant difference in the derived allele frequency of private deleterious and private neutral nonsynonymous mutations in SLC and SLL populations (supplementary fig. 3). In other words, private deleterious alleles are more common than expected within each SLC and SLL population. In SP, private deleterious alleles were not more frequent than expected (supplementary figs. 2-3), confirming that, although some deleterious alleles may be escaping selection, deleterious alleles overall are more efficiently purged out of a population in the wild, than under domestication. Thus, an increase in mutational load during origin of SLC and subsequent tomato domestication seems to be manifested more strongly in private alleles, than those shared between populations.

Deleterious mutations might accumulate near genomic regions under positive selection due to genetic hitchhiking via linkage disequilibrium (Moyers et al. 2017). Also, phylogenetically conserved regions of the genome are expected to harbor lower mutational load, due to the highly negative impact of deleterious mutations in such regions on the organism’s fitness. To test these and provide insights on the genomic factors affecting mutational load in tomato populations, we calculated relative mutational load in genomic windows of 100 kb for all the accessions. For each window, we also estimated recombination rate (*r*), nucleotide diversity (*π*), and signals of selective sweeps, using SweeD likelihood ratio (see Methods).

In all populations, genomic windows with higher nucleotide diversity tend to have higher relative mutational load (Table 1). Also, in all populations, except SLC San Martin and the two SLL groups, windows with lower recombination rates tend to, on average, have higher relative mutational load, suggesting that recombination rates do generally affect the ability to purge deleterious mutations. However, in all populations except SLL Americas, in which, genomic regions with signals of selective sweeps (high SweeD likelihood ratio) tend to have lower relative mutational load.

**Table 1.**
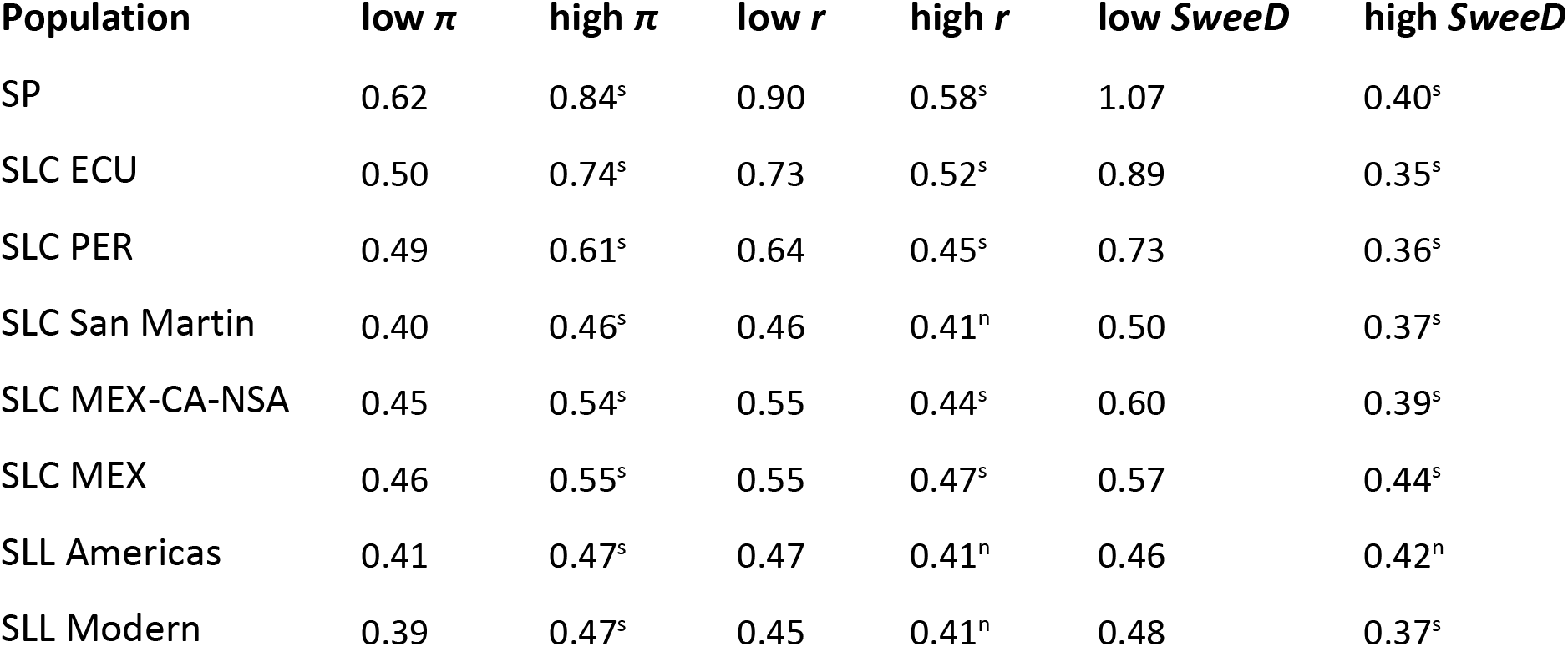
A comparison of average relative mutational load, across accessions, in genomic windows with different levels of nucleotide diversity (*π*), recombination (*r*), and selective sweep signal (*SweeD*). Results of Wilcoxon tests on windows with high and low average *π*, *r*, and *SweeD* are shown with “s” (significant) and “n” (non-significant).

### Structural variants and their impact on mutational load

We identified structural variants using three pipelines: Lumpy (v. 0.3.0; Layer et al., 2014), Delly (v. 0.8.7; Rausch et al., 2012), and Manta (v. 1.6.0; Chen et al., 2016). We then used SURVIVOR (v.1.0.7; Jeffares et al., 2017) to keep SVs present in the output from at least two of the three pipelines. In total, we kept 1,809,695 SVs, with median length of SVs = 418 bp. However, many of these SV were of the same type and had a similar genomic position and length. Therefore, we merged the genotypes from such SVs (see Materials and Methods), leaving 426,624 unique SVs. Further filtering of low-frequency SVs (minimum allele frequency < 0.01) resulted in 97,654 SVs, which were used in the downstream analyses. Derived alleles were determined relative to the genotype observed in the outgroup, *S. pennellii*.

Our estimates of the number of derived SVs from different categories (deletions, translocations, duplications, and inversions) implies a reduction in all categories of derived SVs, relative to *S. pennellii*, through tomato domestication history, (Fig. 3a). This is consistent with the pattern observed for deleterious mutations as well as mutations belonging to other categories (fig. 1d and supplementary fig. 1). We also conducted a comparison between genic and non-genic regions of the tomato genome for the prevalence of SVs. We detected more SVs in non-genic regions (supplementary fig. 4), compared to genic regions (i.e. SVs within, containing, or partially overlapping with genes). Also, among the tomato SVs in genic regions, on average, 61% were detected within intronic and untranslated regions (supplementary fig. 5), which suggests strong selection against exons disrupted by SVs.

**Fig. 3.**
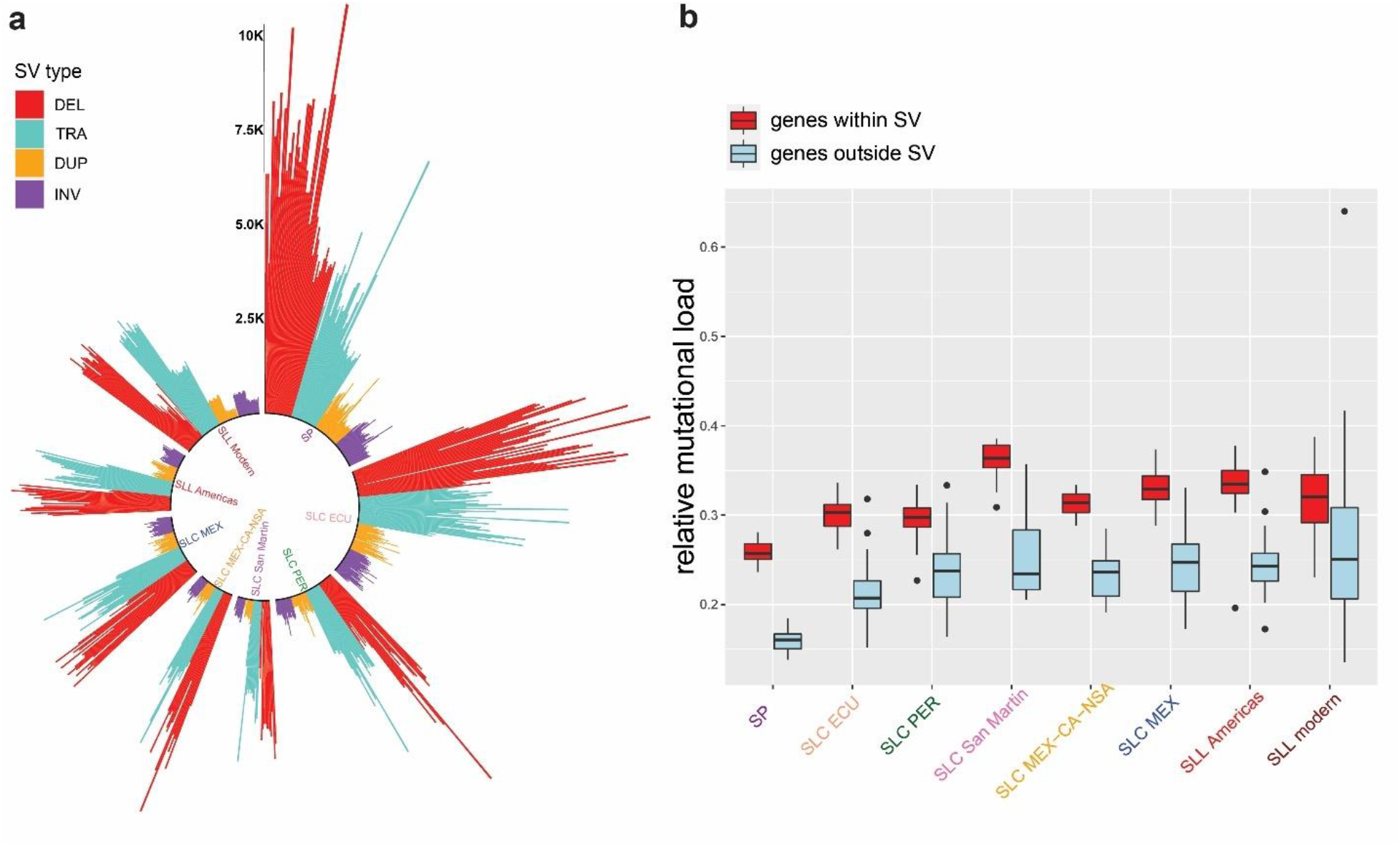
The evolution of structural variants (SVs) during tomato domestication and their association with mutational load. (a) The number of SVs from different categories (DEL: deletions, TRA: translocations, DUP: duplications, and INV: inversions) identified in each accession belonging to different tomato groups. Each colored bar represents one accession (b) A genome-wide comparison between mutational load within and outside SV in each population. The comparisons within all populations are statistically significant (based a T-test; p-value < 0.001).

To assess a potential correlation between relative mutational load and genomic regions where SVs are found in some accessions within a population, we compared relative mutational load in genes where SVs occur in comparison with genes outside SV regions. This analysis revealed a significantly higher relative mutational load in genes within SV regions (Fig. 3b; T-test, p-value < 0.001).

### Deleterious mutations of interest for tomato breeding

To better understand the journeys of each deleterious allele throughout tomato domestication history, we estimated genotype frequencies of all deleterious alleles in modern tomato as well as its closely related populations. These results are provided in a publicly accessible database named TomDel (available from https://github.com/hrazif/TomDel-0.1). Genotype frequency plots for tomato genes can be searched with gene IDs, e.g. Solyc01g005000, using the “Go to file” option. For some well-studied tomato genes, we compared the proportions of deleterious genotypes between different tomato populations (supplementary file 1; supplementary table 1), which revealed that deleterious alleles most often have become less frequent during the transition from SP to SLC and many deleterious alleles are absent in SLL Modern. This might suggest that, at least for these genes, the original domestication of tomato in Ecuador has been highly effective in removing alleles that are detrimental to the fitness of the tomato populations.

We further explored our dataset for sites with deleterious alleles that are present in SLL Modern. These are often the alleles of highest interest for crop improvement. Of 21,162 sites with deleterious alleles in the entire dataset, 1,677 sites contained deleterious alleles in modern SLL. A Gene Ontology (GO) term analysis on deleterious alleles present in modern SLL (supplementary table 2) revealed mostly broad term such as “metabolic process” and “oxidation-reduction process”, which implies that genes with a broad range of functions have been impacted by deleterious mutations. However, we also found more specific GO terms such as “inositol biosynthetic process”, related to ascorbic acid pathway (Calafiore et al. 2016) and “spermidine biosynthetic process”, related to pollen germination under heat stress (Song et al. 2002).

Among sites with deleterious alleles present in modern SLL, 184 sites had considerable deleterious genotype frequency ( > 0.25), and in 103 sites, the deleterious homozygous genotypes were the common genotypes (frequency > 0.5; supplementary tables 3-4). We detected genes with common deleterious genotypes involved in disease resistance, response to heat stress, fruit development, and fruit flavor (Fig. 4, Table 2). Notably, we found a common deleterious allele in *SlAAT4*, which might be involved in regulating tomato fruit flavor through biosynthesis of fruit volatiles esters.

**Fig. 4.**
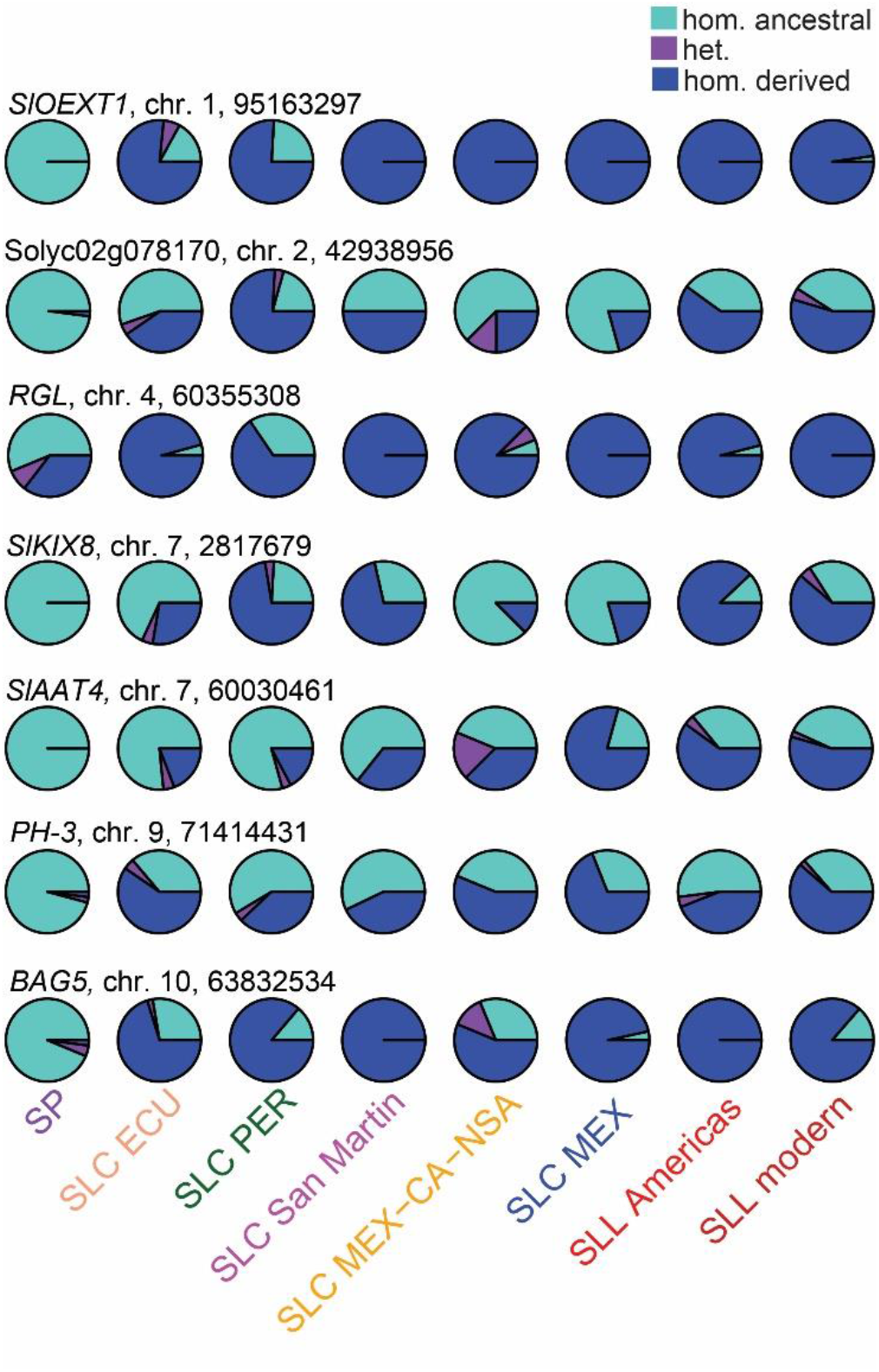
A population-level comparison of genotype frequencies of some example sites with common deleterious alleles (allele frequency > 0.5) in modern SLL. These example sites represent those within well-studied tomato genes or those with a known ortholog in other plants. Genotype frequencies are differentiated as follows: homozygous derived deleterious in blue; homozygous ancestral in turquoise; and heterozygous in purple. The ancestral state is assumed to be the allele present in the outgroup, *S. pennellii* (see Methods).

**Table 2.**
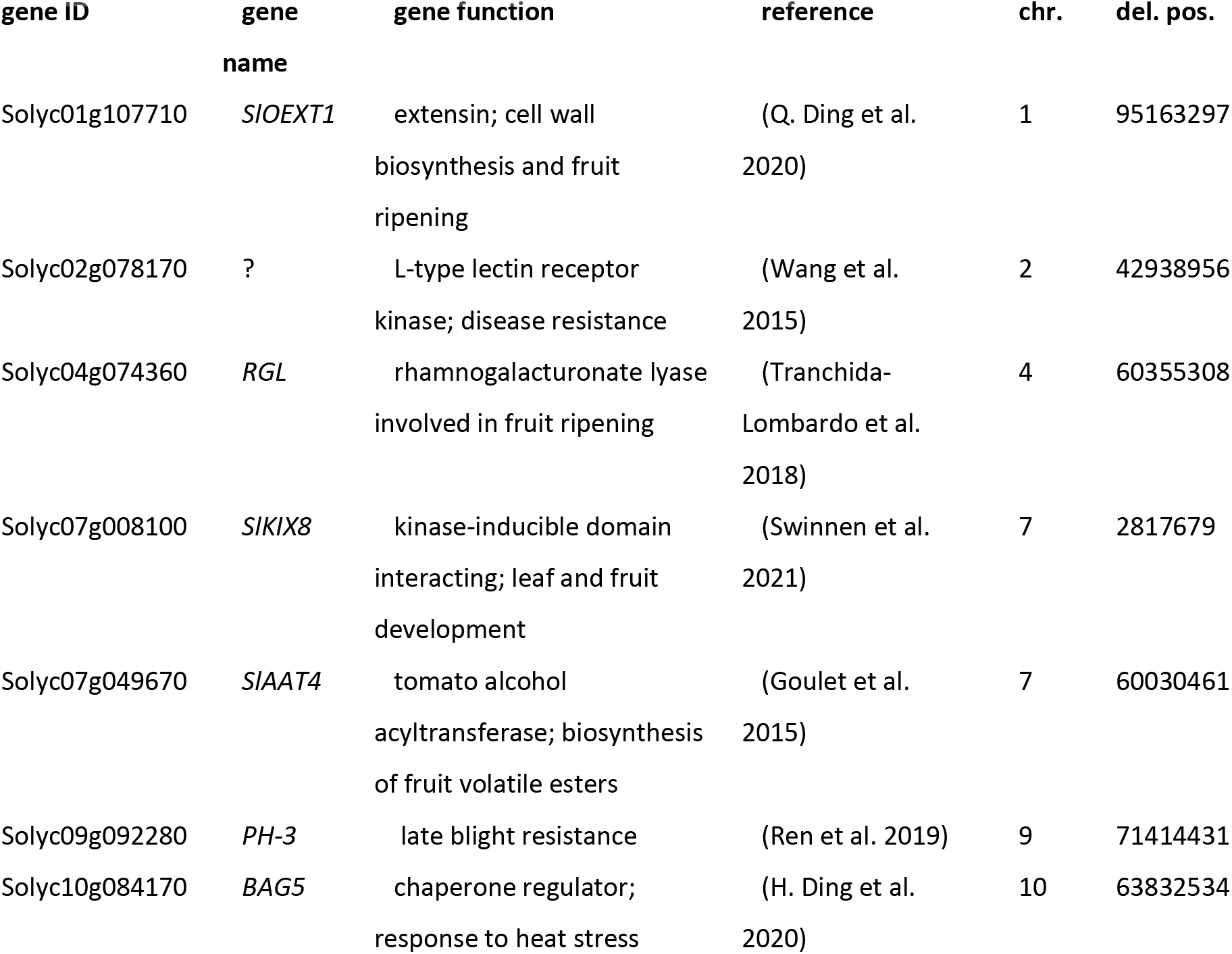
Examples of tomato genes with common deleterious genotypes (frequency > 50%) in SLL Modern. Abbreviations are described as follows: chr., chromosome; del. pos., position of the site with a deleterious allele

Considering the importance of fruit flavor genes for crop improvement, we further investigated the impact of deleterious mutations on fruit volatiles via genome-wide association studies (GWAS) on several volatiles involved in determining fruit flavor (supplementary table 5). We discovered a deleterious allele significantly correlated with methyl salicylate content (Fig. 5a). A deleterious allele on chromosome 1, position 85496207 (fig 4a) is within Solyc01g092950 (STM3), which codes for a MADS-box transcription factor. This deleterious allele was found only in SLL Americas and is negatively correlated with methyl salicylate levels (Fig. 5c and Fig. 5e).

**Fig. 5.**
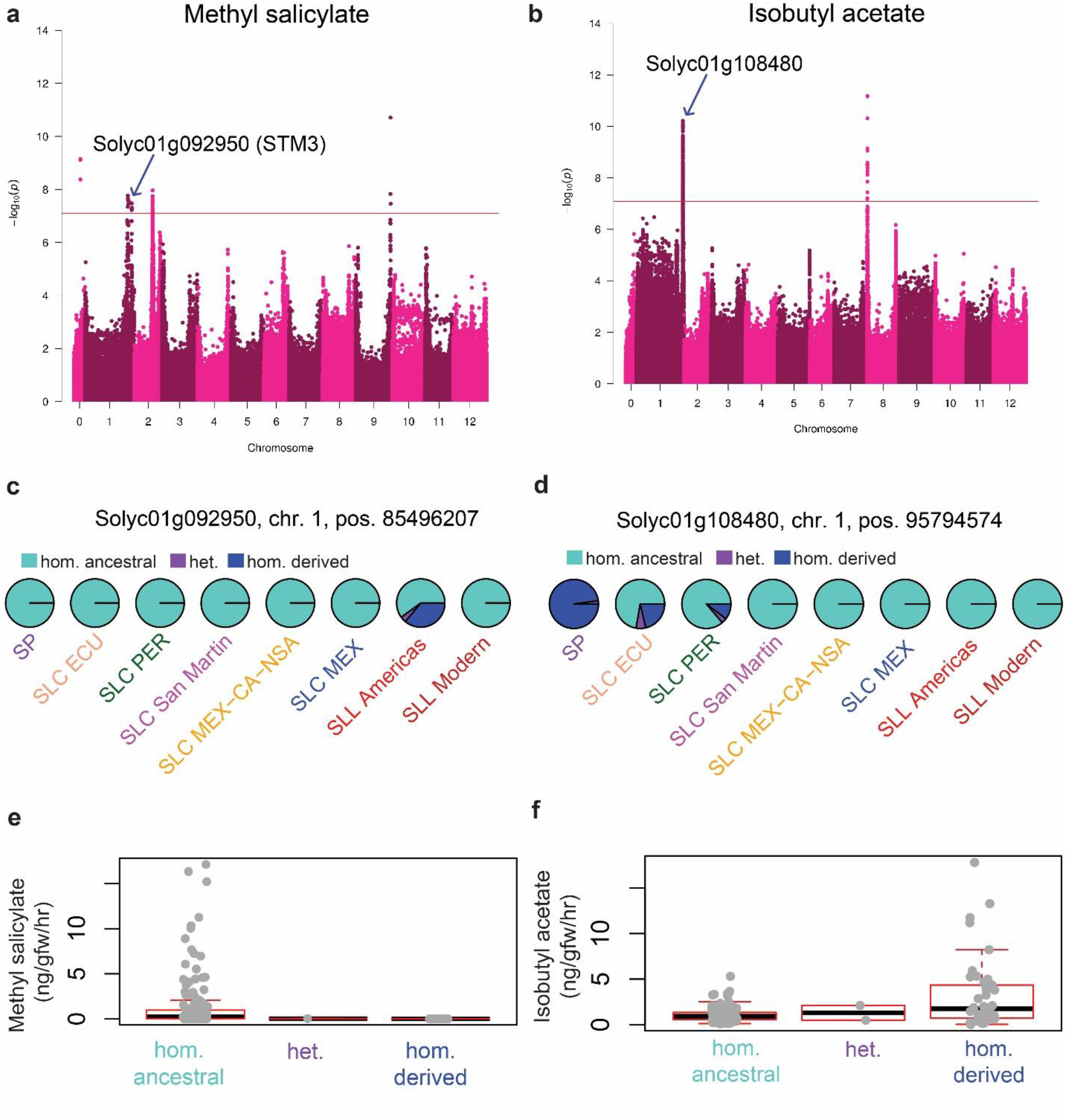
GWAS results on deleterious alleles associated with genes involved in tomato fruit volatile biosynthesis. Results are provided for methyl salicylate (a, c, and e) and isobutyl acetate (b, d, and f). For each volatile, Manhattan plots (a-b), population-level comparisons of genotype frequencies (c-d), and their phenotypic distribution in different genotypes (e-f) are presented.

We also discovered a deleterious allele correlated with isobutyl acetate levels (Fig. 5b). This deleterious allele on chromosome 1, position 95794574, (Fig. 5b) is within Solyc01g108480, which codes for a serine carboxypeptidase. This deleterious allele is common in SP and seems to have diminished in frequency in SLC ECU and SLC PER, then completely lost in remaining SLC populations before the divergence of SLL and SLC (Fig. 5d). Additionally, that allele has a positive correlation with isobutyl acetate (Fig. 5f).

## Discussion

Through this study, we have attempted to create a nuanced view of evolution of deleterious mutations through tomato domestication history. To achieve this goal, we have taken a multipronged approach which involves studying deleterious mutations in the context of demographic and genomic changes.

### Evolution of deleterious mutations through tomato domestication history

We observed an overall reduction in genetic diversity during tomato domestication history (fig. 2c), which is consistent with previous hypotheses on bottleneck events due to artificial selection (Lin et al. 2014; Blanca et al. 2015; Razifard et al. 2020). Also, we observed changes in allele frequency spectrum when comparing cultivated tomato populations with their closely related wild and semi-wild populations (fig. 1c). Several factors might be responsible for such a shift in allele frequency spectrum. One such factor is the transition from outcrossing-inbreeding mating system in SP to a mostly inbreeding mating system in SLC and SLL. However, prevalence of rare alleles in a population can be impacted also by extensive selective sweeps (Caicedo et al. 2007), gene flow (Städler et al. 2005), and crossing of modern tomato varieties with wild varieties for improving favorable traits, such as disease resistance (Hajjar and Hodgkin 2007).

Population expansion, particularly following bottlenecks (Nakazato and Housworth 2011),can also lead to accumulation of new mutations, which seems to have impacted SLL and some SLC populations. Notably, however, SLC ECU and SLC MEX-CA-NSA have median Tajima’s *D* values near zero (similar to SP), which implies an overall mutation-drift equilibrium in these populations.

Our estimates of *Tajma’s D* implied abundance of rare alleles in modern tomato accessions collected from outside South America and Mesoamerica (Fig. 1c). Loss of genetic diversity due to bottleneck could partially explain this pattern. Another important fact might be the intentional crossing of modern tomato varieties with wild tomato accessions to enhance agricually favorable traits such as disease resistance in the modern accessions (Hajjar and Hodgkin 2007).

We also found a correspondence between the average number of deleterious mutations and nucleotide diversity in each tomato population. Overall, we observed a downward trend, throughout tomato domestication history, in both nucleotide diversity and average number of deleterious mutations (Fig. 1c–1d). This observation suggests that bottleneck events combined with transition to inbreeding have had a major impact on the absolute number of deleterious mutations during tomato domestication history. Such demographic factors can remove many mutations from a population, regardless of their impact on fitness. Thus, we advocate for the use of ratios such as *nonsyn./syn.*, relative mutational load and deleterious allele frequency along with absolute number of deleterious mutations to provide more accurate comparisons of mutational load.

Our results imply that SP harbors a relatively low proportion of nonsynonymous and deleterious alleles compared to other populations (Figs. 2a–2b), although SP accessions have the highest numbers of sites with derived deleterious alleles per genome (Fig. 1d). Also, the cultivated tomato seems to harbor more relative mutational load compared to SP, confirming the estimates by (Koenig et al. 2013); although the median per-genome number of sites with derived deleterious alleles is significantly smaller than SP. Our results also suggest that proportion of nonsynonymous alleles have increased during the origin of SLC in Ecuador as well as during the northward spread of SLC, compared to Ecuadorian SLC. However, no significant change seems to have happened during the divergence of SLL and SLC MEX. However, modern tomato accessions (“SLL modern”) seem to harbor a significantly lower proportion of nonsynonymous mutations than SLC MEX.

Relative mutational load also seems to have increased during the origin of SLC (Figs. 2a–2b). Also, SLC San Martin seems to have a significantly higher mutational load than other South American SLC groups. Compared to SLC ECU, SLC MEX also contains a significantly higher relative mutational load. However, no significant difference was observed in relative mutational load in modern SLL, compared with SLC MEX.

These observations altogether suggest that a) drastic changes in proportions of nonsynonymous and deleterious have occurred, not during the final stage of tomato domestication, but rather with the origin of its semi-domesticated relative, SLC; b) many neutral or beneficial nonsynonymous mutations might have been selected for during the northward spread of SLC, represented by SLC MEX-CA-NSA; c) during modern breeding of the cultivated tomato, many neutral or beneficial nonsynonymous mutations have been purged out of modern SLL, although the proportion of its deleterious alleles have remained, more or less, the same. Therefore, in modern cultivated tomato, the cost-of-domestication hypothesis (Lu et al. 2006) is, in part, manifested in loss of non-deleterious alleles.

The genetic cost of tomato domestication was also evident among private alleles. In SLC and SLL, we found a higher proportion of private deleterious alleles than private neutral nonsynonymous alleles (supplementary fig. 2). This observation implies that many deleterious alleles in tomato have a recent origin, thus are less likely to be shared with other populations via gene flow. We also observed as higher than expected average frequency of private deleterious alleles in SLC and SLL populations (supplementary fig. 3). Therefore, the origin of SLC and subsequent tomato domestication seems to have been less effective at purging out deleterious alleles that are unique to each population. This is consistent with the theory that in small populations, selection is less effective in purging deleterious alleles.

Exploring genomic factors affecting the cost of tomato domestication, we found a correlation between mutational load and three genomic factors (Table 1). Overall, a higher mutational load was observed in genomic regions with high average nucleotide diversity, low recombination, and weaker selective sweep signals. Several factors might be involved in creating this pattern. One possible factor is genetic hitchhiking of deleterious alleles on beneficial alleles in areas under “soft” selective sweeps, where a considerable level of nucleotide diversity is still present in a population (McCoy and Akey 2017). In contrast to soft sweeps, “hard” selective sweeps lead to loss of nucleotide diversity, impacting alleles from all fitness categories including deleterious alleles, in the genomic regions under strong or ancient selective sweeps.

Exceptions to the general patterns of association between mutational load and genomic factors was observed in SLC San Martin and the two SLL populations. This is interesting considering large average fruit sizes (Fig. 1a) as well as low genetic diversity (Fig. 1c) and higher frequency of deleterious alleles than expected (Fig. 2c) in these populations. These observations imply that SLC San Martin may have undergone a demographic history comparable with that of the cultivated tomato groups, which has led to decoupling of deleterious allele accumulation and recombination rate.

### The evolution of deleterious load in the context of genomic structural variants

Our results revealed a reduction in the number of SVs through tomato domestication history (fig. 3a), which is similar to the pattern observed for the number of mutations from different categories, i.e. non-coding, synonymous, non-synonymous, and neutral non-synonymous (fig. 1d and supplementary fig. 1). Such reduction can also be attributed to loss of overall genetic diversity (fig. 1c), due to bottleneck events, through tomato domestication history (Lin et al. 2014; Blanca et al. 2015; Razifard et al. 2020).

Furthermore, for most accessions, we discovered a significantly larger number of SVs in non-genic regions of the tomato genome (supplementary fig. 4). This is consistent with the theory that SVs occurring in genic regions might have a strongly negative impact on the fitness of the individuals, and thus will be removed from the population by selection (Vaughn et al. 2021). Among SVs in genic regions, most were found in introns and untranslated regions (supplementary fig. 5). Together, these results suggest strong selection against SVs in genic regions, especially those impacting, or likely disrupting, exons.

Finally, we discovered a larger relative mutational load (*del.*/*neutr. nonsyn.*) within SVs compared to genomic regions not impacted by SVs (Fig. 3b). This observation implies that SVs might lead to gene disruption and pseudogenization, which creates genomic hotspots for accumulation of deleterious mutations. Alternatively, it is possible that previously pseduogenized genes harbor a larger number of SVs and deleterious mutations, due to relaxed selection.

### Functional link between deleterious mutations and desirable traits for tomato breeding

Most methods for detecting deleterious mutations estimate the impact of a mutation based on the degree of the phylogenetic conservation of the gene in which that mutation occurs. This theory is valid for species living in similar environments, but might be difficult to justify for closely related organisms that live in drastically different environments (Tellier et al. 2011). This is often the case for domesticated organisms, which live in a broad range of habitats compared to their wild relatives. Tracking the evolutionary trajectory of deleterious alleles in tomato and its closely related populations, we found that many deleterious alleles have been purged from cultivated tomato, confirming the results discussed in previous sections. Also, our GO-term analyses on genes with deleterious alleles present in modern SLL revealed that genes with a broad spectrum of functions have been impacted by deleterious alleles. However, some genes, for example those involved in ascorbic acid (vitamin C) pathway or pollen germination under heat stress, were overrepresented among the GO terms. Ascorbic acid has an important role in neutralization of reactive oxygen species (Mellidou et al. 2012), thus these GO terms imply adaptations to a diverse range of habitats in the cultivated tomato.

Considering the importance of knowledge on deleterious alleles for crop improvement, we created TomDel, a public database of deleterious alleles present in cultivated tomato and its closely related wild and semi-wild populations. This database provides information on the probable population of origin as well as genotype frequencies of deleterious alleles present in modern SLL and its closely related populations. Thus, it will serve as a guide for breeding purposes with the aim of avoiding unintentional introduction of deleterious mutations to the tomato varieties of interest during experimental crossing.

Among well-studied tomato genes, we found deleterious alleles with high frequency (> 50%) in genes involved in response to biotic and abiotic stress as well as fruit development and flavor regulations (Fig. 4, Table 2). Notably, we found a deleterious allele in *SlAAT4*, a known gene involved in biosynthesis of fruit volatiles, which are essential for fruit flavor. This motivated us to conduct GWAS on several volatiles to search for deleterious alleles that might be associated with fruit volatile biosynthesis.

Our GWAS results (Fig. 5a–b) revealed deleterious alleles associated with methyl salicylate and isobutyl acetate concentrations in tomato fruits. In previous studies, no significant correlation has been found between methyl salicylate and flavor preference (Tieman et al. 2017). However, methyl salicylate is a well-studied compound involved in disease resistance in plants, e.g. see (Min et al. 2018), as well as regulating flowering time (Banday and Nandi 2015).

The deleterious allele associated with methyl salicylate is within STM3, which codes for a transcription factor, putatively involved in regulating flowering time (Jiang et al. unpublished data) and inflorescence branching (Alonge et al. 2020) in tomato. This deleterious allele is negatively correlated with methyl salicylate concentrations and it is only found in SLL Americas. Therefore, it is likely that this deleterious allele has arisen during the origin of SLL, but was selected against during the modern breeding of the cultivated tomato.

Our results also revealed a deleterious allele associated with isobutyl acetate concentrations in tomato fruits (Fig. 5b). Isobutyl acetate was previously reported to be negatively correlated with flavor preference in tomato (“overall liking”; Tieman et al. 2017). This deleterious allele is within Solyc01g108480, which codes for a serine carboxypeptidase, suggested to be involved in response to wounding in tomato leaves (Moura et al. 2001). Also, this allele and seems to absent in SLC San Martin, northern populations of SLC, and both SLL populations. Thus, lower levels of isobutyl acetate probably have been selected for during tomato domestication. This finding implies loss of a deleterious allele that might be beneficial for crop improvement. Therefore, at least in this case, it is crucial to distinguish between truly deleterious alleles and those that are deleterious in the wild, but desirable for breeding purposes.

## Conclusions

In this study, we created novel knowledge on the evolution of deleterious mutations through tomato domestication history, especially in the context of genomic structural variants (SVs). Our results help elucidate the dynamics of deleterious mutations as well as within different stages of tomato domestication, i.e. initial domestication in Ecuador, northward spread of SLC toward Central America and Mexico, and redomestication o SLL in Mexico. Our results also revealed novel insight on the genomic factors, such as SVs and recombination rate, involved in shaping the deleterious load through tomato domestication history.

Through this study, we also attempted to create a functional link between mutations identified as deleterious and the phenotypes impact by such mutations. In many functional genetic studies, the causal allele underlying a phenotype is difficult to pinpoint even after identifying the causal gene. Our approach helps narrow down the list of candidate alleles by identifying those with a potentially high impact on the phenotype of interest. Our findings can also be used for guiding breeding experiments. Using our results, breeders can avoid accidentally introducing deleterious alleles into the tomato variety of interest from its closely related populations.

As our results revealed for isobutyl acetate, some mutations identified as deleterious by currently available methods, might in fact confer a human-desired trait in crops. Therefore, phrases such as “deleterious mutations” and “mutational load” might not fully represent the diverse functional impacts of mutations with high impacts on underlying phenotypes. Therefore, our uses of such phrases throughout this paper, refer to high-impact mutations identified as deleterious by two commonly used algorithms. We acknowledge that some of the “deleterious” mutations reported in this study, might confer desirable traits in domesticated organisms.

## Acknowledgments

We thank members of Buckler Lab at Cornell University, especially Edward Buckler, Baoxing Song, and Michelle Stitzer for their valuable feedback on SV analyses. This study was funded by the National Science Foundation of USA, grant number: IOS 1564366

## Materials and Methods

### Plant material and input dataset

A dataset of genome-wide variants created in a previous study (Razifard et al. 2020) was used to identify deleterious mutations in wild and cultivated tomatoes. The input dataset included accessions of diverse SP (*Solanum pimpinellifolium*) and SLC (*S. lycopersicum* var*. cerasiforme*) from South America, SLL (*S. lycopersicum* var*. lycopersicum*) from the Americas as well as improved material from outside of the Americas, and one accession of *Solanum pennellii* L. (EA00585), serving as outgroup. The reference genome used for calling variants was ‘Heinz 1706’, build SL2.50 (Sato et al. 2012). From the input dataset, we excluded accessions with > 25% missing data as well as sites with insertion-deletions (indels), leaving 15,011,193 SNPs in the final dataset.

The missing genotypes in the final dataset were imputed using LinkImpute version 1.1.4 (reference to paper), with the default settings (“LD-kNNi”, imputation of missing genotypes based on k-nearest neighbors as well as linkage disequilibrium). The outgroup accession was used for determining the ancestral alleles for all SNPs in the final dataset, i.e. considering the allele observed in the outgroup as ancestral and the allele that differed from the outgroup as derived.

### Group Delimitations

The main population were defined in a mostly consistent manner with the phylogenetic relationships provided in Razifard et al. (2020), with the following exceptions. Due to low number of individuals for some populations of SP, we combined all three SP populations -- a population of SP from northern Ecuador (SP NECU), one from southern Ecuador (SP SECU) and one from Peru (SP PER) -- into a single group. We also split the cultivated tomato group (SLL) into two major populations: a clade of SLL accessions from the Americas (“SLL Americas”) and another clade comprising mostly modern SLL accessions from outside the Americas (“SLL modern”), to better distinguish the effects of tomato domestication from later improvement

### Population-level analyses

Nucleotide diversity (*π*) and Tajima’s *D* were calculated for each population using VCFTools within 10-kb windows. A Maximum-likelihood population tree was constructed using TreeMix (Pickrell and Pritchard 2012). The input for TreeMix was the same as the one used for SIFT analysis. To account for the effect of linkage disequilibrium, stretches of 1,200 SNPs were grouped in separate windows (“−k 1200”), representing ~10-Mb windows (Pickrell and Pritchard 2012).

### Identification of SNP categories and evaluation of mutational impacts

Two methods were used to classify SNPs into different categories and predict the impact of derived alleles. First, all sites were evaluated using SIFT (Kumar et al. 2009), which categorized SNPs as non-coding, synonymous, and nonsynonymous; the effect of nonsynonymous SNPs was further predicted as “deleterious” or “deleterious (low-confidence)”. Second, following the procedure of Renaut and Rieseberg (2015), SNPs identified as nonsynonymous by SIFT were further evaluated using PROVEAN (Choi et al. 2012). SNPs with PROVEAN score < −2.5 were considered potentially deleterious, as suggested by Choi et al. 2012. In all downstream analyses, only SNPs identified as deleterious by both SIFT and PROVEAN were considered to be deleterious. Nonsynonymous SNPs not classified as deleterious were considered to be neutral.

To avoid complications due to reference bias, we conducted SIFT runs by substituting reference and alternate alleles with inferred ancestral and derived alleles, respectively. Also, we ran PROVEAN using predicted outgroup protein sequences as input (by modifying reference protein sequences to incorporate amino acids based on the alleles observed in the outgroup).

### Identification of genomic structural variants (SVs)

Genomic structural variants (SVs) were identified for all accessions included in this study, using three independent pipelines, Lumpy (v. 0.3.0; Layer et al., 2014), Delly (v. 0.8.7; Rausch et al., 2012), and Manta (v. 1.6.0; Chen et al., 2016), using default options. Among different SV types, insertions are often difficult to reliably detect using the currently available pipelines, therefore we excluded insertions (only 9 in total) from our analyses. The output from all three pipelines were merged using SURVIVOR (v.1.0.7; Jeffares et al., 2017), keeping SVs present in the outputs from at least two out of three pipelines, with minimum size of 30 bp long. SVs in different accessions but on the same chromosome, with the same SV type, within 100 bp from one another, and with similar length (90% overlap) were considered as the same SV. To avoid false-positive identification due to missing data, rare SVs with minimum allele frequency (MAF) < 0.01 were also excluded from downstream analyses. All SVs were polarized by considering the genotype observed in the outgroup, *S. pennellii*, as the putative ancestral state.

A custom R script (available from Github repository) was developed to count the number of deleterious and neutral nonsynonymous mutations that overlap with different SV types in each accession. Another R script was developed for counting the number of SVs occurring in genic regions.

### TomDel

We created a visual database of all deleterious alleles identified in this study within modern tomato and its closely related populations. To create this database, we used a custom R script (available from GitHub repository) to plot genotype frequencies of all the SNPs with deleterious alleles in all the populations. The database is publicly available on GitHub (https://github.com/hrazif/TomDel-0.1).

### Private alleles

We created a custom R script (available from GitHub repository) to prepare separate lists of derived deleterious mutations and the derived neutral nonsynonymous mutations that were unique to each of the populations. Derived alleles with frequency > 0 in only one population were considered as private to that population. Counting the number of private alleles in each accession, as well as the follow-up plotting and statistical comparisons, were conducted in R (v. 3.6.3; http://www.R-project.org/).

### Statistical comparisons

All statistical comparisons were conducted in R. Kruskal-Wallis tests were used to determine whether populations have a significant effect on the distribution of the ratios (e.g. del/neutr). Then, Dunn’s test was used to perform a pair-wise comparison between the distributions of values between populations. The comparison between private and shared del/neutr was based on a T-test. A Kolmogorov-Smirnov test was conducted to compare the distribution of derived allele frequencies of deleterious and neutral nonsynonymous alleles. In all statistical tests involving multiple testing herein, the p-value significance cutoff was decided at 0.05, after adjusting for multiple testing based on Bonferroni correction.

### Correlation between relative mutational load and other genomic features

We explored potential correlation between del/neutr and three genomic statistics: *π*, recombination rate (based on the genetic map ITAG2.3 from www.solgenomics.net), and SweeD Likelihood ratio (testing for deviation of site frequency spectrum from neutrality; Pavlidis et al. 2013). Each statistic was calculated within 100-kb windows for each population. Correlation between the four statistics and del/neutr was explored separately for each statistic as follows. Windows were sorted by each statistic and divided into “high” and “low” groups, placing windows with top 50% values in the “high” group and vice versa. Wilcoxon tests were conducted separately for each statistic to test for difference in the distributions of relative mutational load across individuals in “high” and “low” groups in each population.

### GO-term enrichment analyses

We conducted GO-term enrichment analyses, using R package topGO (v.2.28.0) (Alexa et al. 2006), on SNPs with deleterious alleles present in modern cultivated tomato (deleterious derived allele frequency > 0). Multiple algorithms -- “classic”, “elim”, “weight”, “weight01”, “lea”, and “parentchild” -- were tested for these analyses, and their results were explored for shared GO terms. Statistical significance was measured using Fisher’s Exact Test and the resulting p-values were independently adjusted for multiple testing, according to the way in which each method treats GO-term relationships (further described in topGO user manual).

### Genome-wide Association Studies (GWAS)

Several tomato fruit volatiles were quantified as described in (Tieman et al. 2017). Briefly, three plants from each accession were grown in a randomize manner in Florida. Volatiles were extracted from ripe fruits and were quantified using gas chromatography combined with mass spectrometry.

The genomic variant dataset used for GWAS included small indels and “chromosome-zero” SNPs, but excluded SNPs with minor allele frequency < 0.05 and missing rate > 10%. Also, to avoid potential false GWAS hits due to strong population differentiation, SP accessions were removed from the GWAS input dataset. The GWAS results were explored for overlap with deleterious mutations. Phenotypes with distributions deviating significantly from normality based on Shapiro test (p-value < 0.01) were normalized in R. GWAS was conducted in GEMMA (v 0.94.1) using a mixed linear model (Zhou and Stephens 2012). The associations were adjusted for population structure according to a genetic relatedness matrix created also using GEMMA. P-values based on Likelihood Ratio Test (LRT) were used for determining significant correlations. A significance cutoff was determined based on the effective number of independent SNPs calculated using the Genetic Type I error calculator (GEC, v0.2) (Li et al. 2012).

## References

Alexa A, Rahnenführer J, Lengauer T. 2006. Improved scoring of functional groups from gene expression data by decorrelating GO graph structure. Bioinformatics 22:1600–1607.

Alonge M, Wang X, Benoit M, Soyk S, Pereira L, Zhang L, Suresh H, Ramakrishnan S, Maumus F, Ciren D, et al. 2020. Major Impacts of Widespread Structural Variation on Gene Expression and Crop Improvement in Tomato. Cell 182:145–161.e23. Available from: https://www.sciencedirect.com/science/article/pii/S0092867420306164

Banday ZZ, Nandi AK. 2015. Interconnection between flowering time control and activation of systemic acquired resistance. Front. Plant Sci. 6:174. Available from: https://www.frontiersin.org/article/10.3389/fpls.2015.00174

Barraclough TG, Fontaneto D, Ricci C, Herniou EA. 2007. Evidence for Inefficient Selection Against Deleterious Mutations in Cytochrome Oxidase I of Asexual Bdelloid Rotifers. Mol. Biol. Evol. 24:1952–1962. Available from: https://doi.org/10.1093/molbev/msm123

Barton NH. 1995. Linkage and the limits to natural selection. Genetics 140:821 LP – 841. Available from: http://www.genetics.org/content/140/2/821.abstract

Blanca J, Montero-Pau J, Sauvage C, Bauchet G, Illa E, Díez MJ, Francis D, Causse M, van der Knaap E, Cañizares J. 2015. Genomic variation in tomato, from wild ancestors to contemporary breeding accessions. BMC Genomics 16:257. Available from: http://www.biomedcentral.com/1471-2164/16/257

Bortoluzzi C, Bosse M, Derks MFL, Crooijmans RPMA, Groenen MAM, Megens H-J. 2020. The type of bottleneck matters: Insights into the deleterious variation landscape of small managed populations. Evol. Appl. 13:330–341. Available from: https://doi.org/10.1111/eva.12872

Byers DL, Waller DM. 1999. Do Plant Populations Purge Their Genetic Load? Effects of Population Size and Mating History on Inbreeding Depression. Annu. Rev. Ecol. Syst. 30:479–513. Available from: https://doi.org/10.1146/annurev.ecolsys.30.1.479

Caicedo AL, Williamson SH, Hernandez RD, Boyko A, Fledel-Alon A, York TL, Polato NR, Olsen KM, Nielsen R, McCouch SR, et al. 2007. Genome-Wide Patterns of Nucleotide Polymorphism in Domesticated Rice. PLOS Genet. 3:e163. Available from: https://doi.org/10.1371/journal.pgen.0030163

Calafiore R, Ruggieri V, Raiola A, Rigano MM, Sacco A, Hassan MI, Frusciante L, Barone A. 2016. Exploiting Genomics Resources to Identify Candidate Genes Underlying Antioxidants Content in Tomato Fruit. Front. Plant Sci. 7:397. Available from: https://www.frontiersin.org/article/10.3389/fpls.2016.00397

Choi Y, Sims GE, Murphy S, Miller JR, Chan AP. 2012. Predicting the functional effect of amino acid substitutions and indels. PLoS One 7:e46688–e46688. Available from: https://pubmed.ncbi.nlm.nih.gov/23056405

Ding Q, Yang X, Pi Y, Li Z, Xue J, Chen H, Li Y, Wu H. 2020. Genome-wide identification and expression analysis of extensin genes in tomato. Genomics 112:4348–4360. Available from: http://www.sciencedirect.com/science/article/pii/S0888754320300914

Elena SF, Lenski RE. 1997. Test of synergistic interactions among deleterious mutations in bacteria. Nature 390:395–398. Available from: https://doi.org/10.1038/37108

Gaut BS, Seymour DK, Liu Q, Zhou Y. 2018. Demography and its effects on genomic variation in crop domestication. Nat. Plants 4:512–520. Available from: http://dx.doi.org/10.1038/s41477-018-0210-1

Goulet C, Kamiyoshihara Y, Lam NB, Richard T, Taylor MG, Tieman DM, Klee HJ. 2015. Divergence in the enzymatic activities of a tomato and *Solanum pennellii* alcohol acyltransferase impacts fruit volatile ester composition. Mol. Plant 8:153–162.

Grossen C, Guillaume F, Keller LF, Croll D. 2020. Purging of highly deleterious mutations through severe bottlenecks in Alpine ibex. Nat. Commun. 11:1001. Available from: https://doi.org/10.1038/s41467-020-14803-1

Hajjar R, Hodgkin T. 2007. The use of wild relatives in crop improvement: A survey of developments over the last 20 years. Euphytica 156:1–13.

Kim M-S, Lozano R, Kim JH, Bae DN, Kim S-T, Park J-H, Choi MS, Kim J, Ok H-C, Park S-K, et al. 2021. The patterns of deleterious mutations during the domestication of soybean. Nat. Commun. 12:97. Available from: https://doi.org/10.1038/s41467-020-20337-3

Klopfstein S, Currat M, Excoffier L. 2006. The Fate of Mutations Surfing on the Wave of a Range Expansion. Mol. Biol. Evol. 23:482–490. Available from: https://doi.org/10.1093/molbev/msj057

Koenig D, Jimenez-Gomez JM, Kimura S, Fulop D, Chitwood DH, Headland LR, Kumar R, Covington MF, Devisetty UK, Tat A V., et al. 2013. Comparative transcriptomics reveals patterns of selection in domesticated and wild tomato. Proc. Natl. Acad. Sci. 110:E2655–E2662. Available from: http://www.pnas.org/cgi/doi/10.1073/pnas.1309606110

Kono TJY, Fu F, Mohammadi M, Hoffman PJ, Liu C, Stupar RM, Smith KP, Tiffin P, Fay JC, Morrell PL. 2016. The Role of Deleterious Substitutions in Crop Genomes. Mol. Biol. Evol. 33:2307–2317. Available from: http://dx.doi.org/10.1093/molbev/msw102

Kumar P, Henikoff S, Ng PC. 2009. Predicting the effects of coding non-synonymous variants on protein function using the SIFT algorithm. Nat. Protoc. 4:1073–1082.

Li MX, Yeung JMY, Cherny SS, Sham PC. 2012. Evaluating the effective numbers of independent tests and significant *p*-value thresholds in commercial genotyping arrays and public imputation reference datasets. Hum. Genet. 131:747–756.

Lin T, Zhu G, Zhang J, Xu X, Yu Q, Zheng Z, Zhang Z, Lun Y, Li S, Wang X, et al. 2014. Genomic analyses provide insights into the history of tomato breeding. Nat. Genet. 46:1220–1226. Available from: http://www.nature.com/doifinder/10.1038/ng.3117

Liu Q, Zhou Y, Morrell PL, Gaut BS, Ge S. 2017. Deleterious variants in Asian Rice and the potential cost of domestication. Mol. Biol. Evol. 34:908–924.

Lozano R, Gazave E, dos Santos JPR, Stetter MG, Valluru R, Bandillo N, Fernandes SB, Brown PJ, Shakoor N, Mockler TC, et al. 2021. Comparative evolutionary genetics of deleterious load in sorghum and maize. Nat. Plants 7:17–24. Available from: https://doi.org/10.1038/s41477-020-00834-5

Lu J, Tang T, Tang H, Huang J, Shi S, Wu C-I. 2006. The accumulation of deleterious mutations in rice genomes: a hypothesis on the cost of domestication. Trends Genet. 22:126–131. Available from: https://www.sciencedirect.com/science/article/pii/S0168952506000229

Makino T, Rubin C-J, Carneiro M, Axelsson E, Andersson L, Webster MT. 2018. Elevated Proportions of Deleterious Genetic Variation in Domestic Animals and Plants. Genome Biol. Evol. 10:276–290.

Marsden CD, Ortega-Del Vecchyo D, O’Brien DP, Taylor JF, Ramirez O, Vilà C, Marques-Bonet T, Schnabel RD, Wayne RK, Lohmueller KE. 2016. Bottlenecks and selective sweeps during domestication have increased deleterious genetic variation in dogs. Proc. Natl. Acad. Sci. 113:152 LP – 157. Available from: http://www.pnas.org/content/113/1/152.abstract

McCoy RC, Akey JM. 2017. Selection plays the hand it was dealt: evidence that human adaptation commonly targets standing genetic variation. Genome Biol. 18:139. Available from: https://doi.org/10.1186/s13059-017-1280-5

Mellidou I, Keulemans J, Kanellis AK, Davey MW. 2012. Regulation of fruit ascorbic acid concentrations during ripening in high and low vitamin C tomato cultivars. BMC Plant Biol. 12:239. Available from: https://doi.org/10.1186/1471-2229-12-239

Min D, Li F, Zhang X, Shu P, Cui X, Dong L, Ren C, Meng D, Li J. 2018. Effect of methyl salicylate in combination with 1-methylcyclopropene on postharvest quality and decay caused by Botrytis cinerea in tomato fruit. J. Sci. Food Agric. 98:3815–3822. Available from: https://doi.org/10.1002/jsfa.8895

Moura DS, Bergey DR, Ryan CA. 2001. Characterization and localization of a wound-inducible type I serine-carboxypeptidase from leaves of tomato plants (Lycopersicon esculentum Mill.). Planta 212:222–230.

Moyers BT, Morrell PL, McKay JK. 2017. Genetic Costs of Domestication and Improvement. J. Hered. 109:103–116. Available from: http://academic.oup.com/jhered/article/doi/10.1093/jhered/esx069/4064635/Genetic-Costs-of-Domestication-and-Improvement

Muller HJ. 1932. Some genetic aspects of sex. Am. Nat. 66:118–138.

Nakazato T, Housworth EA. 2011. Spatial genetics of wild tomato species reveals roles of the Andean geography on demographic history. Am. J. Bot. 98:88–98.

Ng PC, Henikoff S. 2003. SIFT: predicting amino acid changes that affect protein function. Nucleic Acids Res. 31:3812–3814. Available from: https://doi.org/10.1093/nar/gkg509

Pavlidis P, Živković D, Stamatakis A, Alachiotis N. 2013. SweeD: Likelihood-based detection of selective sweeps in thousands of genomes. Mol. Biol. Evol. 30:2224–2234.

Pickrell JK, Pritchard JK. 2012. Inference of population splits and mixtures from genome-wide allele frequency data. PLoS Genet. 8:e1002967.

Ramu P, Esuma W, Kawuki R, Rabbi IY, Egesi C, Bredeson J V, Bart RS, Verma J, Buckler ES, Lu F. 2017. Cassava haplotype map highlights fixation of deleterious mutations during clonal propagation. Nat. Genet. 49:959–963. Available from: https://doi.org/10.1038/ng.3845

Razifard H, Ramos A, Della Valle AL, Bodary C, Goetz E, Manser EJ, Li X, Zhang L, Visa S, Tieman D, et al. 2020. Genomic Evidence for Complex Domestication History of the Cultivated Tomato in Latin America. Mol. Biol. Evol. 37:1118–1132.

Renaut S, Rieseberg LH. 2015. The accumulation of deleterious mutations as a consequence of domestication and improvement in sunflower and other Compositae crops. Mol. Biol. Evol. 32:2273–2283.

Sato S, Tabata S, Hirakawa H, Asamizu E, Shirasawa K, Isobe S, Kaneko T, Nakamura Y, Shibata D, Aoki K, et al. 2012. The tomato genome sequence provides insights into fleshy fruit evolution. Nature 485:635–641. Available from: http://www.nature.com/doifinder/10.1038/nature11119

Schubert M, Jónsson H, Chang D, Der Sarkissian C, Ermini L, Ginolhac A, Albrechtsen A, Dupanloup I, Foucal A, Petersen B, et al. 2014. Prehistoric genomes reveal the genetic foundation and cost of horse domestication. Proc. Natl. Acad. Sci. 111:E5661 LP–E5669. Available from: http://www.pnas.org/content/111/52/E5661.abstract

Song J, Nada K, Tachibana S. 2002. Suppression of S-adenosylmethionine Decarboxylase Activity is a Major Cause for High-Temperature Inhibition of Pollen Germination and Tube Growth in Tomato (Lycopersicon esculentum Mill.). Plant Cell Physiol. 43:619–627. Available from: https://doi.org/10.1093/pcp/pcf078

Städler T, Roselius K, Stephan W. 2005. Genealogical footprints of speciation processes in wild tomatoes: demography and evidence for historical gene flow. Evolution (N. Y). 59:1268–1279. Available from: https://doi.org/10.1111/j.0014-3820.2005.tb01777.x

Tellier A, Fischer I, Merino C, Xia H, Camus-Kulandaivelu L, Städler T, Stephan W. 2011. Fitness effects of derived deleterious mutations in four closely related wild tomato species with spatial structure. Heredity (Edinb). 107:189–199. Available from: https://doi.org/10.1038/hdy.2010.175

Tieman D, Zhu G, Resende MFR, Lin T, Nguyen C, Bies D, Rambla JL, Beltran KSO, Taylor M, Zhang B, et al. 2017. A chemical genetic roadmap to improved tomato flavor. Science (80-.). 355:391–394.

Tranchida-Lombardo V, Aiese Cigliano R, Anzar I, Landi S, Palombieri S, Colantuono C, Bostan H, Termolino P, Aversano R, Batelli G, et al. 2018. Whole-genome re-sequencing of two Italian tomato landraces reveals sequence variations in genes associated with stress tolerance, fruit quality and long shelf-life traits. DNA Res. 25:149–160. Available from: https://doi.org/10.1093/dnares/dsx045

Travis JMJ, Münkemüller T, Burton OJ, Best A, Dytham C, Johst K. 2007. Deleterious Mutations Can Surf to High Densities on the Wave Front of an Expanding Population. Mol. Biol. Evol. 24:2334–2343. Available from: https://doi.org/10.1093/molbev/msm167

United Nations Food and Agriculture Organization. 2013. FAOSTAT. Available from: http://www.fao.org/faostat/en/#data/QC

Wang Y, Weide R, Govers F, Bouwmeester K. 2015. L-type lectin receptor kinases in Nicotiana benthamiana and tomato and their role in Phytophthora resistance. J. Exp. Bot. 66:6731–6743. Available from: https://doi.org/10.1093/jxb/erv379

Zhou X, Stephens M. 2012. Genome-wide efficient mixed-model analysis for association studies. Nat. Genet. 44:821–824. Available from: http://www.nature.com/doifinder/10.1038/ng.2310

Zhou Y, Massonnet M, Sanjak JS, Cantu D, Gaut BS. 2017. Evolutionary genomics of grape (Vitis vinifera ssp. vinifera) domestication. Proc. Natl. Acad. Sci. 114:11715 LP – 11720. Available from: http://www.pnas.org/content/114/44/11715.abstract

