## supplementary fig for "The evolutionary dynamics of genetic mutational load throughout tomato domestication history"

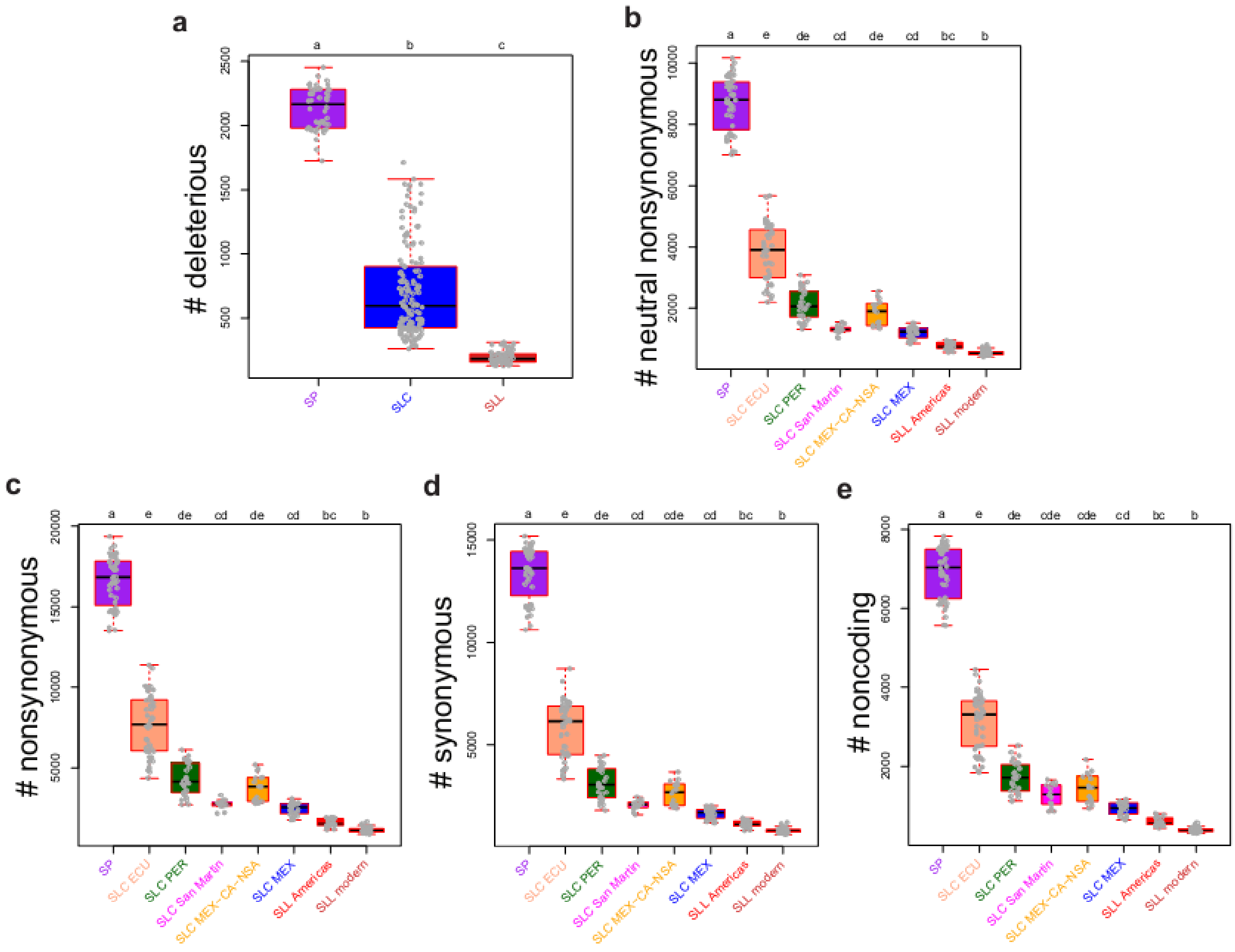


Supplementary fig. 1. The counts of sites with derived alleles in in different categories, showing a downward trend when comparing SLC populations with SP and SLLs with SLC. **(a)** a comparison of the number of sites with deleterious alleles per genome between SP, SLC, and SLL. **(b-e)** Inter-population comparisons of number of sites with alleles in different categories. Statistically non-significant comparisons, according to Dunn's test, have been indicated using lower-case numbers.

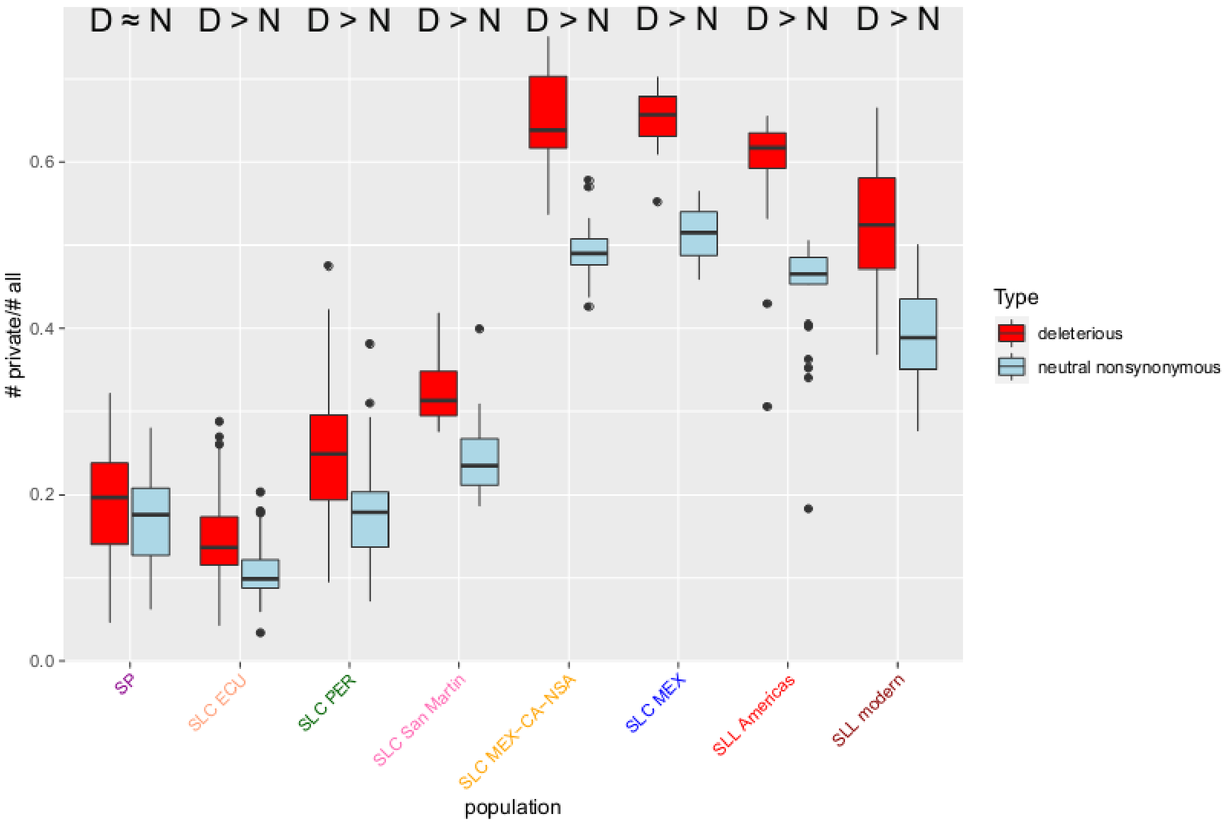


Supplementary fig. 2. Per-accession proportion of private derived deleterious to all derived deleterious (red) and private derived neutral nonsynonymous to all derived neutral nonsynonymous (blue) mutations. For each population, a significant t-test result (Bonferroni-adjusted p-value < 0.05) is shown with D > N, i.e. the ratio of private derived deleterious alleles to all derived deleterious is higher than the ratio of private derived neutral nonsynonymous alleles to all derived neutral nonsynonymous alleles. No significant different was observed for SP (D ≈ N).

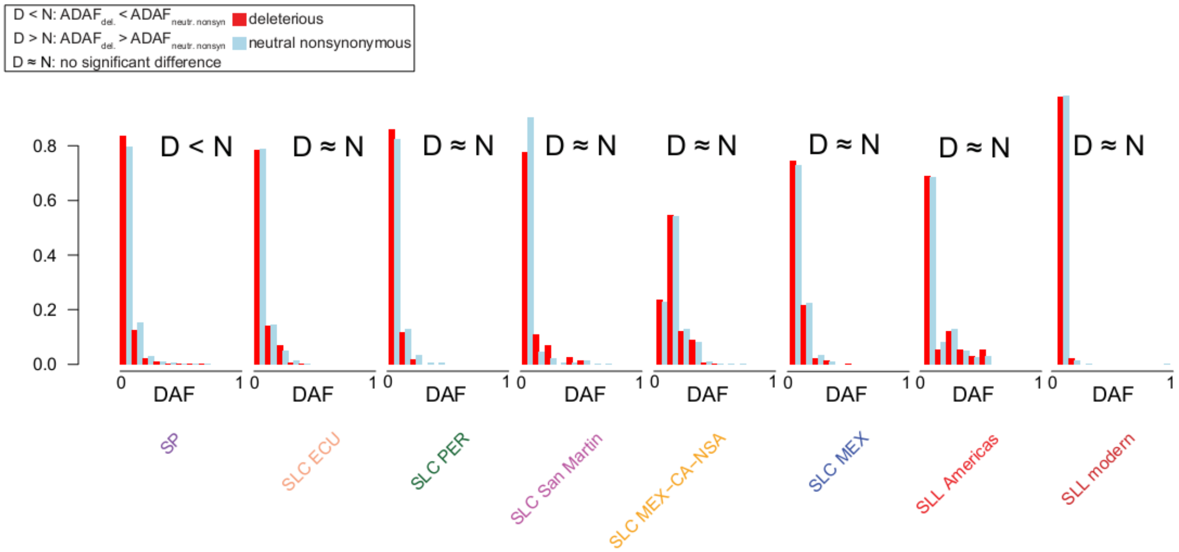


Supplementary fig. 3. Site frequency spectra for private deleterious (red) and private neutral nonsynonymous (blue) mutations in different tomato populations. Results of statistical tests of average derived allele frequency of deleterious alleles ($ADAF_{del}$) versus average derived allele frequency of neutral nonsynonymous alleles ($ADAF_{neutr.\ nonsyn.}$) are shown with D < N ($ADAF_{del}$ is smaller than $ADAF_{neutr.\ nonsyn.}$), D > N ($ADAF_{del}$ is greater than $ADAF_{neutr.\ nonsyn.}$ l), and D ≈ N ($ADAF_{del}$ is not significantly different from $ADAF_{neutr.\ nonsyn.}$).

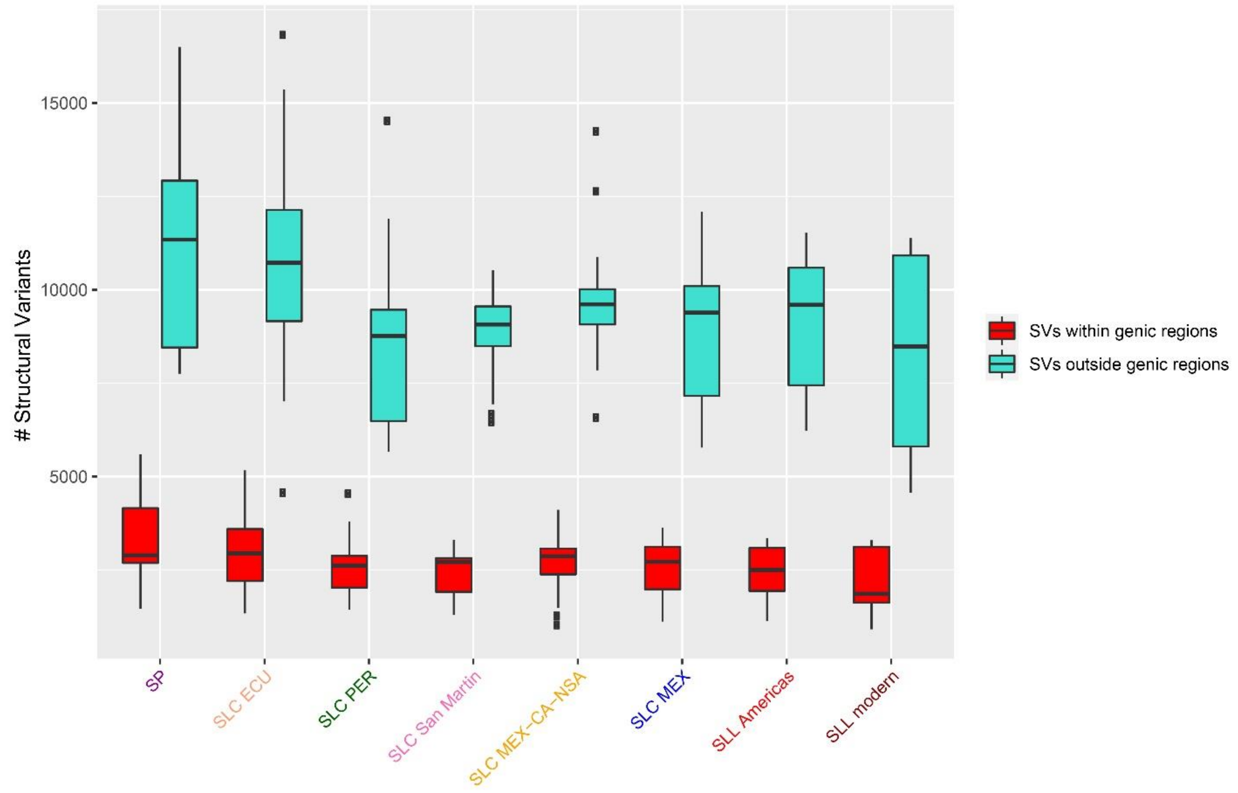


Supplementary fig. 4. A genome-wide comparison of the number of SVs within or outside genic regions in each tomato population. The comparisons within all populations are statistically significant (based a T-test; p-value < 0.001).

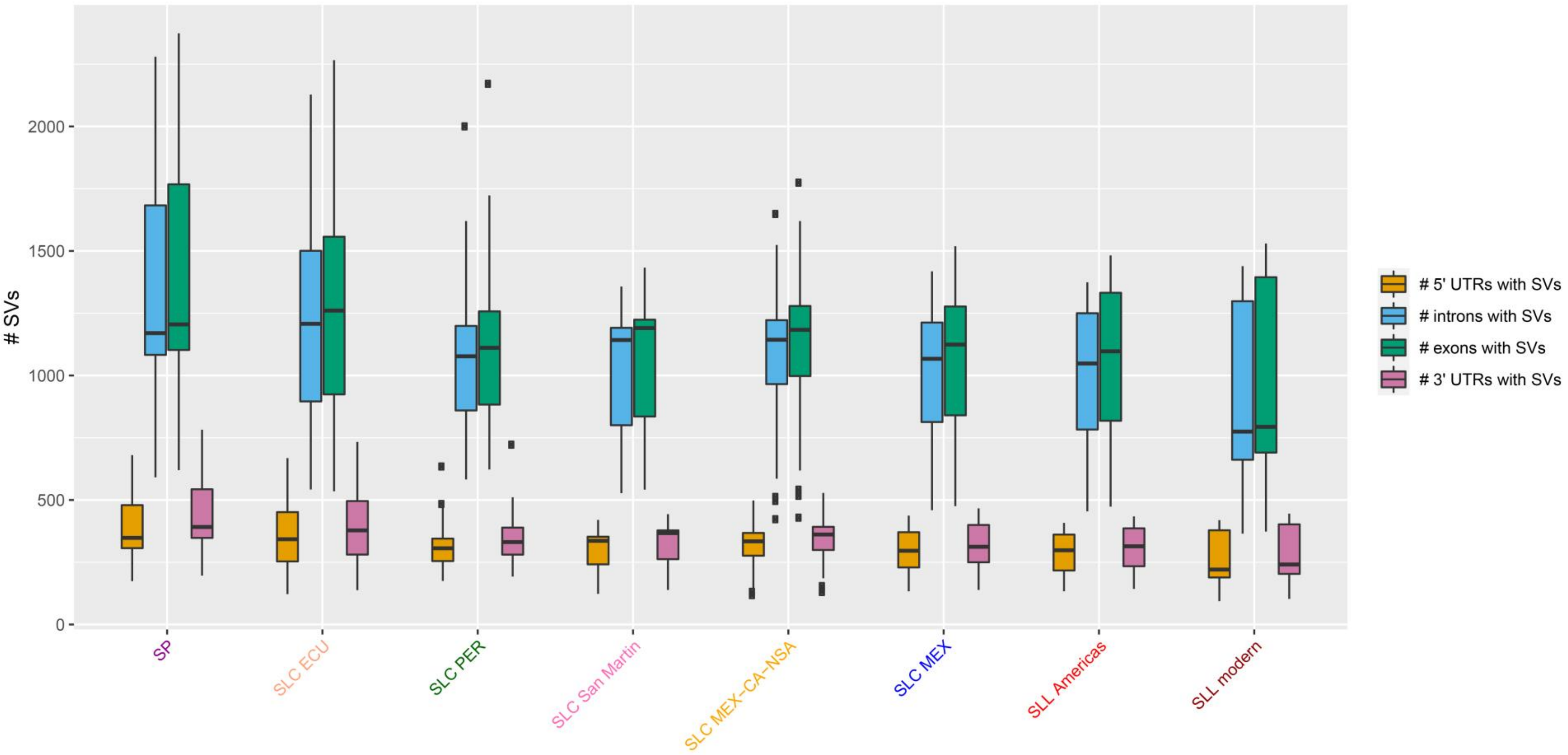


Supplementary fig. 5. A genome-wide comparison of the number of SVs within or outside 5’ UTRS, introns, exons, and 3’ UTRs in each tomato population.
