## supplementary file 1 for "The evolutionary dynamics of genetic mutational load throughout tomato domestication history"

### SlMET1L, SL2.50ch01, 760235

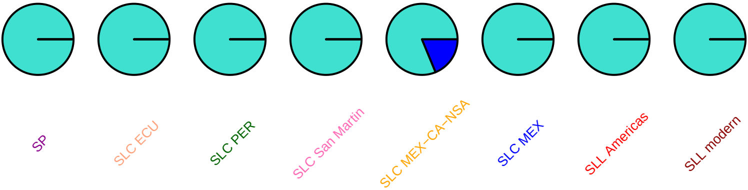

#### cevi1, SL2.50ch01, 889169

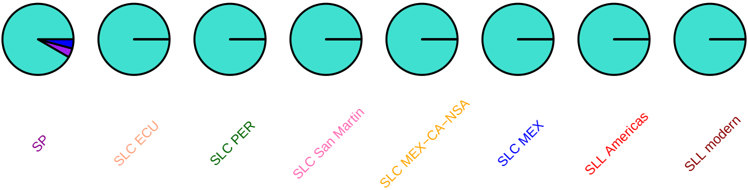

### cevi1, SL2.50ch01, 890331

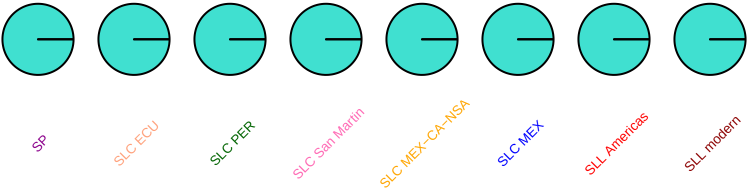

### loxc, SL2.50ch01, 1122500

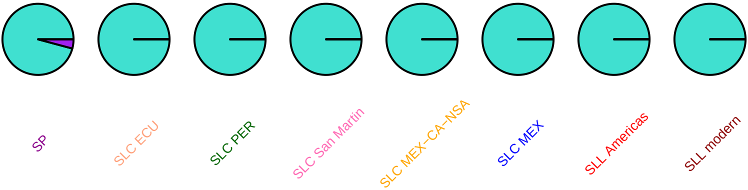

#### SlOFP3, SL2.50ch01, 1971431

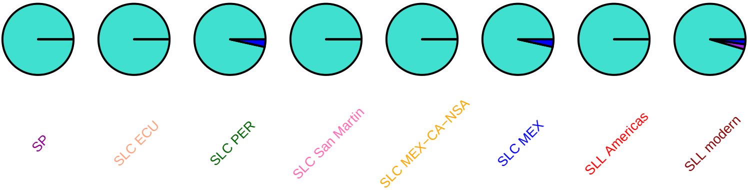

### SlSTP9, SL2.50ch01, 2371273

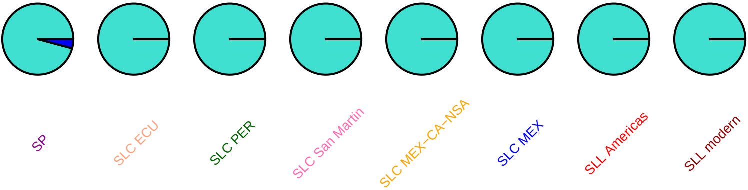

# DQ304483, SL2.50ch01, 2466236

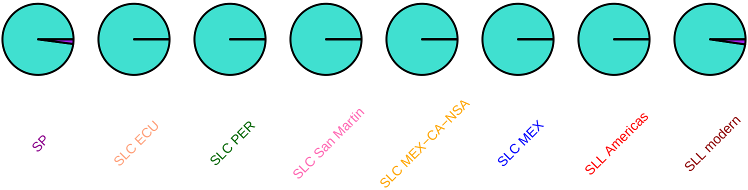

### par1, SL2.50ch01, 2639104

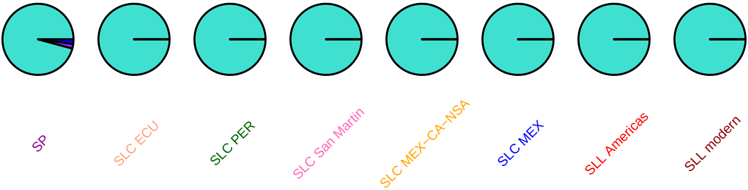

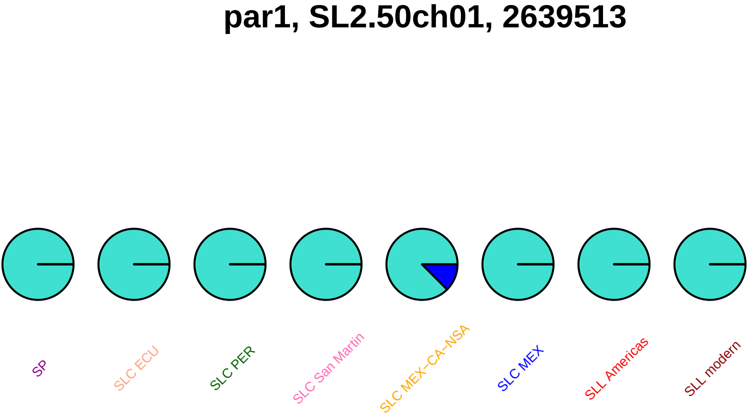
par1, SL2.50ch01, 2639513
SP
SLC ECU
SLC PER
SLC San Martin
SLC MEX−CA−NSA
SLC MEX
SLL Americas
SLL modern

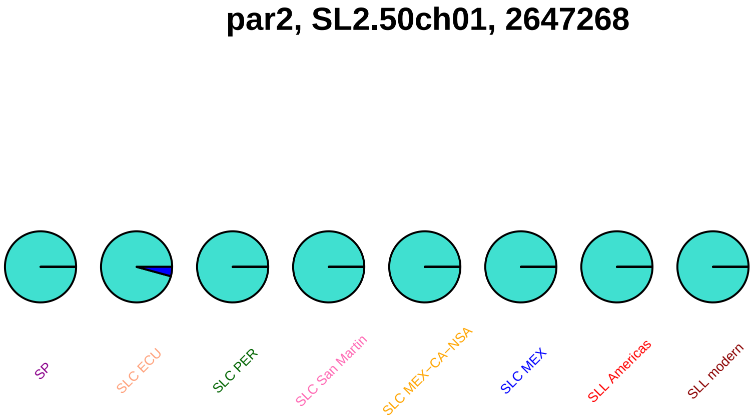

par2, SL2.50ch01, 2647268
SP
SLC ECU
SLC PER
SLC San Martin
SLC MEX−CA−NSA
SLC MEX
SLL Americas
SLL modern

### man4, SL2.50ch01, 2755417

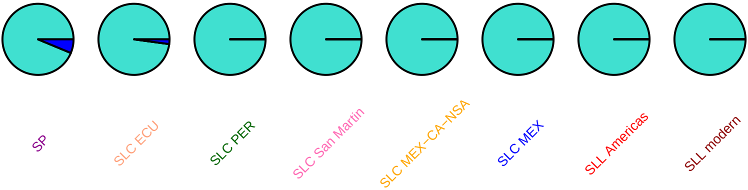

#### man4, SL2.50ch01, 2755599

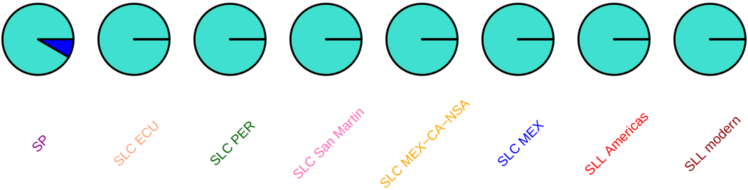

### man4, SL2.50ch01, 2756870

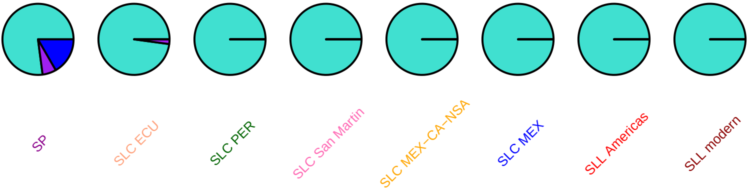

#### man4, SL2.50ch01, 2757708

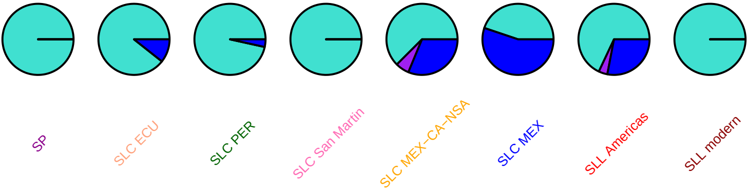

### SlCycB1_4, SL2.50ch01, 3043994

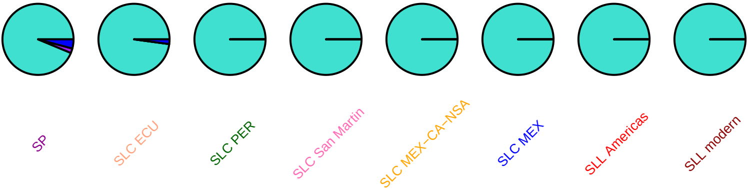

### SlCycB1_4, SL2.50ch01, 3044246

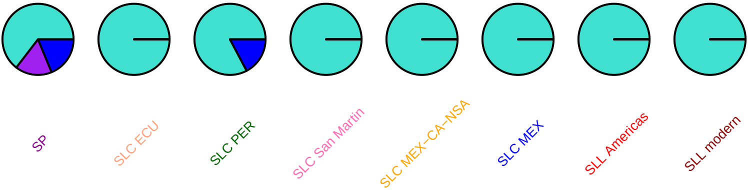

#### eil, SL2.50ch01, 3204841

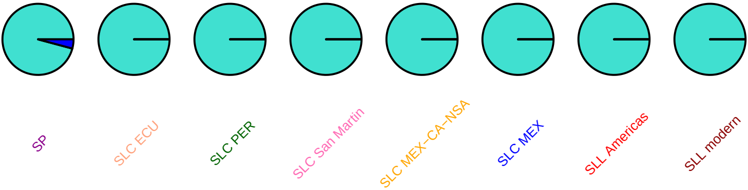

#### sit, SL2.50ch01, 3255759

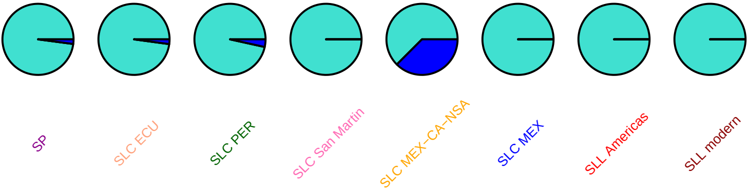

#### SlSTP5, SL2.50ch01, 5549372

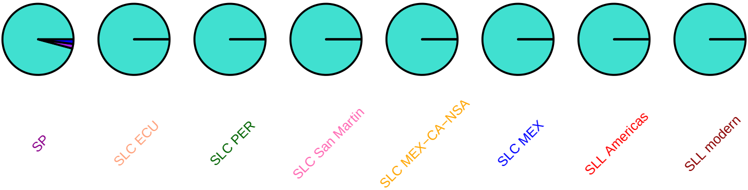

### SlSTP5, SL2.50ch01, 5549924

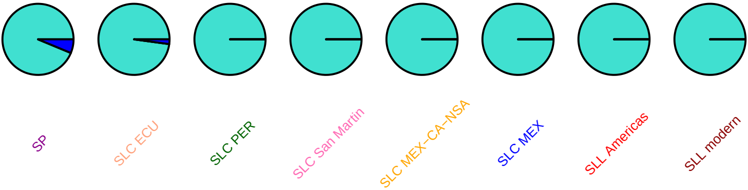

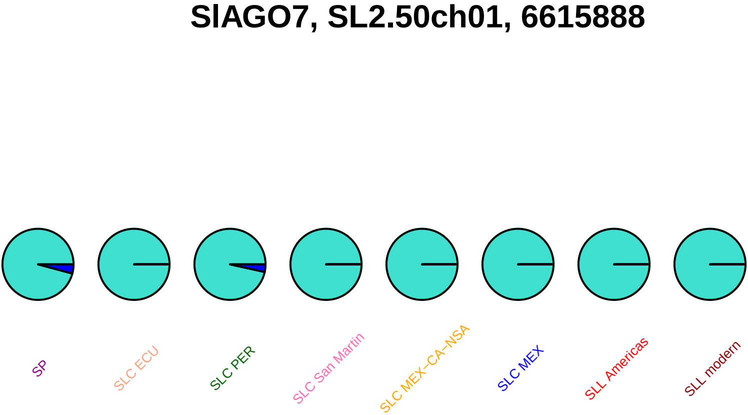

SlAGO7, SL2.50ch01, 6615888
SP
SLC ECU
SLC PER
SLC San Martin
SLC MEX–CA–NSA
SLC MEX
SLL Americas
SLL modern

**shy, SL2.50ch01, 7015274**

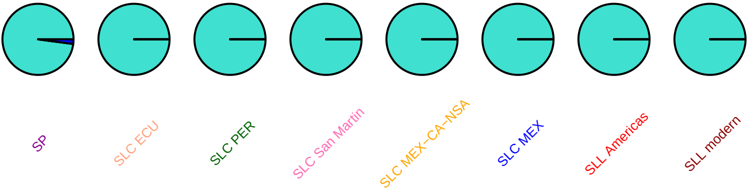

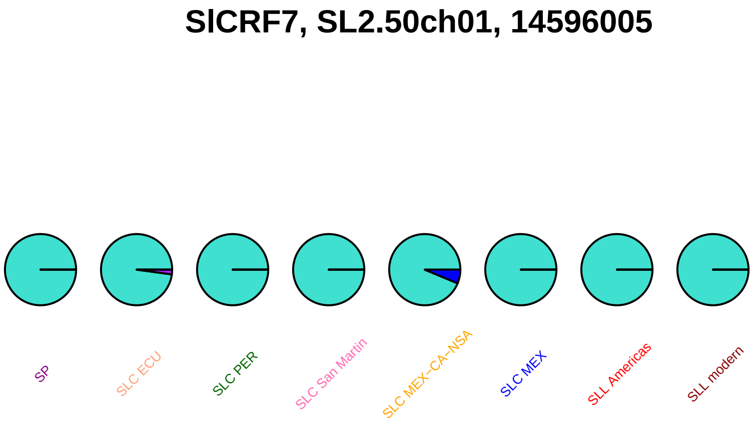
SlCRF7, SL2.50ch01, 14596005
SP
SLC ECU
SLC PER
SLC San Martin
SLC MEX–CA–NSA
SLC MEX
SLL Americas
SLL modern

### SlCRF7, SL2.50ch01, 14596288

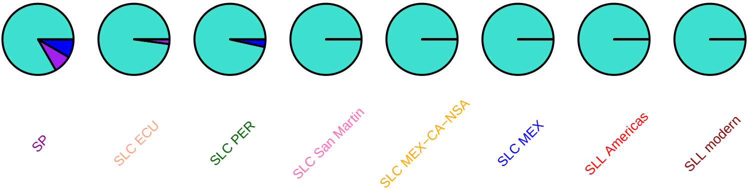

#### SlbHLH002, SL2.50ch01, 60448829

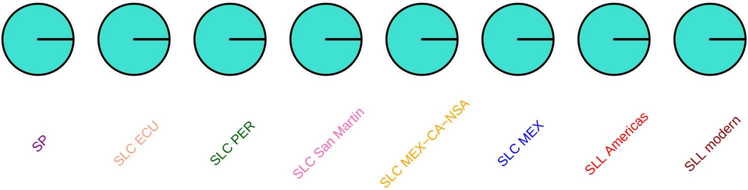

#### SlMYB2, SL2.50ch01, 64321852

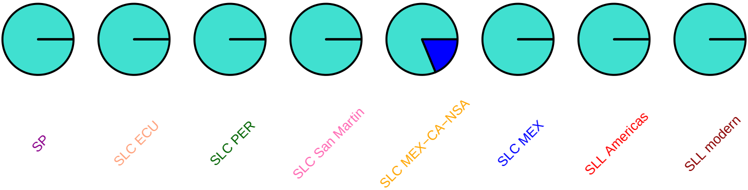

### SlMAPKKK3, SL2.50ch01, 68825685

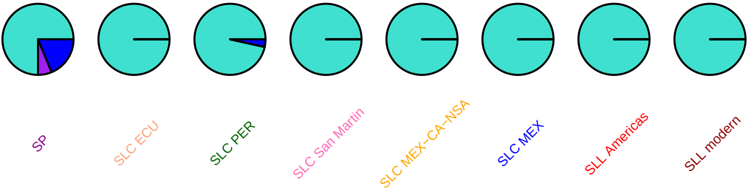

### phyB1, SL2.50ch01, 68866199

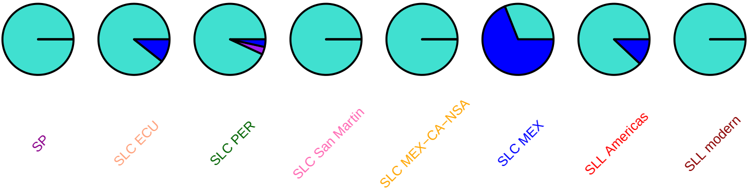

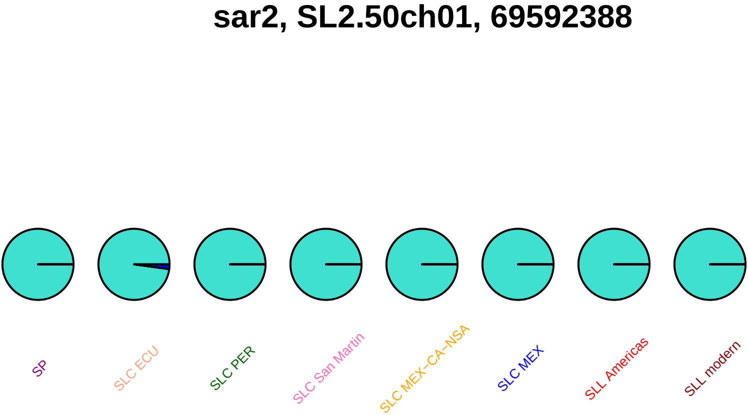
sar2, SL2.50ch01, 69592388
SP
SLC ECU
SLC PER
SLC San Martin
SLC MEX−CA−NSA
SLC MEX
SLL Americas
SLL modern

### SlySBP12a, SL2.50ch01, 77196510

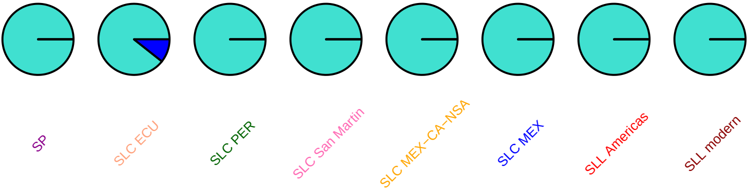

### xegip, SL2.50ch01, 79139722

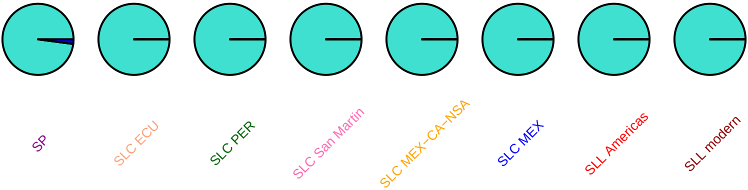

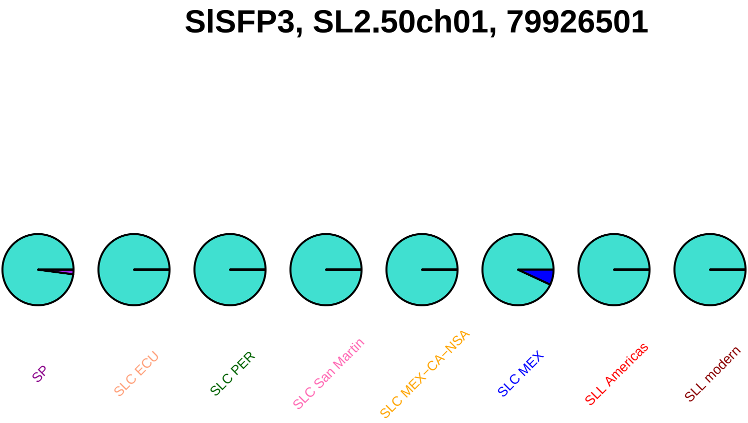
SlSFP3, SL2.50ch01, 79926501
SP
SLC ECU
SLC PER
SLC San Martin
SLC MEX−CA−NSA
SLC MEX
SLL Americas
SLL modern

#### gstt1, SL2.50ch01, 81643980

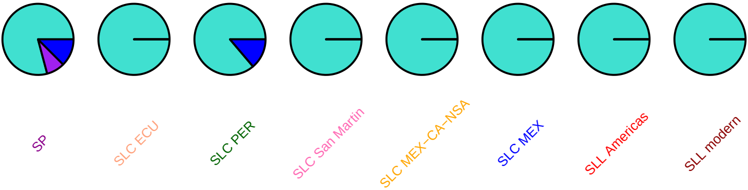

### SlbHLH076, SL2.50ch01, 81798134

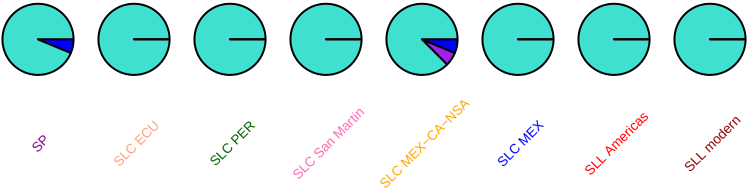

### ccd1b, SL2.50ch01, 82209833

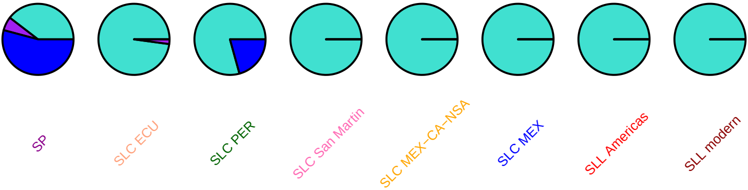

### adh, SL2.50ch01, 82587481

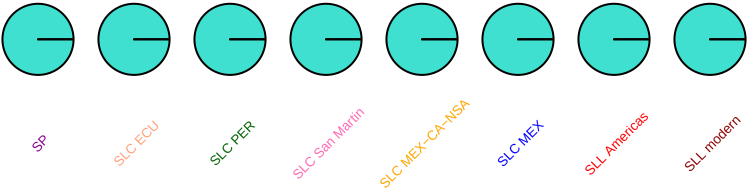

# ao2, SL2.50ch01, 83033820

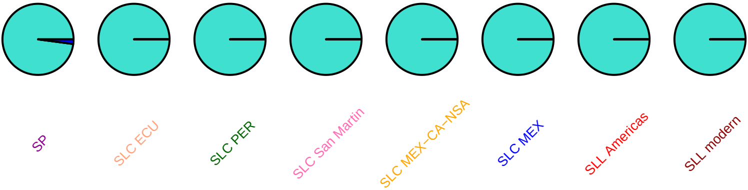

# ao2, SL2.50ch01, 83036473

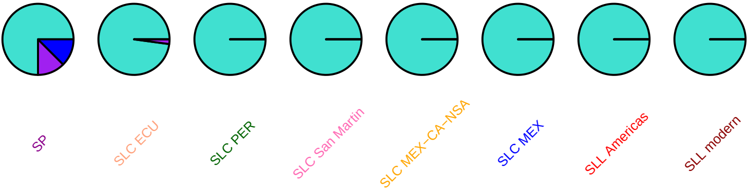

## ao2, SL2.50ch01, 83038544

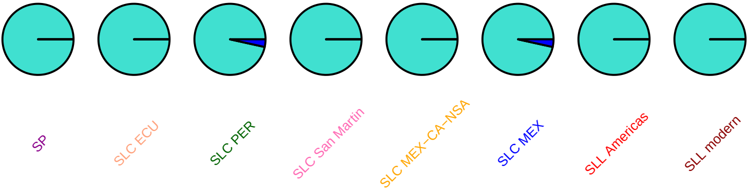

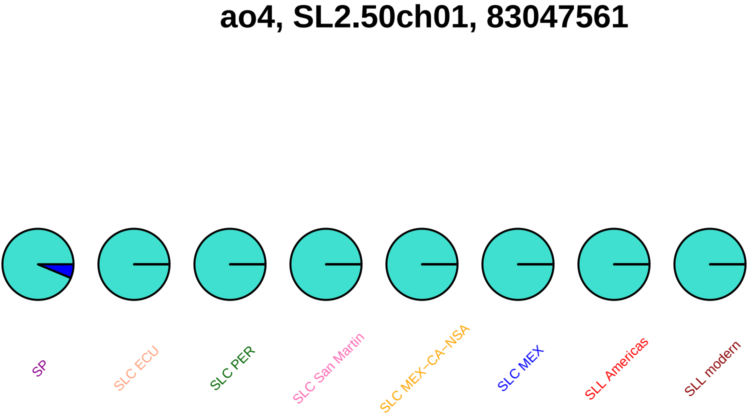
ao4, SL2.50ch01, 83047561
SP
SLC ECU
SLC PER
SLC San Martin
SLC MEX–CA–NSA
SLC MEX
SLL Americas
SLL modern

# ao4, SL2.50ch01, 83048571

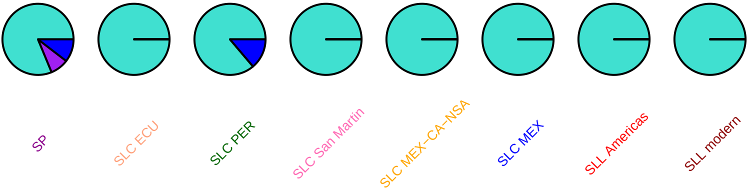

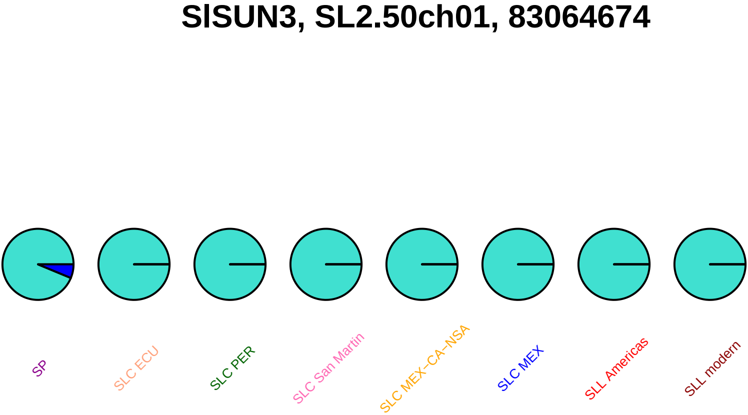
SlSUN3, SL2.50ch01, 83064674
SP
SLC ECU
SLC PER
SLC San Martin
SLC MEX–CA–NSA
SLC MEX
SLL Americas
SLL modern

#### cmdh, SL2.50ch01, 84351522

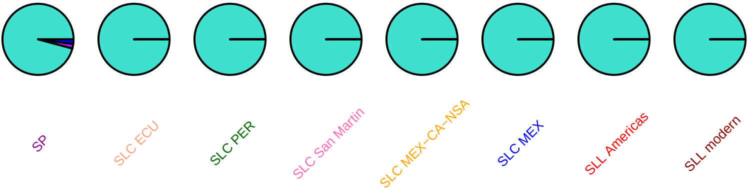

# CD2, SL2.50ch01, 85221979

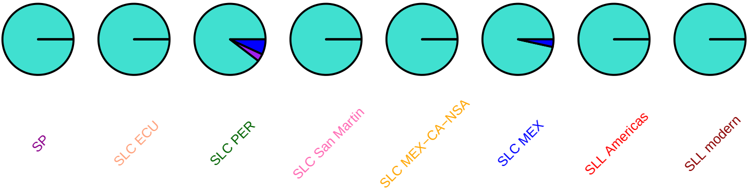

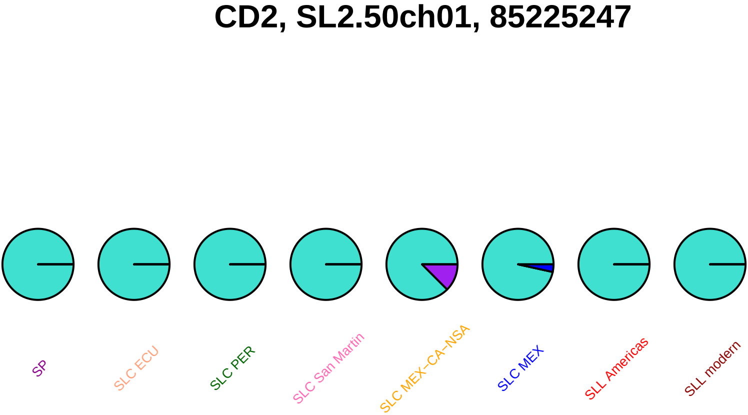
CD2, SL2.50ch01, 85225247
SP
SLC ECU
SLC PER
SLC San Martin
SLC MEX−CA−NSA
SLC MEX
SLL Americas
SLL modern

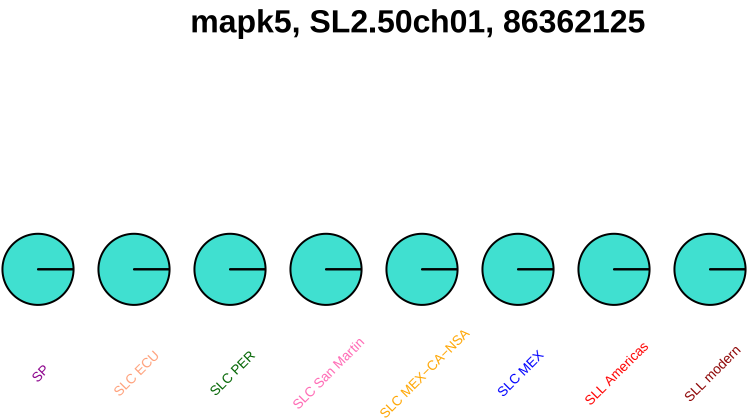
mapk5, SL2.50ch01, 86362125
SP
SLC ECU
SLC PER
SLC San Martin
SLC MEX−CA−NSA
SLC MEX
SLL Americas
SLL modern

#### mapk5, SL2.50ch01, 86362751

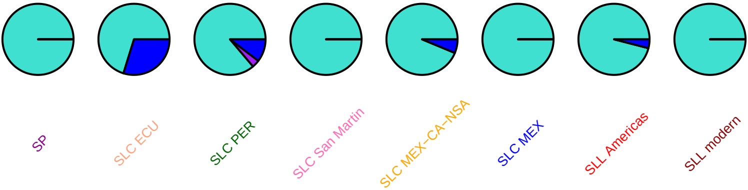

### acc2, SL2.50ch01, 86453656

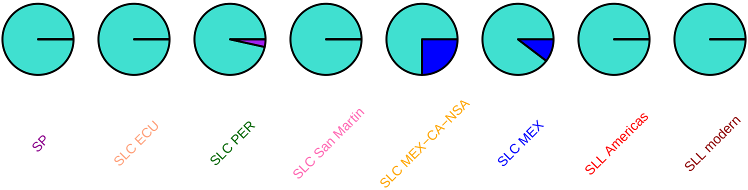

#### SlDEAH3, SL2.50ch01, 86902569

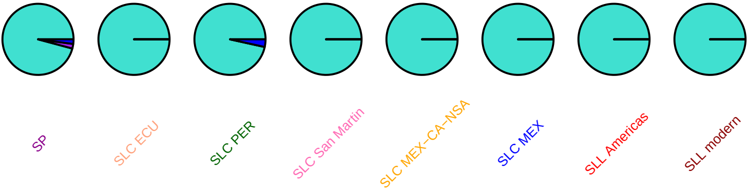

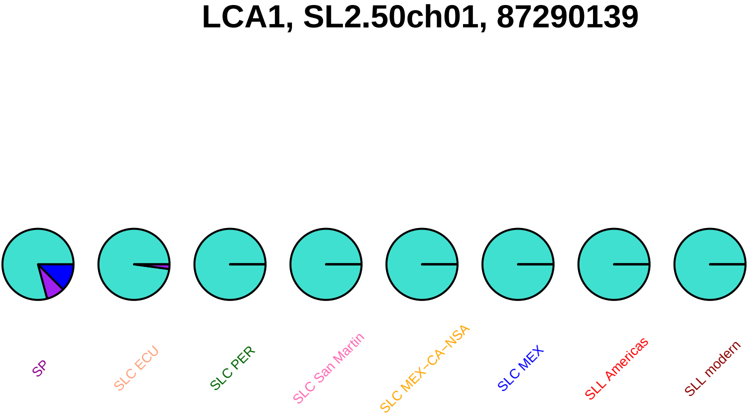

LCA1, SL2.50ch01, 87290139
SP
SLC ECU
SLC PER
SLC San Martin
SLC MEX−CA−NSA
SLC MEX
SLL Americas
SLL modern

### eil3, SL2.50ch01, 87804867

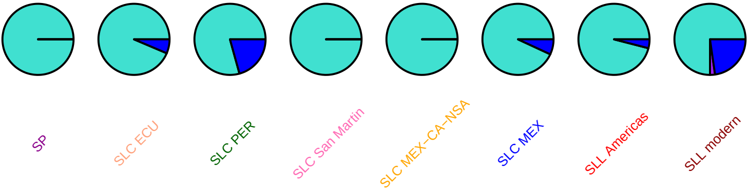

### SlIAA16, SL2.50ch01, 88196770

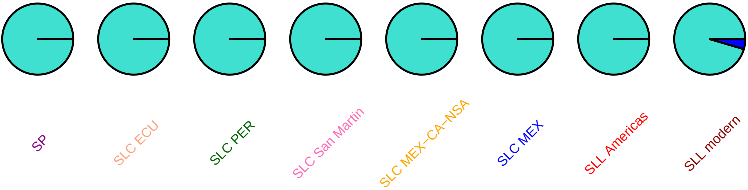

#### AnnSl3, SL2.50ch01, 88337322

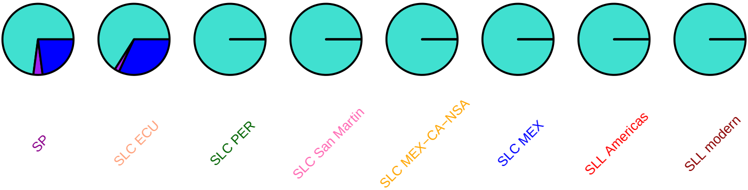

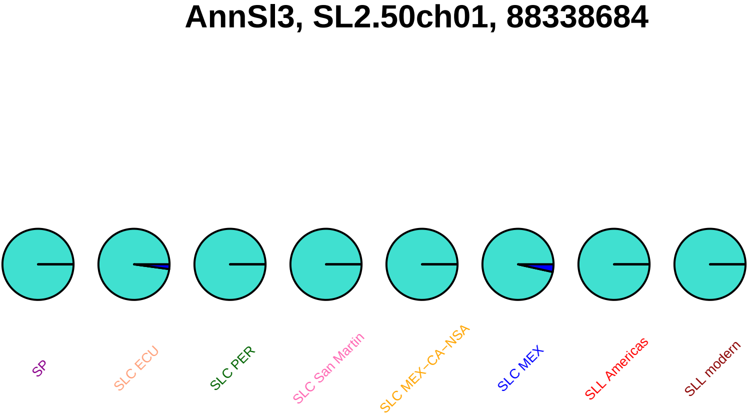
AnnSl3, SL2.50ch01, 88338684
SP
SLC ECU
SLC PER
SLC San Martin
SLC MEX–CA–NSA
SLC MEX
SLL Americas
SLL modern

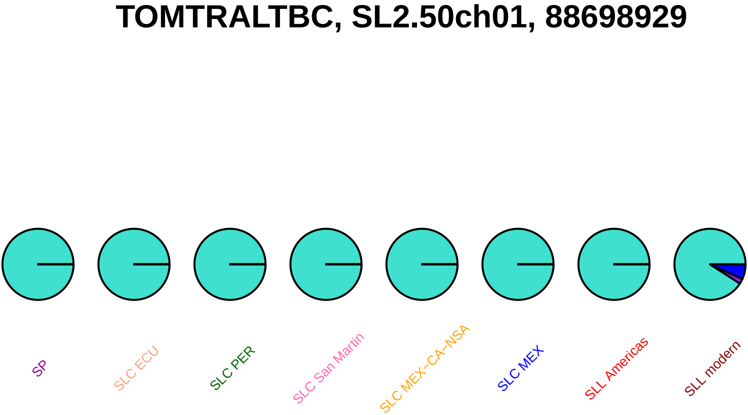
TOMTRALTBC, SL2.50ch01, 88698929
SP
SLC ECU
SLC PER
SLC San Martin
SLC MEX−CA−NSA
SLC MEX
SLL Americas
SLL modern

#### SlSFP1, SL2.50ch01, 88992196

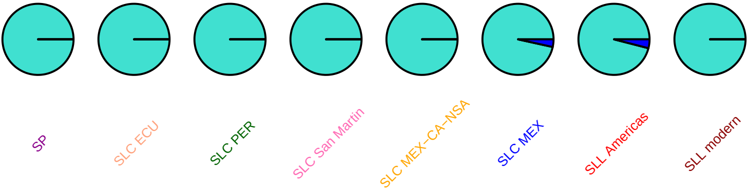

### SlSFP8, SL2.50ch01, 89047594

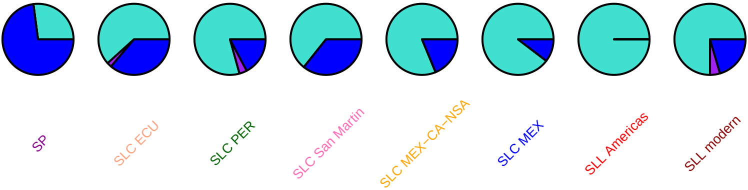

### SlSFP8, SL2.50ch01, 89049163

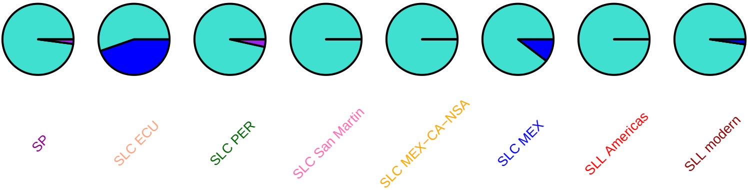

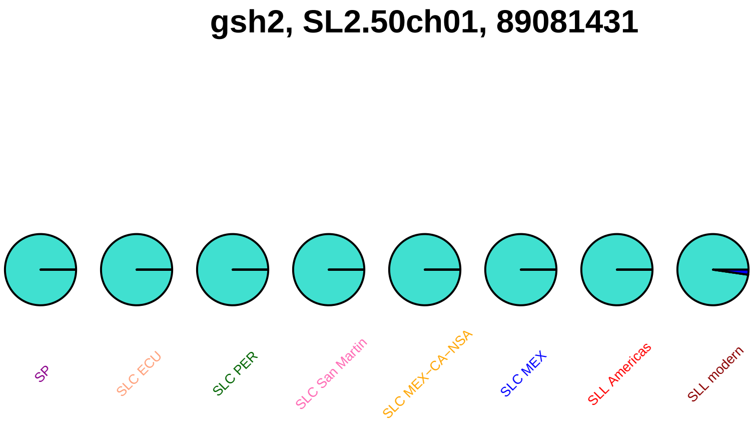
gsh2, SL2.50ch01, 89081431
SP
SLC ECU
SLC PER
SLC San Martin
SLC MEX–CA–NSA
SLC MEX
SLL Americas
SLL modern

### SlMAPKKK8, SL2.50ch01, 89337447

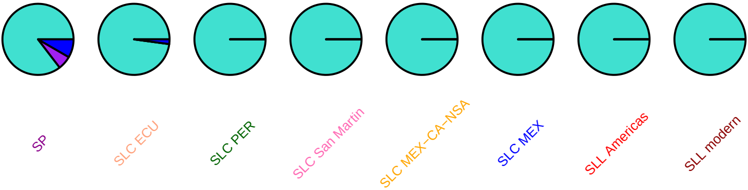

### cevi34, SL2.50ch01, 89475838

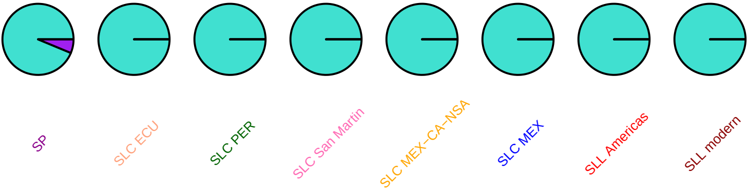

### cevi34, SL2.50ch01, 89481139

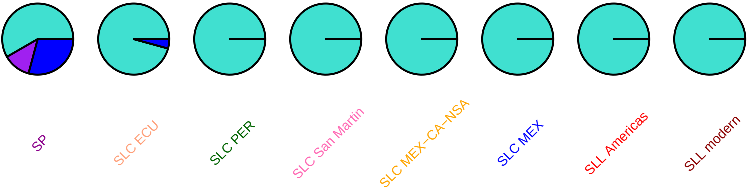

### SlCycH1_1, SL2.50ch01, 89605698

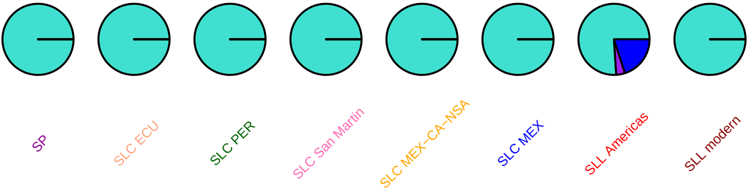

### xth1, SL2.50ch01, 89810516

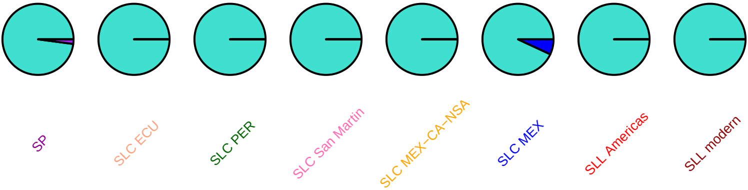

### Interactors of OVATE, SL2.50ch01, 90370976

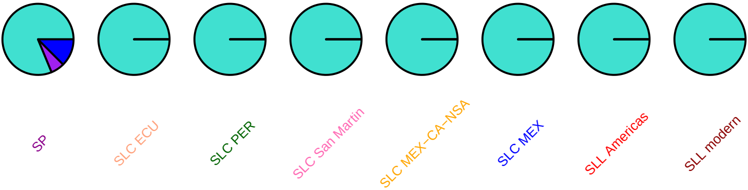

#### sam, SL2.50ch01, 90920106

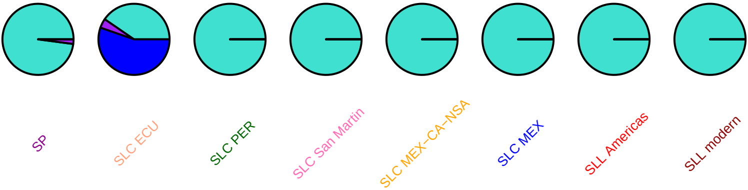

### SlbHLH006, SL2.50ch01, 91125522

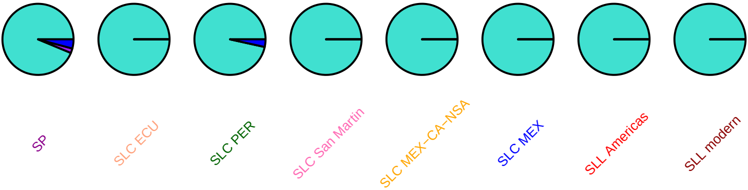

#### SlMYB61, SL2.50ch01, 91166504

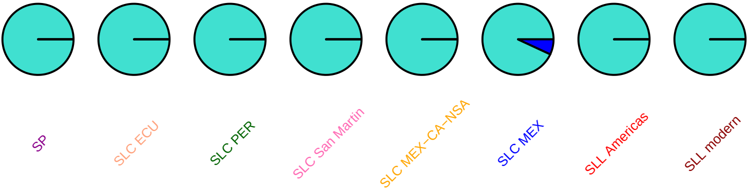

# AF081021, SL2.50ch01, 91325688

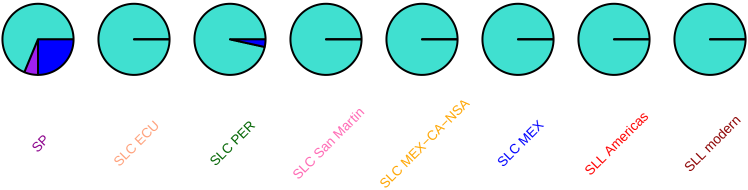

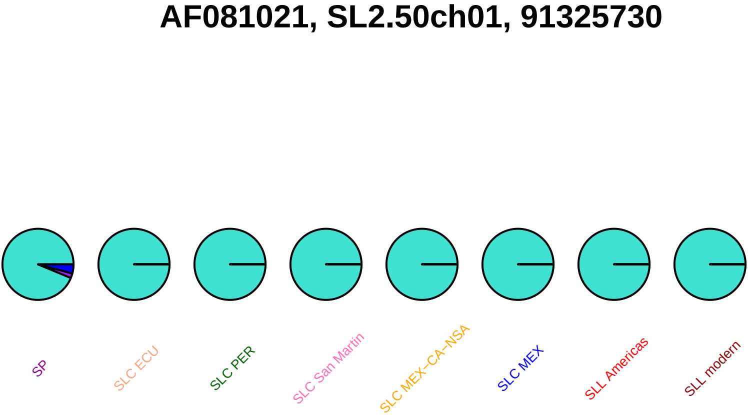
AF081021, SL2.50ch01, 91325730
SP
SLC ECU
SLC PER
SLC San Martin
SLC MEX–CA–NSA
SLC MEX
SLL Americas
SLL modern

## AF081021, SL2.50ch01, 91327992

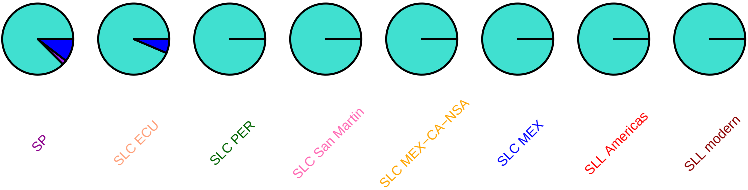

### SlC3H14, SL2.50ch01, 91761883

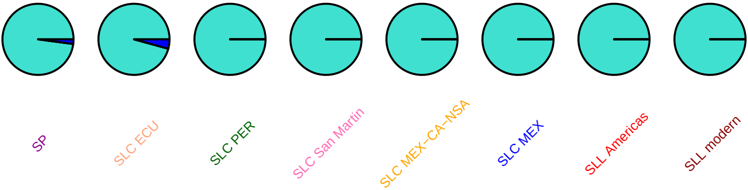

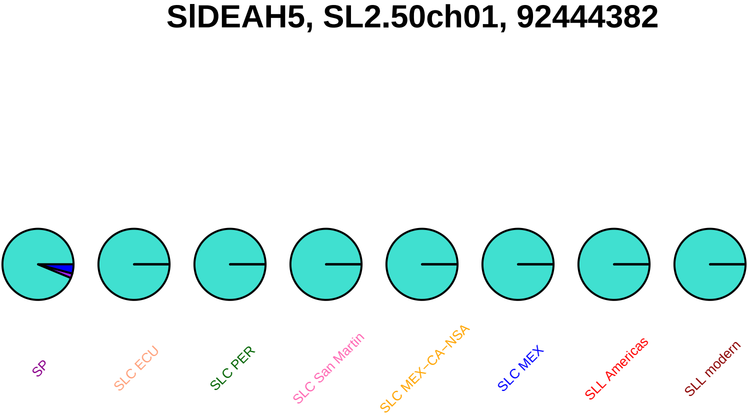
SlDEAH5, SL2.50ch01, 92444382
SP
SLC ECU
SLC PER
SLC San Martin
SLC MEX–CA–NSA
SLC MEX
SLL Americas
SLL modern

# th2, SL2.50ch01, 93495523

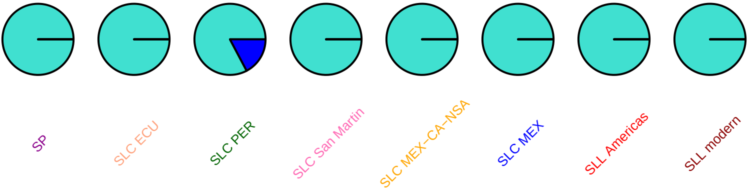

### riba, SL2.50ch01, 93701502

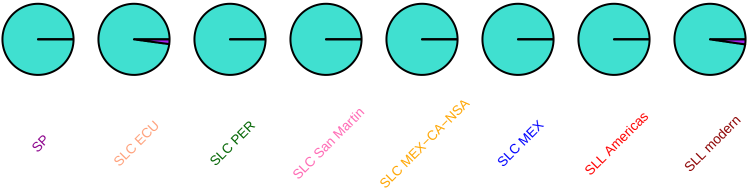

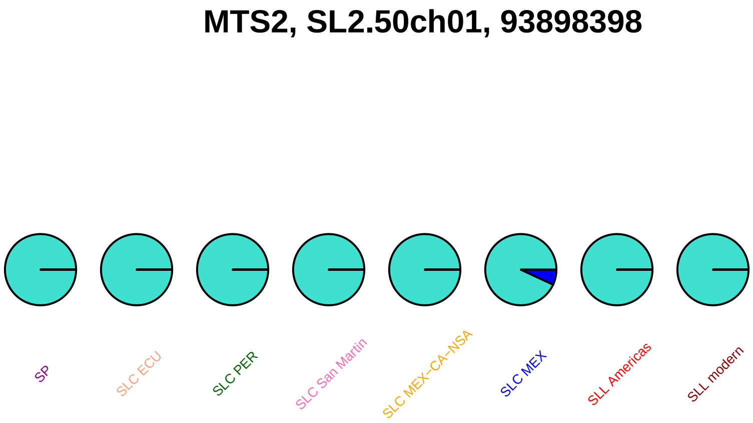
MTS2, SL2.50ch01, 93898398
SP
SLC ECU
SLC PER
SLC San Martin
SLC MEX−CA−NSA
SLC MEX
SLL Americas
SLL modern

### SlbHLH009, SL2.50ch01, 95342070

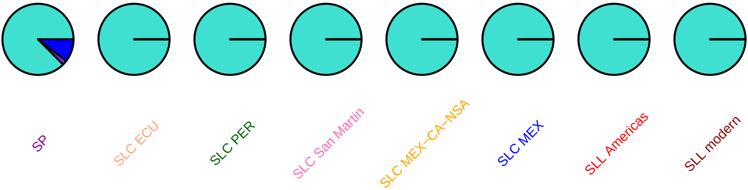

#### dem2, SL2.50ch01, 95622630

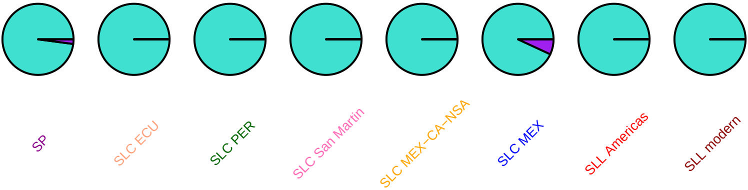

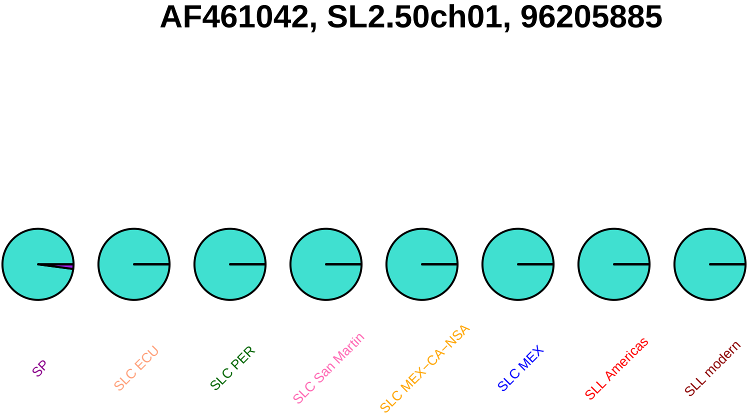
AF461042, SL2.50ch01, 96205885
SP
SLC ECU
SLC PER
SLC San Martin
SLC MEX–CA–NSA
SLC MEX
SLL Americas
SLL modern

### CYP74C4, SL2.50ch01, 96212864

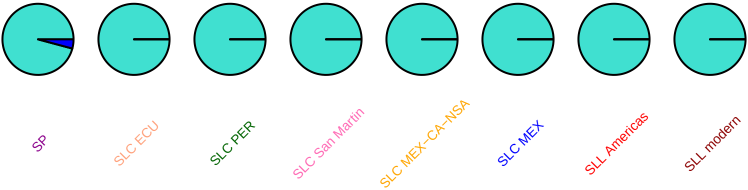

### CYP74C4, SL2.50ch01, 96213779

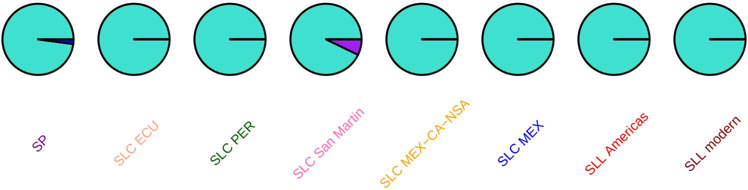

### CYP74C4, SL2.50ch01, 96213996

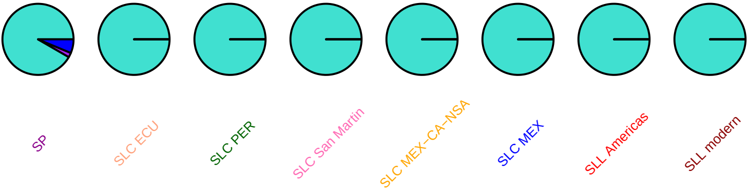

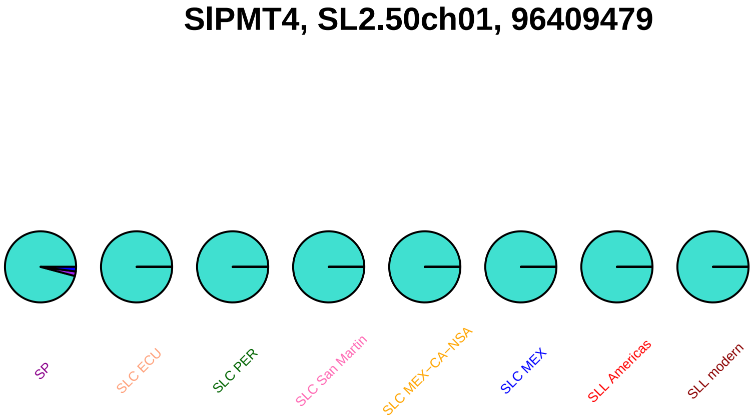
SlPMT4, SL2.50ch01, 96409479
SP
SLC ECU
SLC PER
SLC San Martin
SLC MEX−CA−NSA
SLC MEX
SLL Americas
SLL modern

### agps1, SL2.50ch01, 96637247

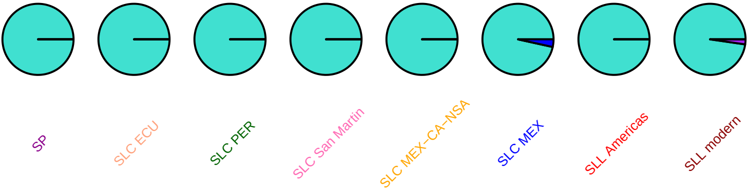

### TOMARGDECA, SL2.50ch01, 97147393

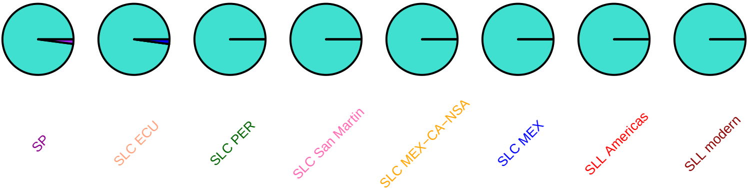

**SlSAUR6, SL2.50ch01, 97240900**

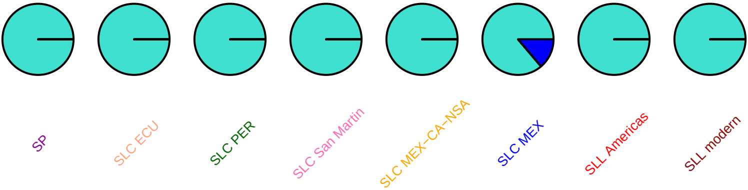

#### SlSAUR10, SL2.50ch01, 97323418

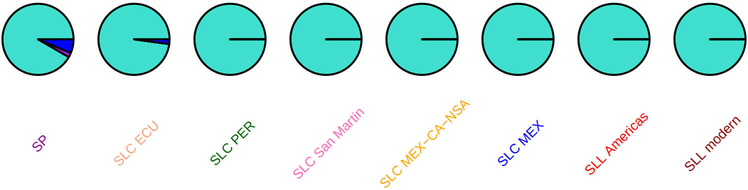

### SlSAUR19, SL2.50ch01, 97349733

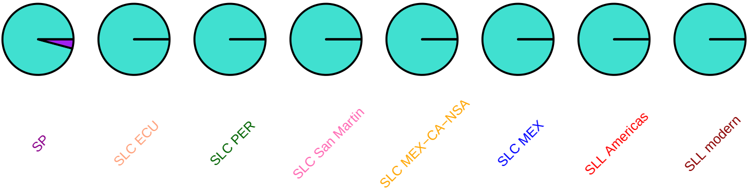

### SlSAUR20, SL2.50ch01, 97363360

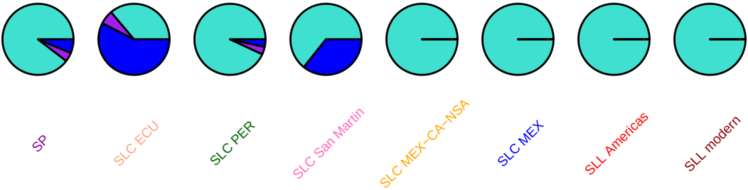

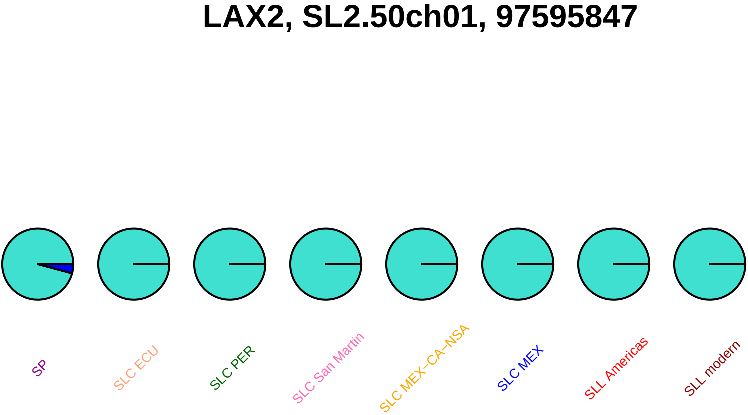
LAX2, SL2.50ch01, 97595847
SP
SLC ECU
SLC PER
SLC San Martin
SLC MEX–CA–NSA
SLC MEX
SLL Americas
SLL modern

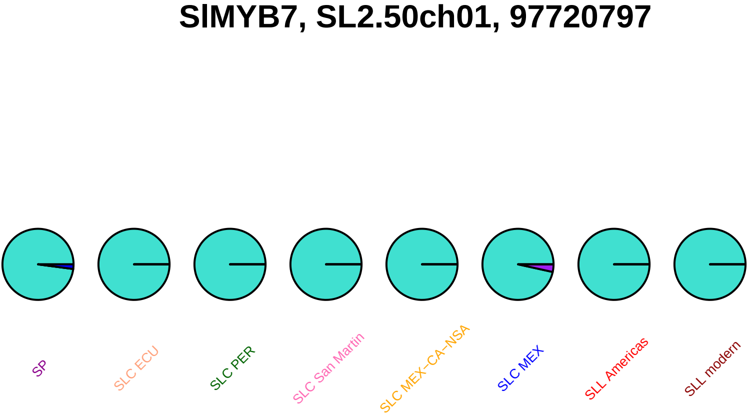

SlMYB7, SL2.50ch01, 97720797
SP
SLC ECU
SLC PER
SLC San Martin
SLC MEX–CA–NSA
SLC MEX
SLL Americas
SLL modern

#### ctd3, SL2.50ch01, 98347436

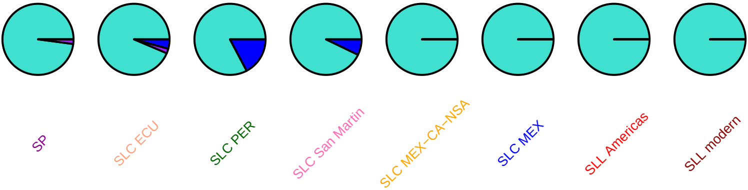

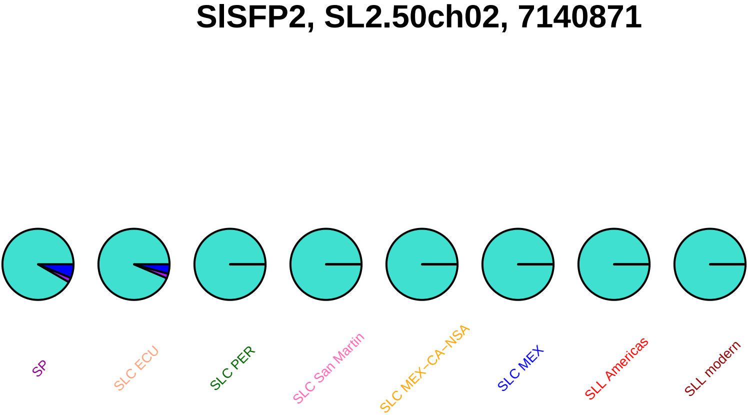
SlSFP2, SL2.50ch02, 7140871
SP
SLC ECU
SLC PER
SLC San Martin
SLC MEX–CA–NSA
SLC MEX
SLL Americas
SLL modern

### scoa2, SL2.50ch02, 8063082

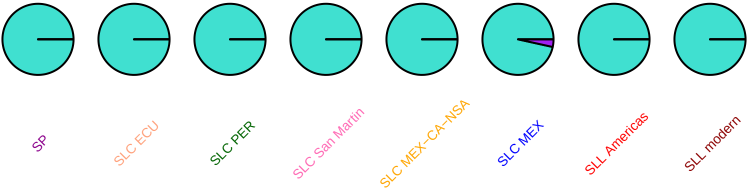

#### CUL4, SL2.50ch02, 22844798

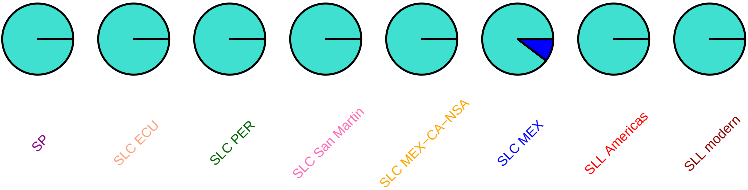

**ddb1, SL2.50ch02, 23364319**

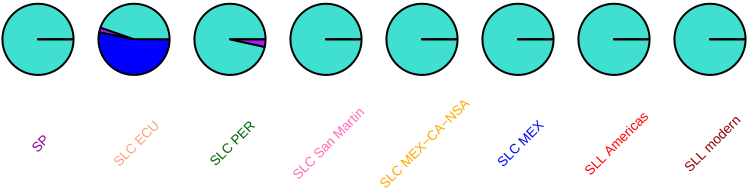

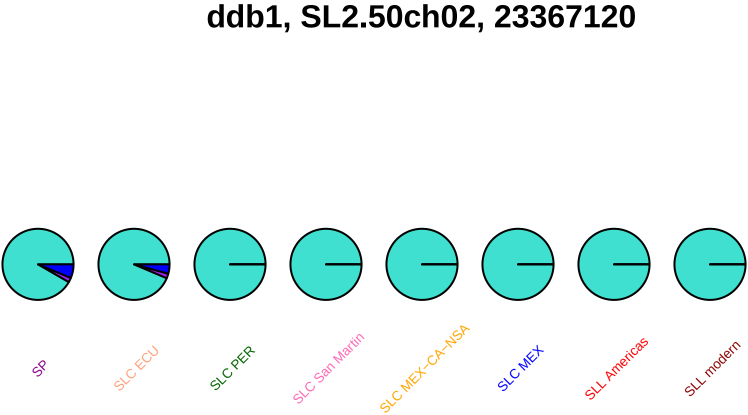

ddb1, SL2.50ch02, 23367120
SP
SLC ECU
SLC PER
SLC San Martin
SLC MEX–CA–NSA
SLC MEX
SLL Americas
SLL modern

### SlMAPKKK12, SL2.50ch02, 27092228

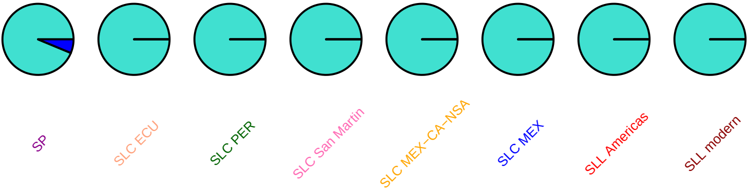

### SlMAPKKK12, SL2.50ch02, 27092315

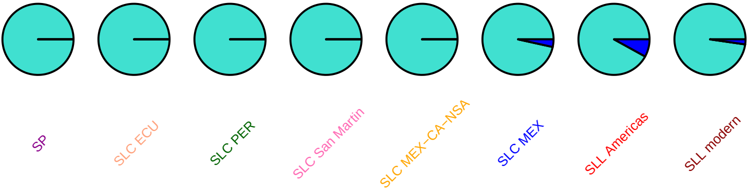

#### SlCycB2_1, SL2.50ch02, 33161685

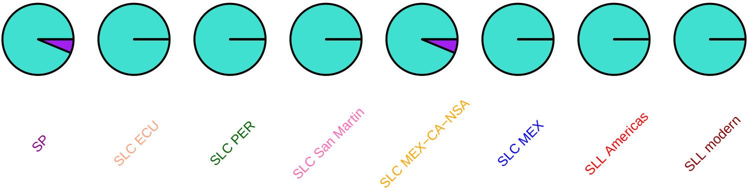

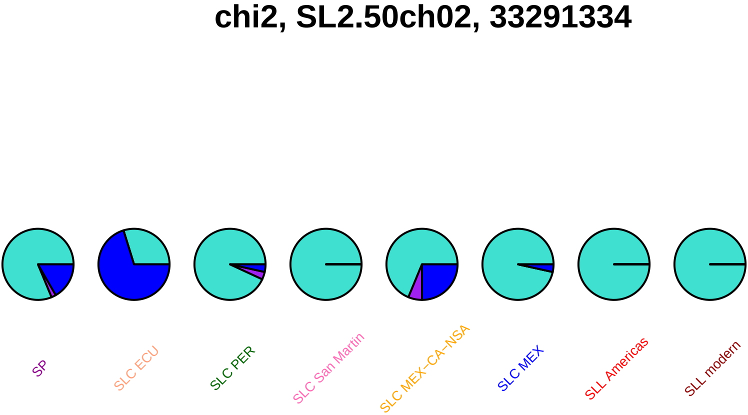

chi2, SL2.50ch02, 33291334
SP
SLC ECU
SLC PER
SLC San Martin
SLC MEX−CA−NSA
SLC MEX
SLL Americas
SLL modern

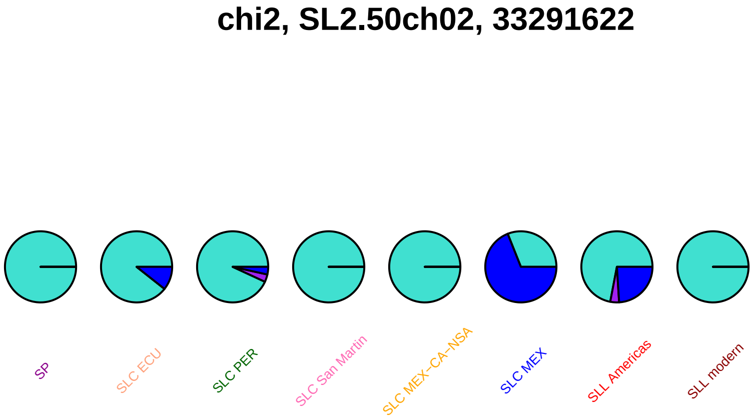
chi2, SL2.50ch02, 33291622
SP
SLC ECU
SLC PER
SLC San Martin
SLC MEX–CA–NSA
SLC MEX
SLL Americas
SLL modern

### grrl2, SL2.50ch02, 34028788

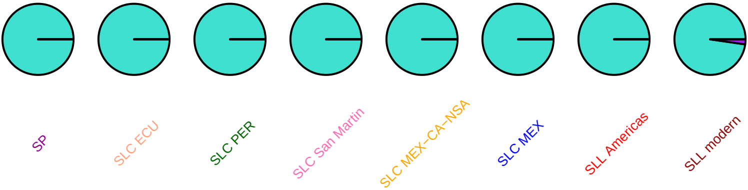

## Tm−1, SL2.50ch02, 34280062

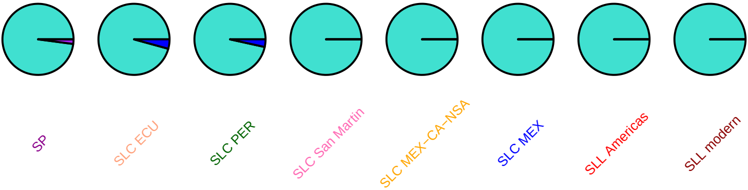

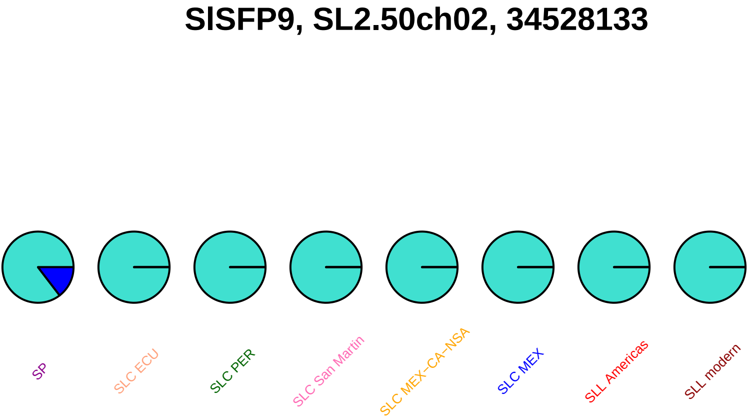
SlSFP9, SL2.50ch02, 34528133
SP
SLC ECU
SLC PER
SLC San Martin
SLC MEX–CA–NSA
SLC MEX
SLL Americas
SLL modern

### gbf12, SL2.50ch02, 34896797

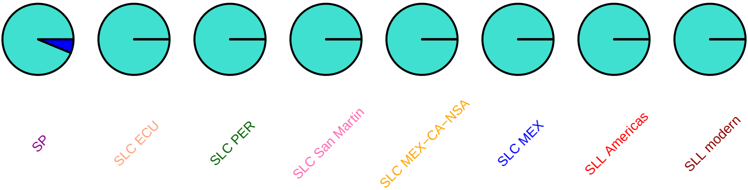

#### app2, SL2.50ch02, 34967425

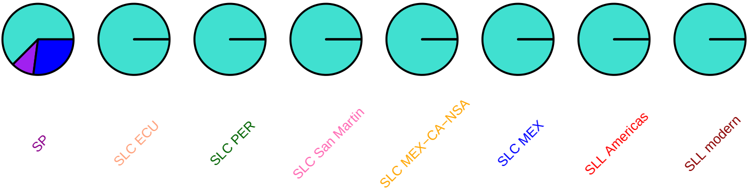

### acs7, SL2.50ch02, 35649434

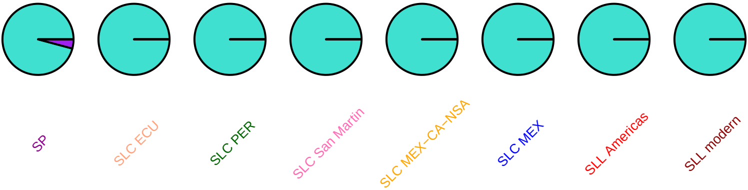

### SlGH3_3, SL2.50ch02, 35957916

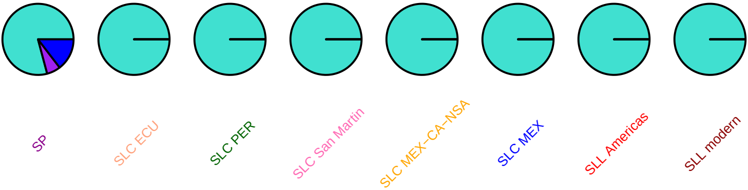

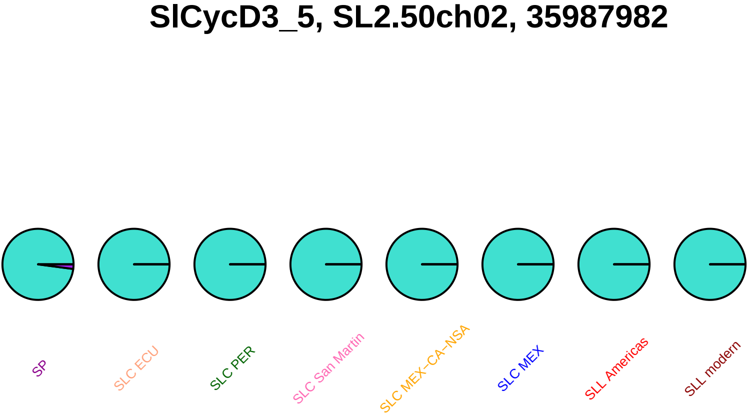
SlCycD3_5, SL2.50ch02, 35987982
SP
SLC ECU
SLC PER
SLC San Martin
SLC MEX−CA−NSA
SLC MEX
SLL Americas
SLL modern

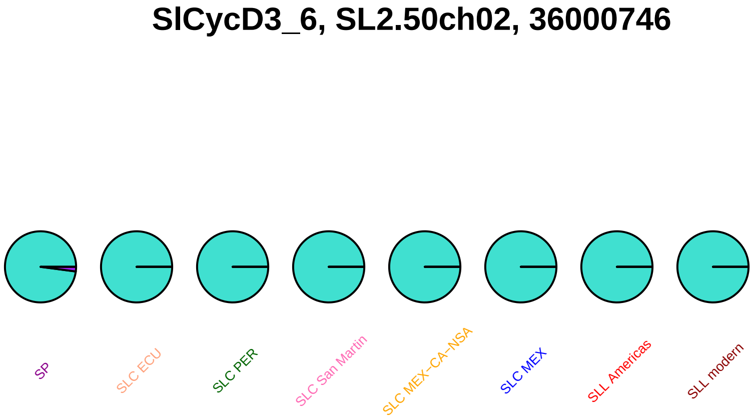
SlCycD3_6, SL2.50ch02, 36000746
SP
SLC ECU
SLC PER
SLC San Martin
SLC MEX−CA−NSA
SLC MEX
SLL Americas
SLL modern

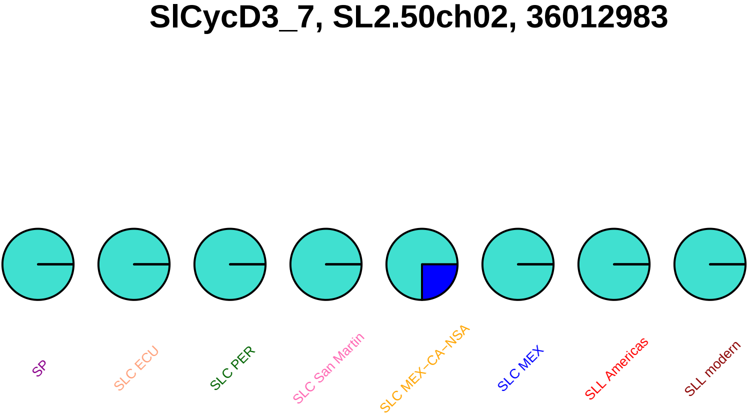
SlCycD3_7, SL2.50ch02, 36012983
SP
SLC ECU
SLC PER
SLC San Martin
SLC MEX–CA–NSA
SLC MEX
SLL Americas
SLL modern

### SlMAPKKK13, SL2.50ch02, 36029439

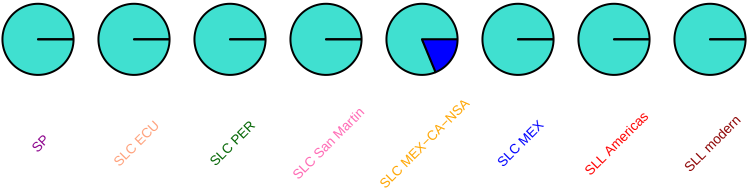

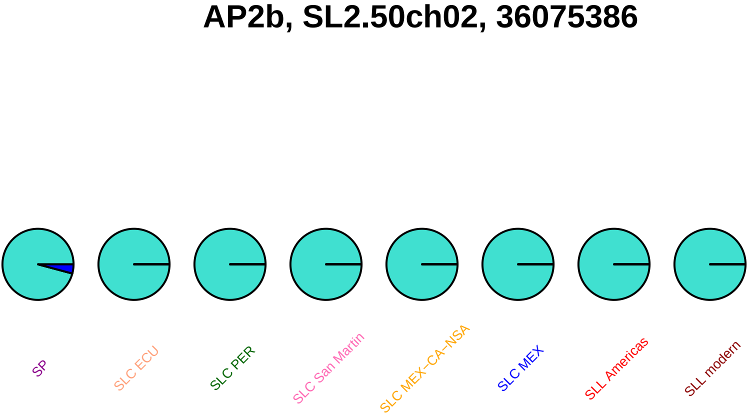
AP2b, SL2.50ch02, 36075386
SP
SLC ECU
SLC PER
SLC San Martin
SLC MEX–CA–NSA
SLC MEX
SLL Americas
SLL modern

## AB190445, SL2.50ch02, 36900335

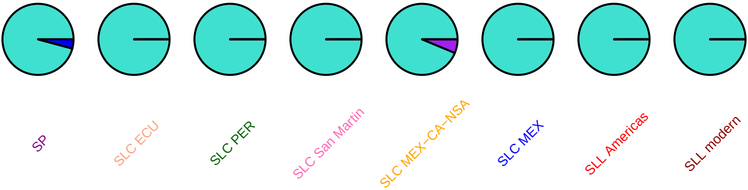

### SlDof3, SL2.50ch02, 37414128

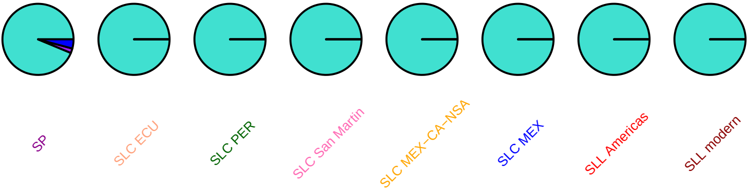

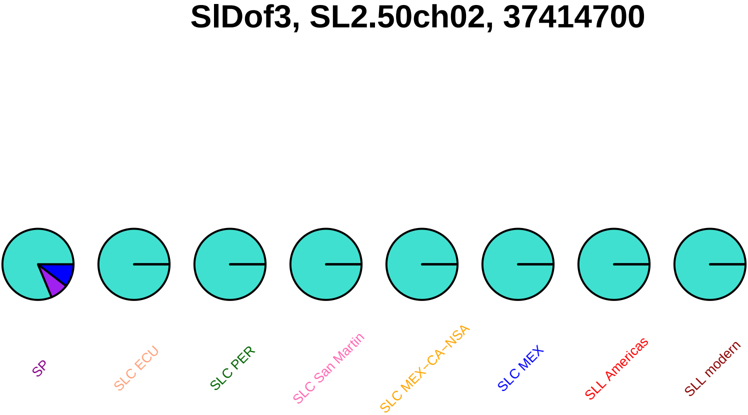
SlDof3, SL2.50ch02, 37414700
SP
SLC ECU
SLC PER
SLC San Martin
SLC MEX−CA−NSA
SLC MEX
SLL Americas
SLL modern

#### SlWRKY72, SL2.50ch02, 37614485

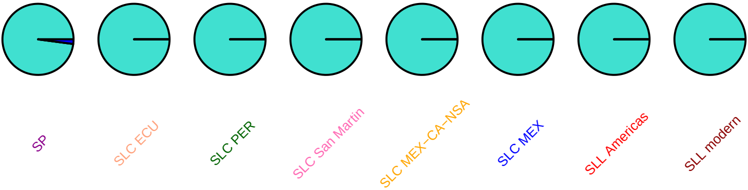

#### tapg2, SL2.50ch02, 37764762

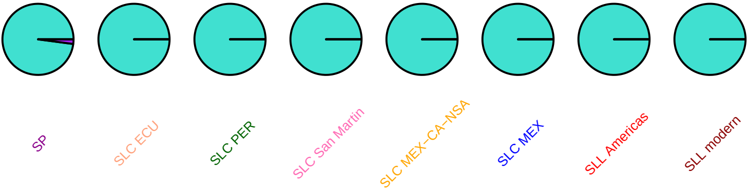

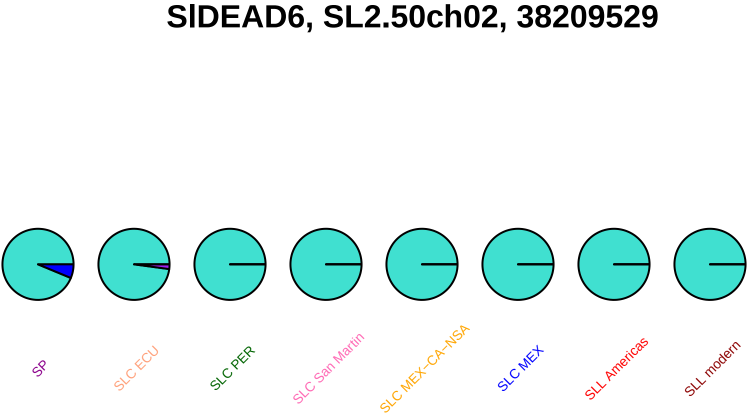
SlDEAD6, SL2.50ch02, 38209529
SP
SLC ECU
SLC PER
SLC San Martin
SLC MEX−CA−NSA
SLC MEX
SLL Americas
SLL modern

### SlDEAD6, SL2.50ch02, 38210045

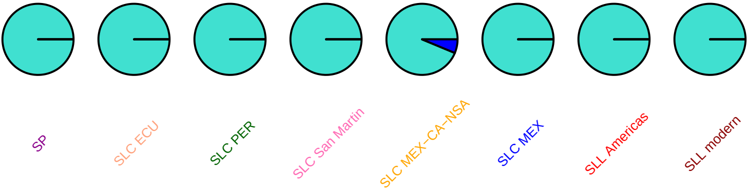

#### irt2, SL2.50ch02, 39155264

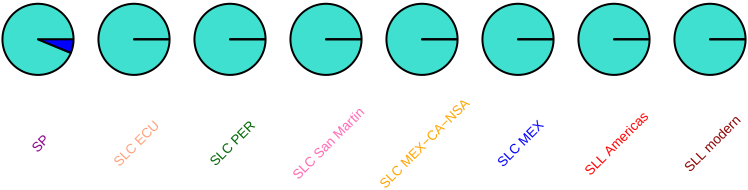

### SlAGO2b, SL2.50ch02, 39215379

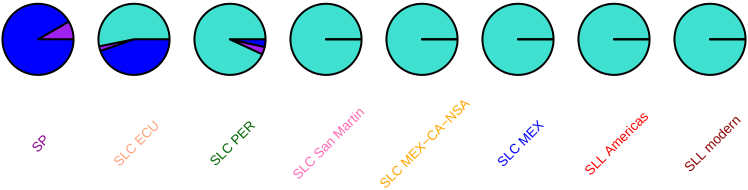

### SlAGO2b, SL2.50ch02, 39215605

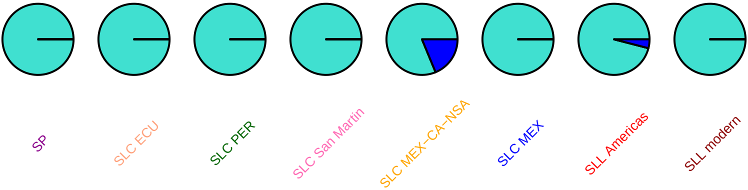

### SlAGO2b, SL2.50ch02, 39217636

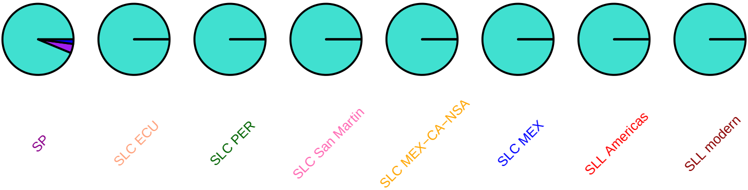

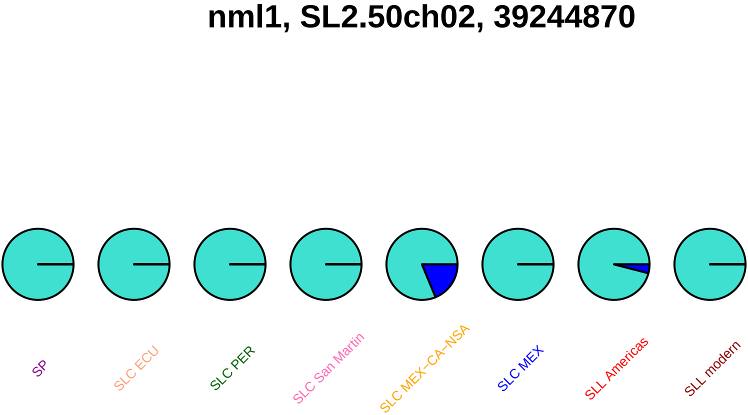
nml1, SL2.50ch02, 39244870
SP
SLC ECU
SLC PER
SLC San Martin
SLC MEX–CA–NSA
SLC MEX
SLL Americas
SLL modern

#### pip1, SL2.50ch02, 42114157

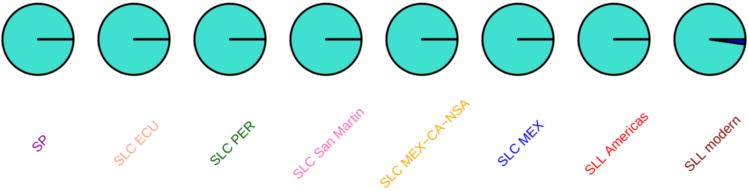

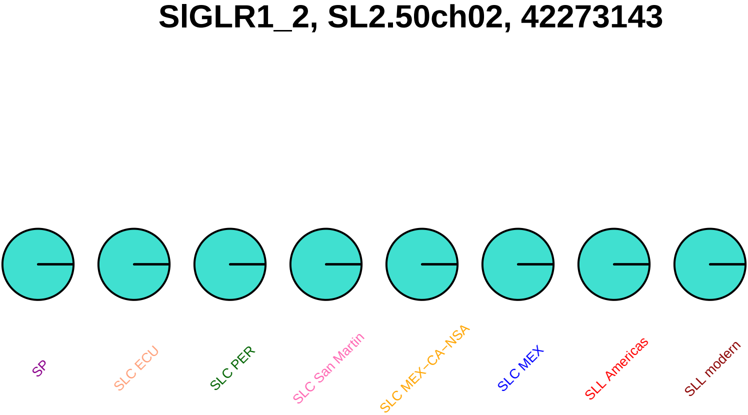
SlGLR1_2, SL2.50ch02, 42273143
SP
SLC ECU
SLC PER
SLC San Martin
SLC MEX−CA−NSA
SLC MEX
SLL Americas
SLL modern

#### SlGLR1_2, SL2.50ch02, 42276097

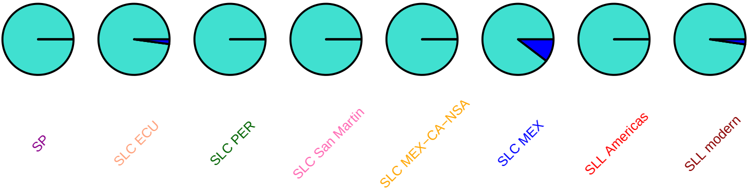

### SlDEAD8, SL2.50ch02, 43522738

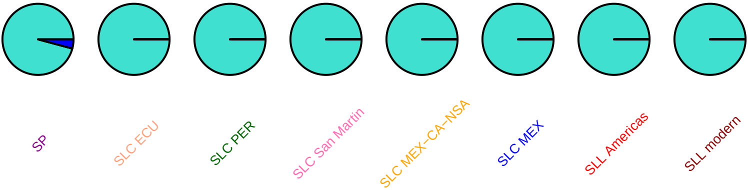

#### psr, SL2.50ch02, 44320064

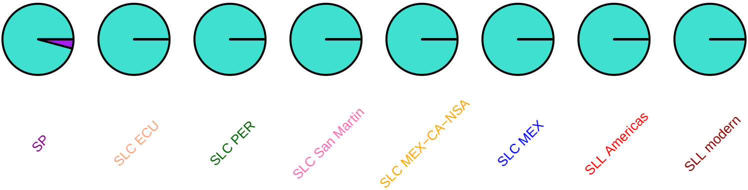

### SlDHS, SL2.50ch02, 44917170

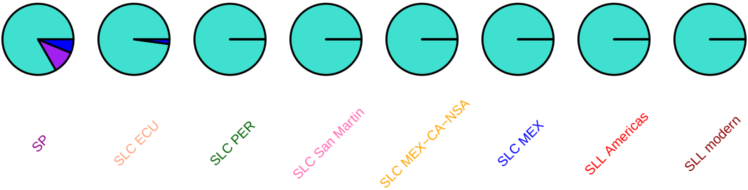

### SlWRKY6, SL2.50ch02, 44992022

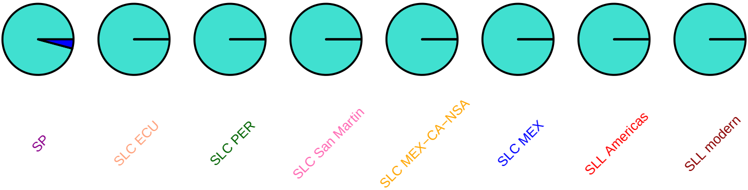

### ftsh6, SL2.50ch02, 45464971

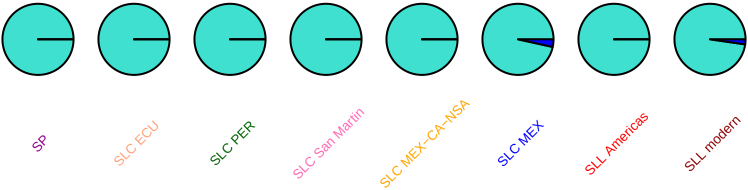

### ftsh6, SL2.50ch02, 45466124

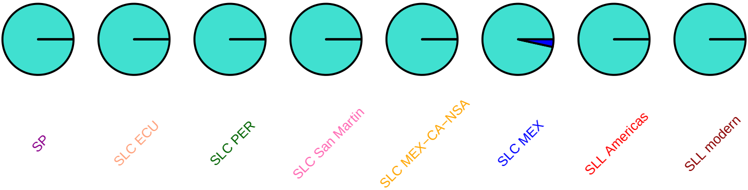

# AF420477, SL2.50ch02, 45952422

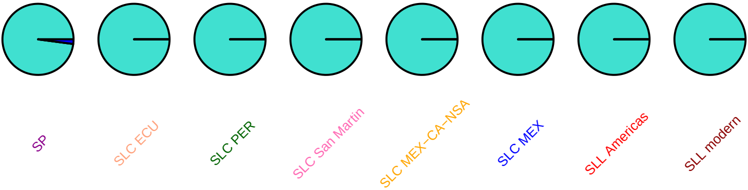

### SlTMT3, SL2.50ch02, 46102538

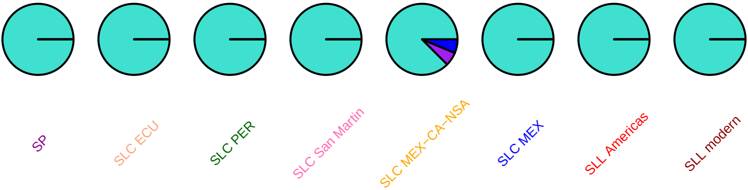

### TOMDHQS, SL2.50ch02, 46938529

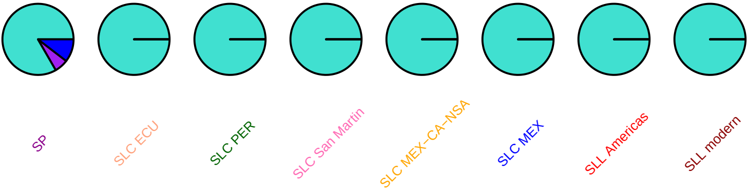

### TOMDHQS, SL2.50ch02, 46939930

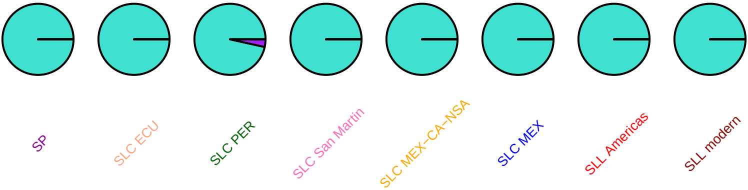

#### SlC3H19, SL2.50ch02, 47375961

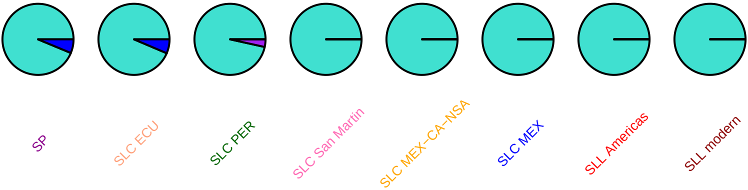

### SlC3H19, SL2.50ch02, 47376186

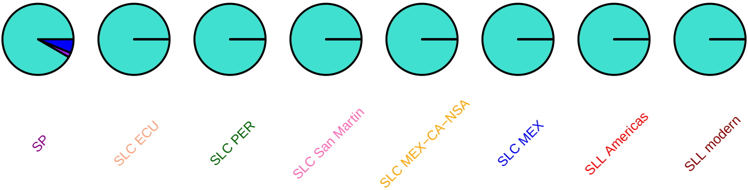

### ggps, SL2.50ch02, 48523169

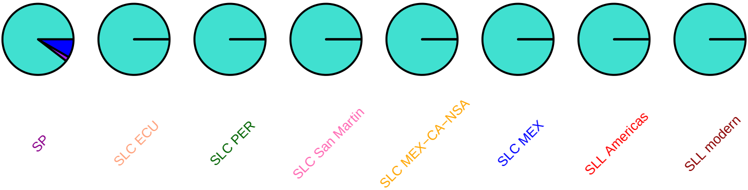

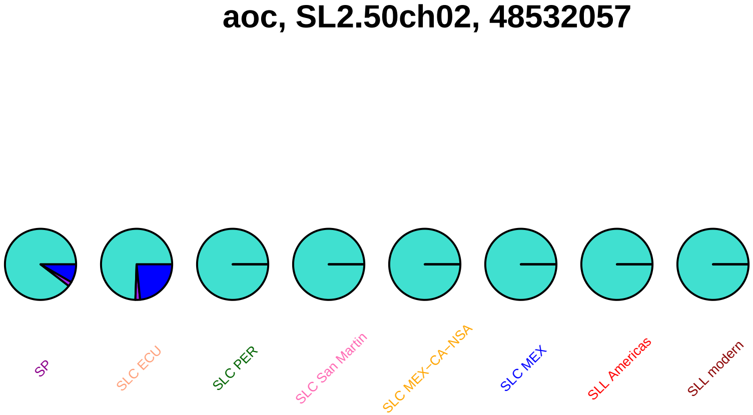

aoc, SL2.50ch02, 48532057
SP
SLC ECU
SLC PER
SLC San Martin
SLC MEX−CA−NSA
SLC MEX
SLL Americas
SLL modern

#### CER6, SL2.50ch02, 48651913

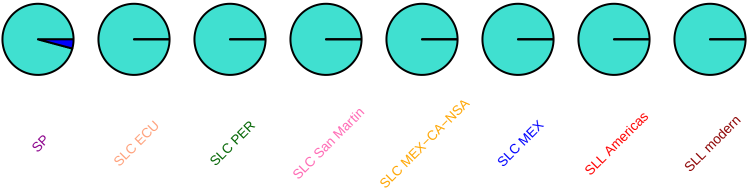

### SlC3H22, SL2.50ch02, 49186088

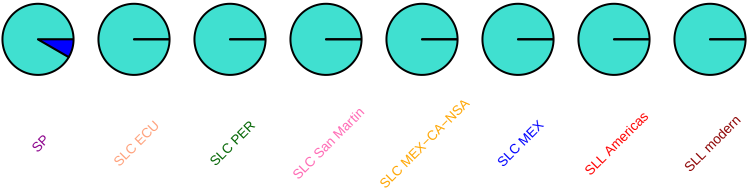

### SlDEAD10, SL2.50ch02, 49307900

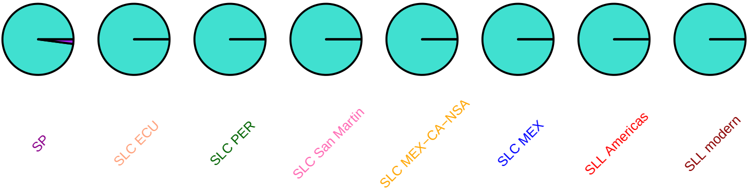

## AY344540, SL2.50ch02, 49612882

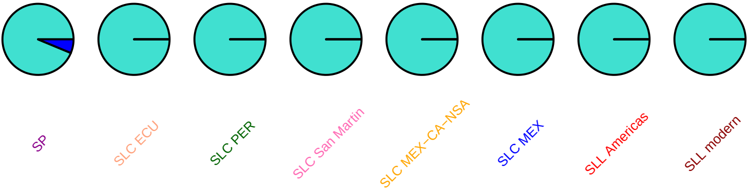

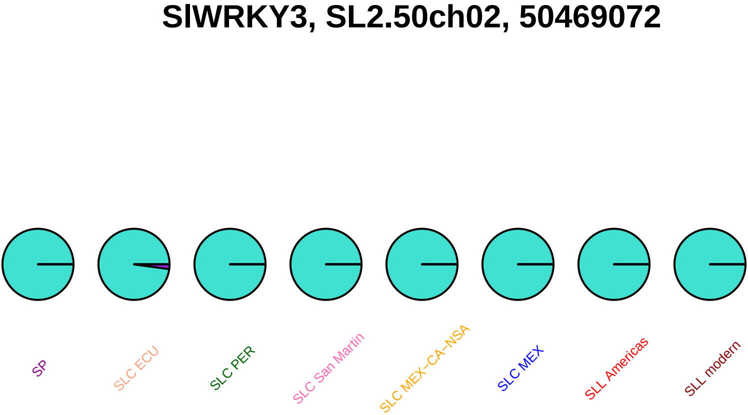
SlWRKY3, SL2.50ch02, 50469072
SP
SLC ECU
SLC PER
SLC San Martin
SLC MEX−CA−NSA
SLC MEX
SLL Americas
SLL modern

# fw2.2, SL2.50ch02, 52253009

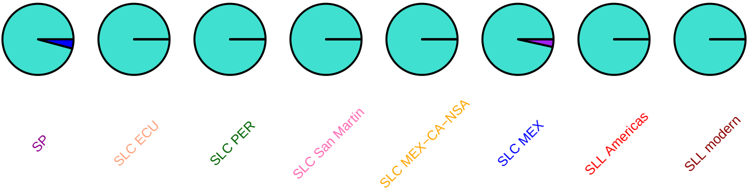

### zep, SL2.50ch02, 52374025

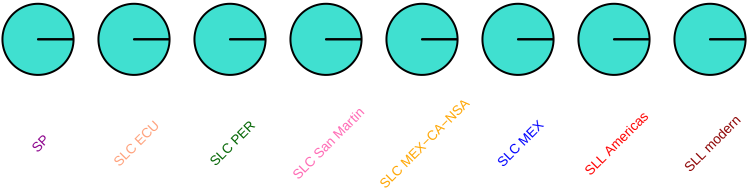

### SlMAPKKK21, SL2.50ch02, 52424941

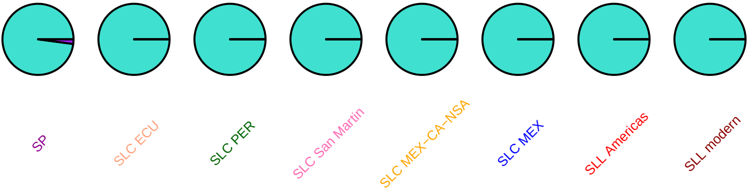

### SlNIP4.1, SL2.50ch02, 52713974

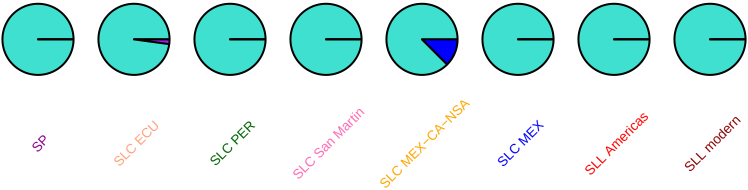

#### SlbHLH016, SL2.50ch02, 53043302

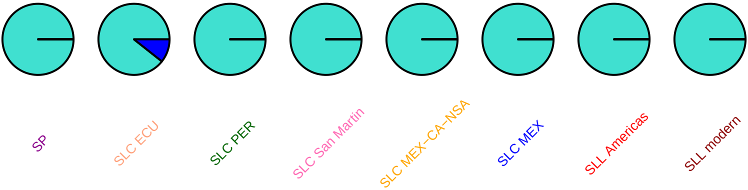

### LeAGP−1, SL2.50ch02, 53762420

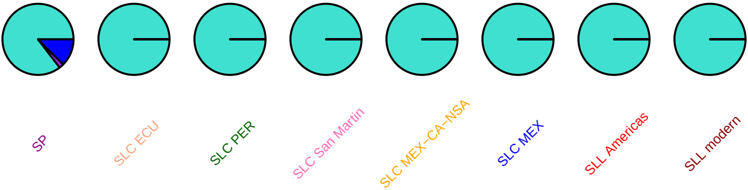

#### SlGH3_4, SL2.50ch02, 53792833

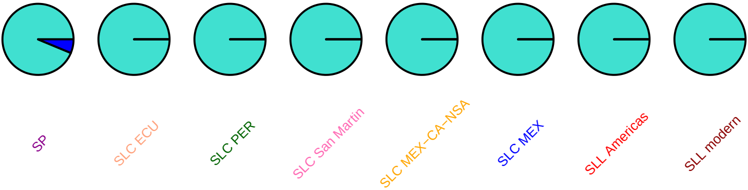

### SlMYB44, SL2.50ch02, 53877910

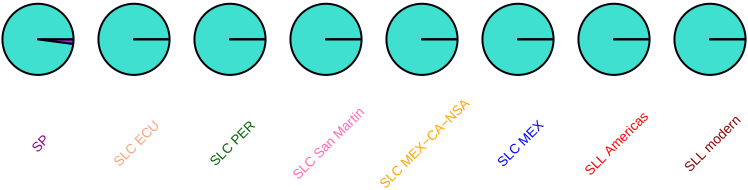

### SlWRKY8, SL2.50ch02, 53998708

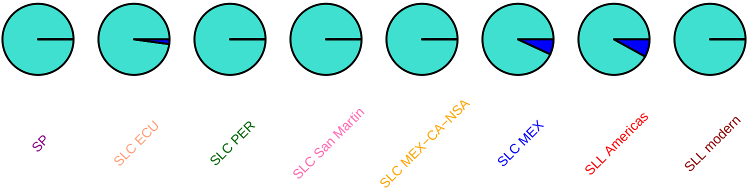

#### SlSTP7, SL2.50ch03, 92027

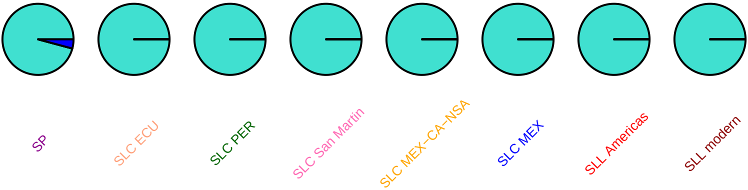

#### SlbHLH018, SL2.50ch03, 207078

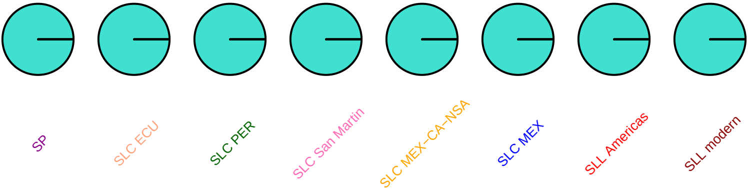

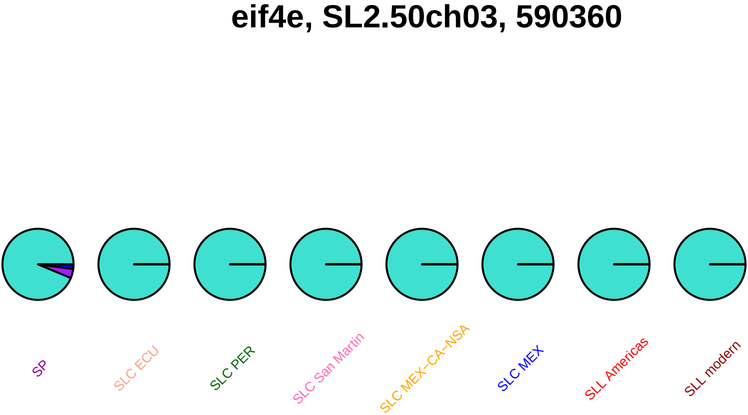
eif4e, SL2.50ch03, 590360
SP
SLC ECU
SLC PER
SLC San Martin
SLC MEX–CA–NSA
SLC MEX
SLL Americas
SLL modern

### eif4e, SL2.50ch03, 592089

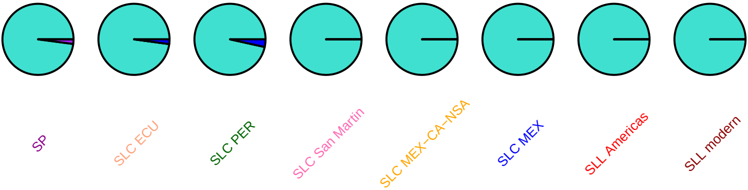

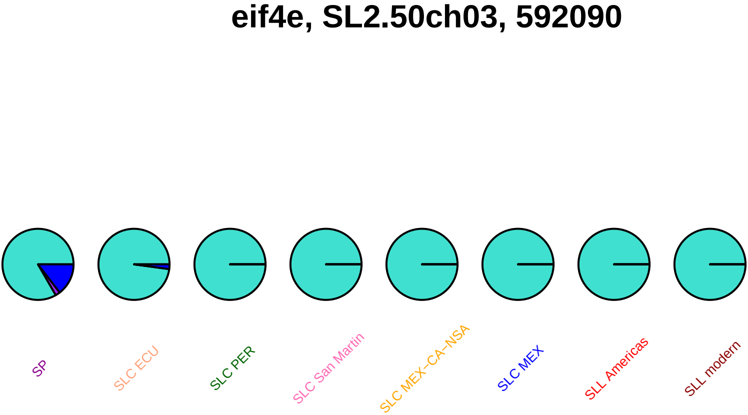
eif4e, SL2.50ch03, 592090
SP
SLC ECU
SLC PER
SLC San Martin
SLC MEX–CA–NSA
SLC MEX
SLL Americas
SLL modern

#### SolycHsfA4a, SL2.50ch03, 679011

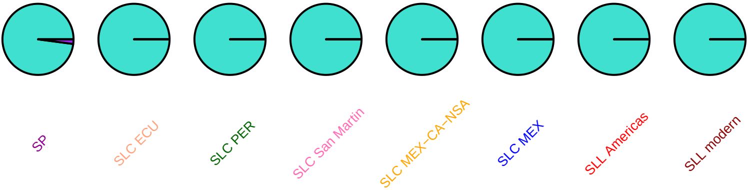

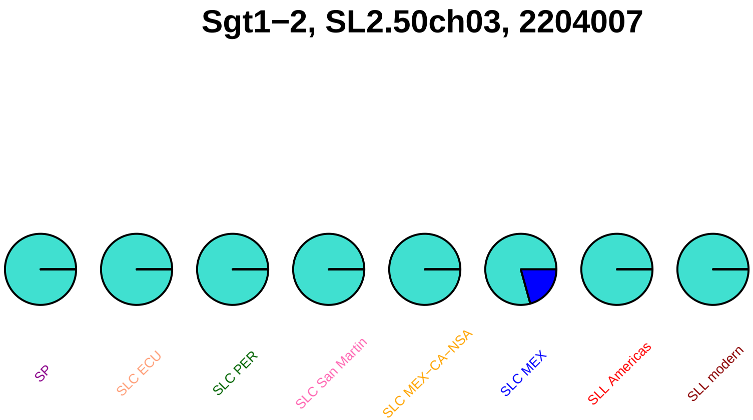
Sgt1−2, SL2.50ch03, 2204007
SP
SLC ECU
SLC PER
SLC San Martin
SLC MEX−CA−NSA
SLC MEX
SLL Americas
SLL modern

### OSP, SL2.50ch03, 2338648

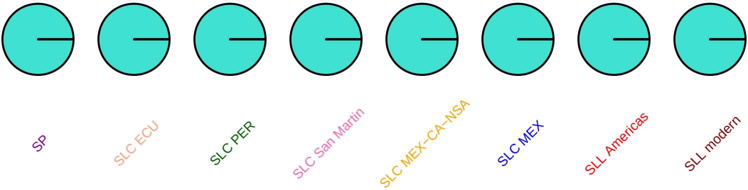

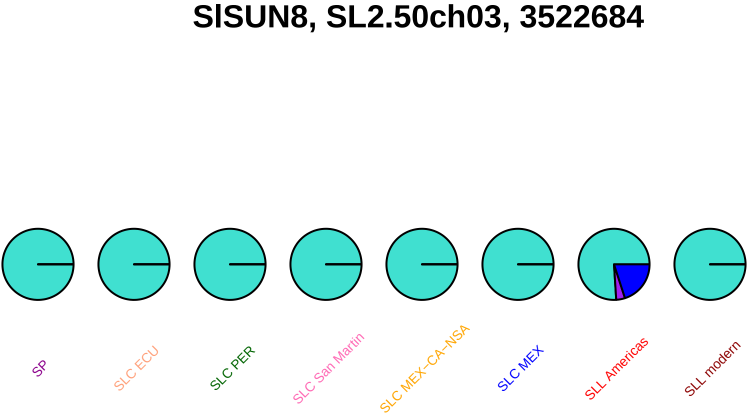
SlSUN8, SL2.50ch03, 3522684
SP
SLC ECU
SLC PER
SLC San Martin
SLC MEX–CA–NSA
SLC MEX
SLL Americas
SLL modern

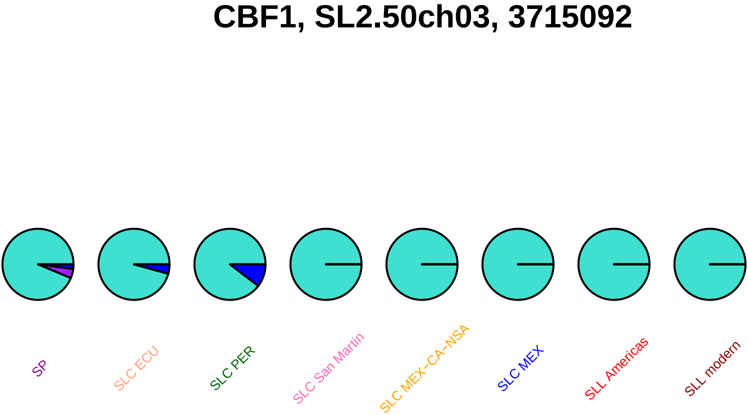
CBF1, SL2.50ch03, 3715092
SP
SLC ECU
SLC PER
SLC San Martin
SLC MEX–CA–NSA
SLC MEX
SLL Americas
SLL modern

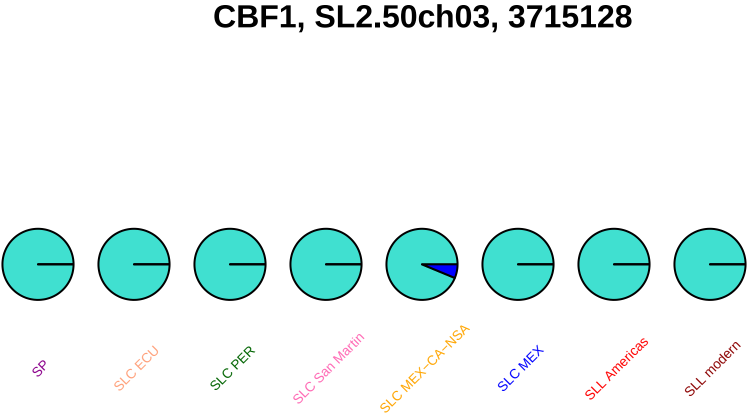
CBF1, SL2.50ch03, 3715128
SP
SLC ECU
SLC PER
SLC San Martin
SLC MEX−CA−NSA
SLC MEX
SLL Americas
SLL modern

#### uref, SL2.50ch03, 3986093

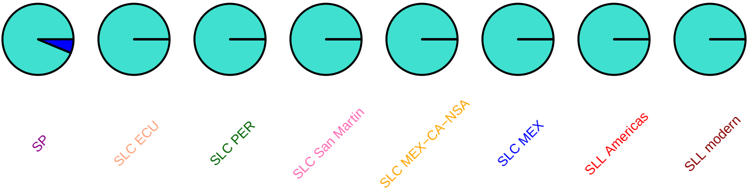

### LeCPK2, SL2.50ch03, 5078653

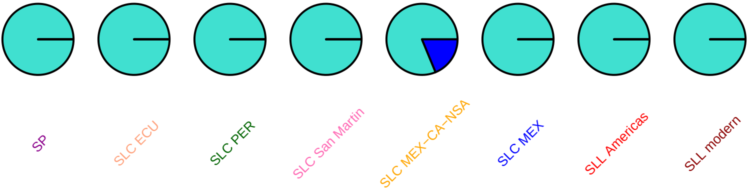

### SlbHLH020, SL2.50ch03, 5701023

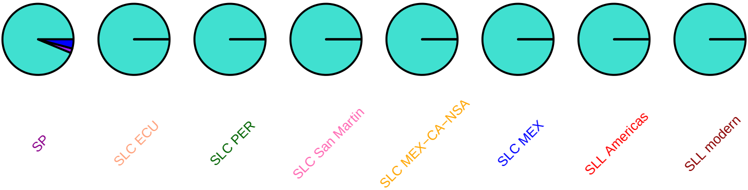

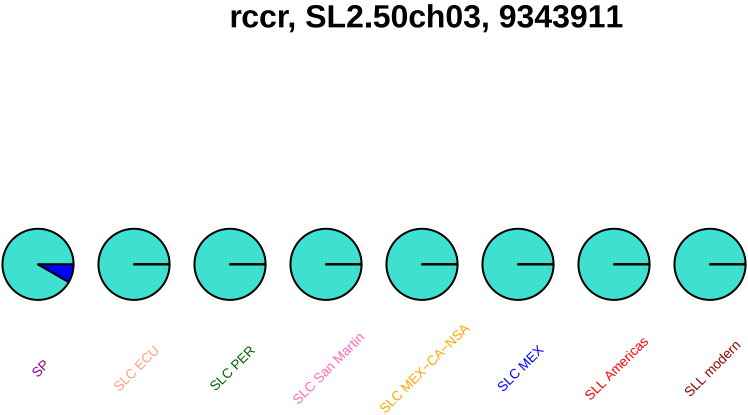
rccr, SL2.50ch03, 9343911
SP
SLC ECU
SLC PER
SLC San Martin
SLC MEX−CA−NSA
SLC MEX
SLL Americas
SLL modern

### rccr, SL2.50ch03, 9344232

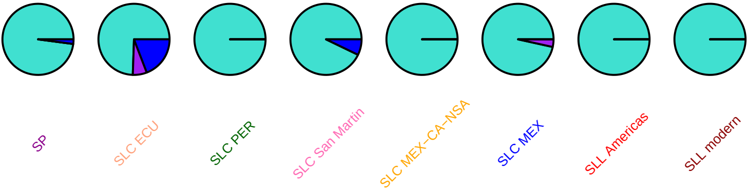

### rccr, SL2.50ch03, 9344409

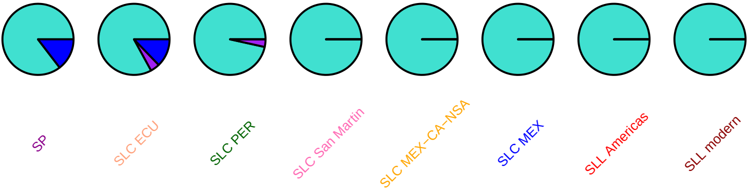

#### imp2, SL2.50ch03, 10780003

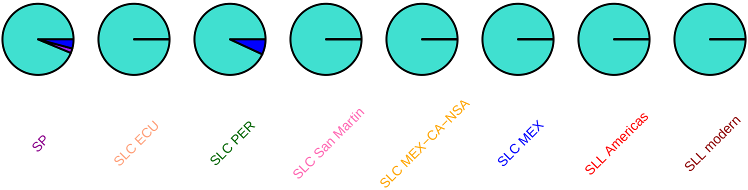

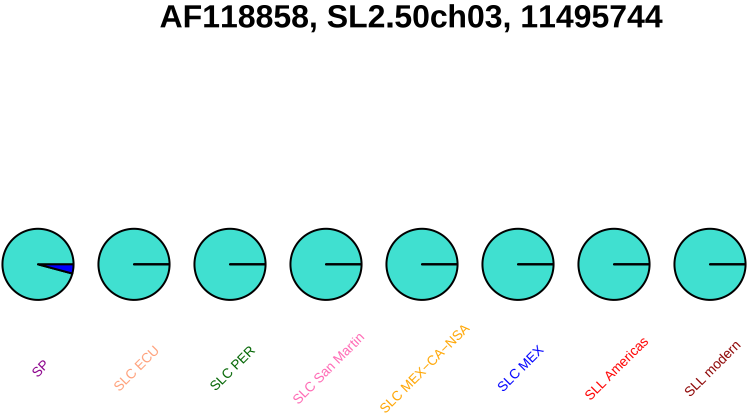

AF118858, SL2.50ch03, 11495744
SP
SLC ECU
SLC PER
SLC San Martin
SLC MEX−CA−NSA
SLC MEX
SLL Americas
SLL modern

### SlC3H26, SL2.50ch03, 35373372

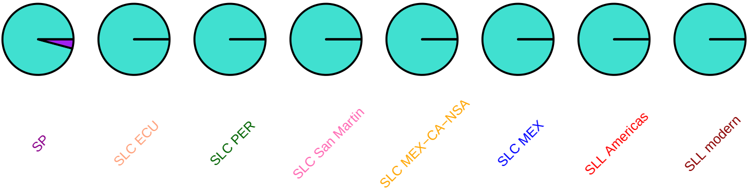

#### SlC3H26, SL2.50ch03, 35373718

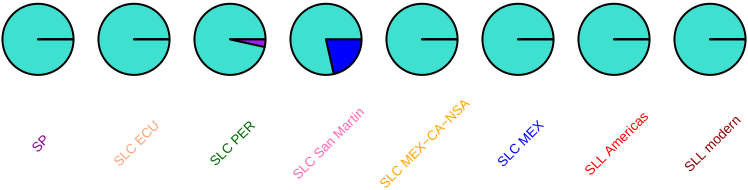

### SlC3H26, SL2.50ch03, 35374471

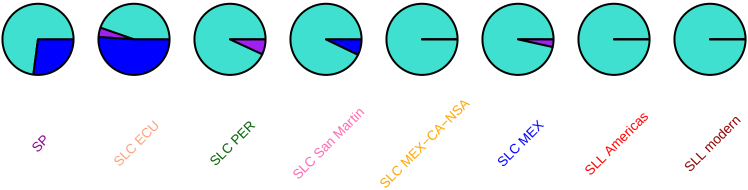

#### SlSTP14, SL2.50ch03, 51230091

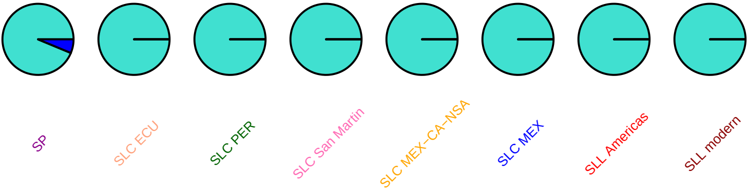

# pk23, SL2.50ch03, 52432076

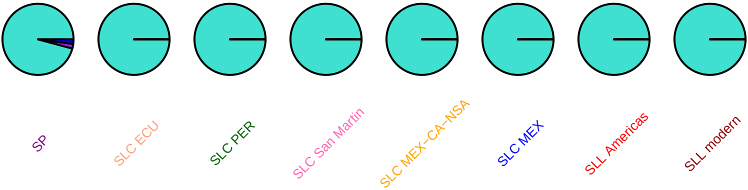

#### SlSAUR36, SL2.50ch03, 52456644

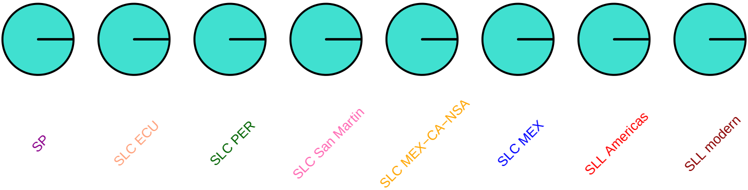

### TIV1, SL2.50ch03, 53853364

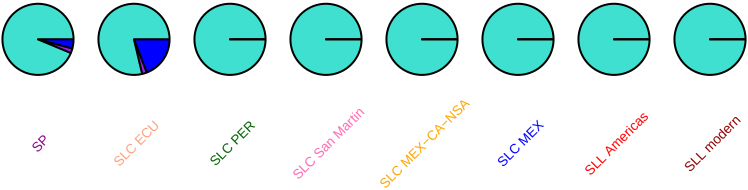

## e86, SL2.50ch03, 57582356

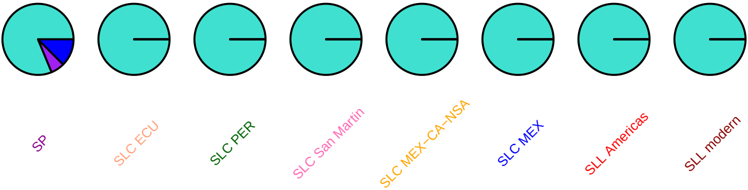

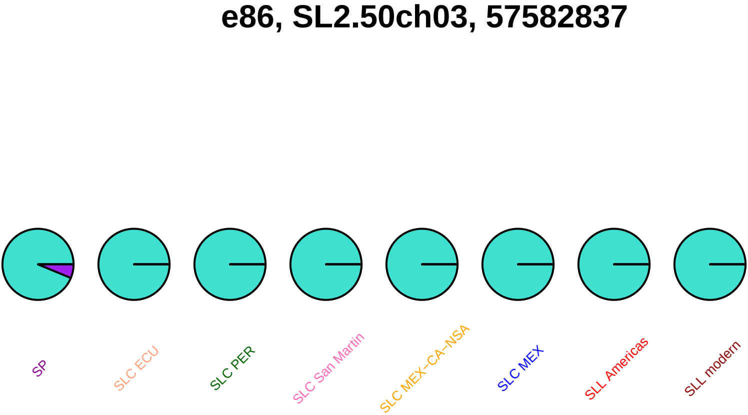
e86, SL2.50ch03, 57582837
SP
SLC ECU
SLC PER
SLC San Martin
SLC MEX−CA−NSA
SLC MEX
SLL Americas
SLL modern

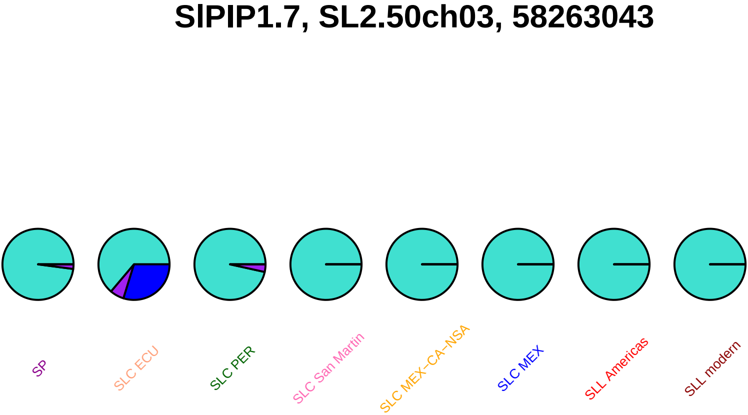
SlPIP1.7, SL2.50ch03, 58263043
SP
SLC ECU
SLC PER
SLC San Martin
SLC MEX–CA–NSA
SLC MEX
SLL Americas
SLL modern

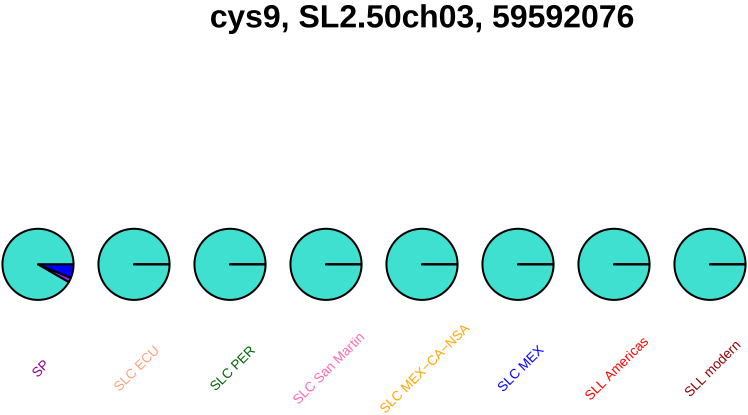

cys9, SL2.50ch03, 59592076
SP
SLC ECU
SLC PER
SLC San Martin
SLC MEX−CA−NSA
SLC MEX
SLL Americas
SLL modern

### TOMMRFRAB, SL2.50ch03, 59614966

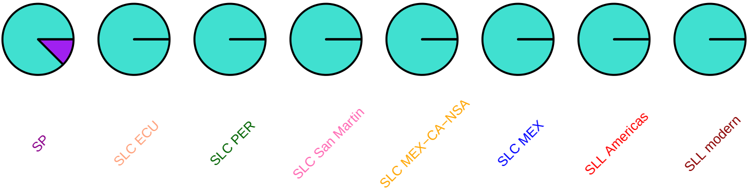

### TOMMRFRAB, SL2.50ch03, 59616567

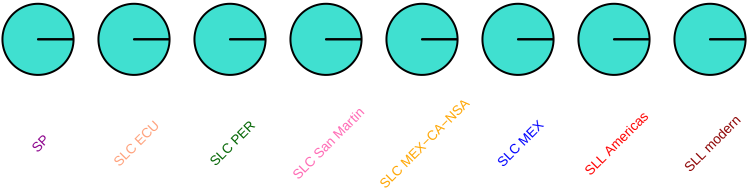

### TOMMRFRAB, SL2.50ch03, 59616615

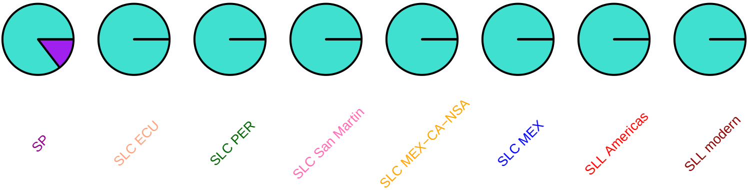

### TOMMRFRAB, SL2.50ch03, 59622949

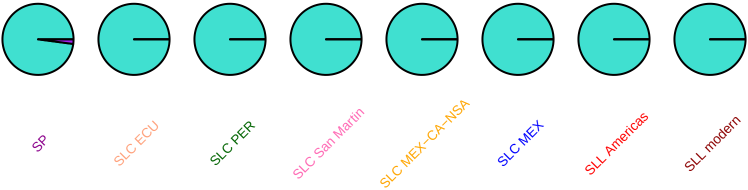

### TOMMRFRAB, SL2.50ch03, 59626511

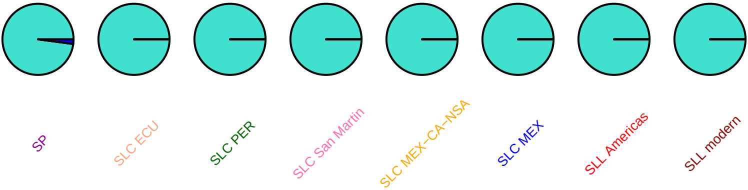

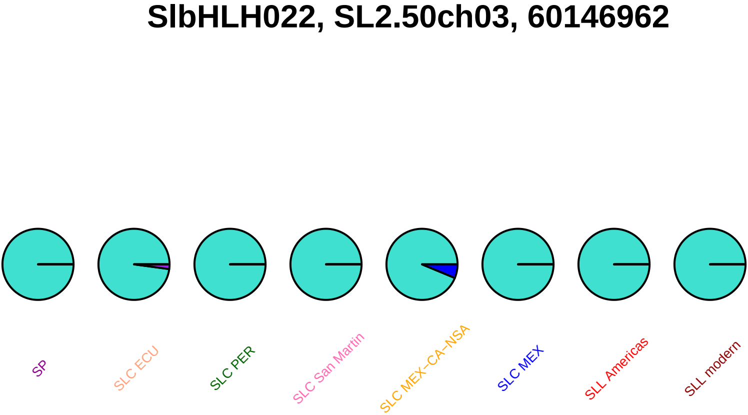
SlbHLH022, SL2.50ch03, 60146962
SP
SLC ECU
SLC PER
SLC San Martin
SLC MEX−CA−NSA
SLC MEX
SLL Americas
SLL modern

### GAD, SL2.50ch03, 60598907

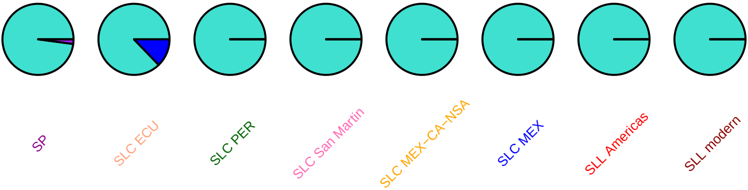

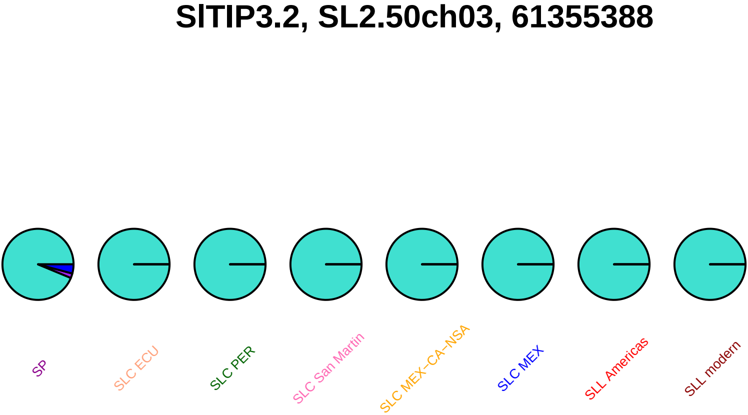
SlTIP3.2, SL2.50ch03, 61355388
SP
SLC ECU
SLC PER
SLC San Martin
SLC MEX−CA−NSA
SLC MEX
SLL Americas
SLL modern

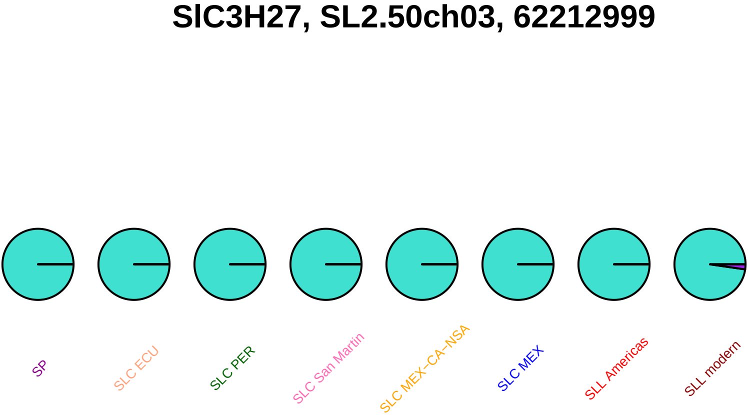
SlC3H27, SL2.50ch03, 62212999
SP
SLC ECU
SLC PER
SLC San Martin
SLC MEX−CA−NSA
SLC MEX
SLL Americas
SLL modern

#### CysEP, SL2.50ch03, 62379908

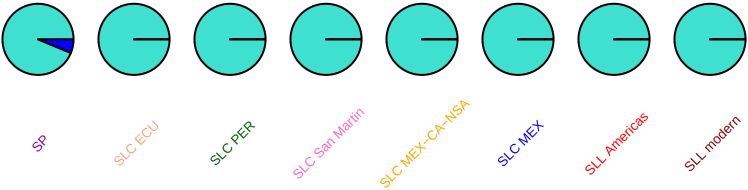

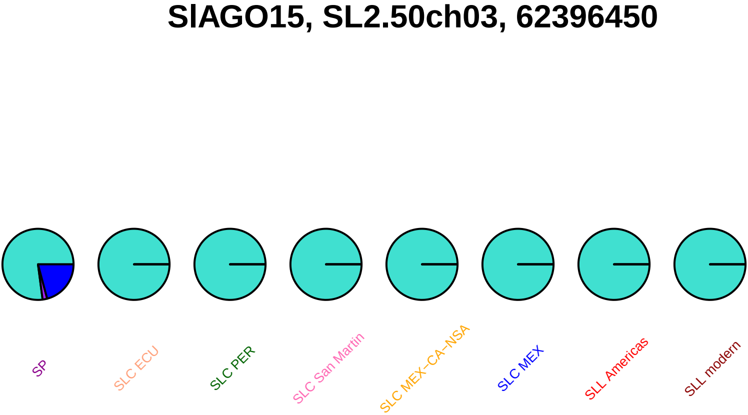
SlAGO15, SL2.50ch03, 62396450
SP
SLC ECU
SLC PER
SLC San Martin
SLC MEX–CA–NSA
SLC MEX
SLL Americas
SLL modern

#### SlAGO15, SL2.50ch03, 62398204

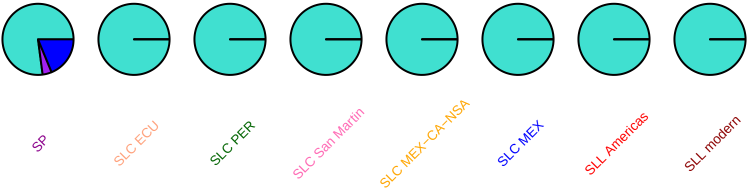

#### SlbHLH082, SL2.50ch03, 64219900

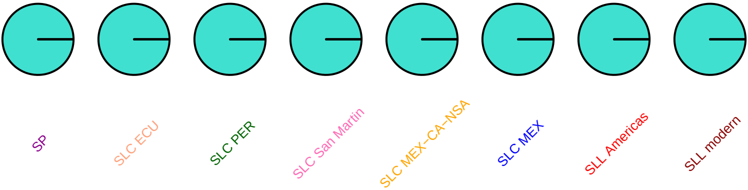

#### SlDEAH9, SL2.50ch03, 64672053

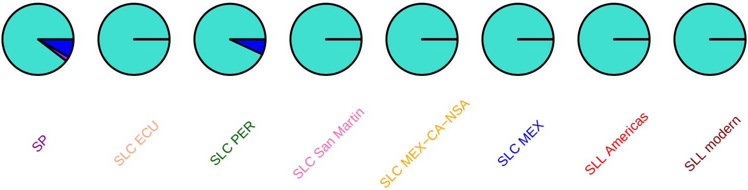

## AY222455, SL2.50ch03, 65341178

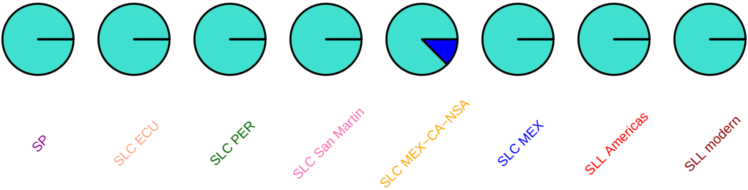

#### SlMYB31, SL2.50ch03, 65610011

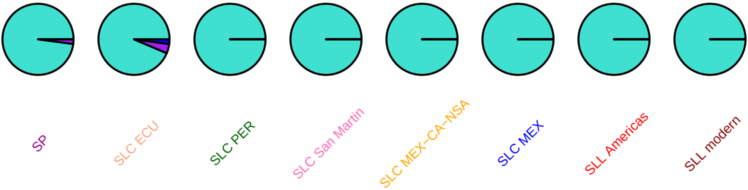

### SlWRKY39, SL2.50ch03, 66140537

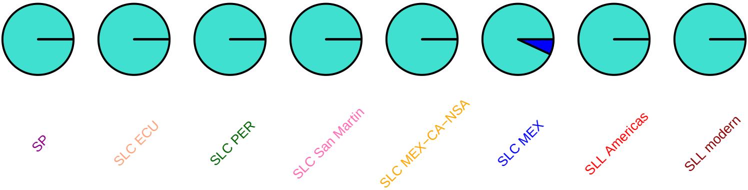

### ccr2, SL2.50ch03, 66179579

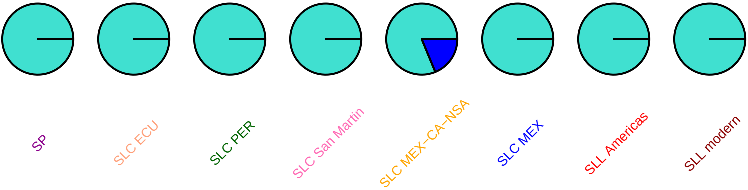

# fa, SL2.50ch03, 67110539

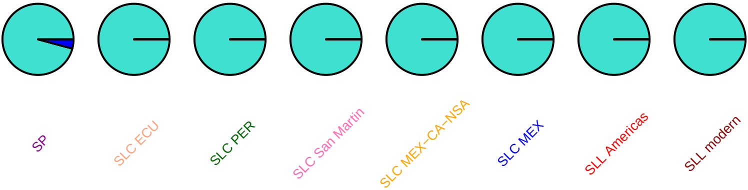

# AF328858, SL2.50ch03, 67581809

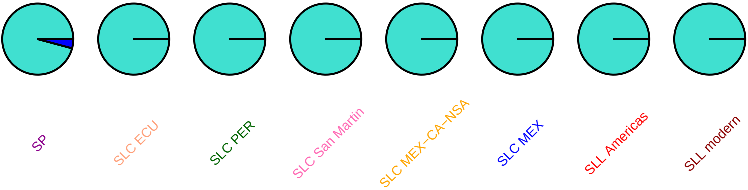

# AF328858, SL2.50ch03, 67583101

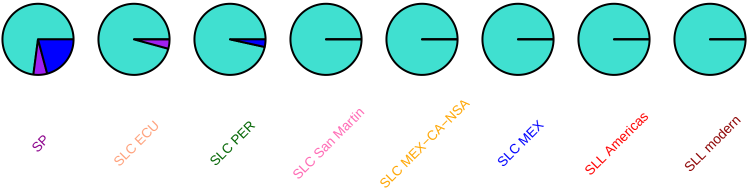

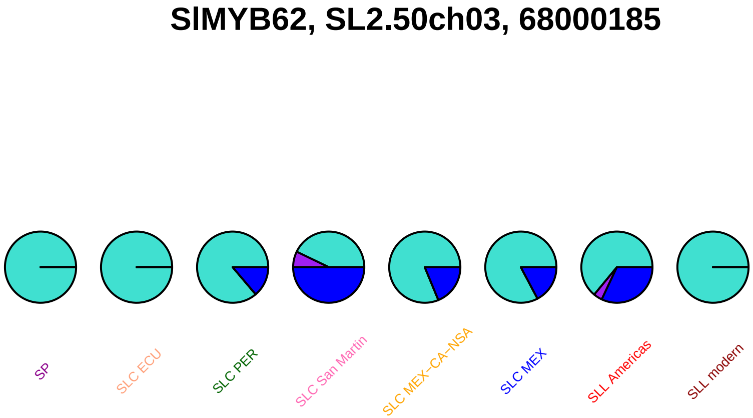
SlMYB62, SL2.50ch03, 68000185
SP
SLC ECU
SLC PER
SLC San Martin
SLC MEX–CA–NSA
SLC MEX
SLL Americas
SLL modern

#### CYP734A7, SL2.50ch03, 68566633

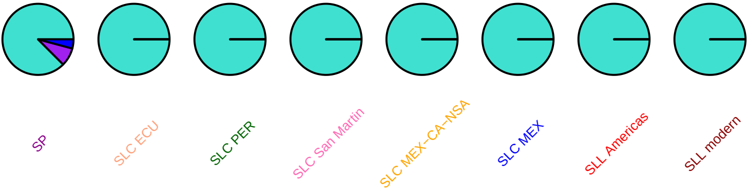

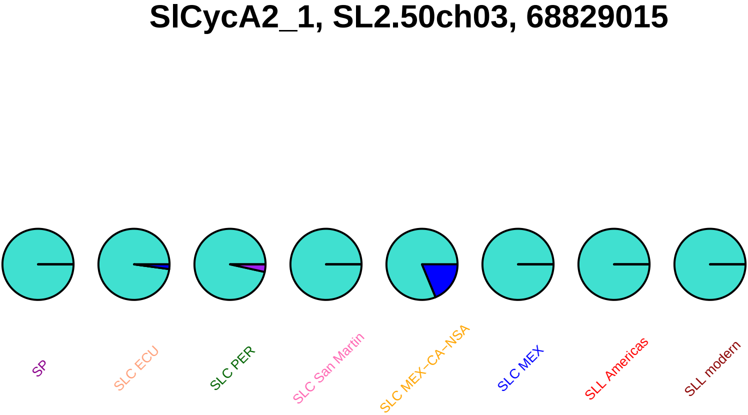

SlCycA2_1, SL2.50ch03, 68829015
SP
SLC ECU
SLC PER
SLC San Martin
SLC MEX−CA−NSA
SLC MEX
SLL Americas
SLL modern

#### SlTIP2.2, SL2.50ch03, 68851468

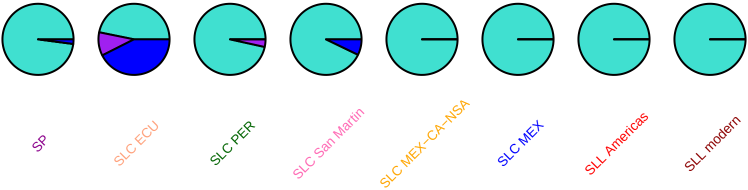

### kpp, SL2.50ch03, 68980852

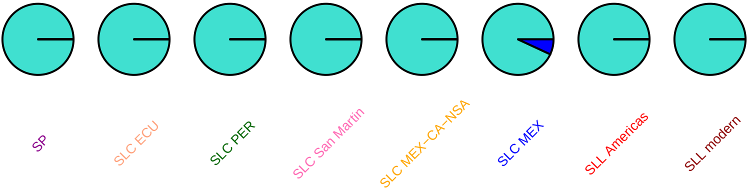

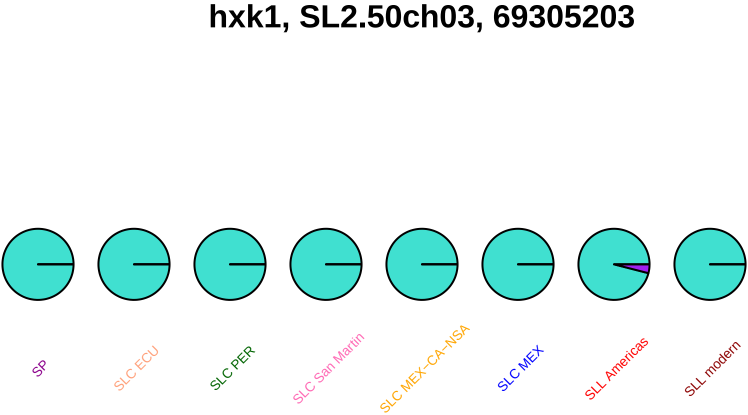

hxk1, SL2.50ch03, 69305203
SP
SLC ECU
SLC PER
SLC San Martin
SLC MEX−CA−NSA
SLC MEX
SLL Americas
SLL modern

### loxd, SL2.50ch03, 70222223

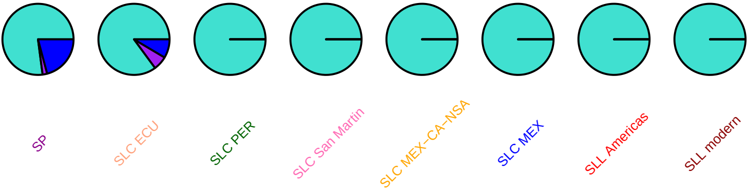

### SlDML4, SL2.50ch03, 70291768

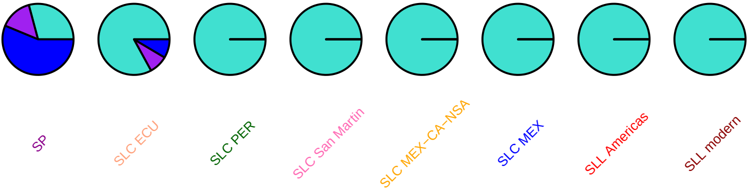

### SlDML4, SL2.50ch03, 70292055

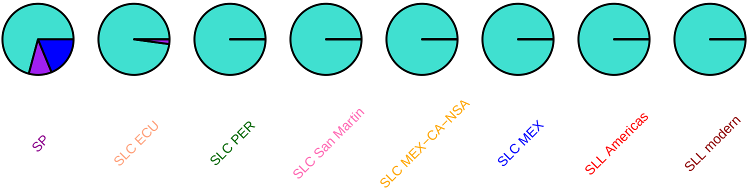

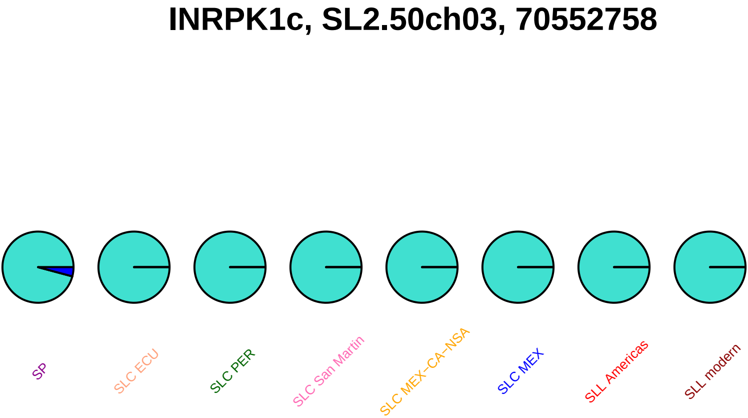

INRPK1c, SL2.50ch03, 70552758
SP
SLC ECU
SLC PER
SLC San Martin
SLC MEX−CA−NSA
SLC MEX
SLL Americas
SLL modern

#### INRPK1c, SL2.50ch03, 70553007

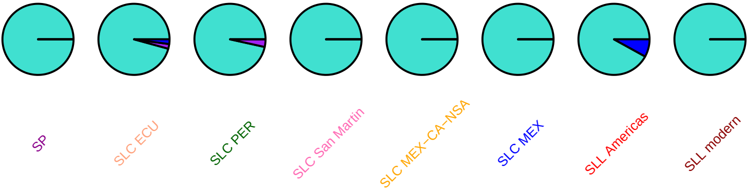

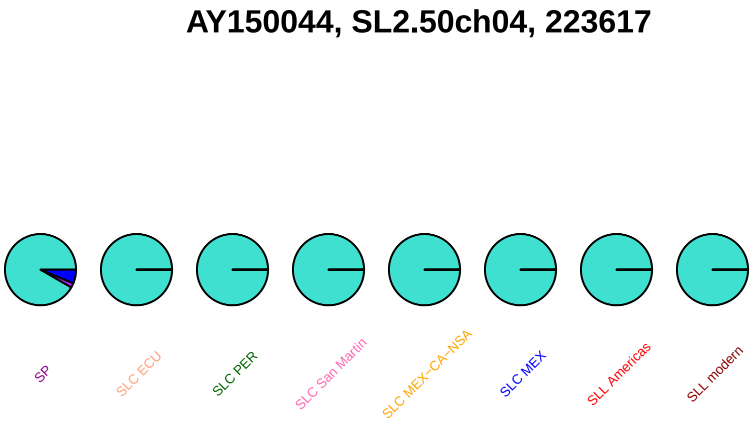
AY150044, SL2.50ch04, 223617
SP
SLC ECU
SLC PER
SLC San Martin
SLC MEX−CA−NSA
SLC MEX
SLL Americas
SLL modern

#### SlbHLH029, SL2.50ch04, 708154

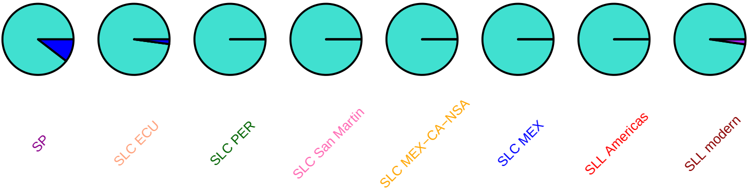

#### Znf, SL2.50ch04, 1149021

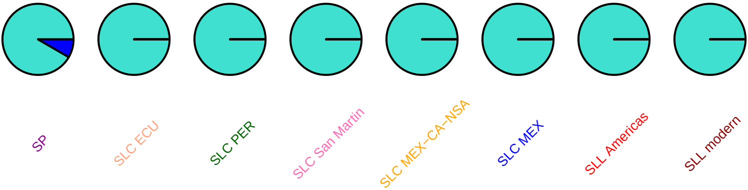

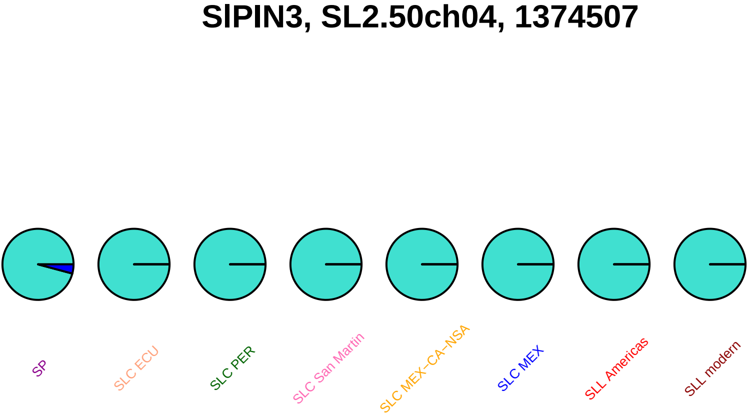
SlPIN3, SL2.50ch04, 1374507
SP
SLC ECU
SLC PER
SLC San Martin
SLC MEX−CA−NSA
SLC MEX
SLL Americas
SLL modern

### help2, SL2.50ch04, 1841212

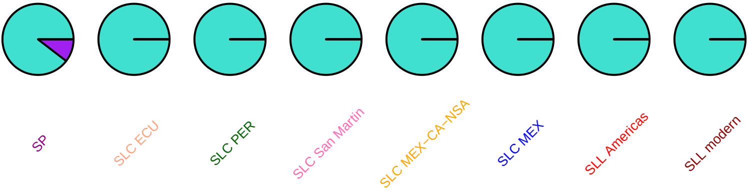

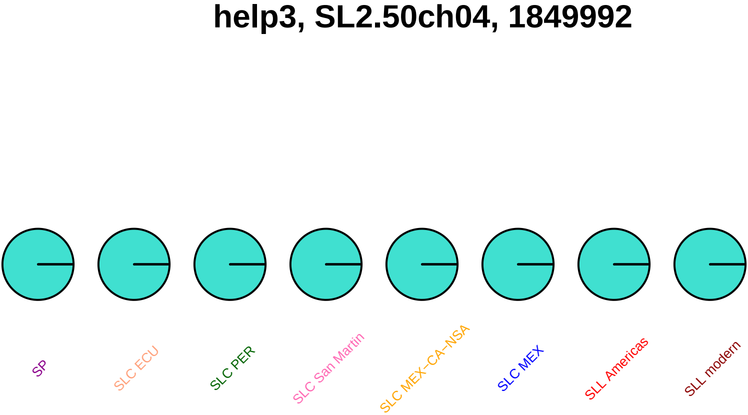
help3, SL2.50ch04, 1849992
SP
SLC ECU
SLC PER
SLC San Martin
SLC MEX−CA−NSA
SLC MEX
SLL Americas
SLL modern

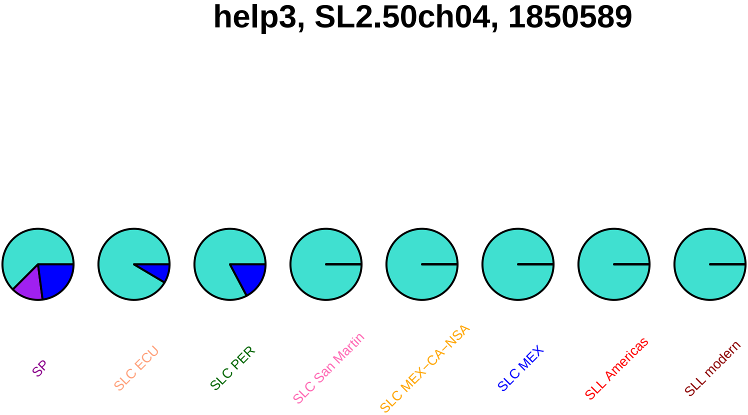
help3, SL2.50ch04, 1850589
SP
SLC ECU
SLC PER
SLC San Martin
SLC MEX−CA−NSA
SLC MEX
SLL Americas
SLL modern

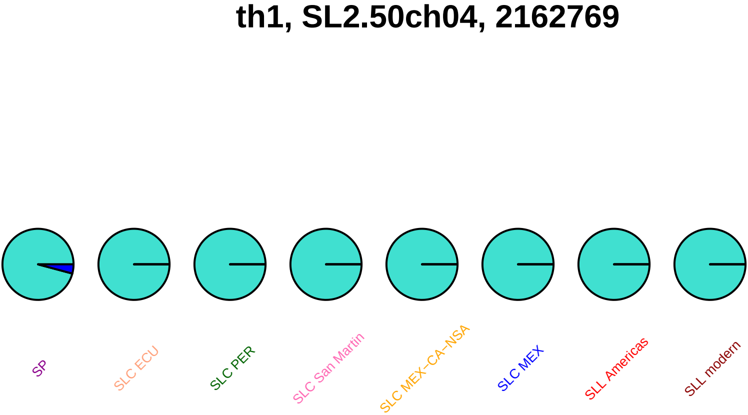
th1, SL2.50ch04, 2162769
SP
SLC ECU
SLC PER
SLC San Martin
SLC MEX−CA−NSA
SLC MEX
SLL Americas
SLL modern

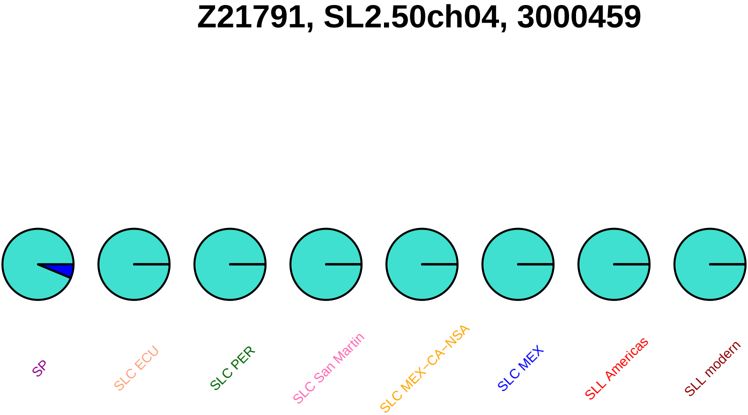
Z21791, SL2.50ch04, 3000459
SP
SLC ECU
SLC PER
SLC San Martin
SLC MEX−CA−NSA
SLC MEX
SLL Americas
SLL modern

## Z21791, SL2.50ch04, 3004950

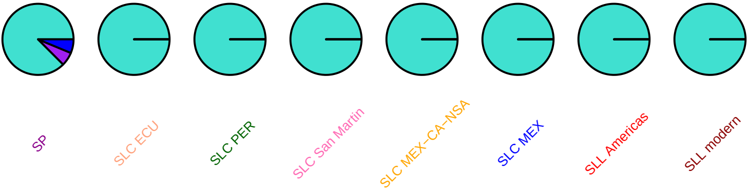

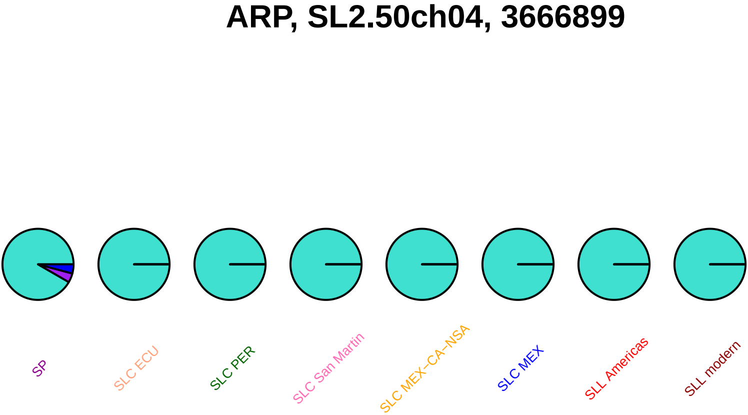
ARP, SL2.50ch04, 3666899
SP
SLC ECU
SLC PER
SLC San Martin
SLC MEX−CA−NSA
SLC MEX
SLL Americas
SLL modern

## er21, SL2.50ch04, 3896881

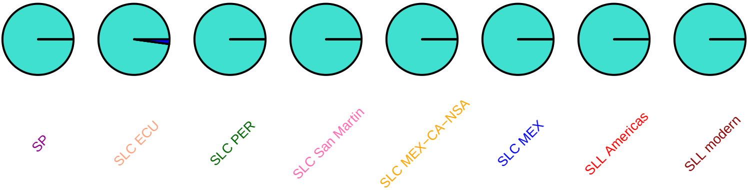

### SlMYB20, SL2.50ch04, 4710760

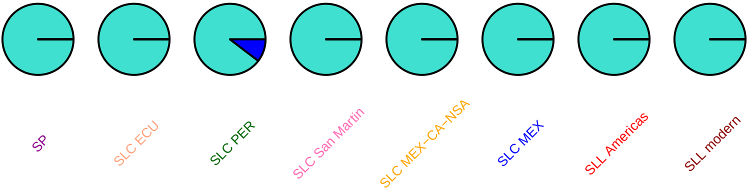

### Sl−ERF_C_2, SL2.50ch04, 4802478

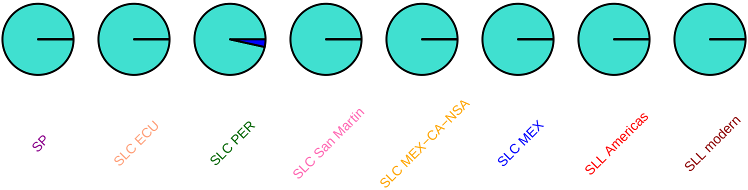

### SlMAPKKK32, SL2.50ch04, 4951629

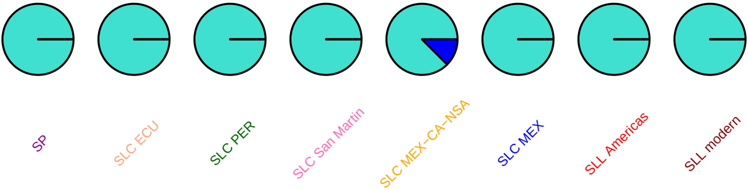

## X55684, SL2.50ch04, 5652381

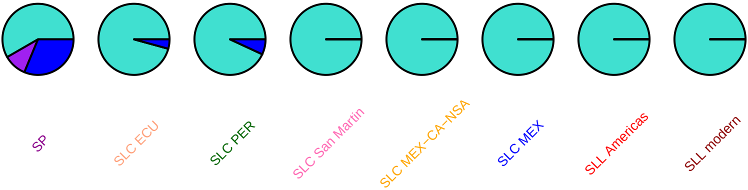

#### ide, SL2.50ch04, 5963991

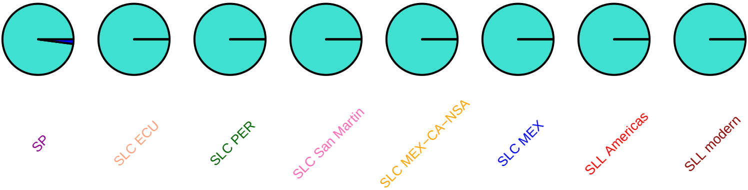

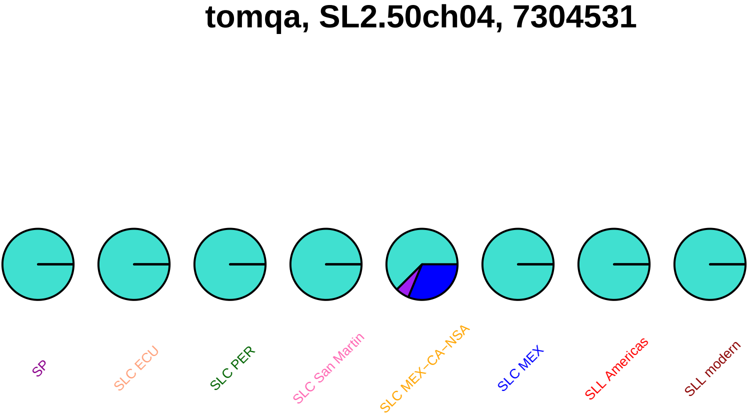
tomqa, SL2.50ch04, 7304531
SP
SLC ECU
SLC PER
SLC San Martin
SLC MEX−CA−NSA
SLC MEX
SLL Americas
SLL modern

### SlSUN12, SL2.50ch04, 7307999

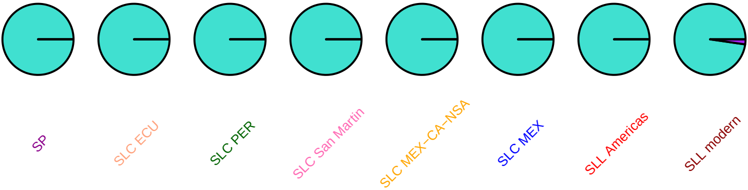

### sgs3, SL2.50ch04, 24325592

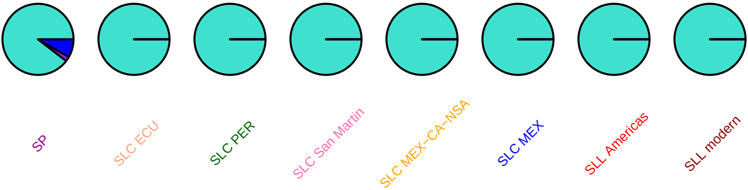

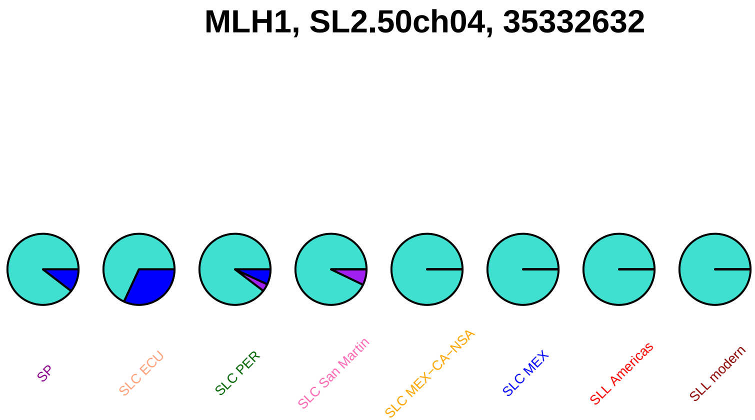
MLH1, SL2.50ch04, 35332632
SP
SLC ECU
SLC PER
SLC San Martin
SLC MEX−CA−NSA
SLC MEX
SLL Americas
SLL modern

### ein57, SL2.50ch04, 39196747

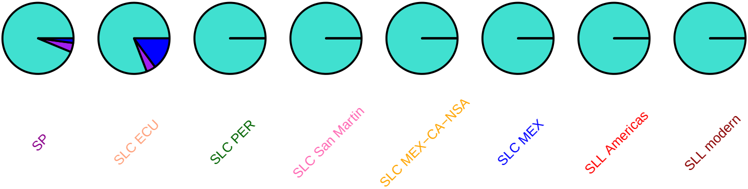

# AY425708, SL2.50ch04, 41623880

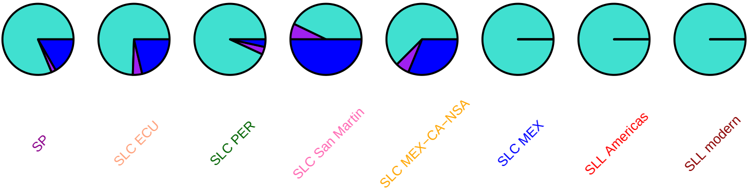

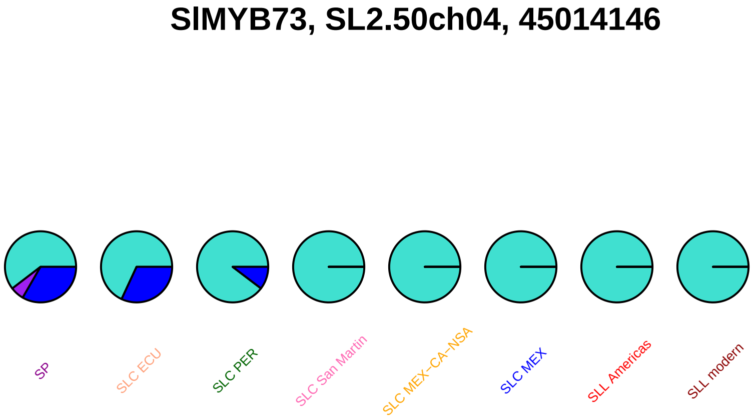
SlMYB73, SL2.50ch04, 45014146
SP
SLC ECU
SLC PER
SLC San Martin
SLC MEX–CA–NSA
SLC MEX
SLL Americas
SLL modern

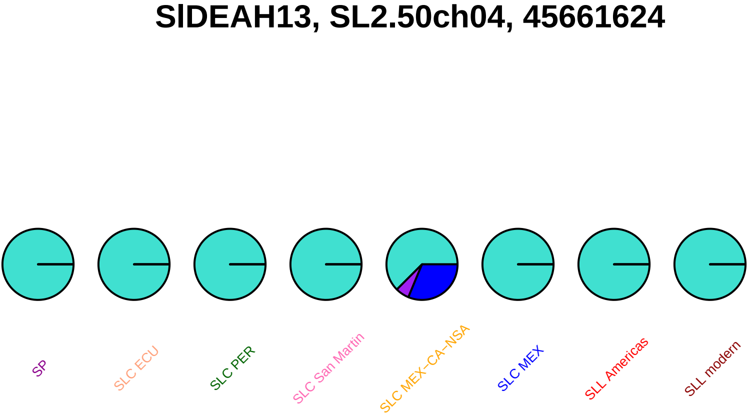
SlDEAH13, SL2.50ch04, 45661624
SP
SLC ECU
SLC PER
SLC San Martin
SLC MEX−CA−NSA
SLC MEX
SLL Americas
SLL modern

### SlWRKY69, SL2.50ch04, 46148530

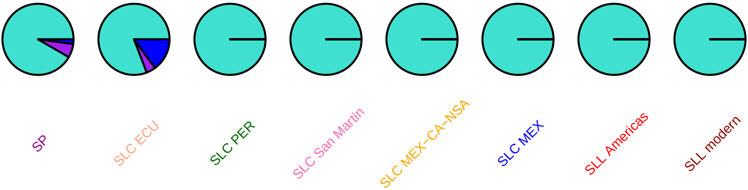

# X63560, SL2.50ch04, 51171303

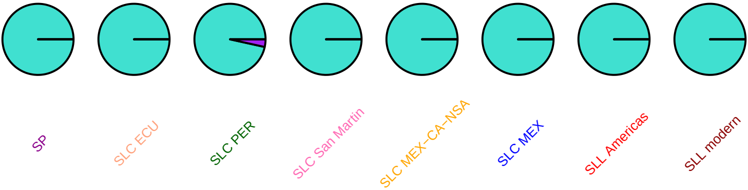

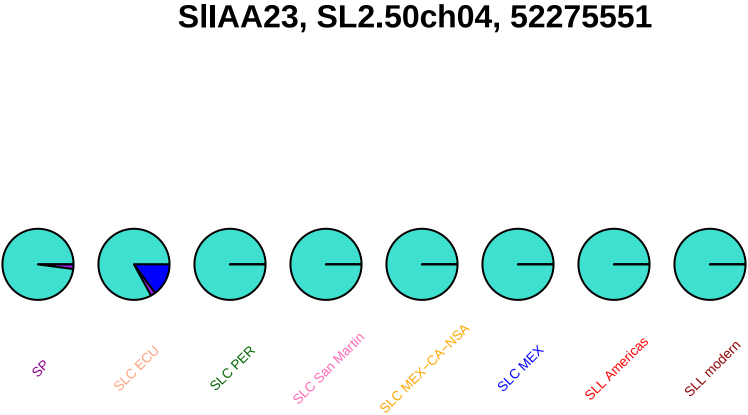

SlIAA23, SL2.50ch04, 52275551
SP
SLC ECU
SLC PER
SLC San Martin
SLC MEX−CA−NSA
SLC MEX
SLL Americas
SLL modern

### SlIAA23, SL2.50ch04, 52275578

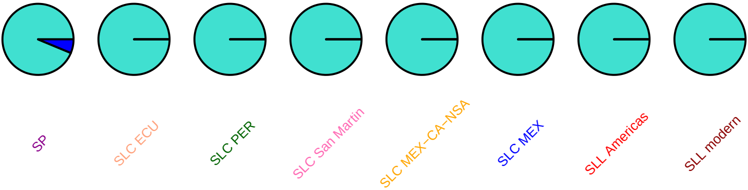

## ao, SL2.50ch04, 53052212

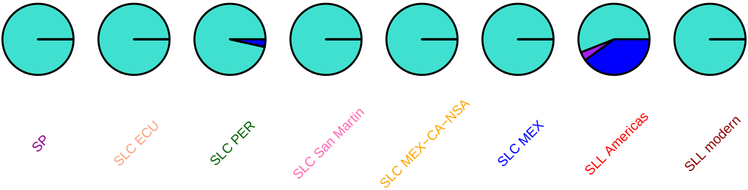

#### inps, SL2.50ch04, 53182836

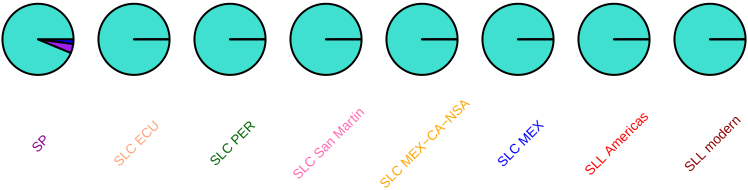

### TOMTRALTBO, SL2.50ch04, 53795864

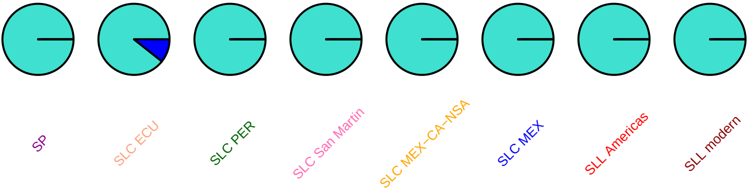

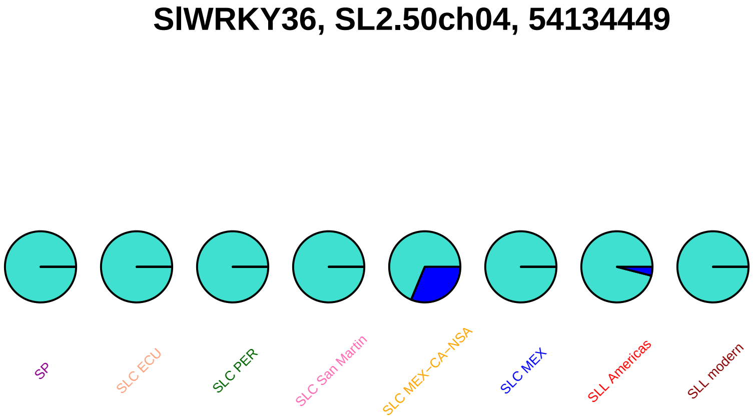
SlWRKY36, SL2.50ch04, 54134449
SP
SLC ECU
SLC PER
SLC San Martin
SLC MEX−CA−NSA
SLC MEX
SLL Americas
SLL modern

**SlC3H33, SL2.50ch04, 55109708**

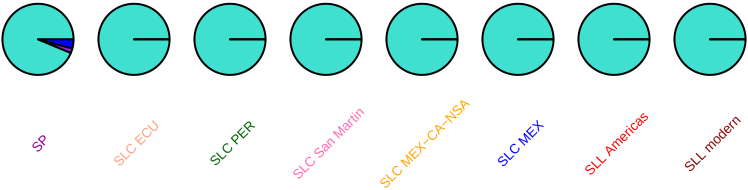

### cas1, SL2.50ch04, 57825074

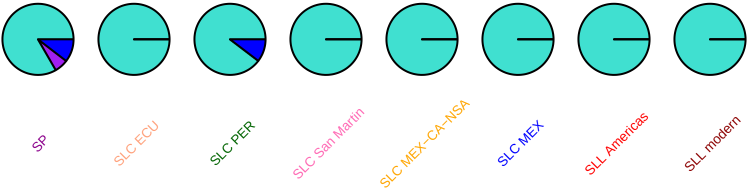

#### SlCycH1_2, SL2.50ch04, 59835918

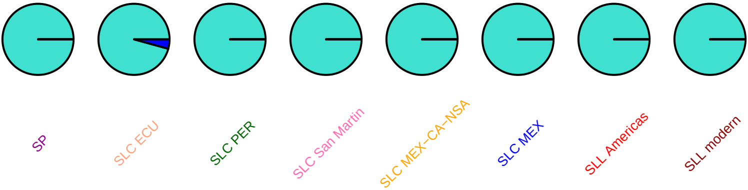

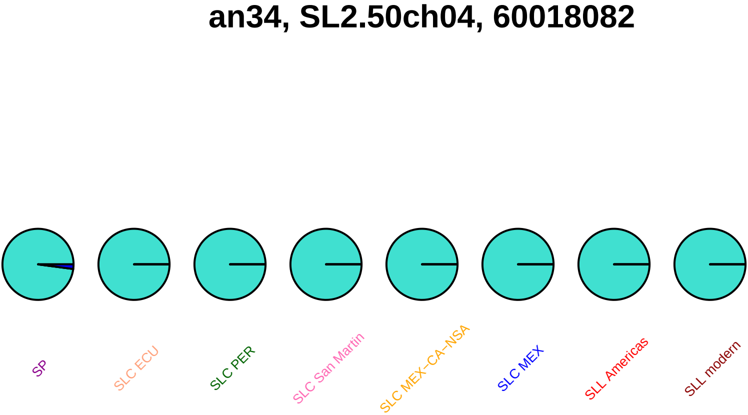
an34, SL2.50ch04, 60018082
SP
SLC ECU
SLC PER
SLC San Martin
SLC MEX–CA–NSA
SLC MEX
SLL Americas
SLL modern

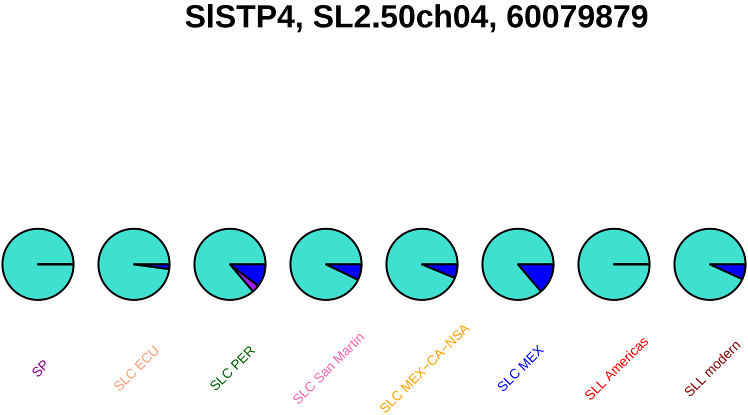
SlSTP4, SL2.50ch04, 60079879
SP
SLC ECU
SLC PER
SLC San Martin
SLC MEX−CA−NSA
SLC MEX
SLL Americas
SLL modern

### SlCycD5_1, SL2.50ch04, 61005485

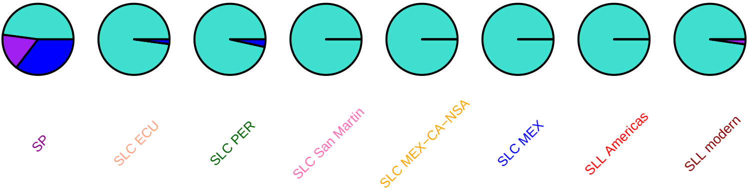

### gpi, SL2.50ch04, 61045738

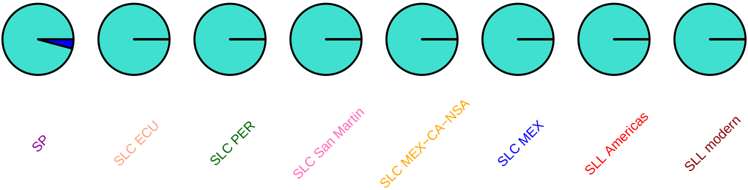

### SlMAPKKK34, SL2.50ch04, 61344290

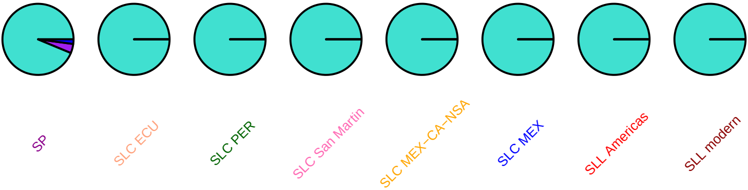

### SlMAPKKK34, SL2.50ch04, 61346327

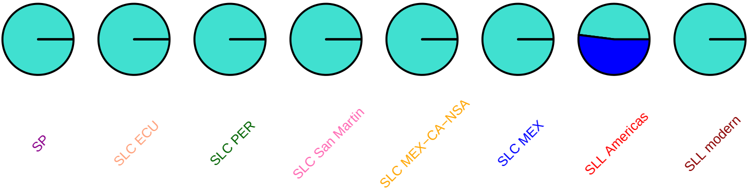

### SlMAPKKK34, SL2.50ch04, 61346328

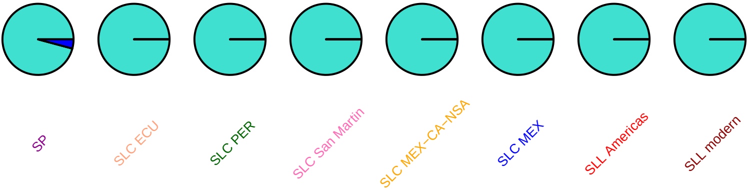

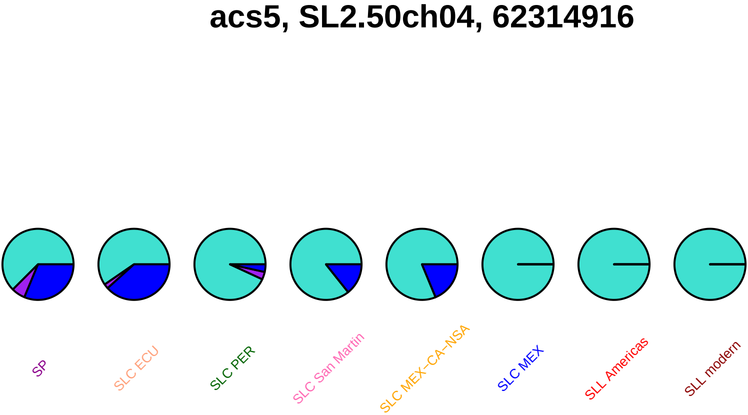
acs5, SL2.50ch04, 62314916
SP
SLC ECU
SLC PER
SLC San Martin
SLC MEX−CA−NSA
SLC MEX
SLL Americas
SLL modern

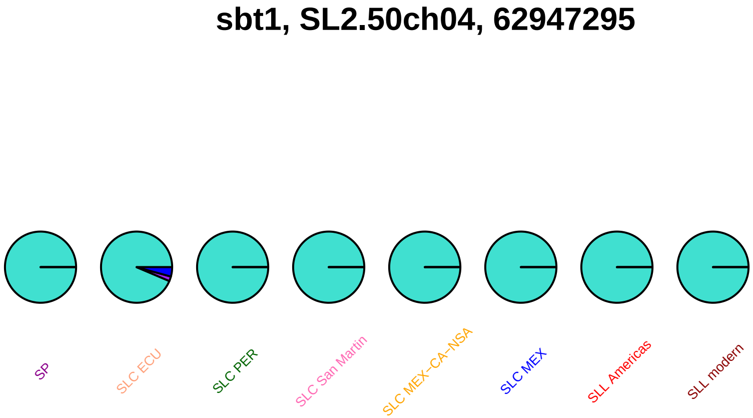
sbt1, SL2.50ch04, 62947295
SP
SLC ECU
SLC PER
SLC San Martin
SLC MEX−CA−NSA
SLC MEX
SLL Americas
SLL modern

### sbt1, SL2.50ch04, 62947343

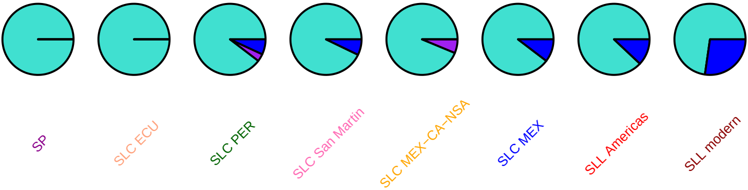

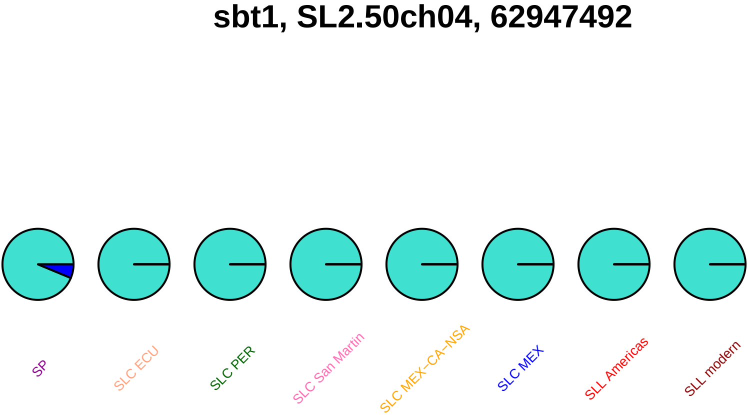
sbt1, SL2.50ch04, 62947492
SP
SLC ECU
SLC PER
SLC San Martin
SLC MEX–CA–NSA
SLC MEX
SLL Americas
SLL modern

### cycd3c3, SL2.50ch04, 63202824

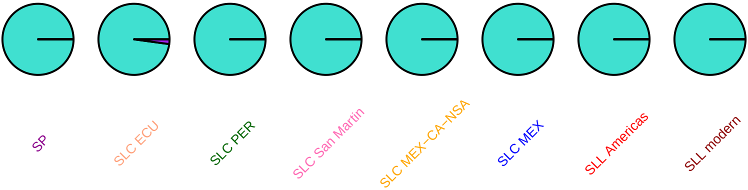

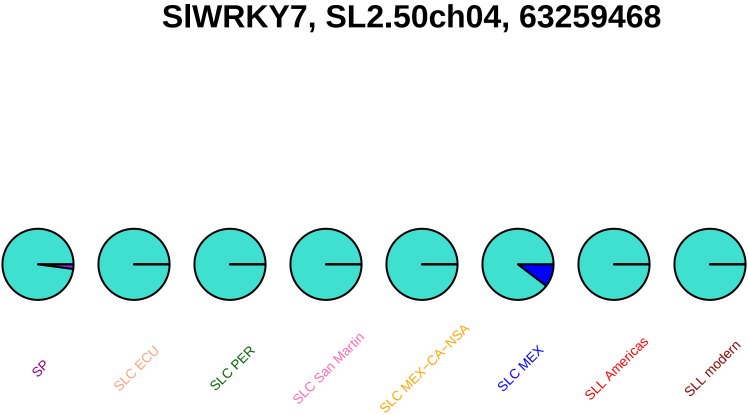
SlWRKY7, SL2.50ch04, 63259468
SP
SLC ECU
SLC PER
SLC San Martin
SLC MEX–CA–NSA
SLC MEX
SLL Americas
SLL modern

### SlWRKY7, SL2.50ch04, 63260653

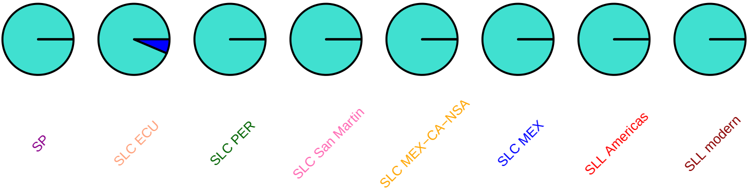

### aos, SL2.50ch04, 64094814

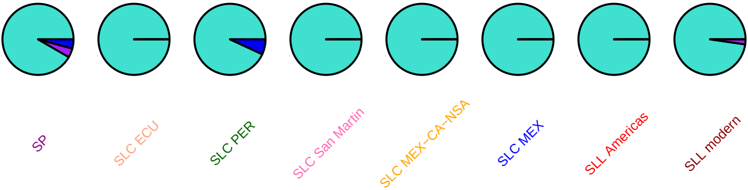

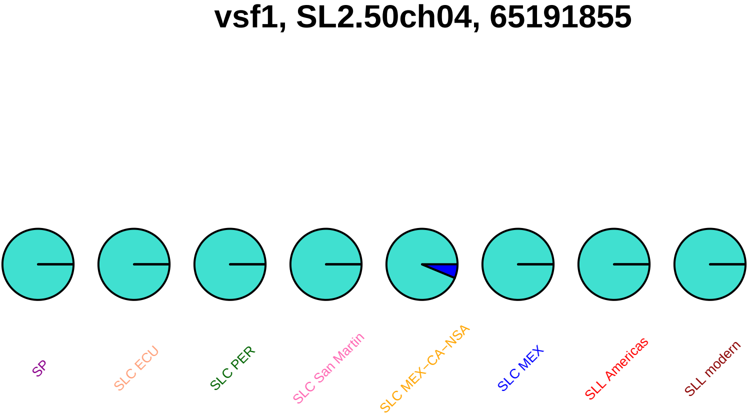
vsf1, SL2.50ch04, 65191855
SP
SLC ECU
SLC PER
SLC San Martin
SLC MEX−CA−NSA
SLC MEX
SLL Americas
SLL modern

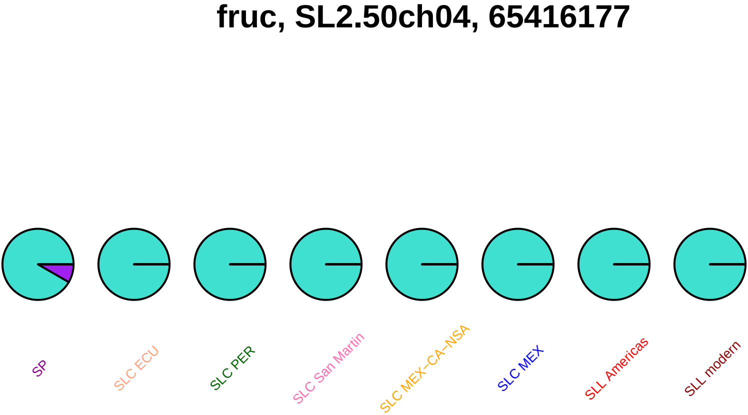

fruc, SL2.50ch04, 65416177
SP
SLC ECU
SLC PER
SLC San Martin
SLC MEX−CA−NSA
SLC MEX
SLL Americas
SLL modern

#### SlTMT2, SL2.50ch04, 66291053

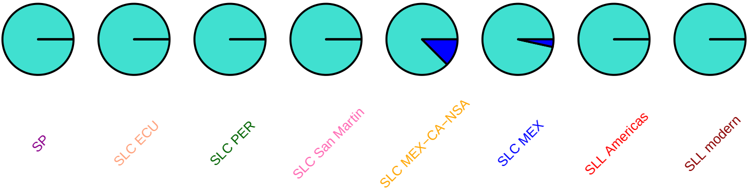

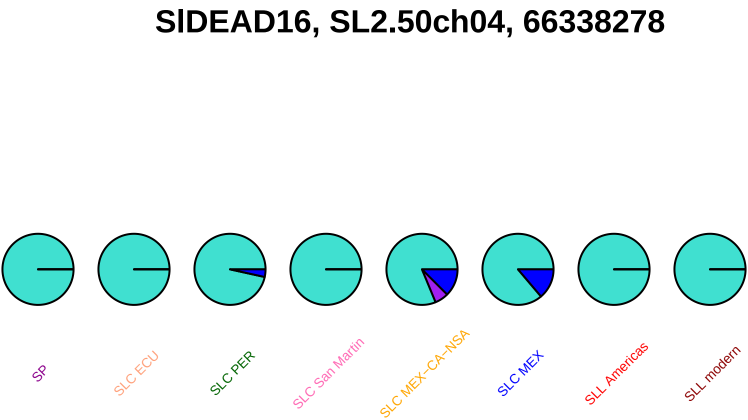
SlDEAD16, SL2.50ch04, 66338278
SP
SLC ECU
SLC PER
SLC San Martin
SLC MEX−CA−NSA
SLC MEX
SLL Americas
SLL modern

# AB211526, SL2.50ch05, 197063

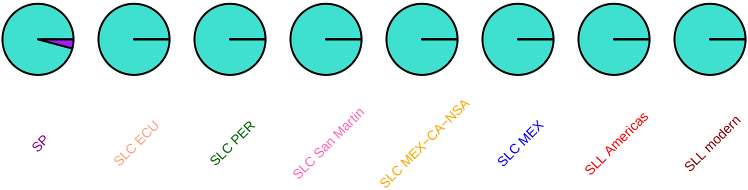

#### SlbHLH084, SL2.50ch05, 242353

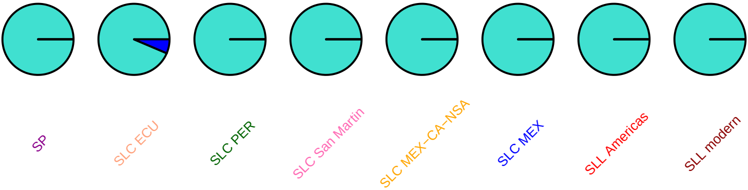

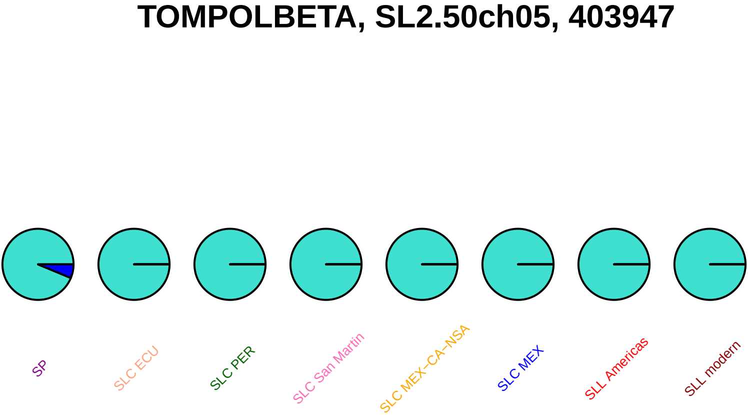
TOMPOLBETA, SL2.50ch05, 403947
SP
SLC ECU
SLC PER
SLC San Martin
SLC MEX−CA−NSA
SLC MEX
SLL Americas
SLL modern

**SlDEAD17, SL2.50ch05, 839314**

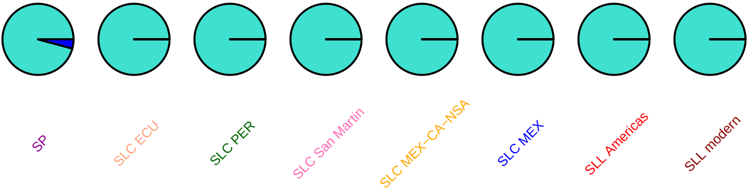

### LeCEP1, SL2.50ch05, 853202

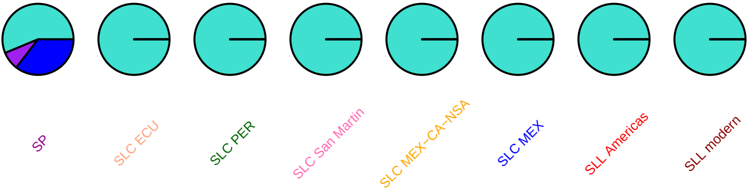

#### SlMYB76, SL2.50ch05, 2635240

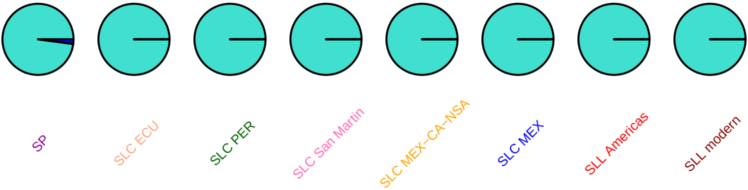

#### SlIAA32, SL2.50ch05, 3080077

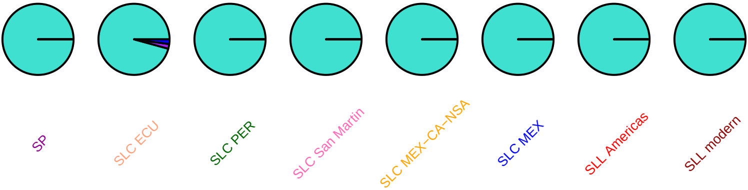

### cab4a, SL2.50ch05, 3329429

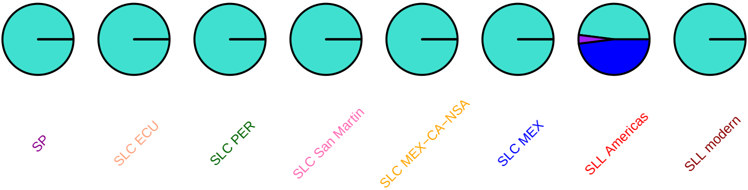

### rin, SL2.50ch05, 5230699

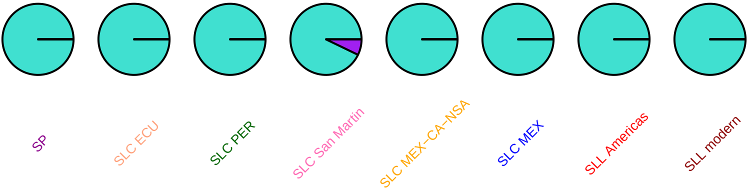

### ddtfr7, SL2.50ch05, 5787633

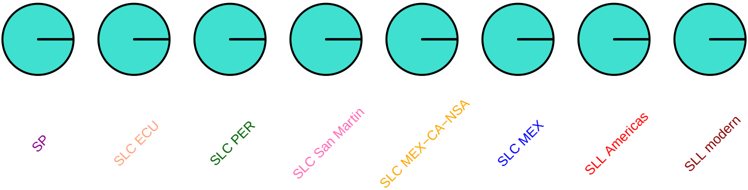

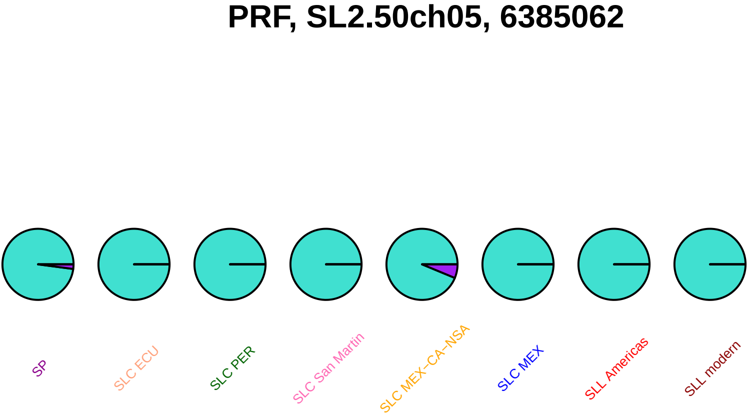
PRF, SL2.50ch05, 6385062
SP
SLC ECU
SLC PER
SLC San Martin
SLC MEX−CA−NSA
SLC MEX
SLL Americas
SLL modern

### Fen, SL2.50ch05, 6387700

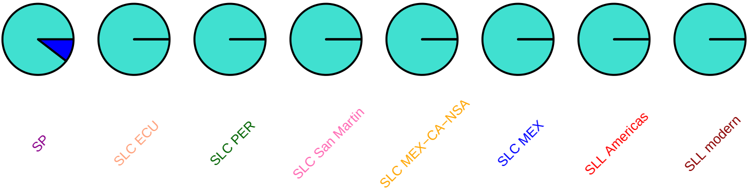

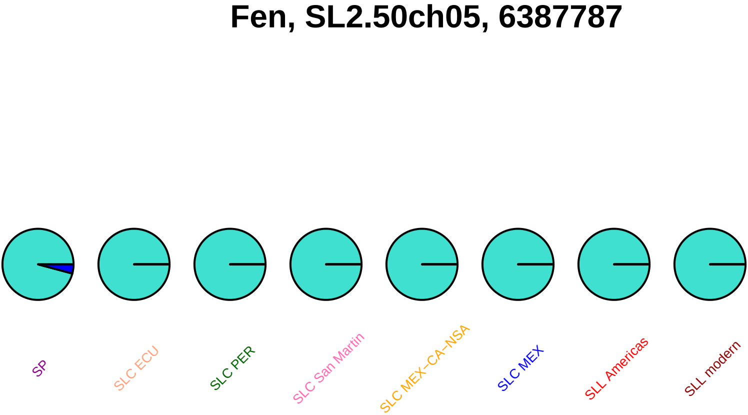
Fen, SL2.50ch05, 6387787
SP
SLC ECU
SLC PER
SLC San Martin
SLC MEX−CA−NSA
SLC MEX
SLL Americas
SLL modern

### Fen, SL2.50ch05, 6387893

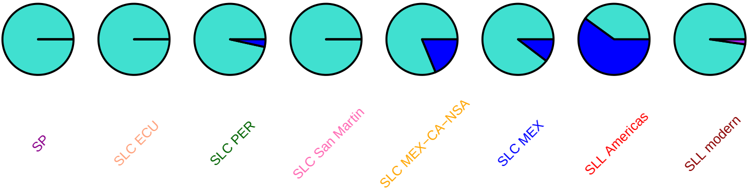

### Fen, SL2.50ch05, 6387898

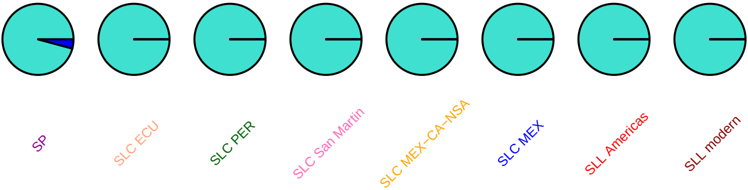

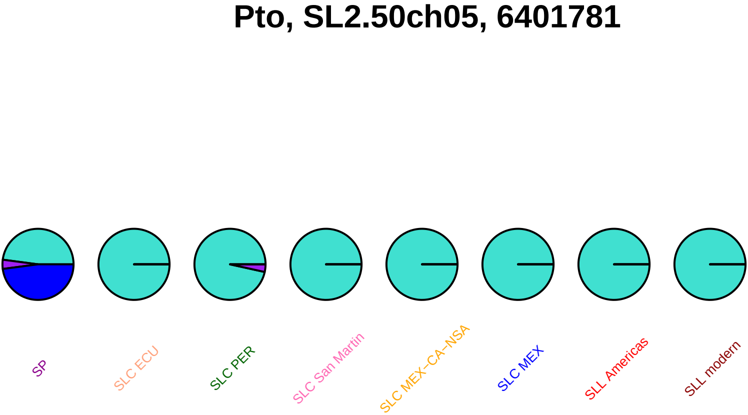
Pto, SL2.50ch05, 6401781
SP
SLC ECU
SLC PER
SLC San Martin
SLC MEX–CA–NSA
SLC MEX
SLL Americas
SLL modern

#### SlDEAH14, SL2.50ch05, 8824455

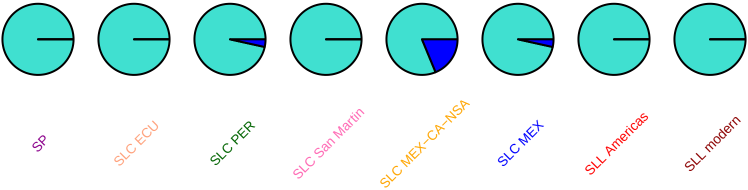

#### aml1, SL2.50ch05, 29173865

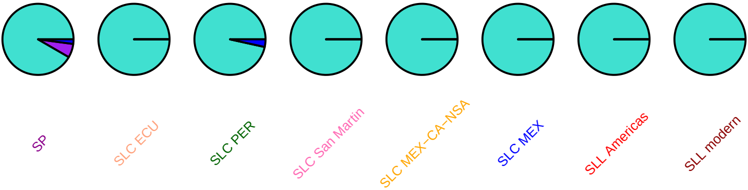

### prk3, SL2.50ch05, 36715385

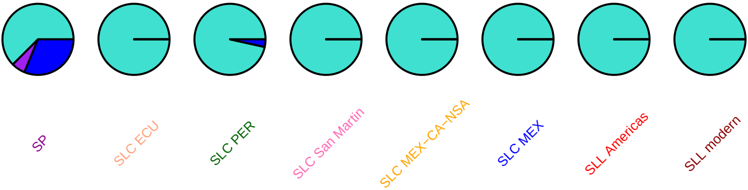

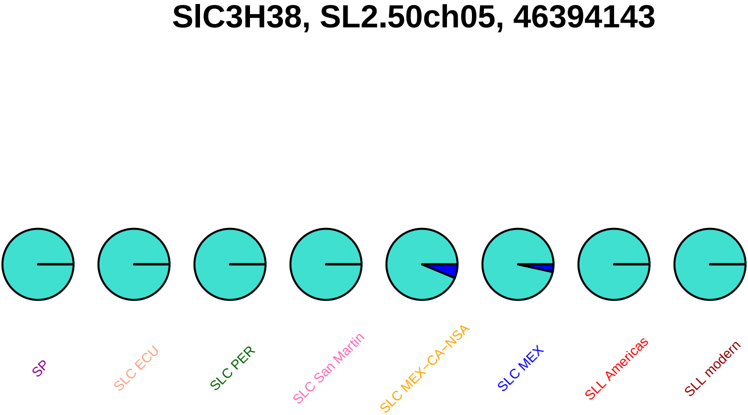
SlC3H38, SL2.50ch05, 46394143
SP
SLC ECU
SLC PER
SLC San Martin
SLC MEX−CA−NSA
SLC MEX
SLL Americas
SLL modern

### SlC3H38, SL2.50ch05, 46394678

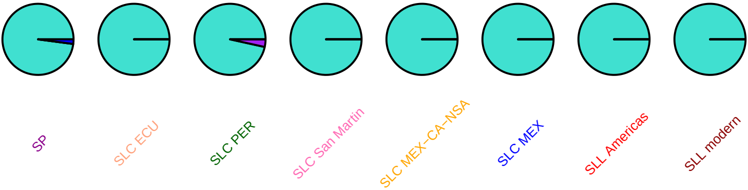

### SlDEAH17, SL2.50ch05, 56699083

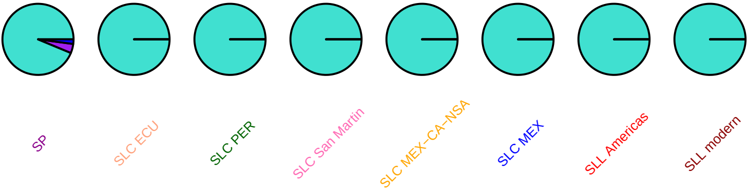

### SlDEAH17, SL2.50ch05, 56699751

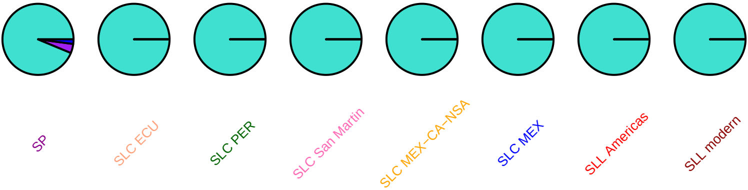

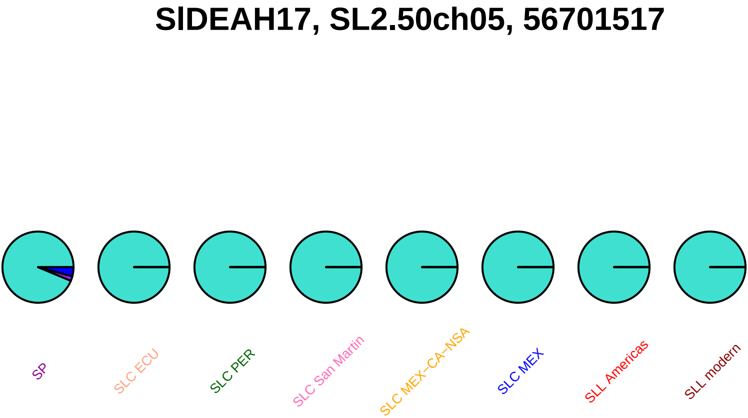
SlDEAH17, SL2.50ch05, 56701517
SP
SLC ECU
SLC PER
SLC San Martin
SLC MEX−CA−NSA
SLC MEX
SLL Americas
SLL modern

### SlDEAH17, SL2.50ch05, 56701811

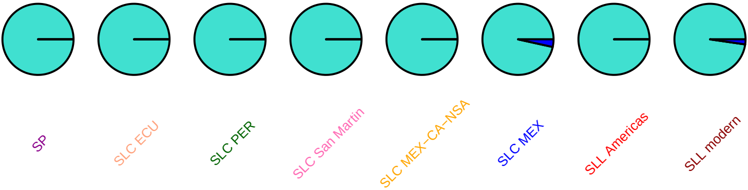

### SlWRKY65, SL2.50ch05, 57557681

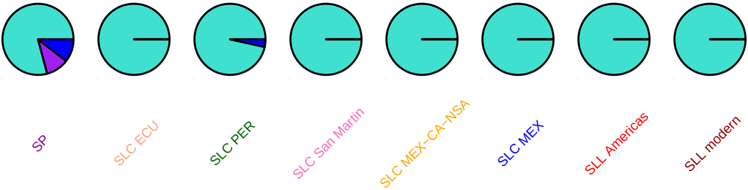

#### SlWRKY65, SL2.50ch05, 57557986

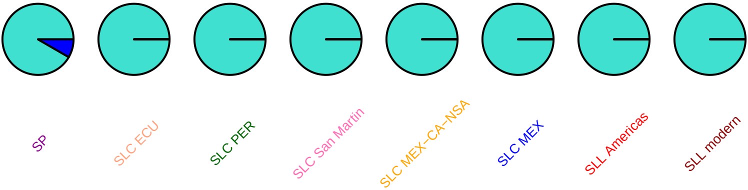

### SlWRKY65, SL2.50ch05, 57558022

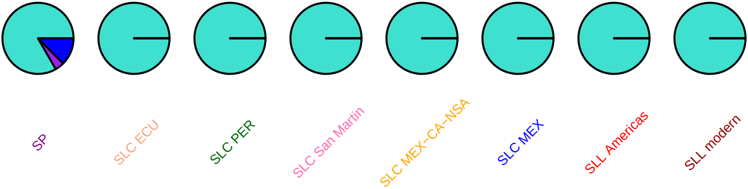

### SlWRKY65, SL2.50ch05, 57558245

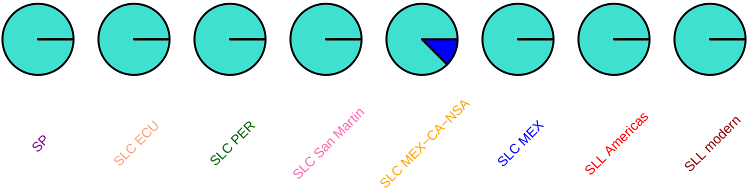

### SlWRKY67, SL2.50ch05, 57587881

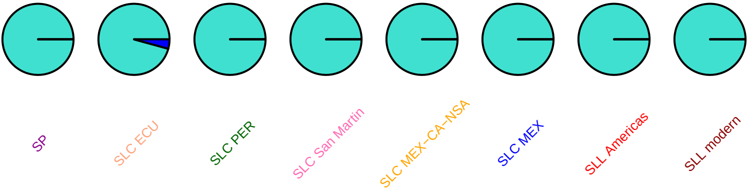

### SlWRKY66, SL2.50ch05, 57609343

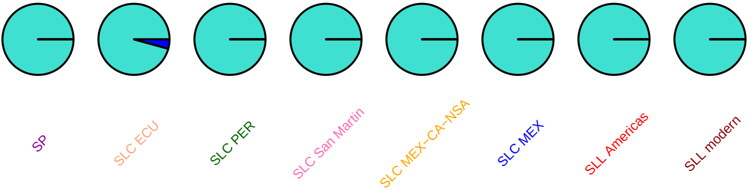

### SlSAUR55, SL2.50ch05, 58550615

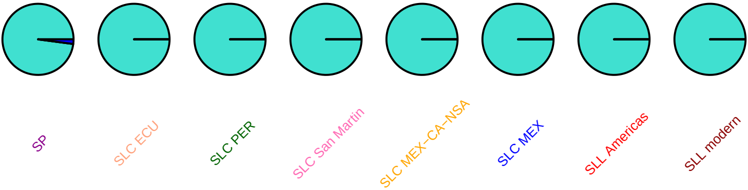

### Sl−ARF7B, SL2.50ch05, 58908971

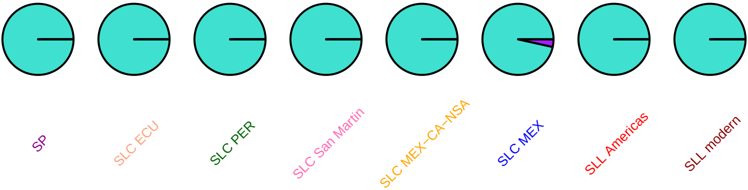

## Z70216, SL2.50ch05, 59013768

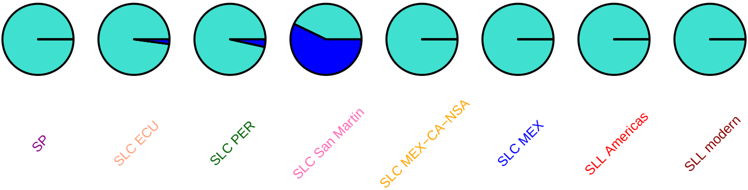

# Z70216, SL2.50ch05, 59014360

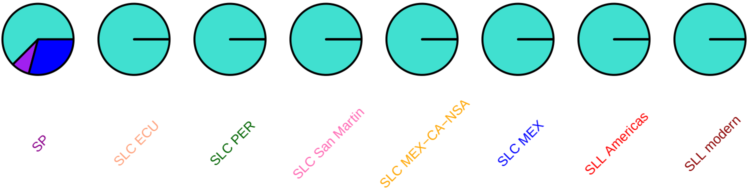

### SlDEAD18, SL2.50ch05, 59572633

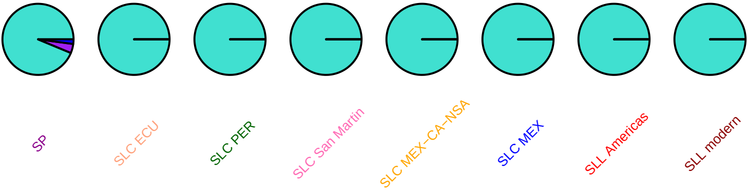

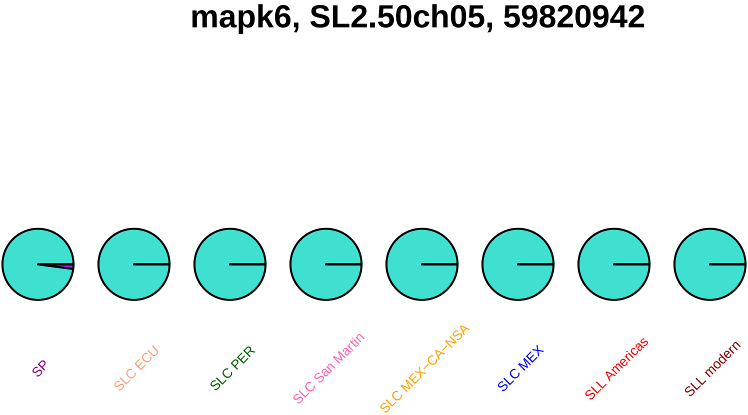
mapk6, SL2.50ch05, 59820942
SP
SLC ECU
SLC PER
SLC San Martin
SLC MEX−CA−NSA
SLC MEX
SLL Americas
SLL modern

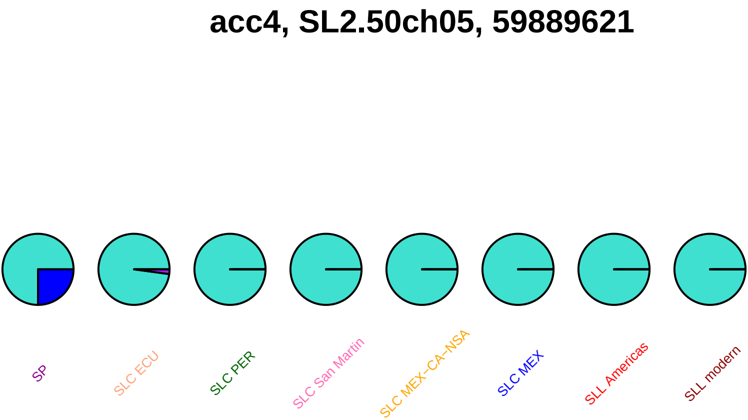
acc4, SL2.50ch05, 59889621
SP
SLC ECU
SLC PER
SLC San Martin
SLC MEX−CA−NSA
SLC MEX
SLL Americas
SLL modern

### acc4, SL2.50ch05, 59889719

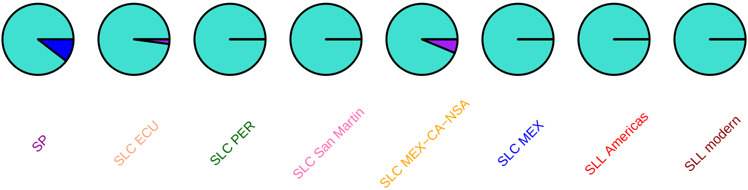

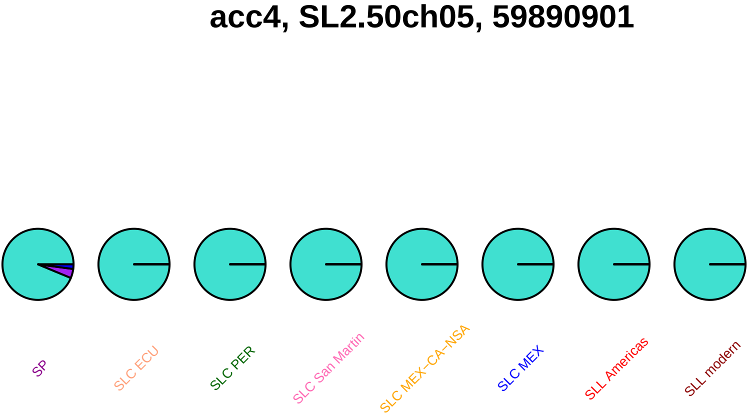
acc4, SL2.50ch05, 59890901
SP
SLC ECU
SLC PER
SLC San Martin
SLC MEX–CA–NSA
SLC MEX
SLL Americas
SLL modern

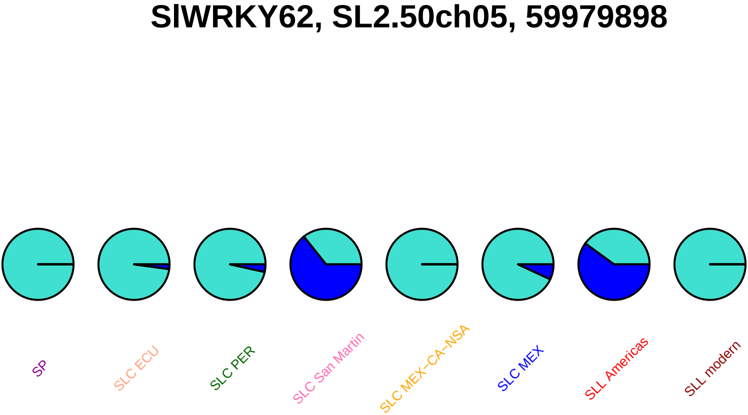
SlWRKY62, SL2.50ch05, 59979898
SP
SLC ECU
SLC PER
SLC San Martin
SLC MEX−CA−NSA
SLC MEX
SLL Americas
SLL modern

### SlWRKY62, SL2.50ch05, 59979900

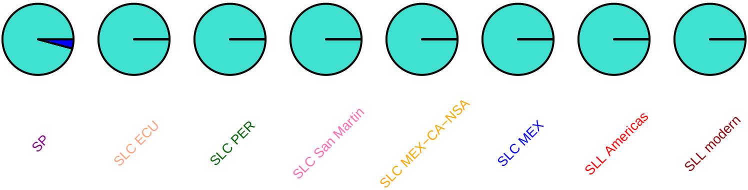

### SlWRKY63, SL2.50ch05, 59986428

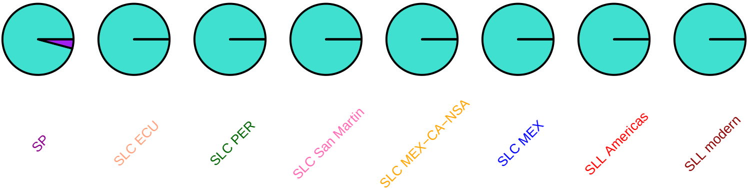

### SlWRKY60, SL2.50ch05, 60451966

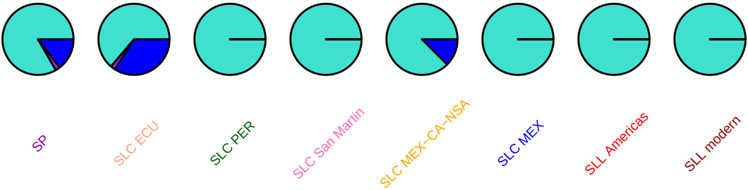

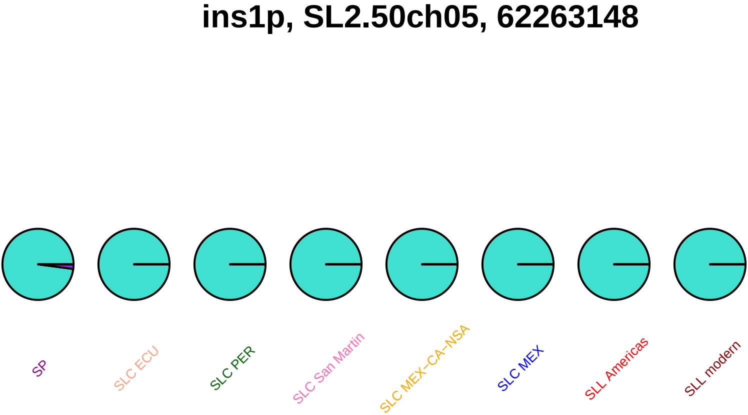
ins1p, SL2.50ch05, 62263148
SP
SLC ECU
SLC PER
SLC San Martin
SLC MEX−CA−NSA
SLC MEX
SLL Americas
SLL modern

### SlC3H39, SL2.50ch05, 62781133

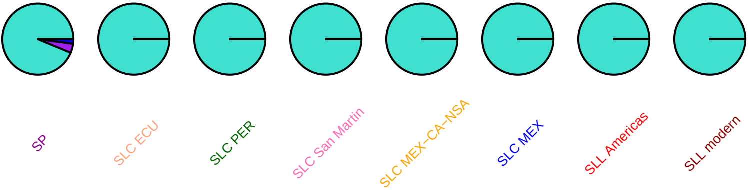

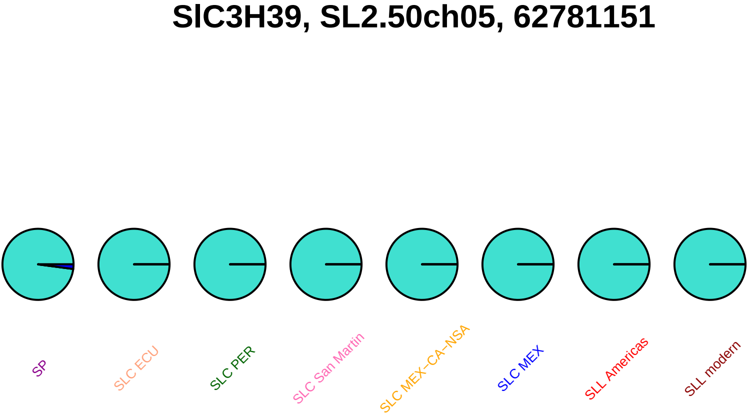
SlC3H39, SL2.50ch05, 62781151
SP
SLC ECU
SLC PER
SLC San Martin
SLC MEX−CA−NSA
SLC MEX
SLL Americas
SLL modern

### SlMYB109, SL2.50ch05, 63038048

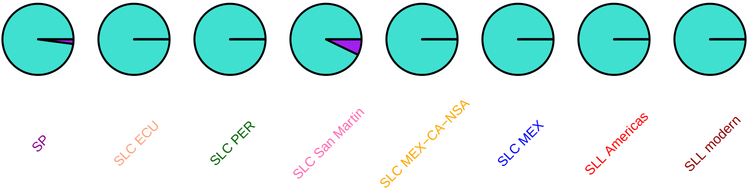

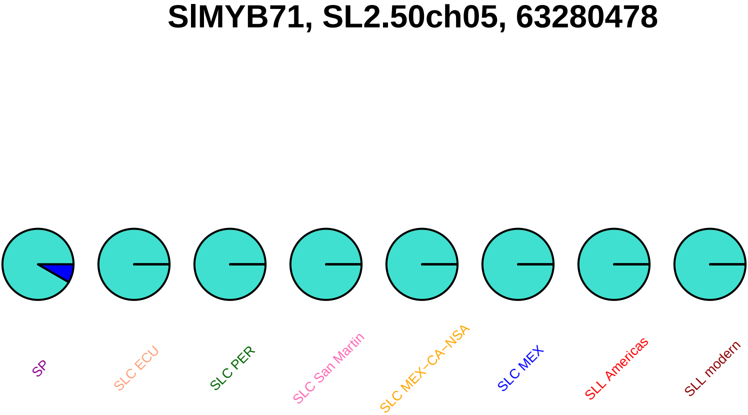
SlMYB71, SL2.50ch05, 63280478
SP
SLC ECU
SLC PER
SLC San Martin
SLC MEX–CA–NSA
SLC MEX
SLL Americas
SLL modern

#### SlMYB71, SL2.50ch05, 63280749

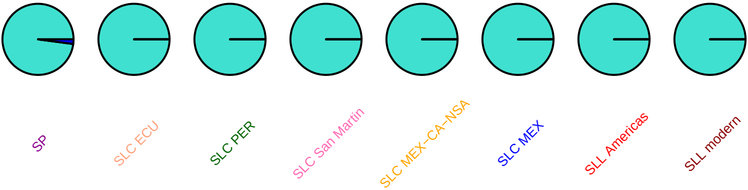

#### tbp1, SL2.50ch05, 64340467

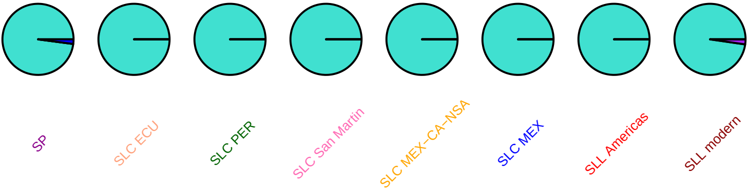

#### tbp1, SL2.50ch05, 64340907

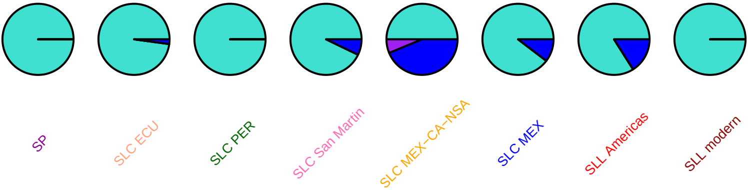

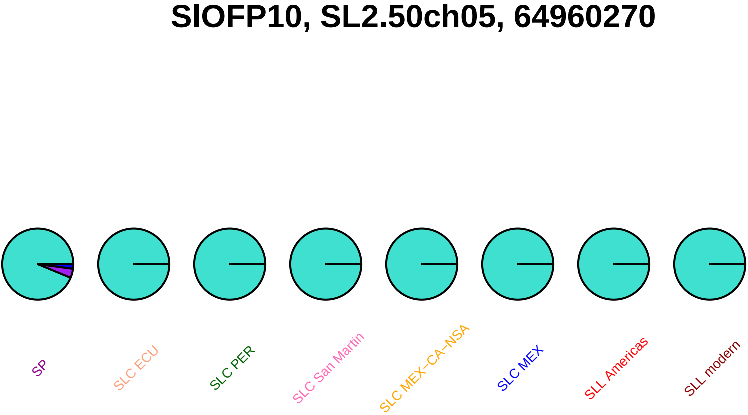
SlOFP10, SL2.50ch05, 64960270
SP
SLC ECU
SLC PER
SLC San Martin
SLC MEX−CA−NSA
SLC MEX
SLL Americas
SLL modern

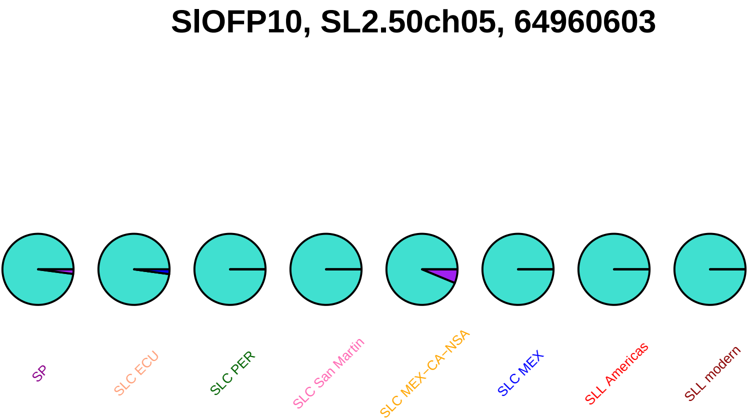
SlOFP10, SL2.50ch05, 64960603
SP
SLC ECU
SLC PER
SLC San Martin
SLC MEX−CA−NSA
SLC MEX
SLL Americas
SLL modern

#### ipi, SL2.50ch05, 65269905

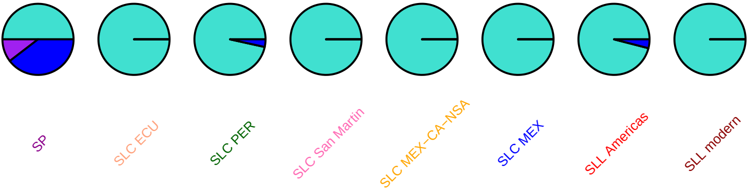

### SlPIP2.12, SL2.50ch05, 65393587

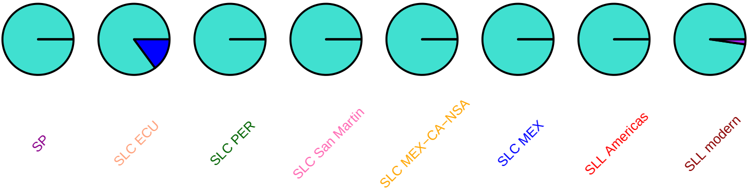

### SlPIP2.12, SL2.50ch05, 65394452

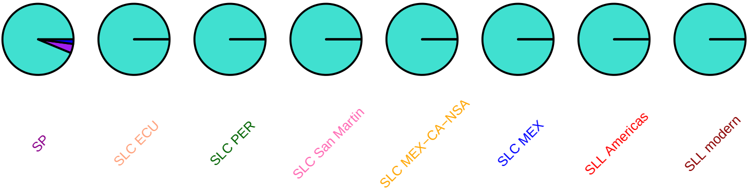

### Sl−ARF24, SL2.50ch05, 65427386

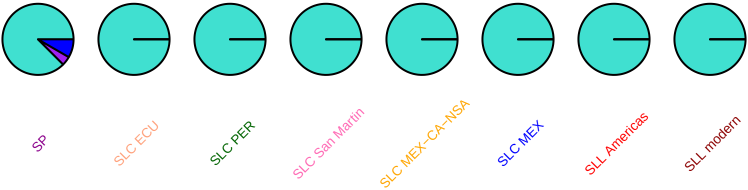

#### tpk1b, SL2.50ch06, 527459

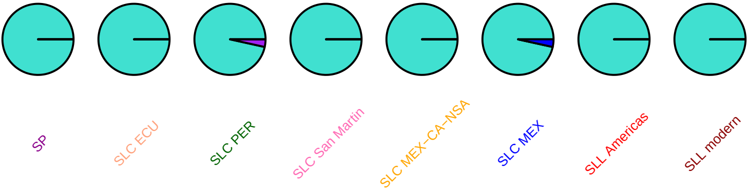

### PLC4, SL2.50ch06, 1179532

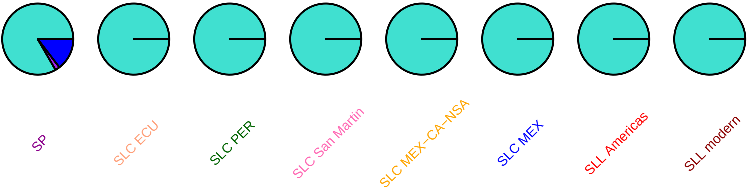

# Mi1_6, SL2.50ch06, 2355704

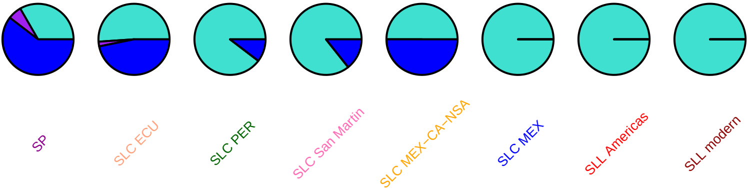

# Mi1_6, SL2.50ch06, 2356501

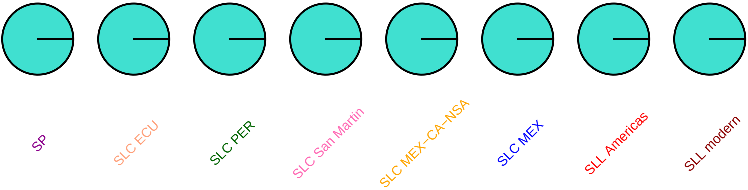

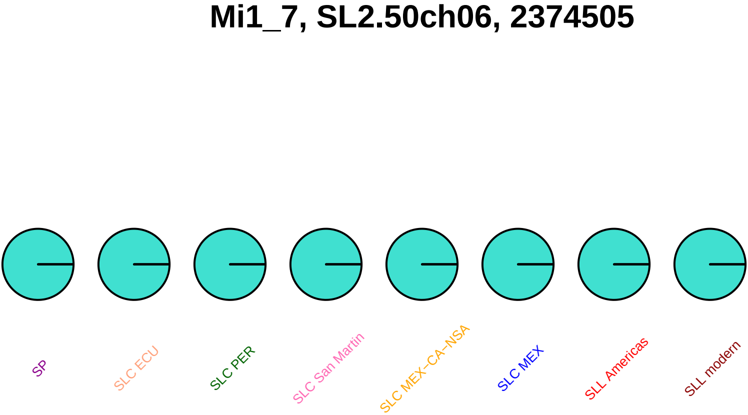
Mi1_7, SL2.50ch06, 2374505
SP
SLC ECU
SLC PER
SLC San Martin
SLC MEX−CA−NSA
SLC MEX
SLL Americas
SLL modern

### SlIAA17, SL2.50ch06, 2500967

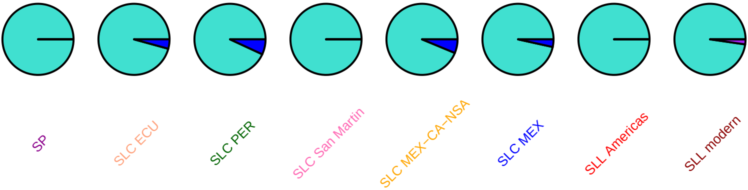

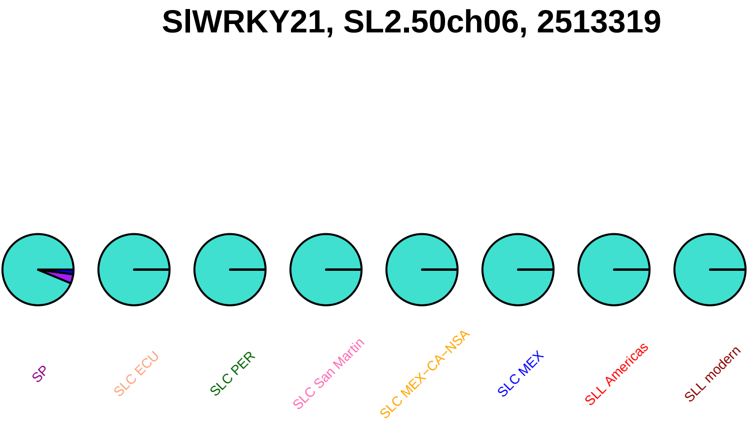
SlWRKY21, SL2.50ch06, 2513319
SP
SLC ECU
SLC PER
SLC San Martin
SLC MEX–CA–NSA
SLC MEX
SLL Americas
SLL modern

### SlWRKY21, SL2.50ch06, 2514589

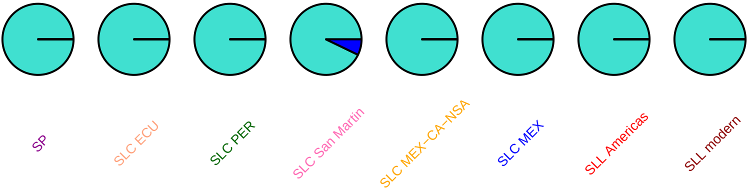

### SlC3H40, SL2.50ch06, 2664926

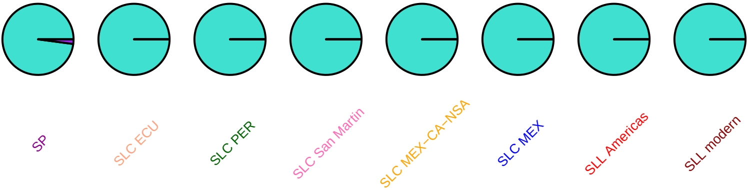

#### CNL4, SL2.50ch06, 2705599

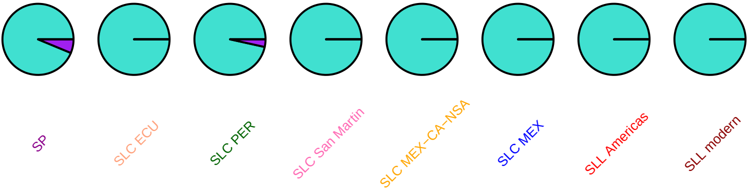

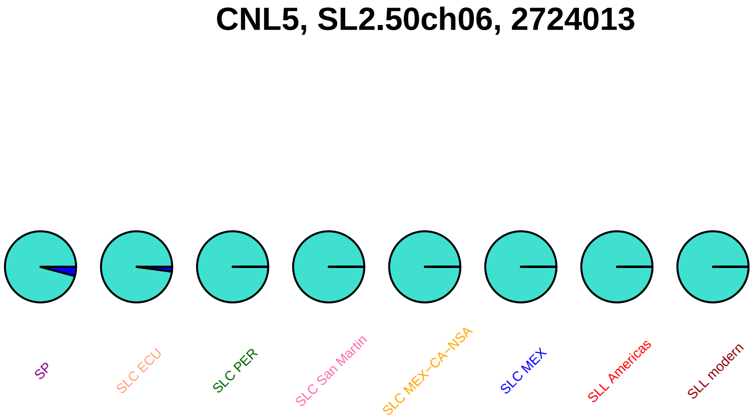
CNL5, SL2.50ch06, 2724013
SP
SLC ECU
SLC PER
SLC San Martin
SLC MEX–CA–NSA
SLC MEX
SLL Americas
SLL modern

### CNL5, SL2.50ch06, 2724548

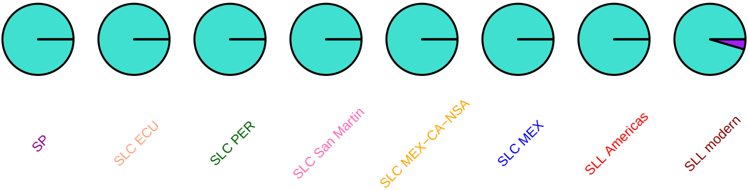

### nhx1, SL2.50ch06, 2766345

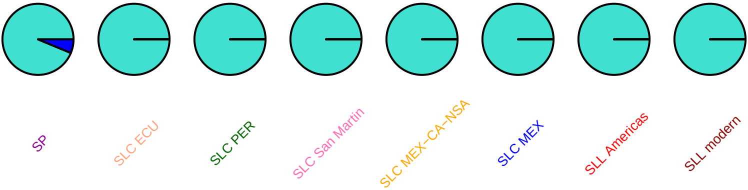

### nhx1, SL2.50ch06, 2769565

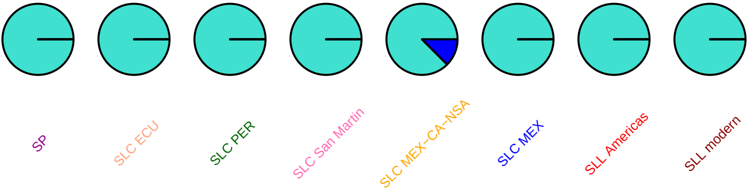

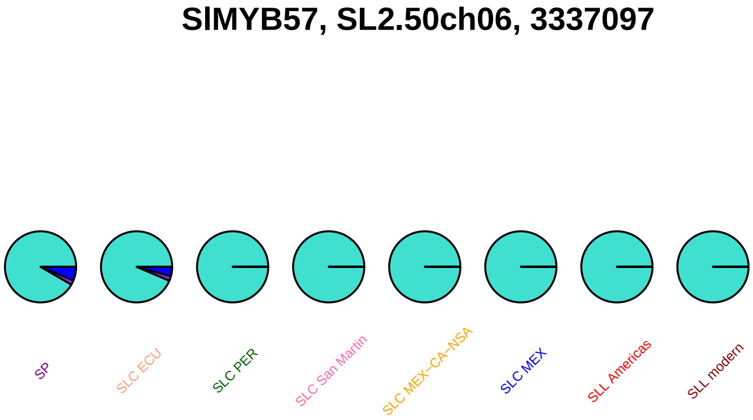
SlMYB57, SL2.50ch06, 3337097
SP
SLC ECU
SLC PER
SLC San Martin
SLC MEX−CA−NSA
SLC MEX
SLL Americas
SLL modern

#### SlMYB57, SL2.50ch06, 3337120

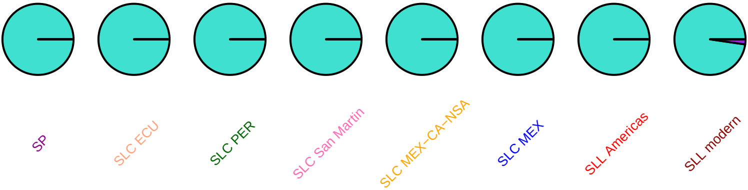

### SlMYB24, SL2.50ch06, 3426526

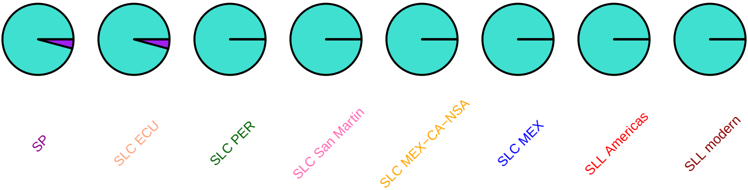

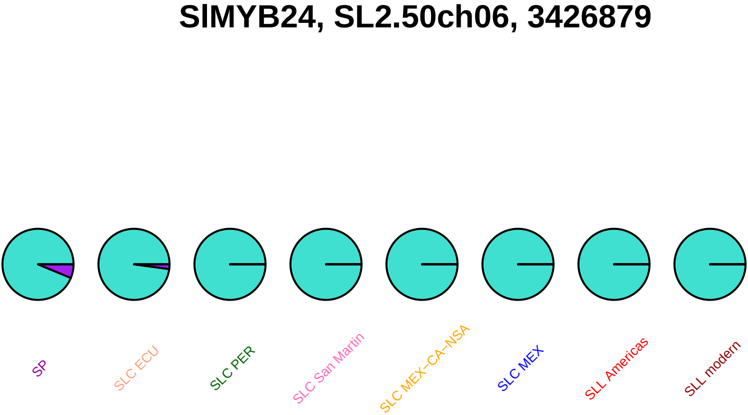
SlMYB24, SL2.50ch06, 3426879
SP
SLC ECU
SLC PER
SLC San Martin
SLC MEX−CA−NSA
SLC MEX
SLL Americas
SLL modern

#### SlMAPKKK37, SL2.50ch06, 25474015

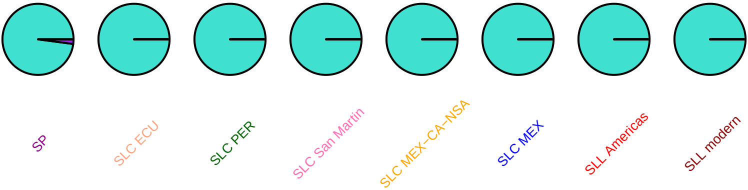

#### gras9, SL2.50ch06, 25639621

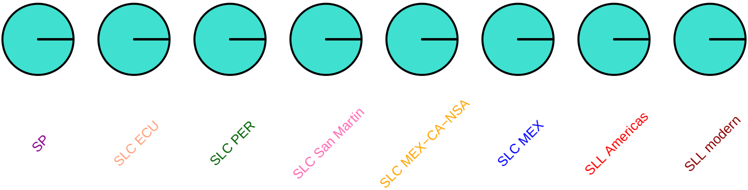

#### SlWRKY19, SL2.50ch06, 31848254

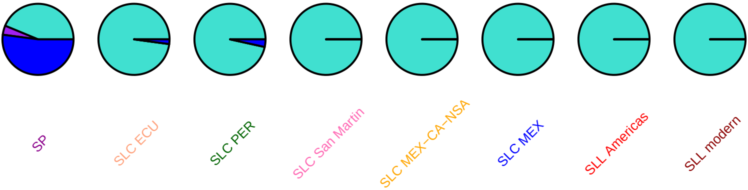

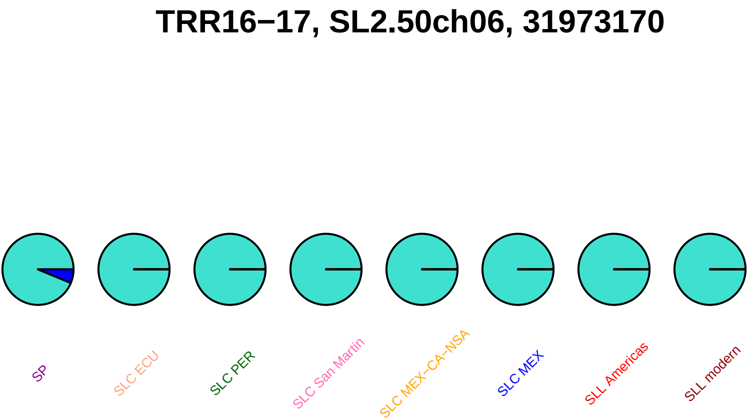
TRR16–17, SL2.50ch06, 31973170
SP
SLC ECU
SLC PER
SLC San Martin
SLC MEX–CA–NSA
SLC MEX
SLL Americas
SLL modern

### SlDCL2a, SL2.50ch06, 32062080

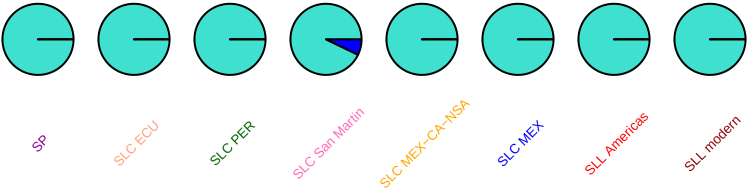

### SlDCL2a, SL2.50ch06, 32064188

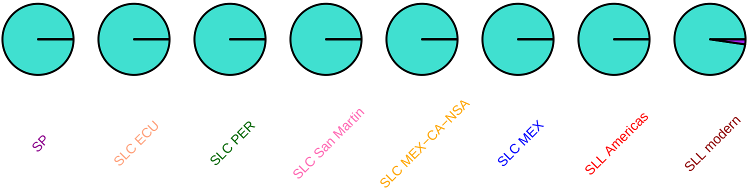

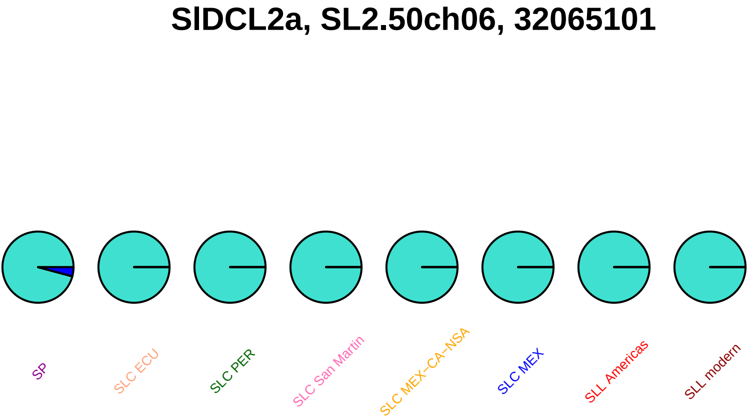
SlDCL2a, SL2.50ch06, 32065101
SP
SLC ECU
SLC PER
SLC San Martin
SLC MEX−CA−NSA
SLC MEX
SLL Americas
SLL modern

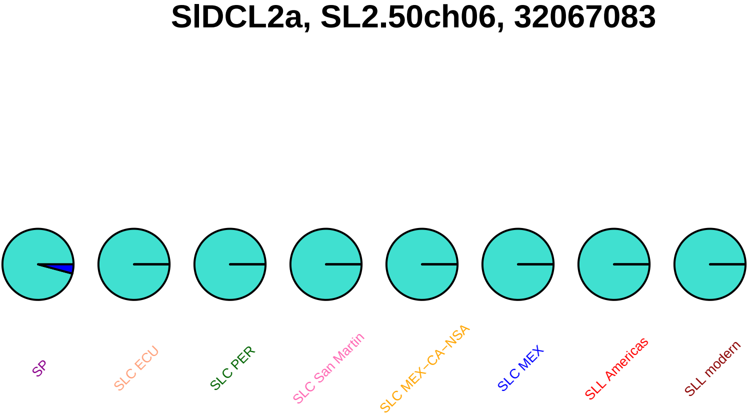

SlDCL2a, SL2.50ch06, 32067083
SP
SLC ECU
SLC PER
SLC San Martin
SLC MEX−CA−NSA
SLC MEX
SLL Americas
SLL modern

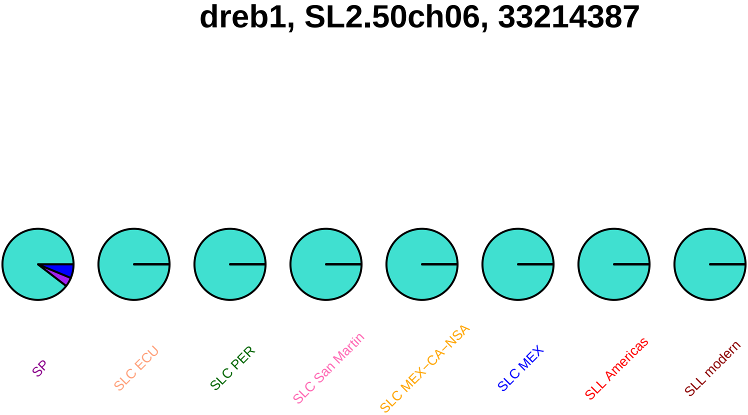
dreb1, SL2.50ch06, 33214387
SP
SLC ECU
SLC PER
SLC San Martin
SLC MEX−CA−NSA
SLC MEX
SLL Americas
SLL modern

#### SlbHLH043, SL2.50ch06, 34496466

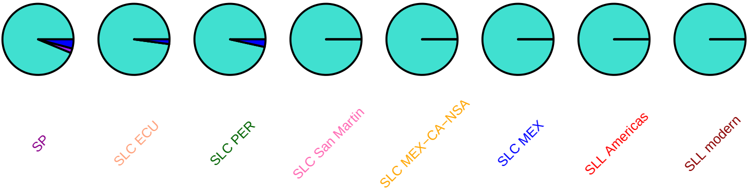

#### SlSUN16, SL2.50ch06, 35676627

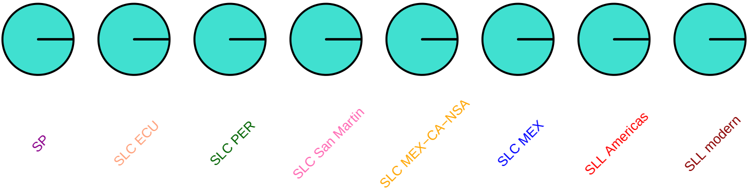

#### ppck2, SL2.50ch06, 36454098

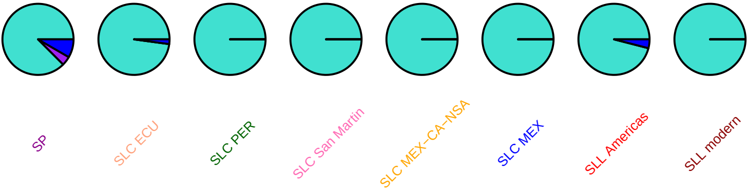

### etr4, SL2.50ch06, 36556893

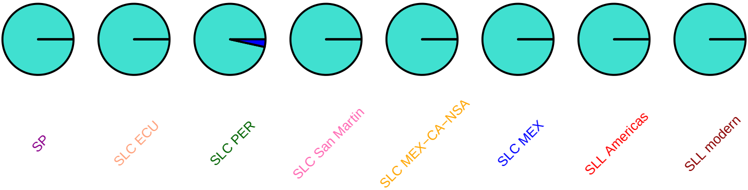

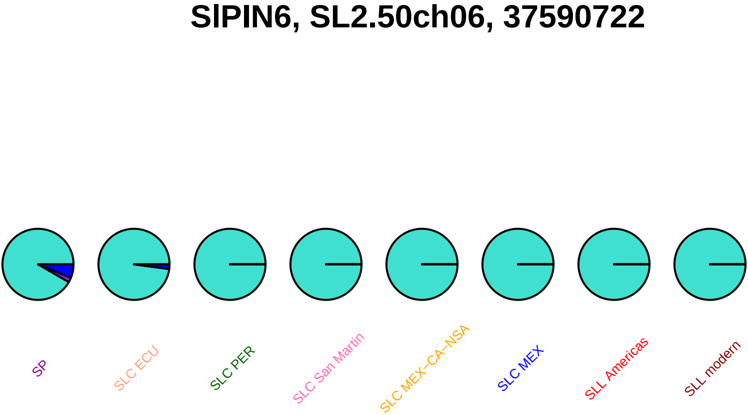
SlPIN6, SL2.50ch06, 37590722
SP
SLC ECU
SLC PER
SLC San Martin
SLC MEX−CA−NSA
SLC MEX
SLL Americas
SLL modern

## Z94180, SL2.50ch06, 37730437

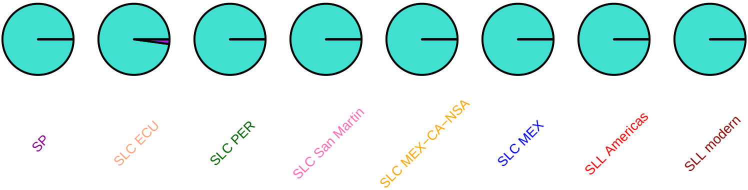

#### sstle1, SL2.50ch06, 37834388

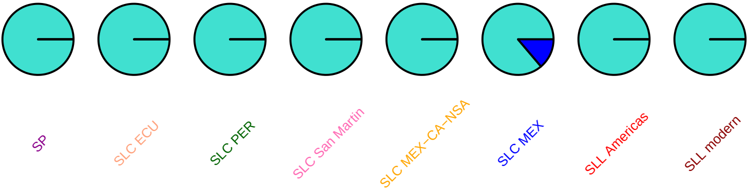

### sstle1, SL2.50ch06, 37834427

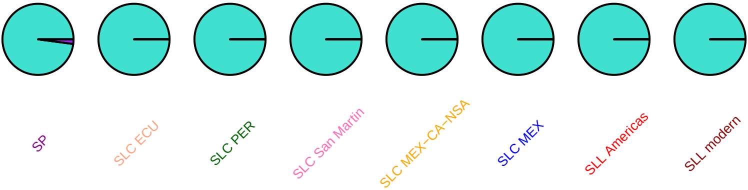

### sstle1, SL2.50ch06, 37862766

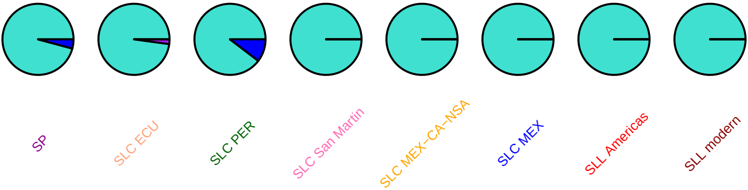

#### sstle1, SL2.50ch06, 37863225

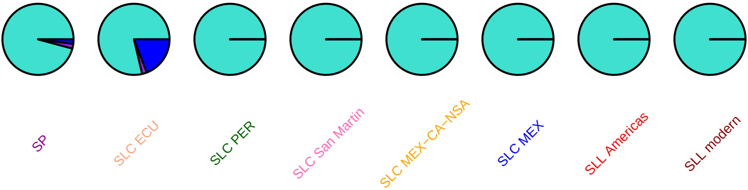

### psd, SL2.50ch06, 38800575

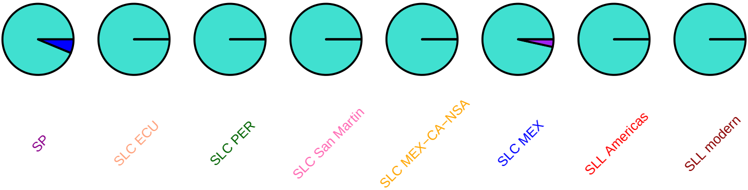

### SlbHLH136, SL2.50ch06, 39408050

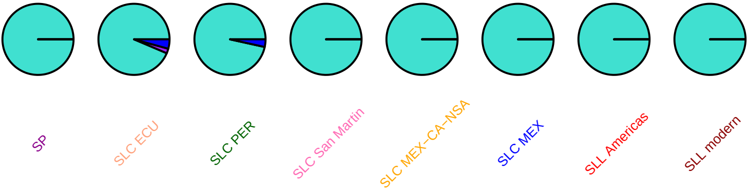

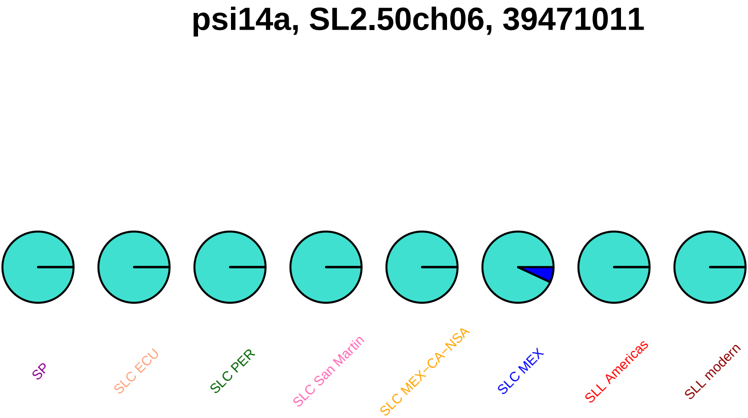

psi14a, SL2.50ch06, 39471011
SP
SLC ECU
SLC PER
SLC San Martin
SLC MEX−CA−NSA
SLC MEX
SLL Americas
SLL modern

### Sl−ERF_A_3, SL2.50ch06, 39817807

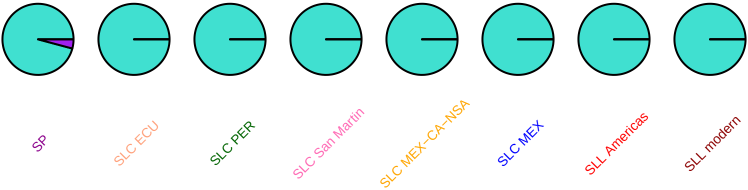

#### Sl−ERF_A_3, SL2.50ch06, 39818224

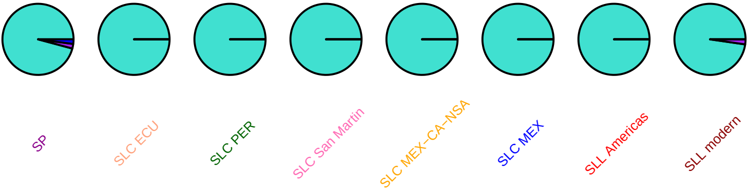

### Sl−ERF_A_3, SL2.50ch06, 39818340

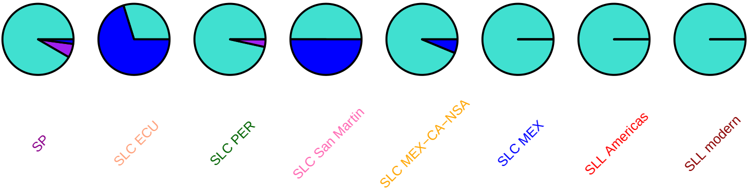

#### SlGLR2_3, SL2.50ch06, 39912715

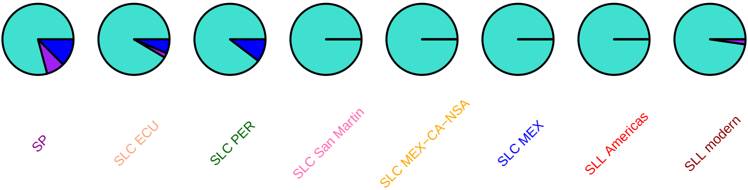

### SlGLR2_3, SL2.50ch06, 39912996

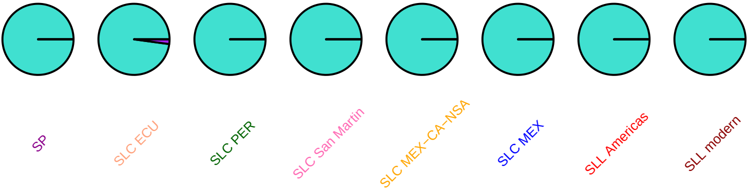

### SlGLR2_4, SL2.50ch06, 39924728

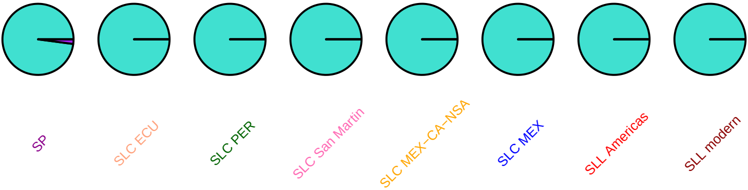

### SLGLR2_5, SL2.50ch06, 39926977

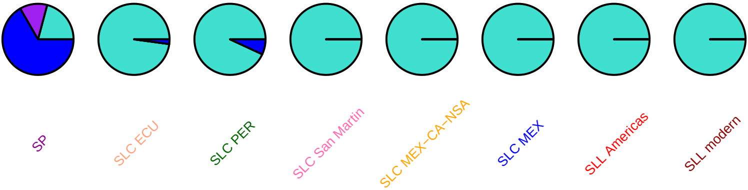

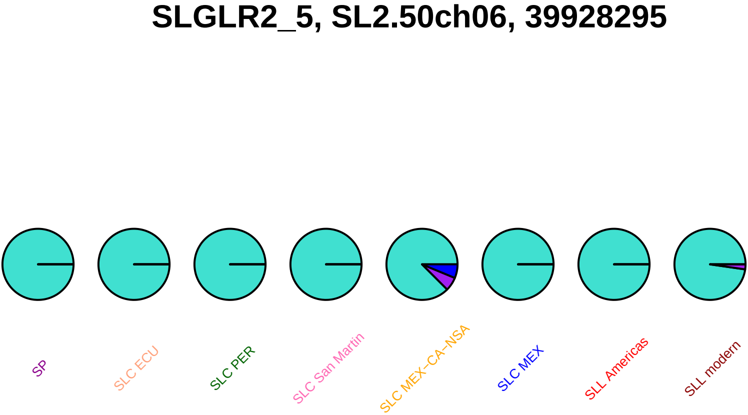
SLGLR2_5, SL2.50ch06, 39928295
SP
SLC ECU
SLC PER
SLC San Martin
SLC MEX−CA−NSA
SLC MEX
SLL Americas
SLL modern

### SLGLR2_5, SL2.50ch06, 39928601

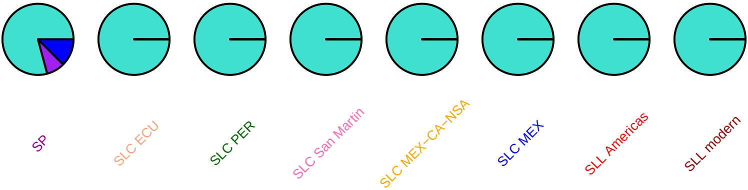

### man2, SL2.50ch06, 40185215

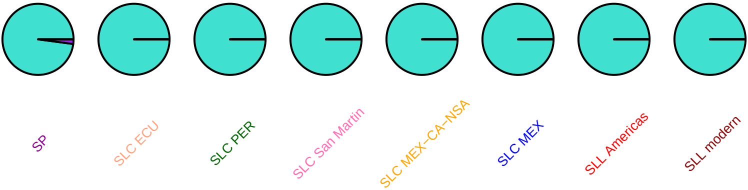

#### SlbHLH044, SL2.50ch06, 40246869

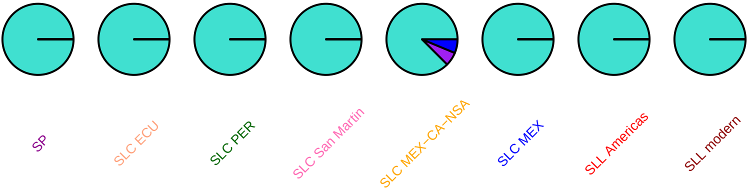

#### SlIAA36, SL2.50ch06, 41377379

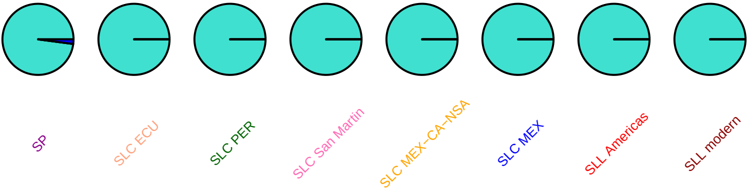

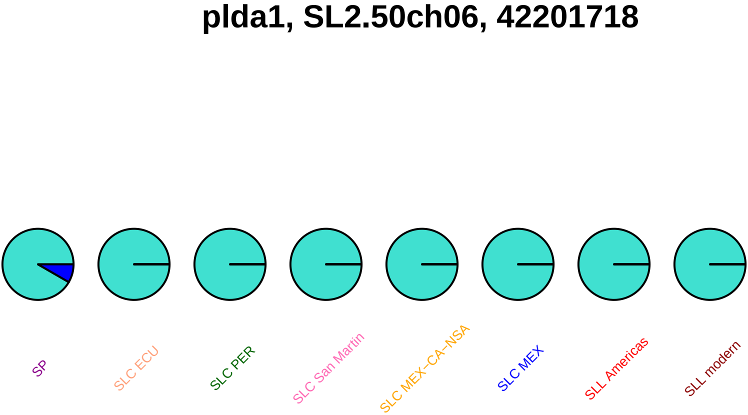
plda1, SL2.50ch06, 42201718
SP
SLC ECU
SLC PER
SLC San Martin
SLC MEX−CA−NSA
SLC MEX
SLL Americas
SLL modern

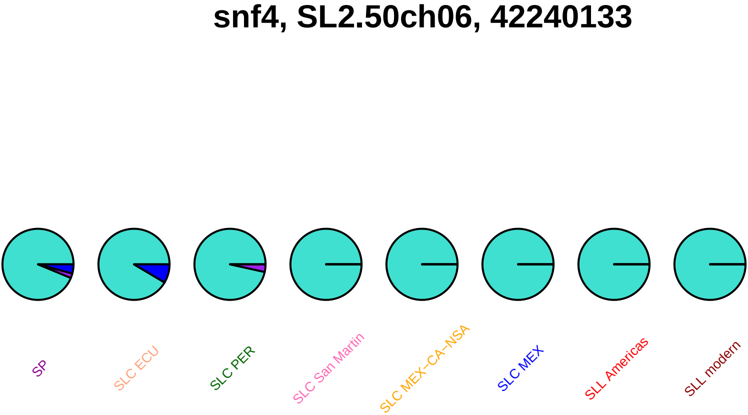
snf4, SL2.50ch06, 42240133
SP
SLC ECU
SLC PER
SLC San Martin
SLC MEX−CA−NSA
SLC MEX
SLL Americas
SLL modern

### SlMAPKKK38, SL2.50ch06, 42464388

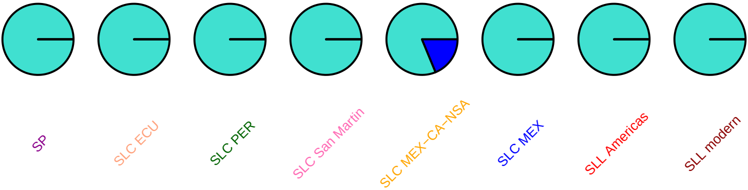

### fruitfull1, SL2.50ch06, 43177170

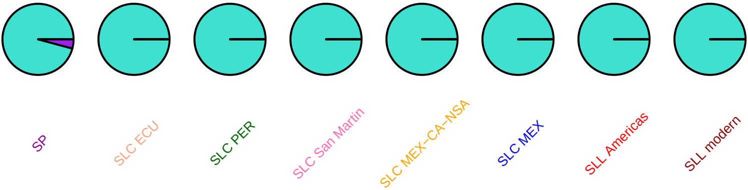

### SlbHLH047, SL2.50ch06, 43346636

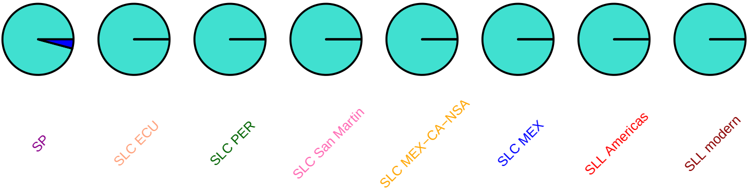

# kd1, SL2.50ch06, 44704112

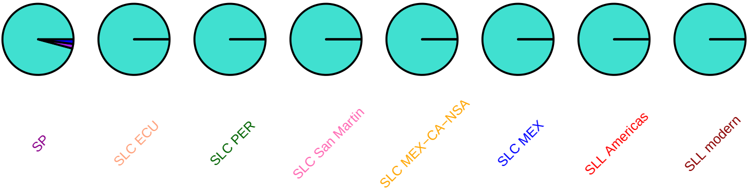

## AY568722, SL2.50ch06, 45028836

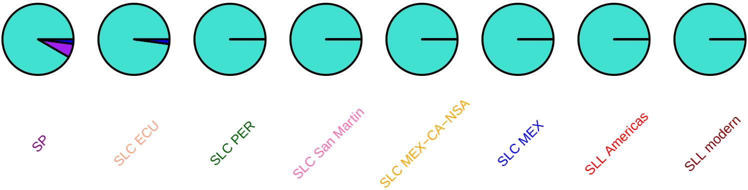

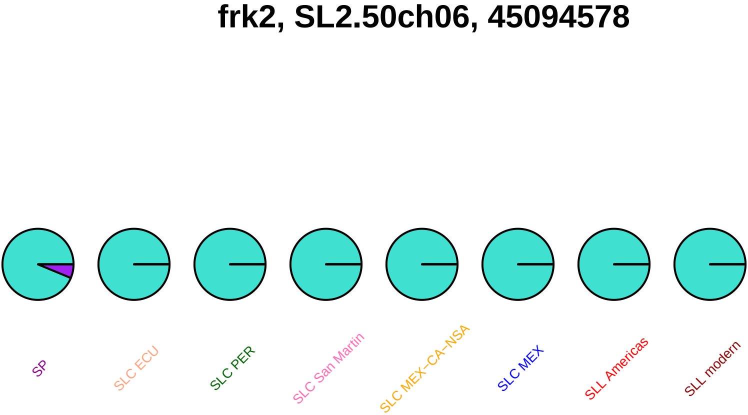
frk2, SL2.50ch06, 45094578
SP
SLC ECU
SLC PER
SLC San Martin
SLC MEX–CA–NSA
SLC MEX
SLL Americas
SLL modern

## AY301280, SL2.50ch06, 45275666

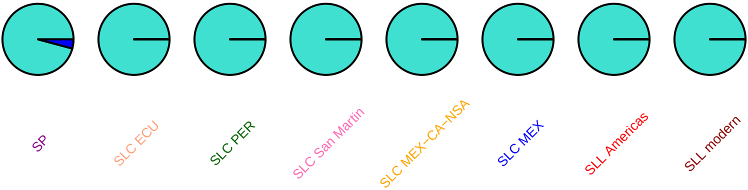

### SlAGO4c, SL2.50ch06, 45327577

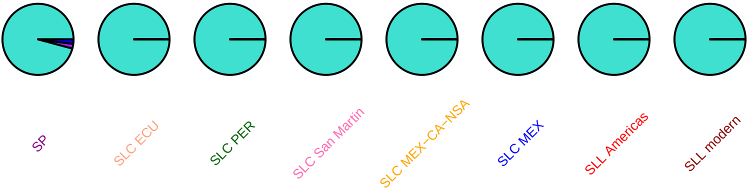

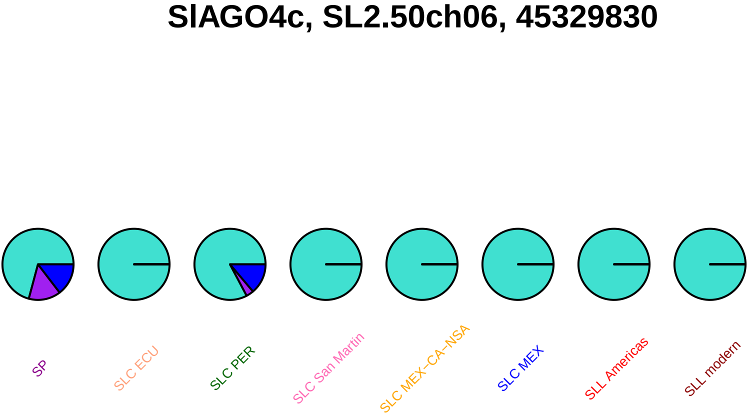
SlAGO4c, SL2.50ch06, 45329830
SP
SLC ECU
SLC PER
SLC San Martin
SLC MEX−CA−NSA
SLC MEX
SLL Americas
SLL modern

#### SlAGO4b, SL2.50ch06, 45335639

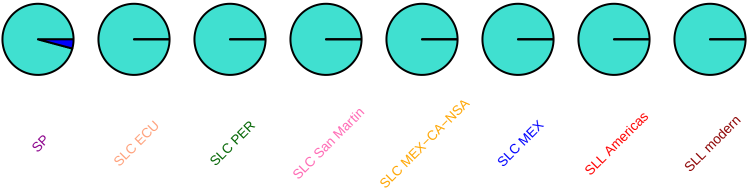

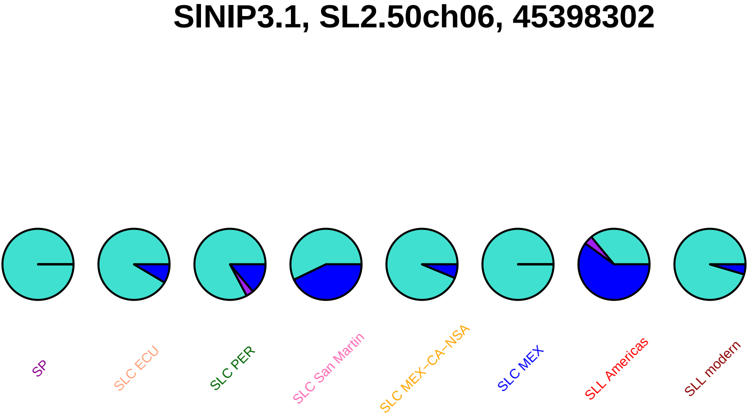
SlNIP3.1, SL2.50ch06, 45398302
SP
SLC ECU
SLC PER
SLC San Martin
SLC MEX−CA−NSA
SLC MEX
SLL Americas
SLL modern

### cycb1, SL2.50ch06, 45420259

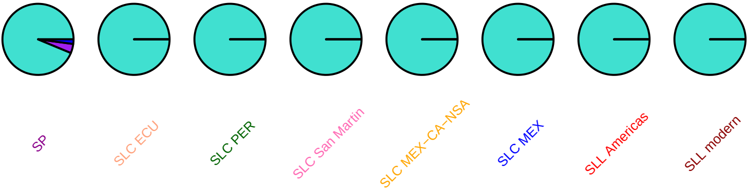

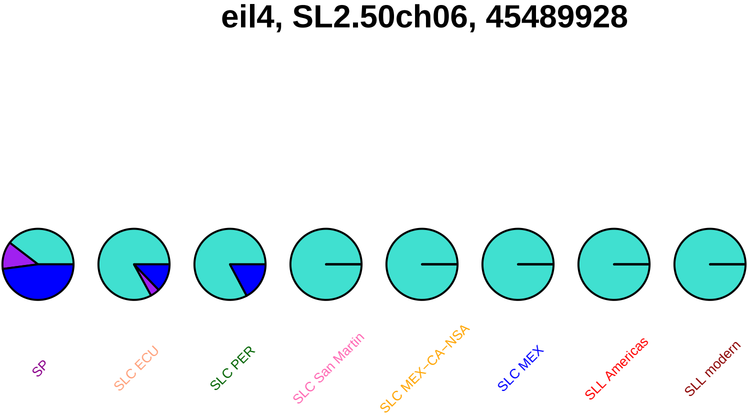
eil4, SL2.50ch06, 45489928
SP
SLC ECU
SLC PER
SLC San Martin
SLC MEX–CA–NSA
SLC MEX
SLL Americas
SLL modern

#### cyc−b, SL2.50ch06, 45898715

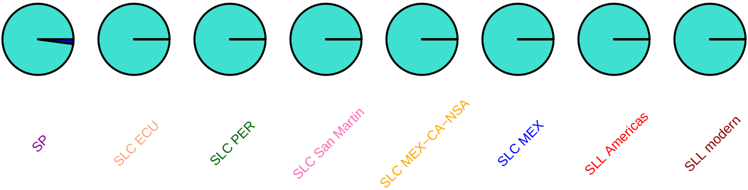

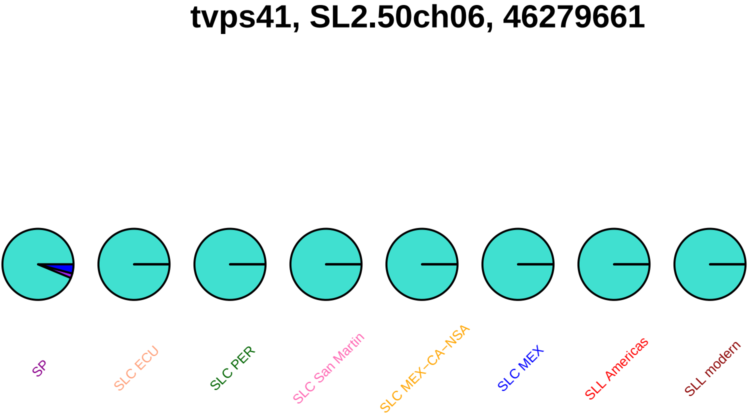
tvps41, SL2.50ch06, 46279661
SP
SLC ECU
SLC PER
SLC San Martin
SLC MEX−CA−NSA
SLC MEX
SLL Americas
SLL modern

### tvps41, SL2.50ch06, 46280306

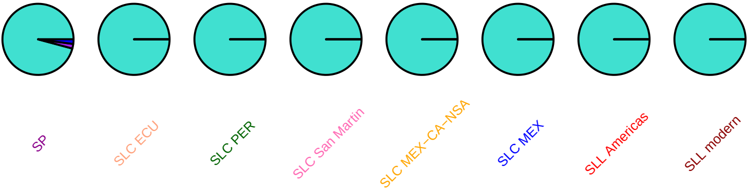

### bli6, SL2.50ch06, 46422610

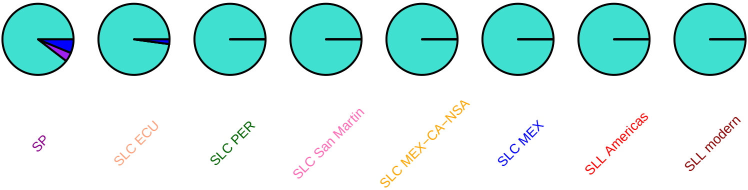

### bli6, SL2.50ch06, 46422639

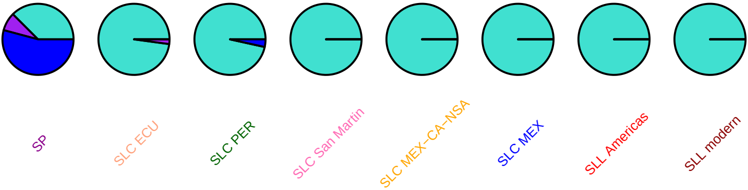

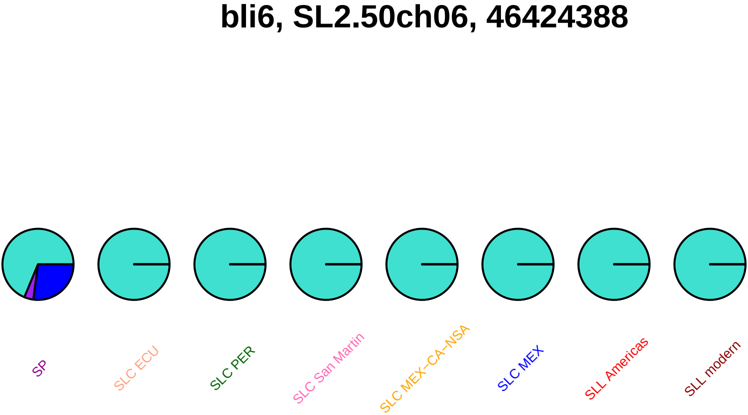
bli6, SL2.50ch06, 46424388
SP
SLC ECU
SLC PER
SLC San Martin
SLC MEX−CA−NSA
SLC MEX
SLL Americas
SLL modern

# AP2e, SL2.50ch06, 46865538

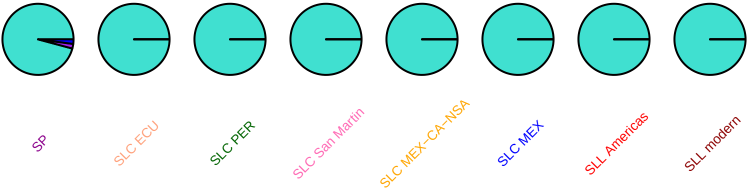

### pcl1, SL2.50ch06, 47438957

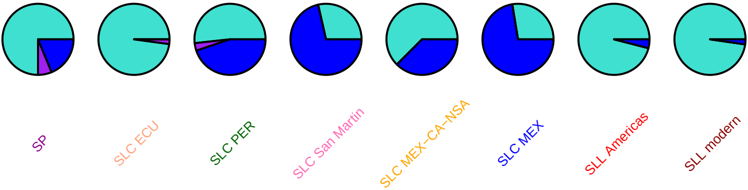

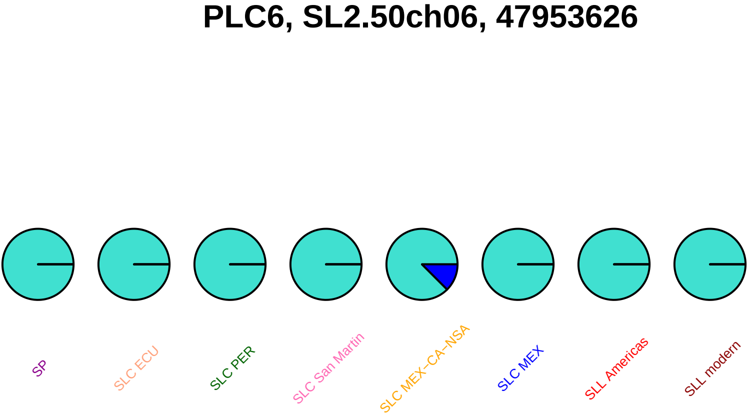
PLC6, SL2.50ch06, 47953626
SP
SLC ECU
SLC PER
SLC San Martin
SLC MEX−CA−NSA
SLC MEX
SLL Americas
SLL modern

### PLC6, SL2.50ch06, 47956055

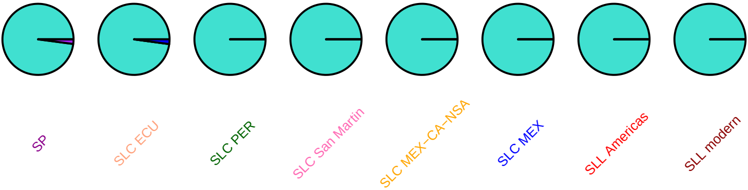

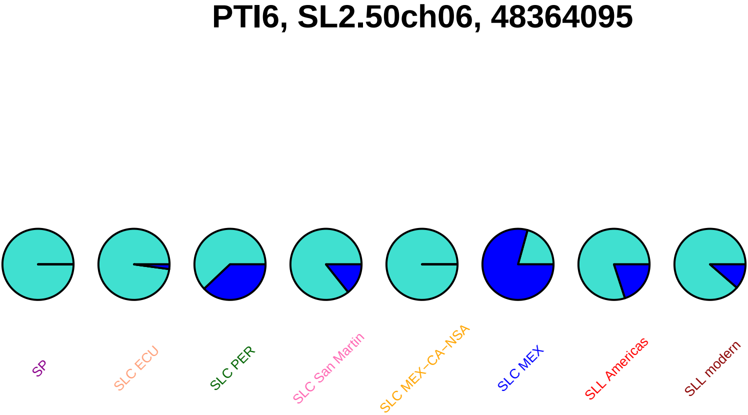
PTI6, SL2.50ch06, 48364095
SP
SLC ECU
SLC PER
SLC San Martin
SLC MEX–CA–NSA
SLC MEX
SLL Americas
SLL modern

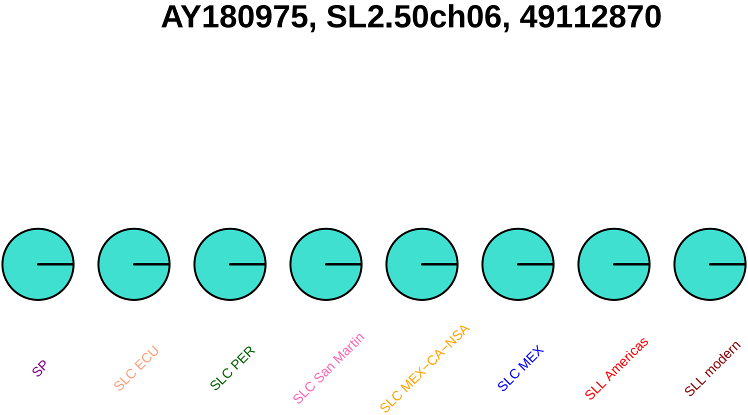
AY180975, SL2.50ch06, 49112870
SP
SLC ECU
SLC PER
SLC San Martin
SLC MEX−CA−NSA
SLC MEX
SLL Americas
SLL modern

#### SlIAA2, SL2.50ch06, 49327285

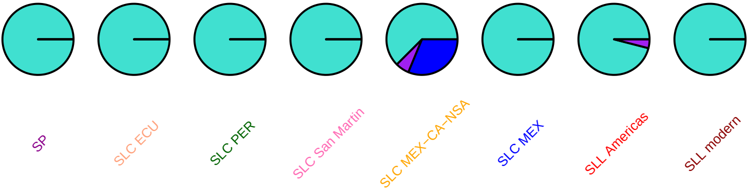

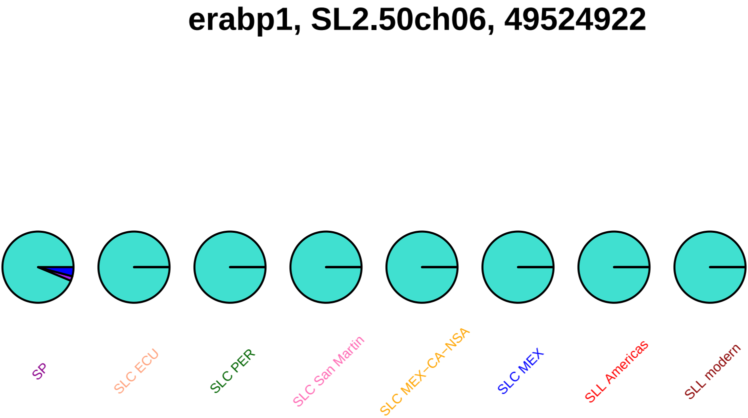
erabp1, SL2.50ch06, 49524922
SP
SLC ECU
SLC PER
SLC San Martin
SLC MEX−CA−NSA
SLC MEX
SLL Americas
SLL modern

### erabp1, SL2.50ch06, 49525714

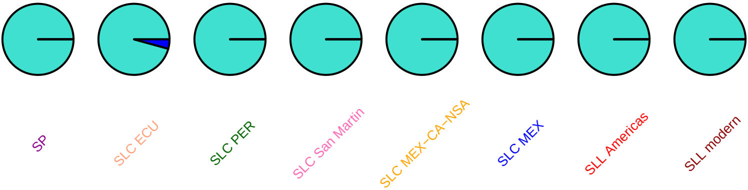

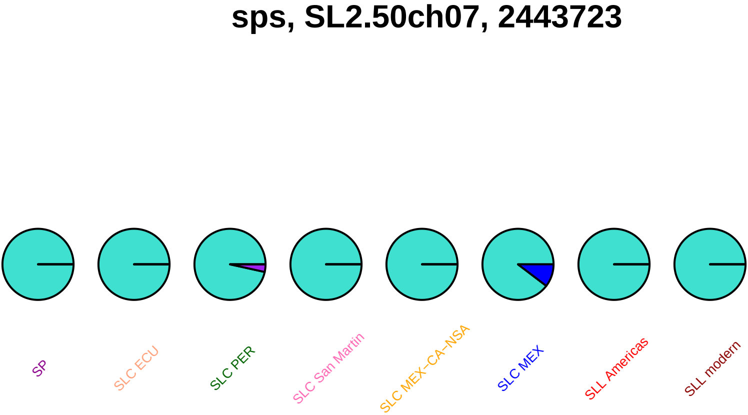
sps, SL2.50ch07, 2443723
SP
SLC ECU
SLC PER
SLC San Martin
SLC MEX−CA−NSA
SLC MEX
SLL Americas
SLL modern

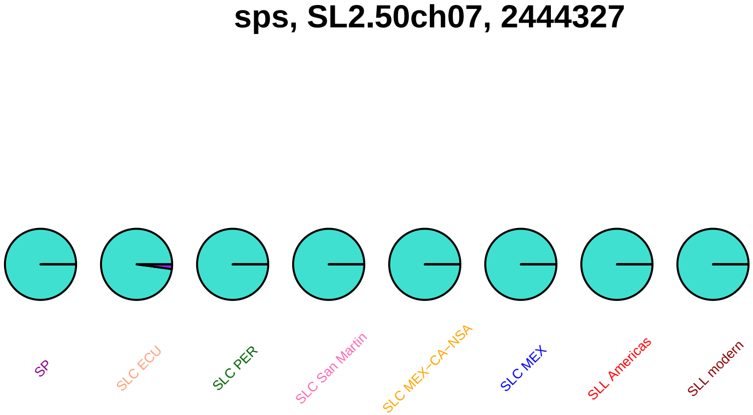
sps, SL2.50ch07, 2444327
SP
SLC ECU
SLC PER
SLC San Martin
SLC MEX−CA−NSA
SLC MEX
SLL Americas
SLL modern

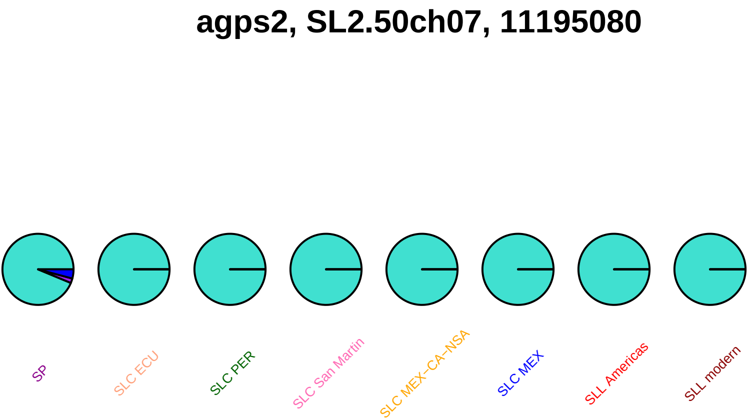

agps2, SL2.50ch07, 11195080
SP
SLC ECU
SLC PER
SLC San Martin
SLC MEX–CA–NSA
SLC MEX
SLL Americas
SLL modern

### SlpGlcT3, SL2.50ch07, 13485575

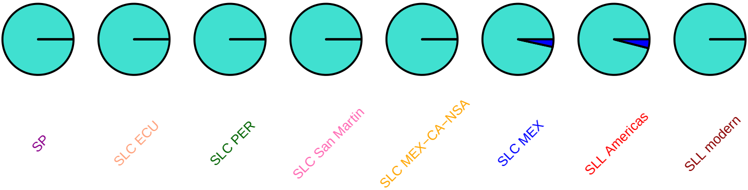

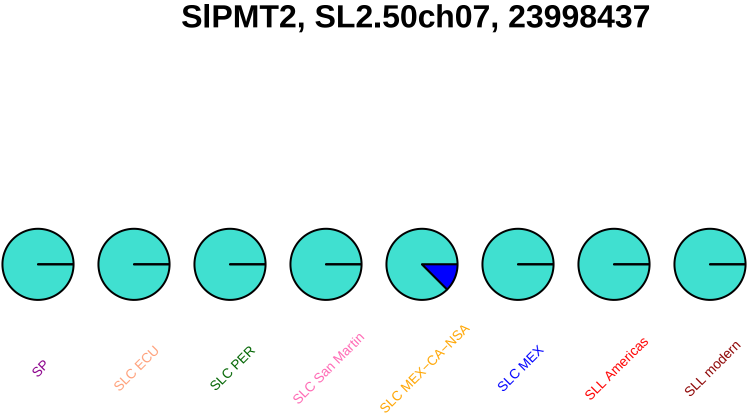
SlPMT2, SL2.50ch07, 23998437
SP
SLC ECU
SLC PER
SLC San Martin
SLC MEX−CA−NSA
SLC MEX
SLL Americas
SLL modern

### SlDEAD21, SL2.50ch07, 49534235

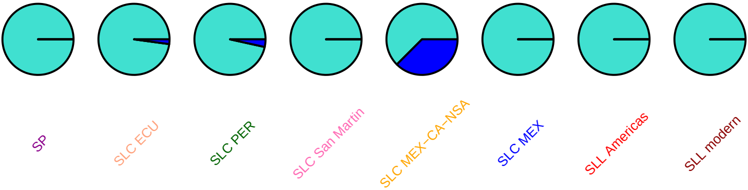

### SlDEAD21, SL2.50ch07, 49535463

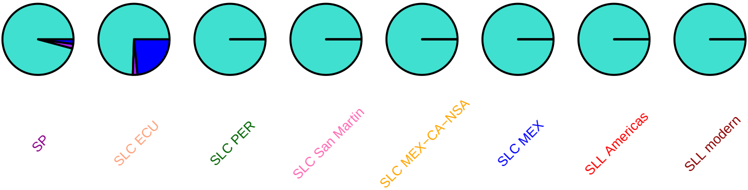

### GABA−TP1, SL2.50ch07, 56994739

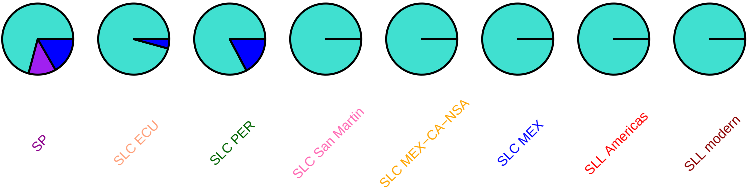

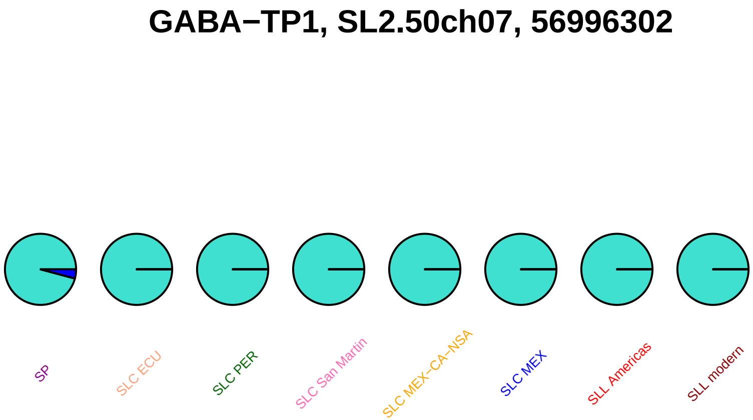
GABA−TP1, SL2.50ch07, 56996302
SP
SLC ECU
SLC PER
SLC San Martin
SLC MEX−CA−NSA
SLC MEX
SLL Americas
SLL modern

### GAME2, SL2.50ch07, 57130250

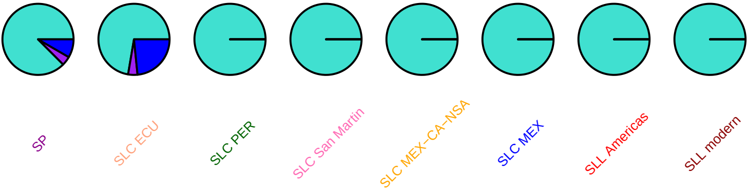

### GAME1, SL2.50ch07, 57318809

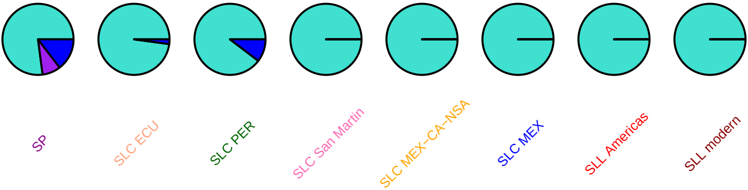

#### GAME18, SL2.50ch07, 57338604

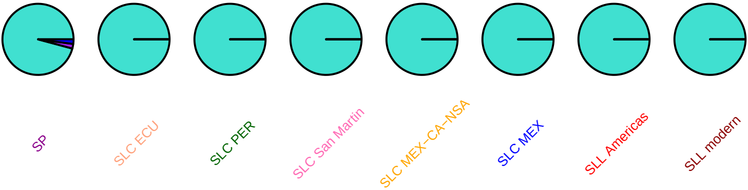

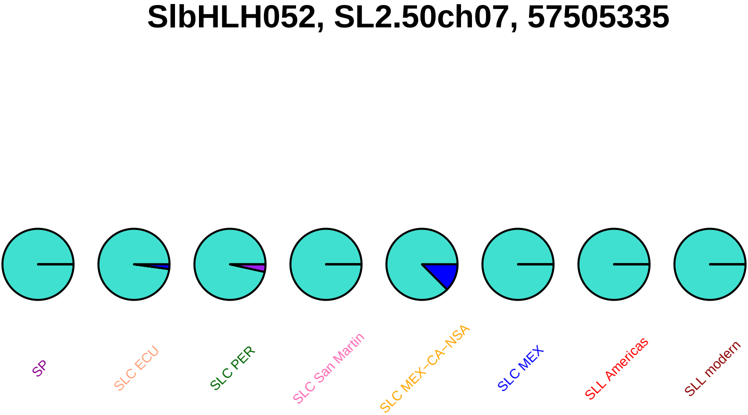
SlbHLH052, SL2.50ch07, 57505335
SP
SLC ECU
SLC PER
SLC San Martin
SLC MEX−CA−NSA
SLC MEX
SLL Americas
SLL modern

### Sl−ARF6B, SL2.50ch07, 57561325

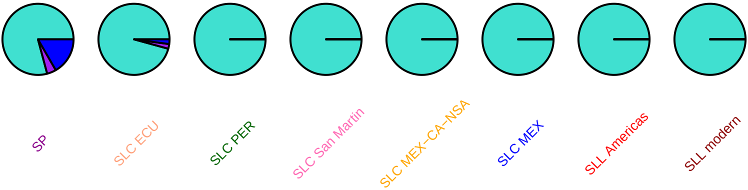

### Sl−ARF6B, SL2.50ch07, 57562147

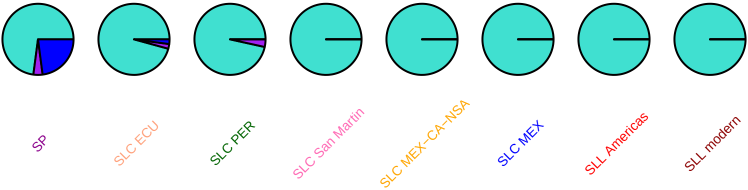

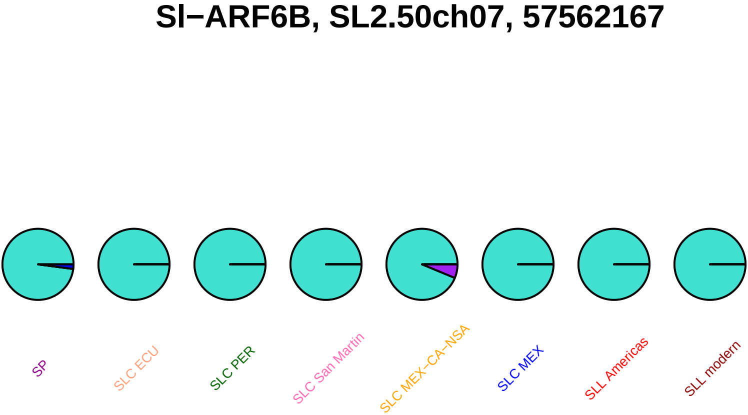

Sl−ARF6B, SL2.50ch07, 57562167
SP
SLC ECU
SLC PER
SLC San Martin
SLC MEX−CA−NSA
SLC MEX
SLL Americas
SLL modern

### Sl−ARF6B, SL2.50ch07, 57563981

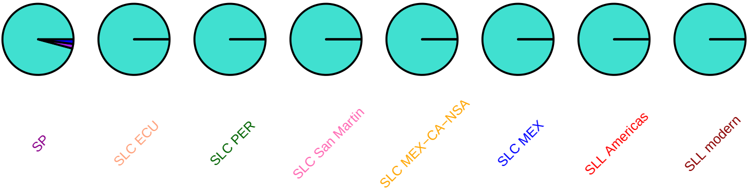

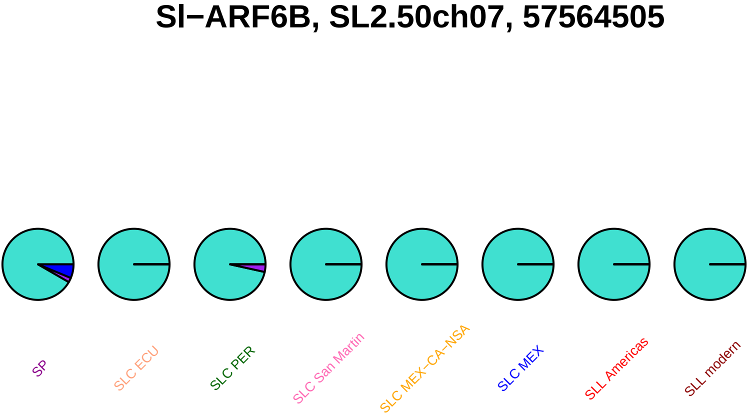
Sl−ARF6B, SL2.50ch07, 57564505
SP
SLC ECU
SLC PER
SLC San Martin
SLC MEX−CA−NSA
SLC MEX
SLL Americas
SLL modern

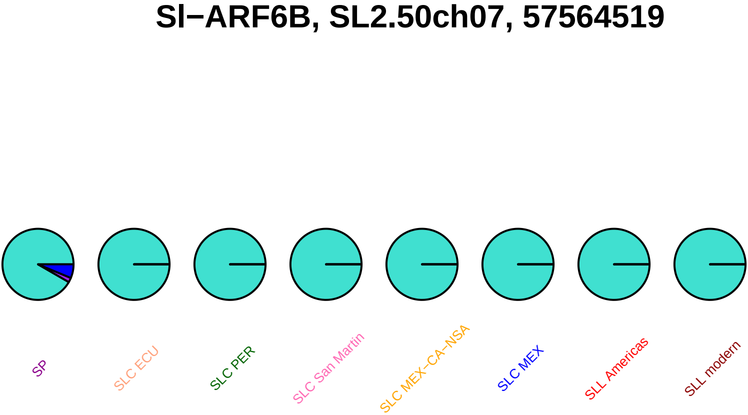
Sl−ARF6B, SL2.50ch07, 57564519
SP
SLC ECU
SLC PER
SLC San Martin
SLC MEX−CA−NSA
SLC MEX
SLL Americas
SLL modern

#### Sl−ARF6B, SL2.50ch07, 57565451

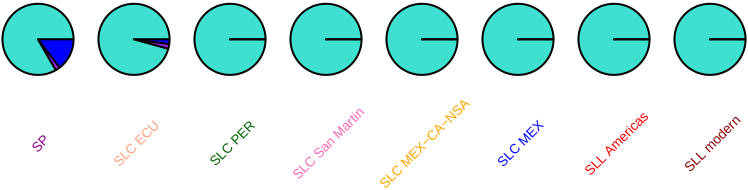

### Sl−ARF6B, SL2.50ch07, 57566479

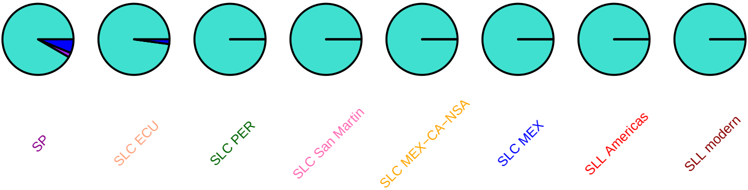

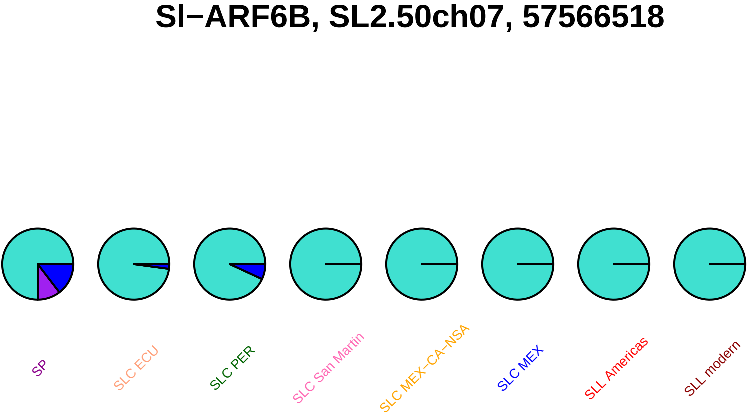
Sl−ARF6B, SL2.50ch07, 57566518
SP
SLC ECU
SLC PER
SLC San Martin
SLC MEX−CA−NSA
SLC MEX
SLL Americas
SLL modern

### psbx, SL2.50ch07, 57922580

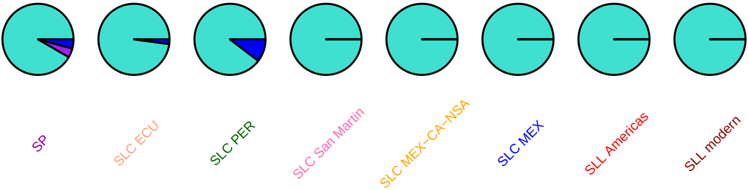

#### psbx, SL2.50ch07, 57922737

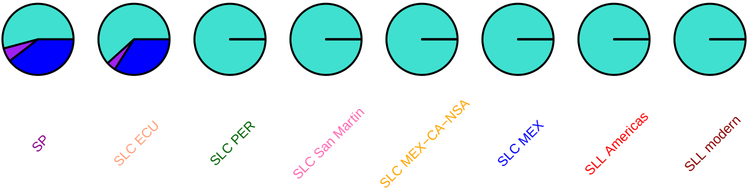

#### nml2, SL2.50ch07, 58027697

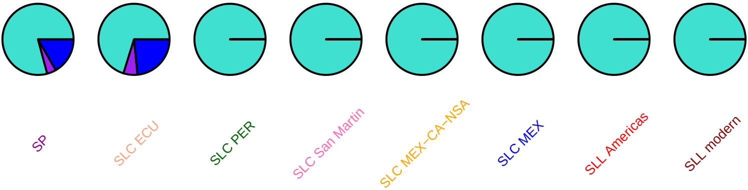

#### SlAGO6, SL2.50ch07, 59778345

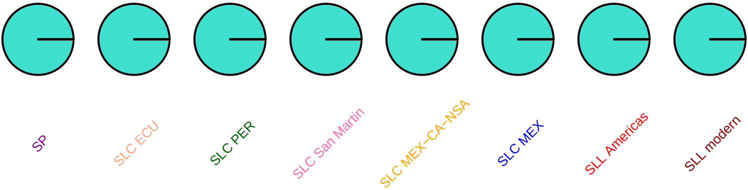

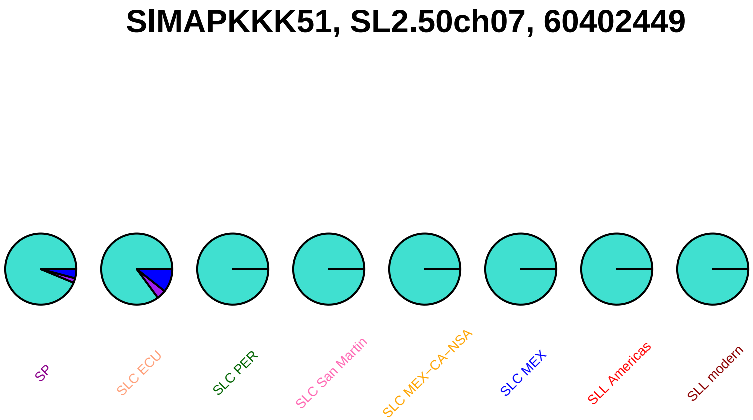
SlMAPKKK51, SL2.50ch07, 60402449
SP
SLC ECU
SLC PER
SLC San Martin
SLC MEX–CA–NSA
SLC MEX
SLL Americas
SLL modern

#### SlMAPKKK54, SL2.50ch07, 60469242

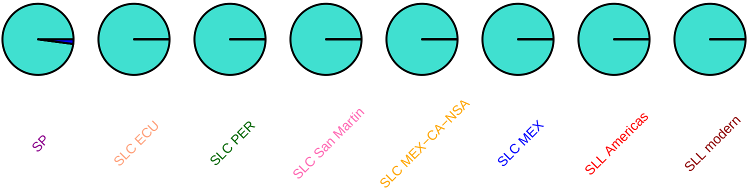

### SlMAPKKK55, SL2.50ch07, 60475184

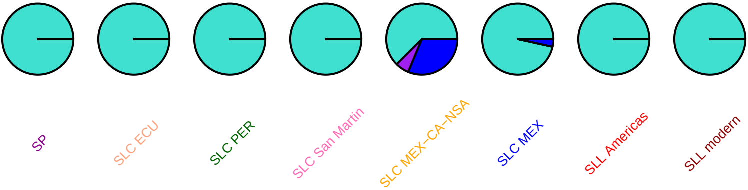

#### SlMYB101, SL2.50ch07, 60797144

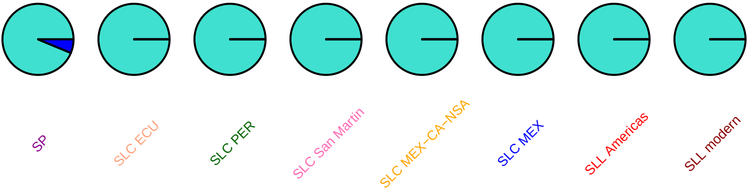

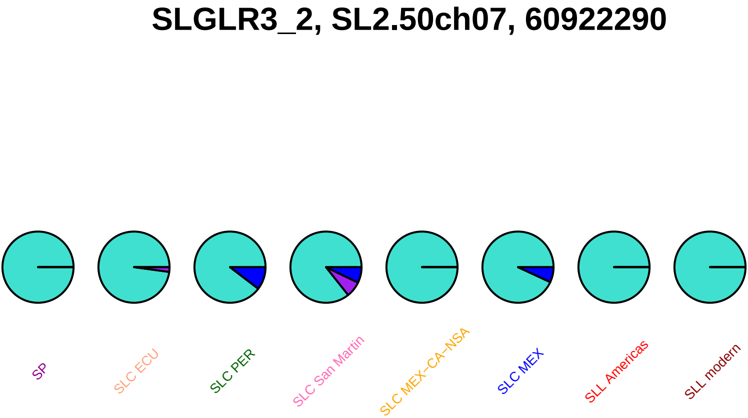
SLGLR3_2, SL2.50ch07, 60922290
SP
SLC ECU
SLC PER
SLC San Martin
SLC MEX−CA−NSA
SLC MEX
SLL Americas
SLL modern

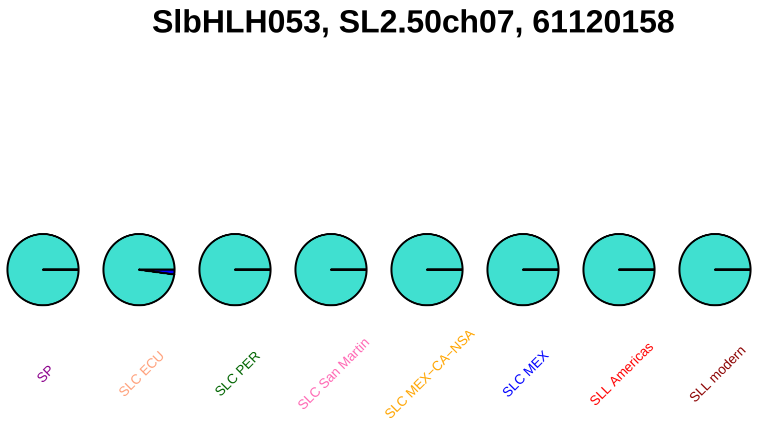
SlbHLH053, SL2.50ch07, 61120158
SP
SLC ECU
SLC PER
SLC San Martin
SLC MEX–CA–NSA
SLC MEX
SLL Americas
SLL modern

## AY150043, SL2.50ch07, 61187653

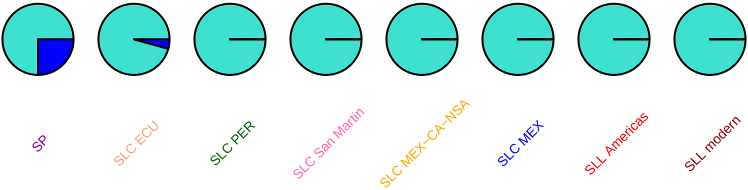

#### SCL32−like, SL2.50ch07, 61367535

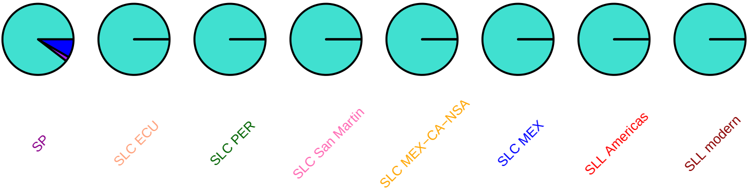

### SlMAPKKK56, SL2.50ch07, 61617841

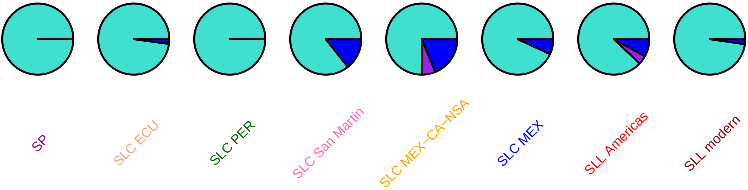

### tft9, SL2.50ch07, 61736577

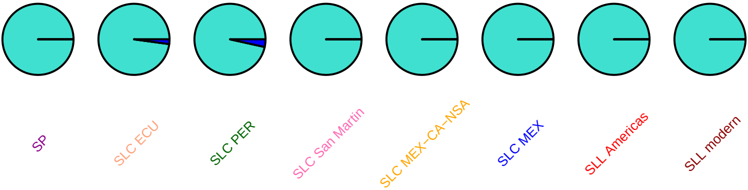

#### diagk1, SL2.50ch07, 63009658

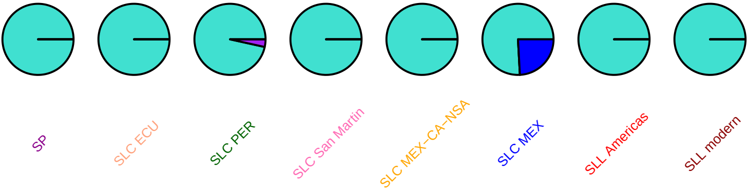

### SlCycD6_1, SL2.50ch07, 63103490

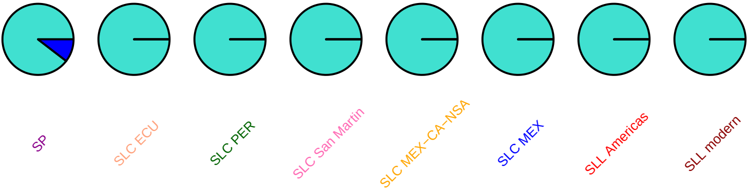

#### SlMYB85, SL2.50ch07, 63136052

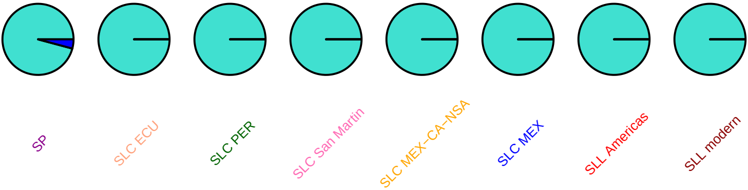

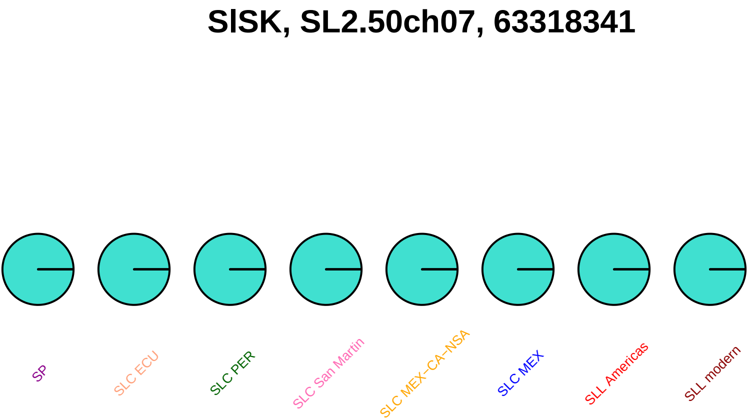
SlSK, SL2.50ch07, 63318341
SP
SLC ECU
SLC PER
SLC San Martin
SLC MEX−CA−NSA
SLC MEX
SLL Americas
SLL modern

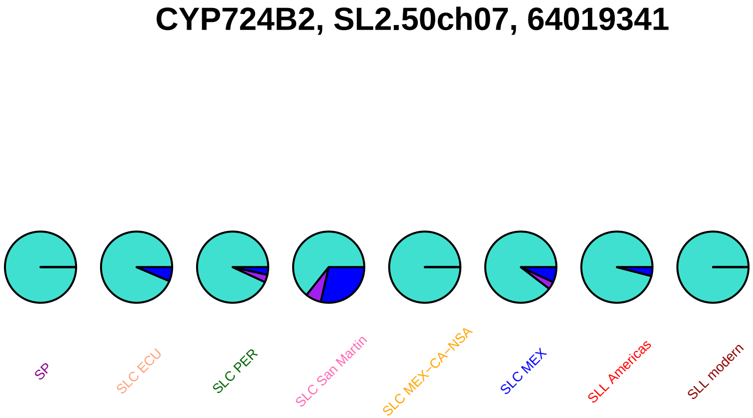
CYP724B2, SL2.50ch07, 64019341
SP
SLC ECU
SLC PER
SLC San Martin
SLC MEX−CA−NSA
SLC MEX
SLL Americas
SLL modern

#### mapk12, SL2.50ch07, 64227380

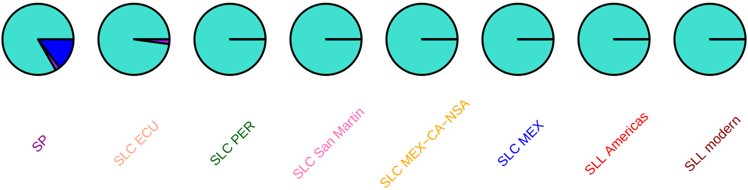

### bigst−gpx, SL2.50ch07, 64299909

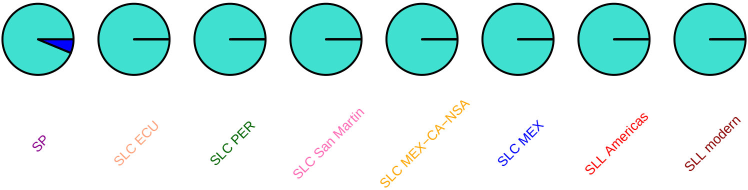

## ga2ox−4, SL2.50ch07, 64681673

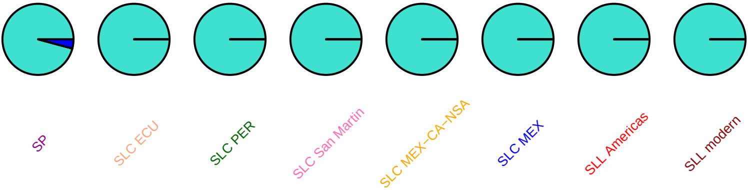

# ga2ox−5, SL2.50ch07, 64694384

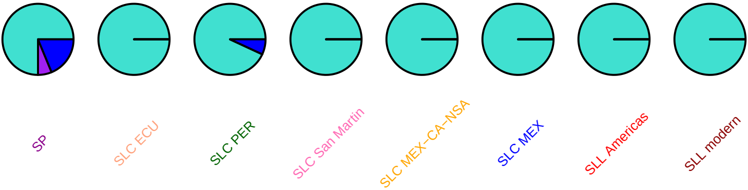

### GAME7, SL2.50ch07, 65223487

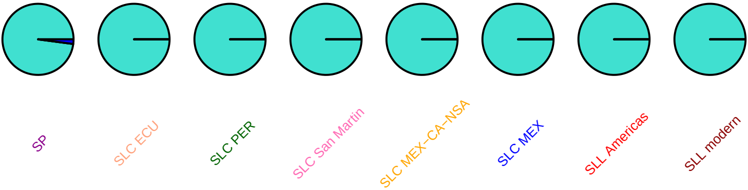

### gras2, SL2.50ch07, 66265139

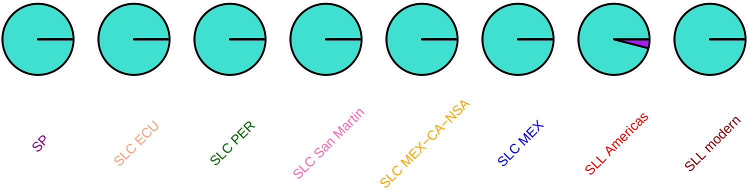

#### gras2, SL2.50ch07, 66265874

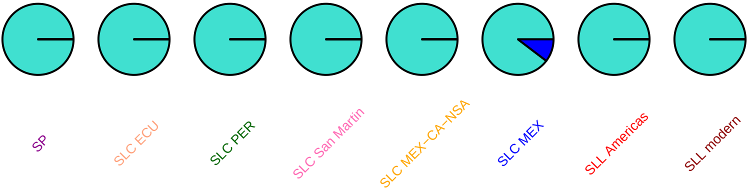

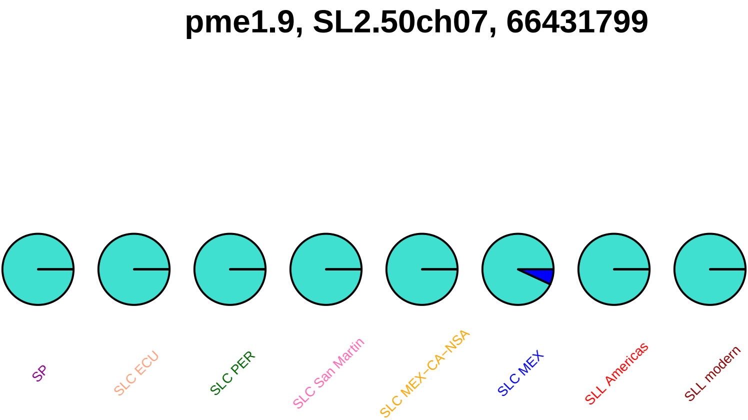
pme1.9, SL2.50ch07, 66431799
SP
SLC ECU
SLC PER
SLC San Martin
SLC MEX–CA–NSA
SLC MEX
SLL Americas
SLL modern

### pme1.9, SL2.50ch07, 66431895

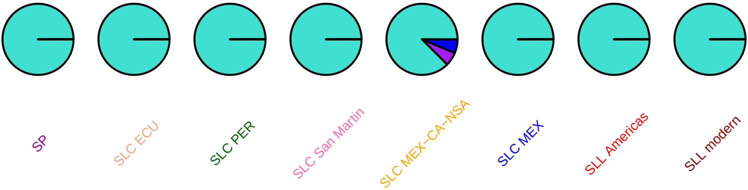

#### pme1.9, SL2.50ch07, 66431918

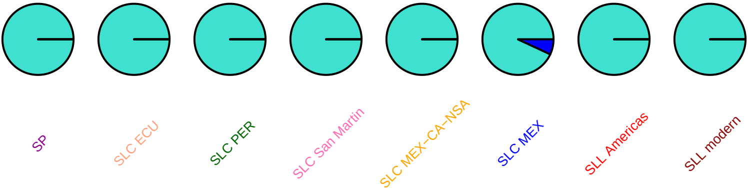

#### pme2.1, SL2.50ch07, 66440708

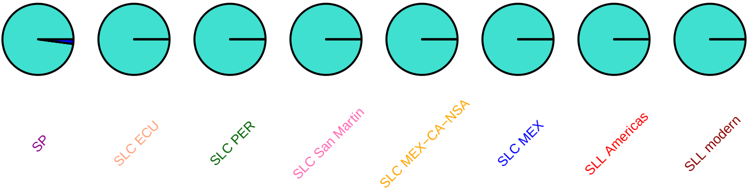

### pme2.1, SL2.50ch07, 66440783

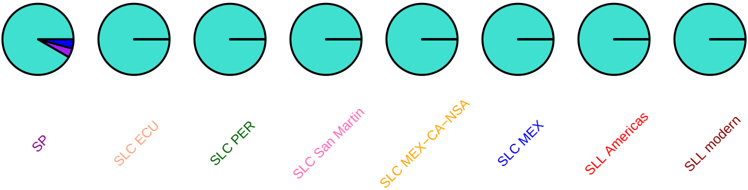

#### pme2.1, SL2.50ch07, 66440995

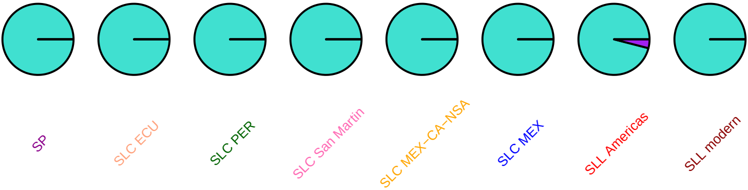

### SlMAPKKK59, SL2.50ch07, 66806317

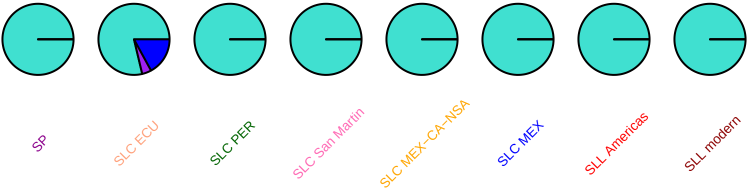

### SlWRKY18, SL2.50ch07, 67095543

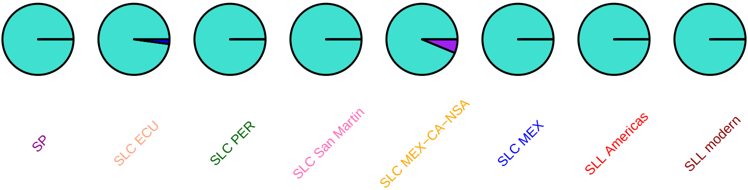

### SlC3H51, SL2.50ch07, 67109117

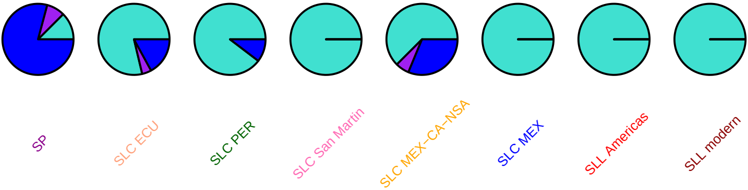

### aox1c, SL2.50ch08, 409946

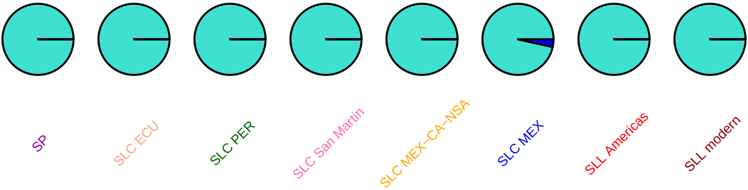

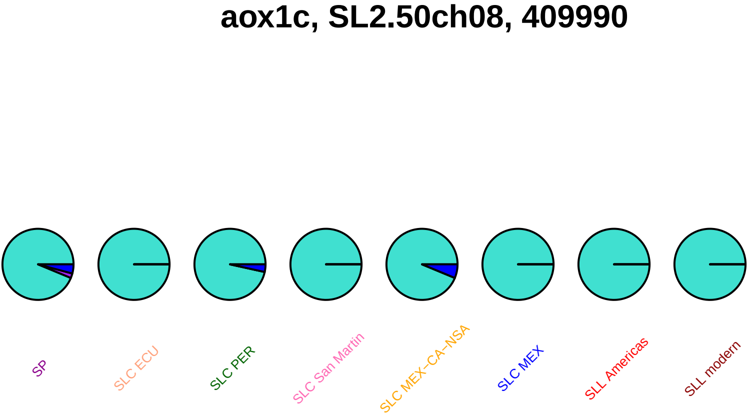
aox1c, SL2.50ch08, 409990
SP
SLC ECU
SLC PER
SLC San Martin
SLC MEX–CA–NSA
SLC MEX
SLL Americas
SLL modern

### aox1c, SL2.50ch08, 410199

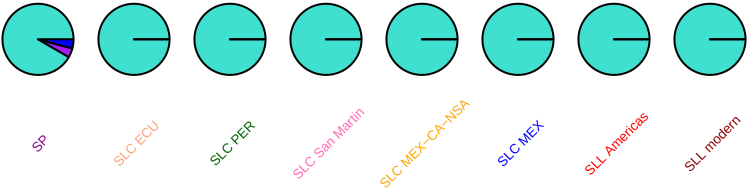

#### aox1c, SL2.50ch08, 410334

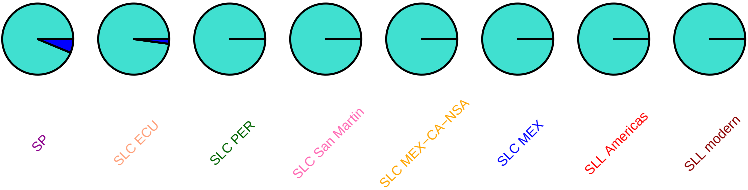

### xth5, SL2.50ch08, 479147

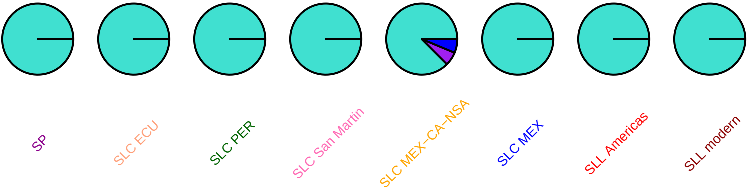

### xth5, SL2.50ch08, 479324

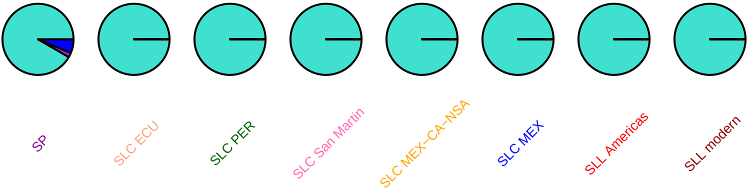

### xth5, SL2.50ch08, 479393

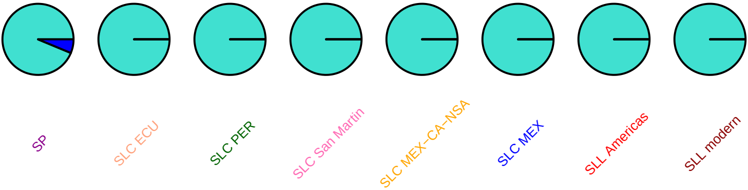

## AY534531, SL2.50ch08, 604233

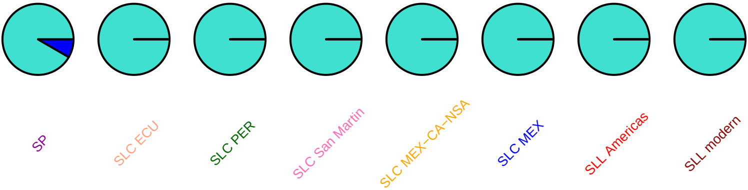

### hmgs, SL2.50ch08, 2296369

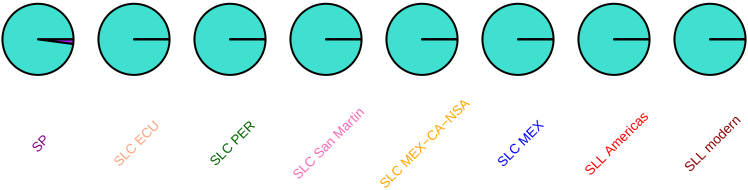

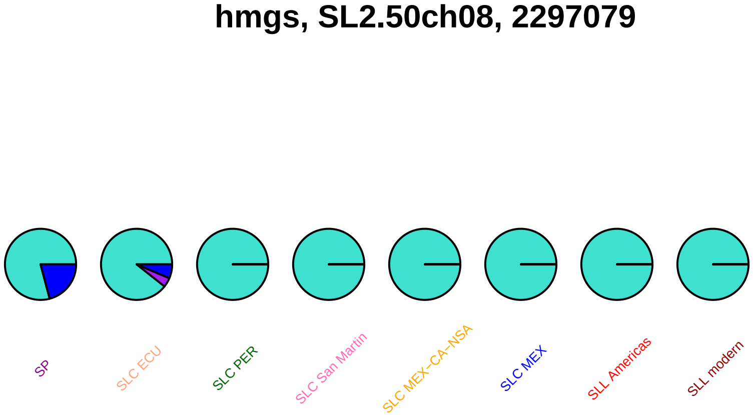
hmgs, SL2.50ch08, 2297079
SP
SLC ECU
SLC PER
SLC San Martin
SLC MEX−CA−NSA
SLC MEX
SLL Americas
SLL modern

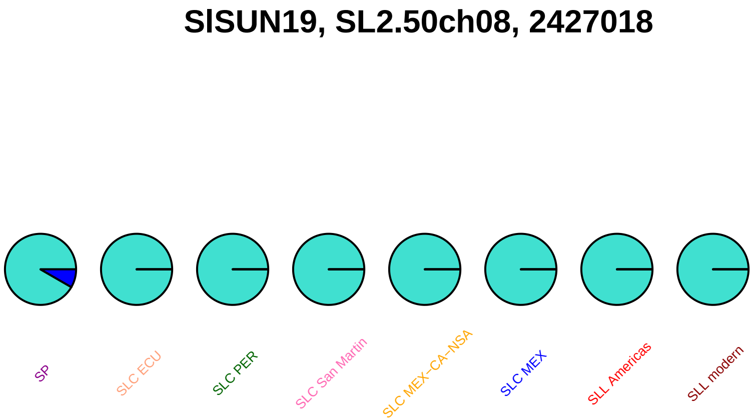

SlSUN19, SL2.50ch08, 2427018
SP
SLC ECU
SLC PER
SLC San Martin
SLC MEX−CA−NSA
SLC MEX
SLL Americas
SLL modern

### SlSUN20, SL2.50ch08, 2436799

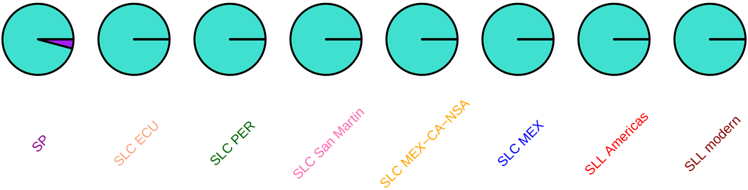

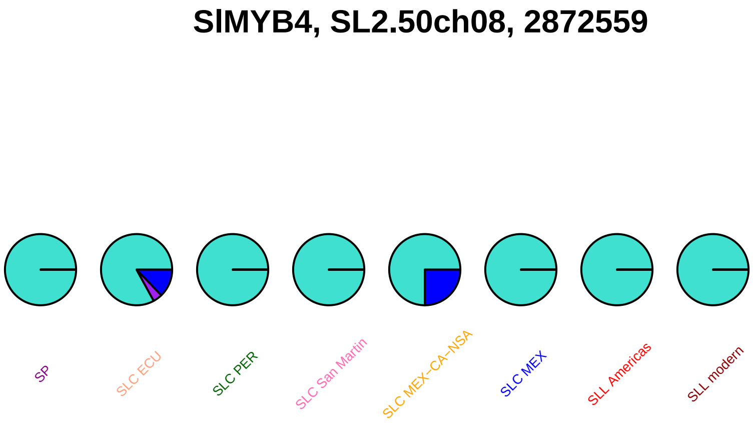
SlMYB4, SL2.50ch08, 2872559
SP
SLC ECU
SLC PER
SLC San Martin
SLC MEX–CA–NSA
SLC MEX
SLL Americas
SLL modern

### SlbHLH055, SL2.50ch08, 2998472

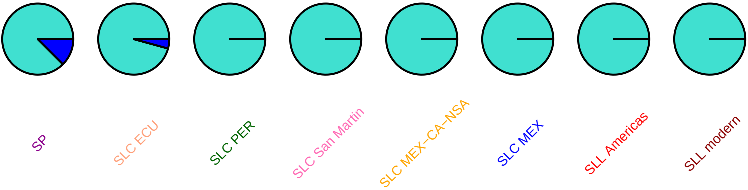

### loxa, SL2.50ch08, 3518092

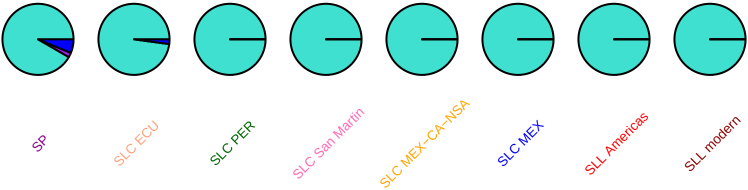

### SlTCP29, SL2.50ch08, 13379701

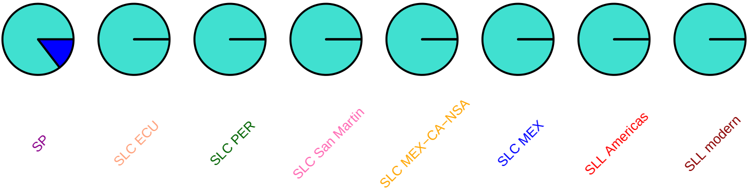

### SlINT2, SL2.50ch08, 14318782

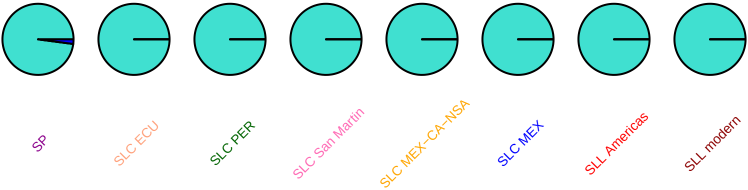

#### SlINT2, SL2.50ch08, 14318971

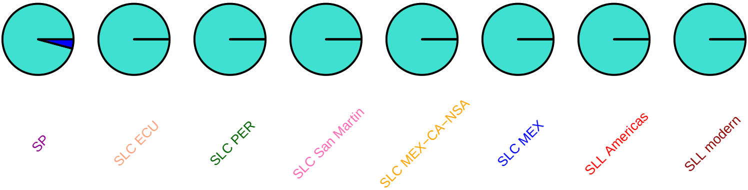

#### SlINT2, SL2.50ch08, 14320333

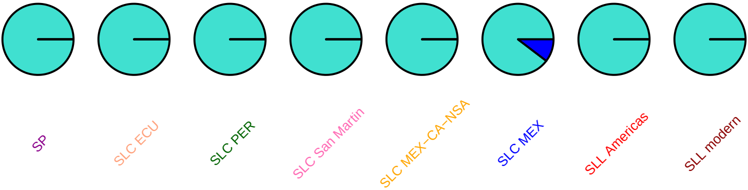

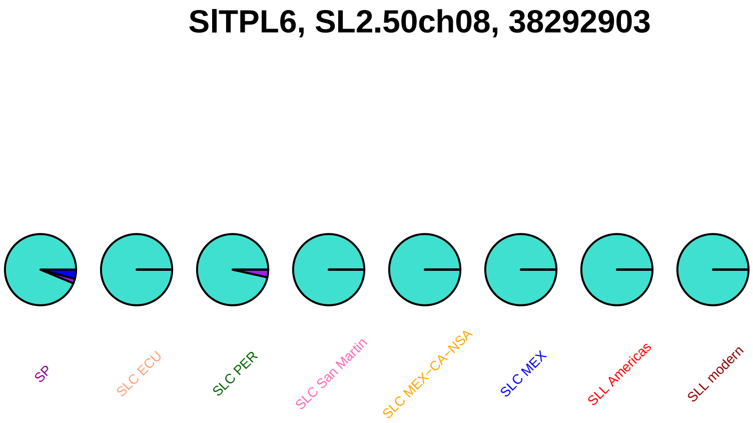
SlTPL6, SL2.50ch08, 38292903
SP
SLC ECU
SLC PER
SLC San Martin
SLC MEX−CA−NSA
SLC MEX
SLL Americas
SLL modern

#### SlTPL6, SL2.50ch08, 38293045

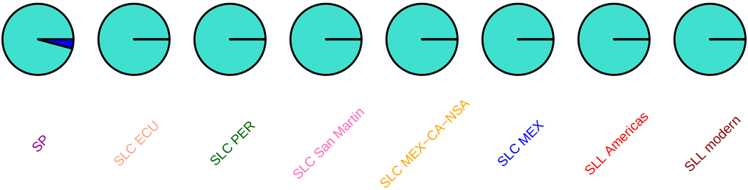

### SlTPL6, SL2.50ch08, 38293063

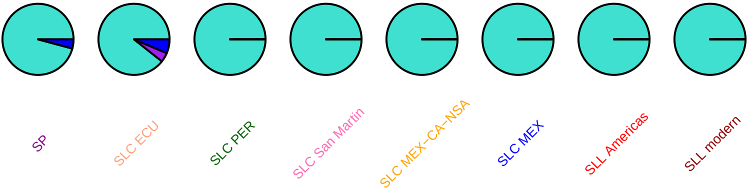

#### SlbHLH089, SL2.50ch08, 51900158

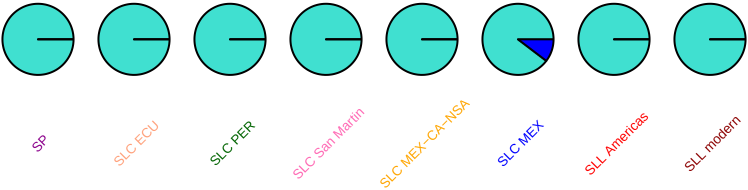

## AJ270963, SL2.50ch08, 53086023

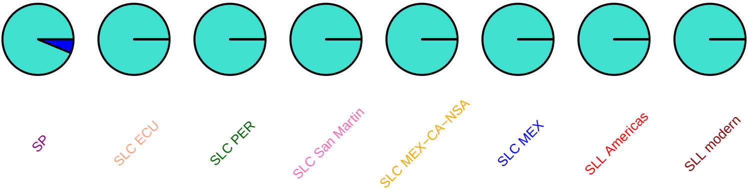

#### tpld, SL2.50ch08, 55661594

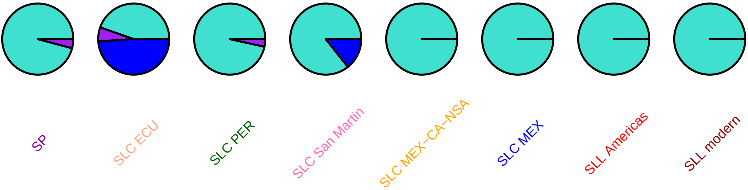

### SlMETL, SL2.50ch08, 55992921

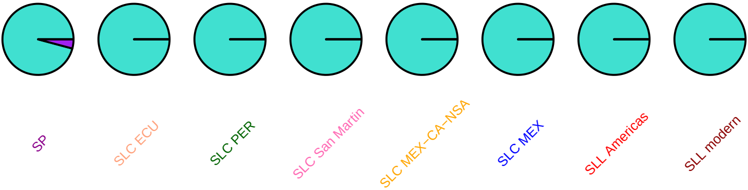

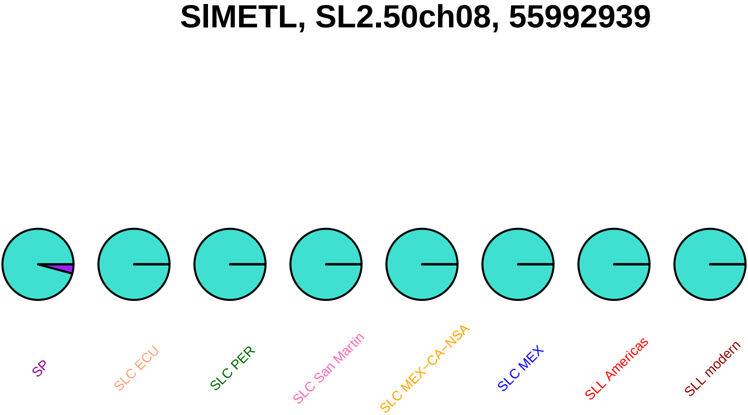
SlMETL, SL2.50ch08, 55992939
SP
SLC ECU
SLC PER
SLC San Martin
SLC MEX−CA−NSA
SLC MEX
SLL Americas
SLL modern

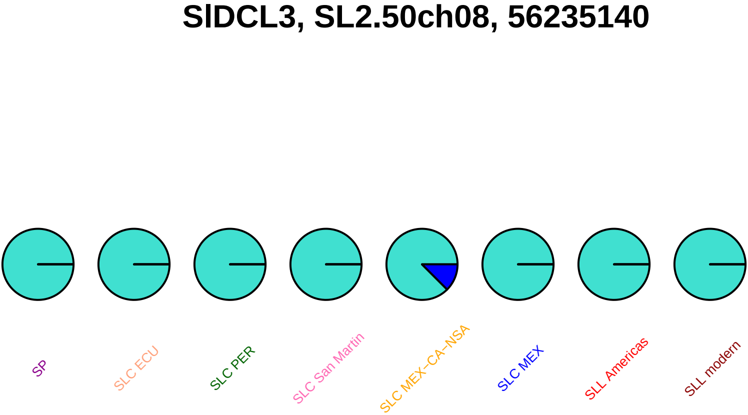
SlDCL3, SL2.50ch08, 56235140
SP
SLC ECU
SLC PER
SLC San Martin
SLC MEX−CA−NSA
SLC MEX
SLL Americas
SLL modern

### ppo, SL2.50ch08, 58841288

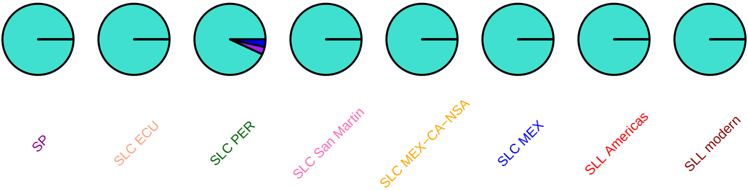

### aox1b, SL2.50ch08, 59716169

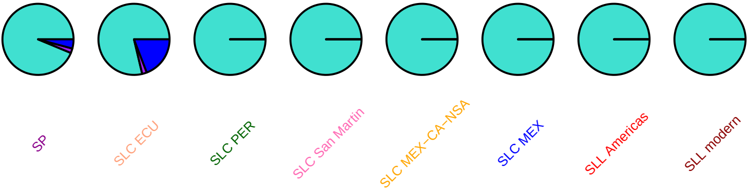

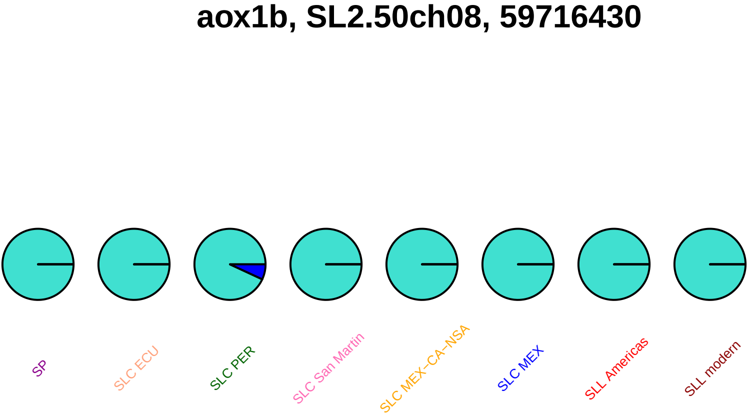
aox1b, SL2.50ch08, 59716430
SP
SLC ECU
SLC PER
SLC San Martin
SLC MEX−CA−NSA
SLC MEX
SLL Americas
SLL modern

### aox1b, SL2.50ch08, 59716576

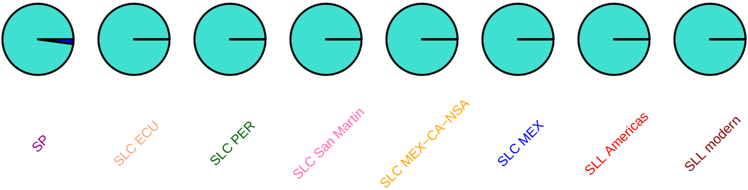

### aox1b, SL2.50ch08, 59718366

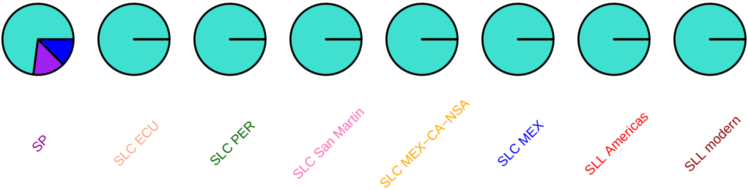

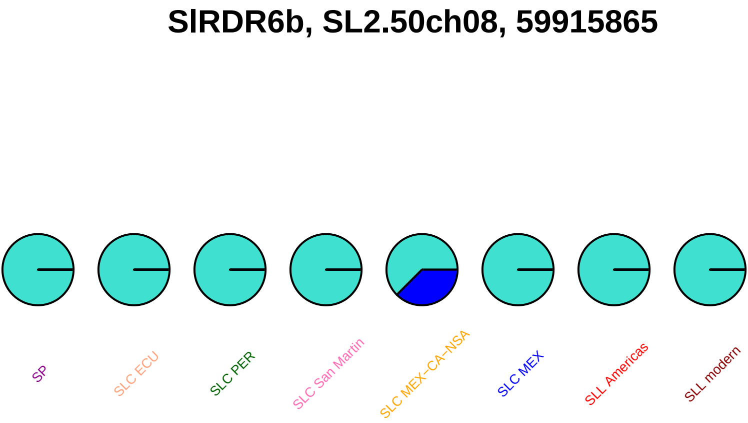
SlRDR6b, SL2.50ch08, 59915865
SP
SLC ECU
SLC PER
SLC San Martin
SLC MEX−CA−NSA
SLC MEX
SLL Americas
SLL modern

### SlRDR6b, SL2.50ch08, 59915904

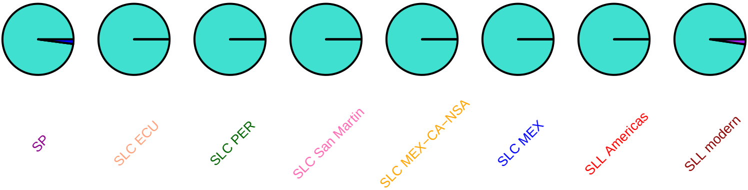

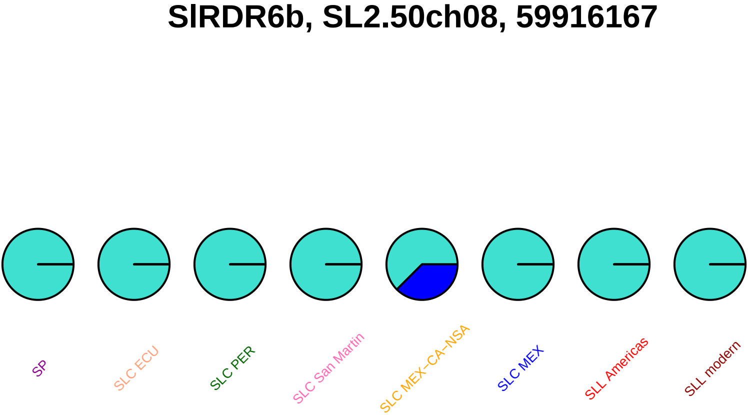
SlRDR6b, SL2.50ch08, 59916167
SP
SLC ECU
SLC PER
SLC San Martin
SLC MEX−CA−NSA
SLC MEX
SLL Americas
SLL modern

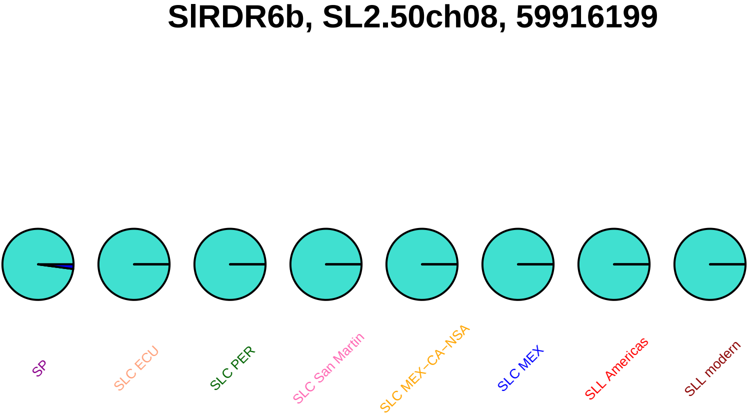

SlRDR6b, SL2.50ch08, 59916199
SP
SLC ECU
SLC PER
SLC San Martin
SLC MEX–CA–NSA
SLC MEX
SLL Americas
SLL modern

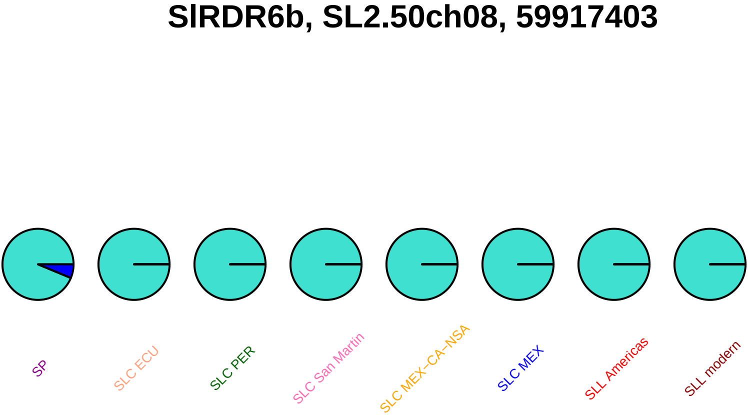

SlRDR6b, SL2.50ch08, 59917403
SP
SLC ECU
SLC PER
SLC San Martin
SLC MEX−CA−NSA
SLC MEX
SLL Americas
SLL modern

SlTPL2, SL2.50ch08, 60085623

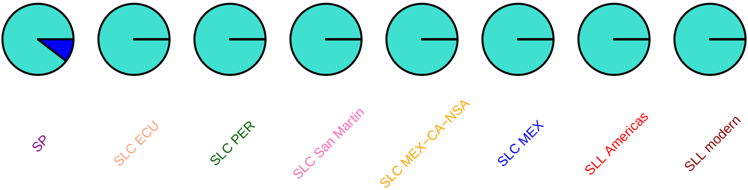

### SlTPL2, SL2.50ch08, 60086470

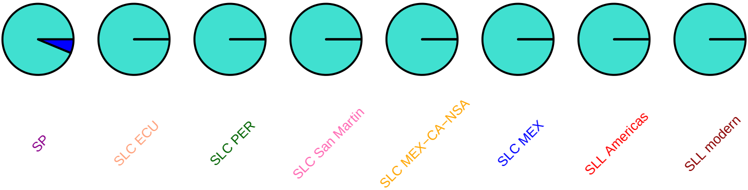

### gpat, SL2.50ch08, 60484720

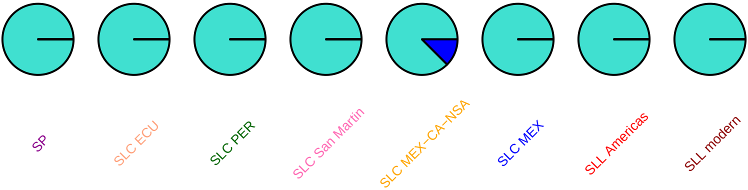

#### SlMYB39, SL2.50ch08, 60653408

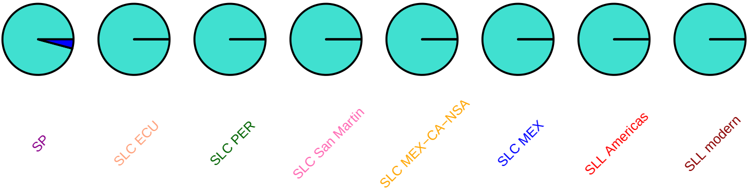

### SlbHLH146, SL2.50ch08, 60732402

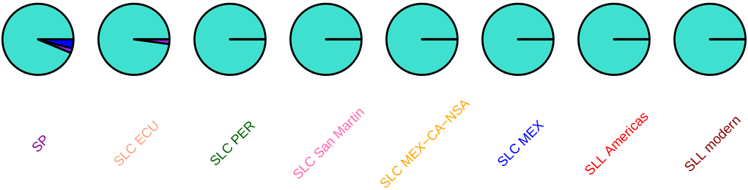

### SlbHLH146, SL2.50ch08, 60733309

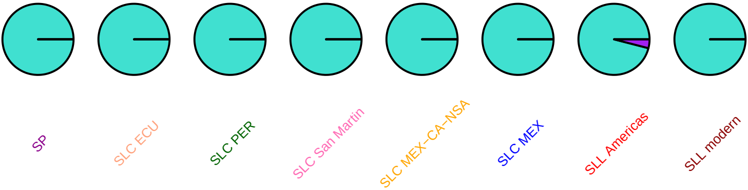

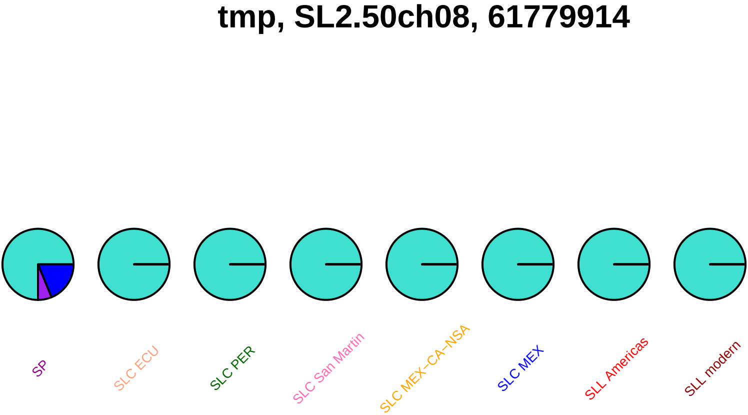
tmp, SL2.50ch08, 61779914
SP
SLC ECU
SLC PER
SLC San Martin
SLC MEX−CA−NSA
SLC MEX
SLL Americas
SLL modern

### lin9, SL2.50ch08, 62722977

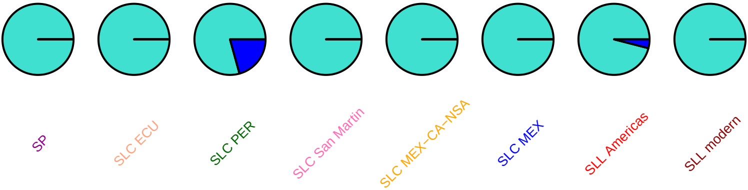

#### lin9, SL2.50ch08, 62722985

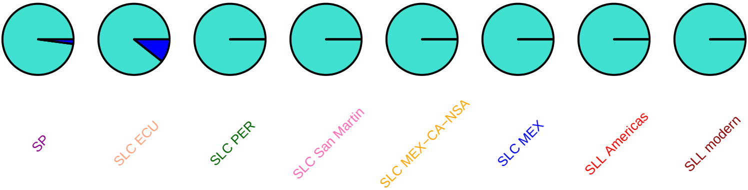

### ccs, SL2.50ch08, 63275938

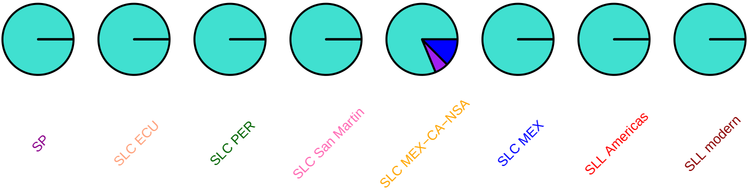

#### SlCCS52A, SL2.50ch08, 63411628

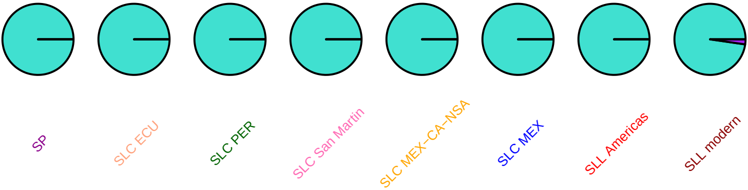

#### pldb1, SL2.50ch08, 63471763

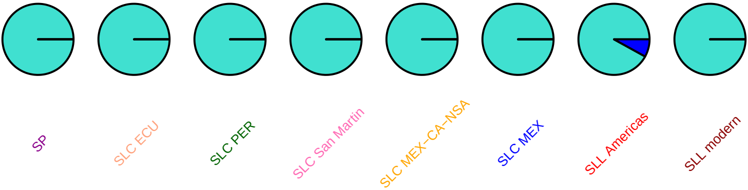

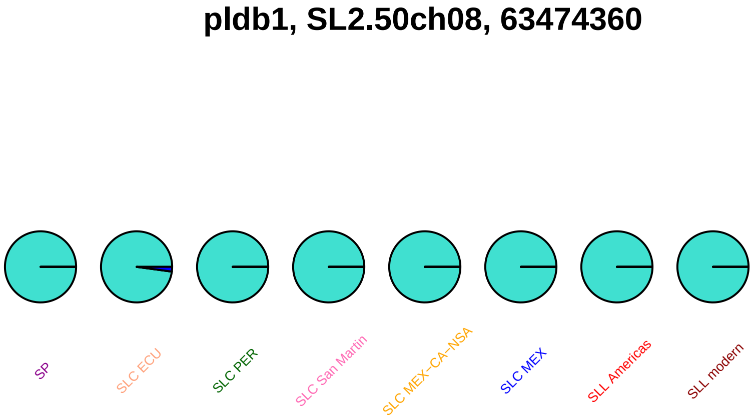
pldb1, SL2.50ch08, 63474360
SP
SLC ECU
SLC PER
SLC San Martin
SLC MEX−CA−NSA
SLC MEX
SLL Americas
SLL modern

### SlSTP6, SL2.50ch08, 63616124

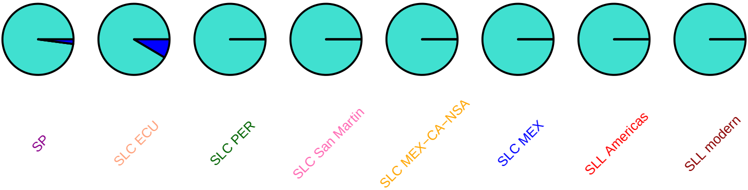

### SlSTP6, SL2.50ch08, 63616224

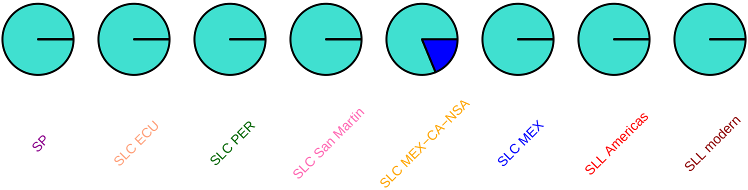

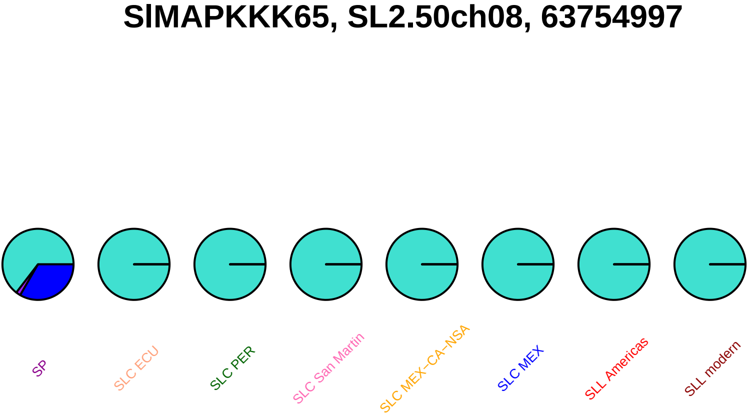
SlMAPKKK65, SL2.50ch08, 63754997
SP
SLC ECU
SLC PER
SLC San Martin
SLC MEX−CA−NSA
SLC MEX
SLL Americas
SLL modern

### SlMAPKKK65, SL2.50ch08, 63755209

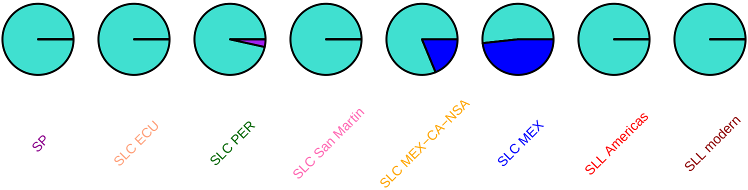

#### SlMAPKKK65, SL2.50ch08, 63755374

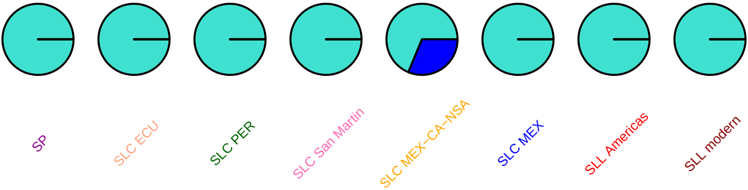

# pr5, SL2.50ch08, 63855225

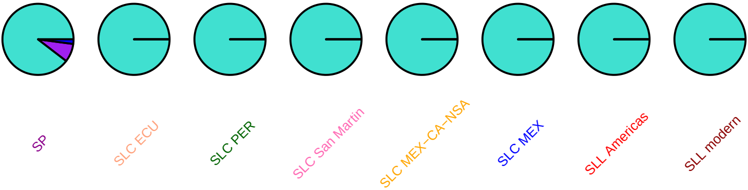

# pr5, SL2.50ch08, 63855260

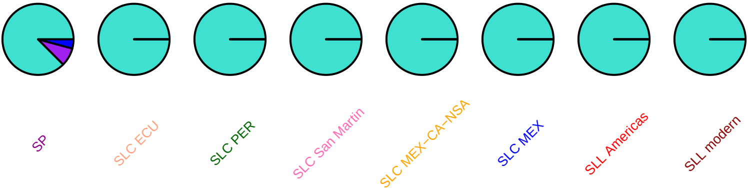

## pr5, SL2.50ch08, 63855502

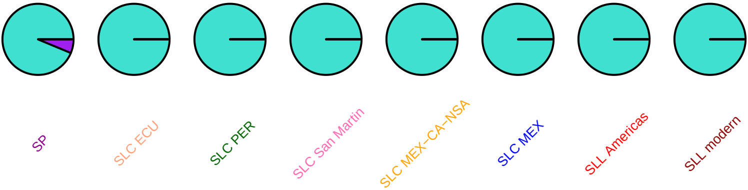

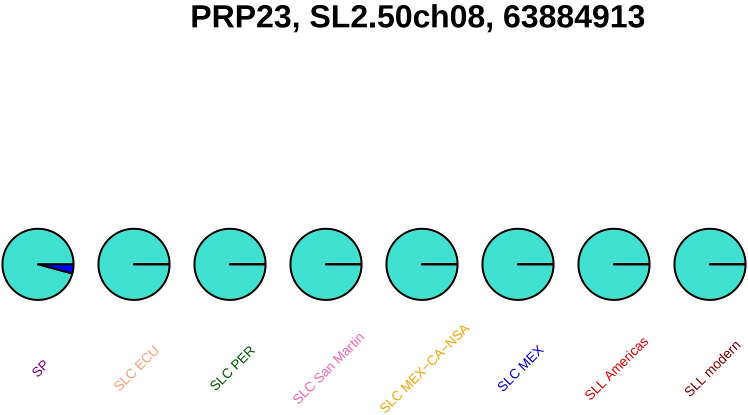
PRP23, SL2.50ch08, 63884913
SP
SLC ECU
SLC PER
SLC San Martin
SLC MEX–CA–NSA
SLC MEX
SLL Americas
SLL modern

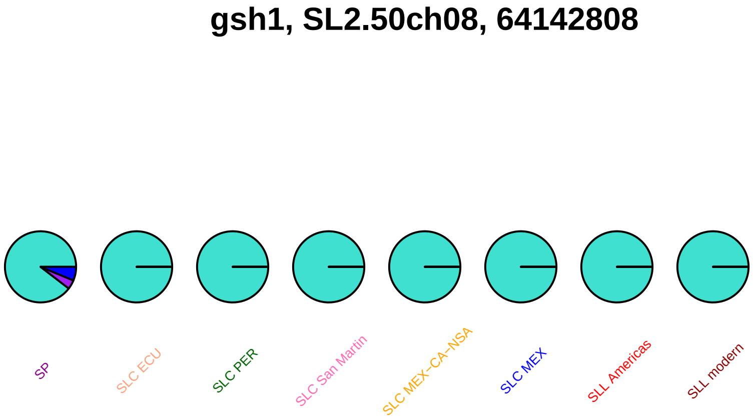
gsh1, SL2.50ch08, 64142808
SP
SLC ECU
SLC PER
SLC San Martin
SLC MEX–CA–NSA
SLC MEX
SLL Americas
SLL modern

#### gsh1, SL2.50ch08, 64147850

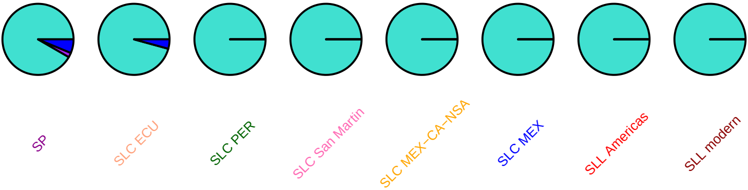

### SlbHLH090, SL2.50ch08, 64224338

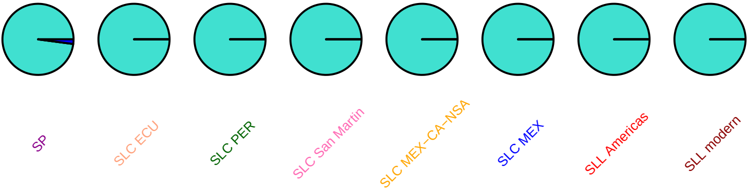

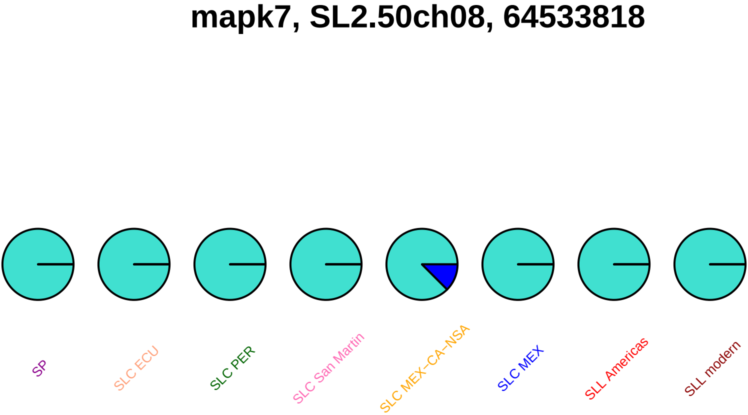
mapk7, SL2.50ch08, 64533818
SP
SLC ECU
SLC PER
SLC San Martin
SLC MEX−CA−NSA
SLC MEX
SLL Americas
SLL modern

### cel1, SL2.50ch08, 64614298

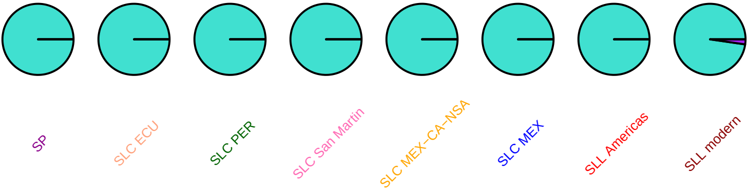

## AJ457975, SL2.50ch08, 64834358

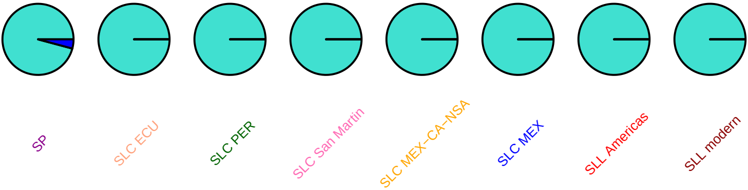

### cel8, SL2.50ch08, 65097223

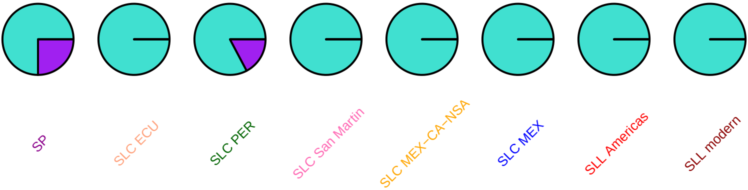

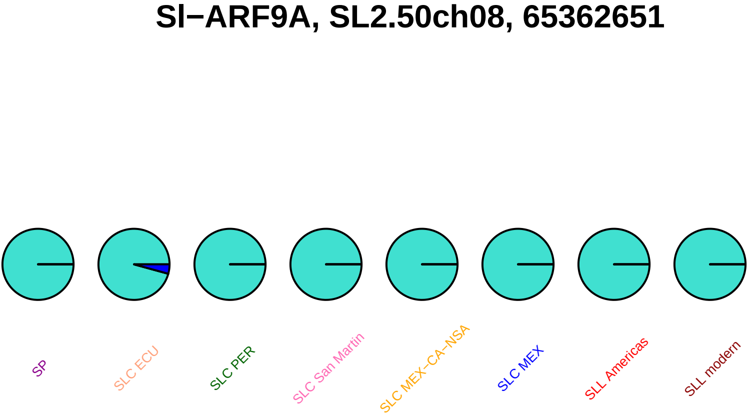
Sl−ARF9A, SL2.50ch08, 65362651
SP
SLC ECU
SLC PER
SLC San Martin
SLC MEX−CA−NSA
SLC MEX
SLL Americas
SLL modern

#### SlbHLH056, SL2.50ch08, 65712304

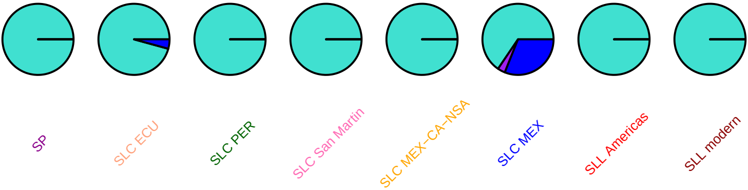

### SlbHLH056, SL2.50ch08, 65712919

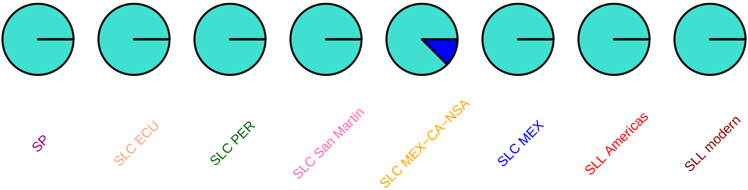

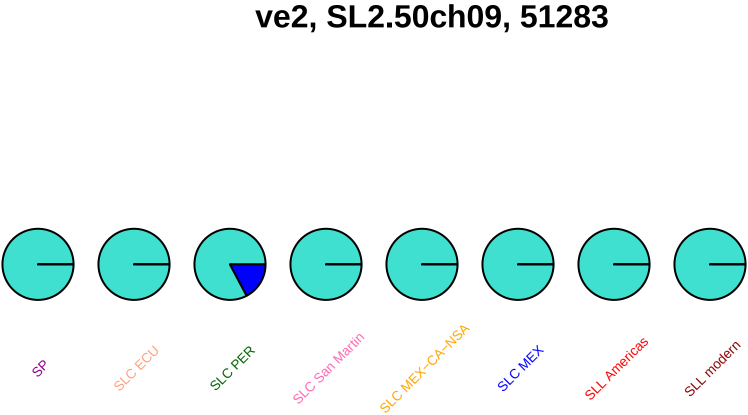
ve2, SL2.50ch09, 51283
SP
SLC ECU
SLC PER
SLC San Martin
SLC MEX−CA−NSA
SLC MEX
SLL Americas
SLL modern

### aci49, SL2.50ch09, 574114

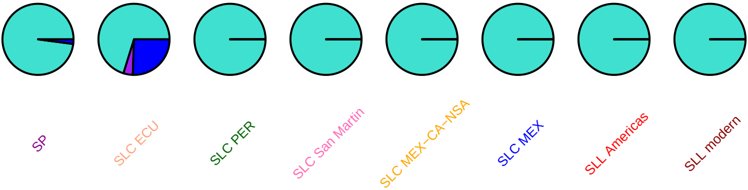

### SlSUN25, SL2.50ch09, 985619

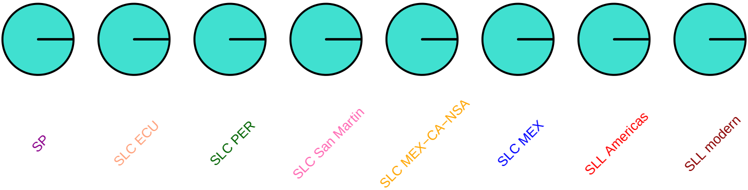

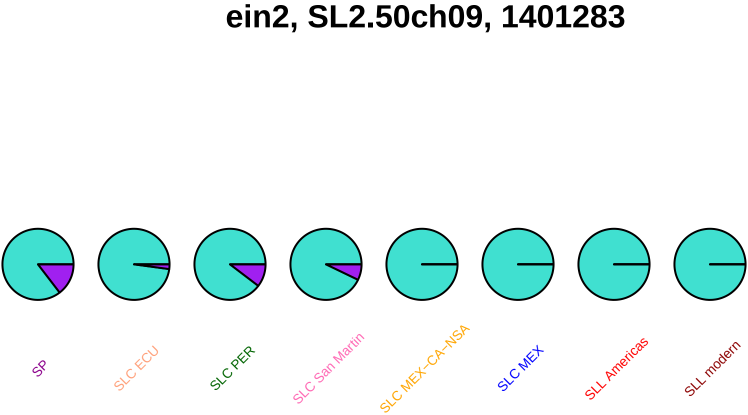

ein2, SL2.50ch09, 1401283
SP
SLC ECU
SLC PER
SLC San Martin
SLC MEX–CA–NSA
SLC MEX
SLL Americas
SLL modern

### ein2, SL2.50ch09, 1401301

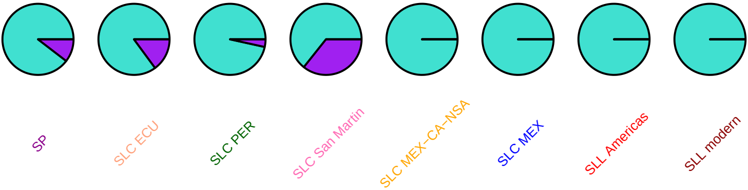

### ein2, SL2.50ch09, 1404092

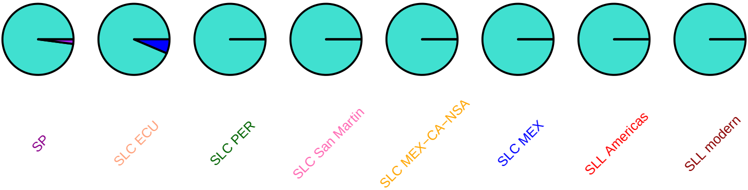

**ein2, SL2.50ch09, 1405090**

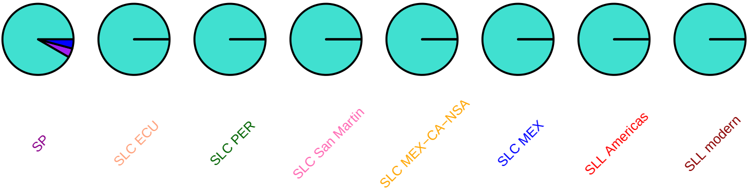

#### SlMYB22, SL2.50ch09, 1852405

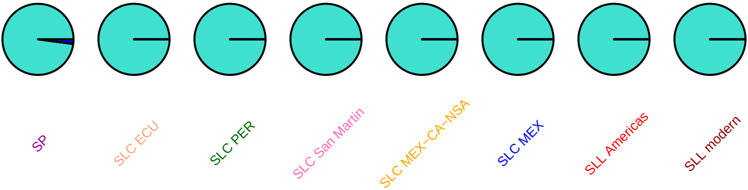

## td, SL2.50ch09, 2124260

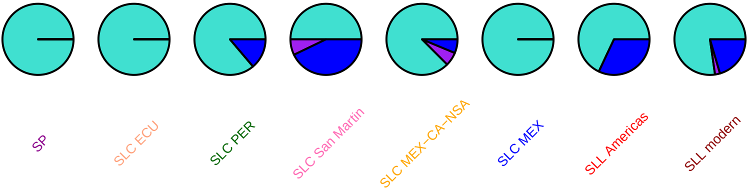

# td, SL2.50ch09, 2124965

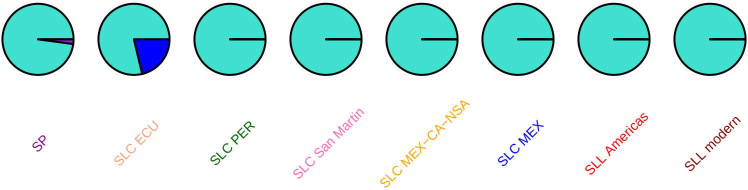

# er28, SL2.50ch09, 2373896

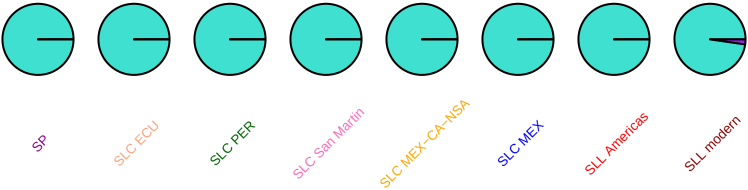

#### lin5, SL2.50ch09, 3477835

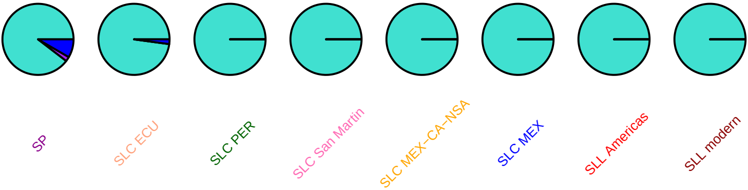

#### lin5, SL2.50ch09, 3478052

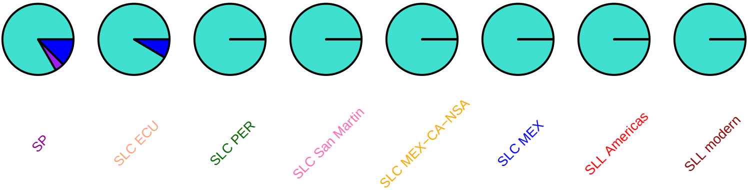

#### cel2, SL2.50ch09, 3600451

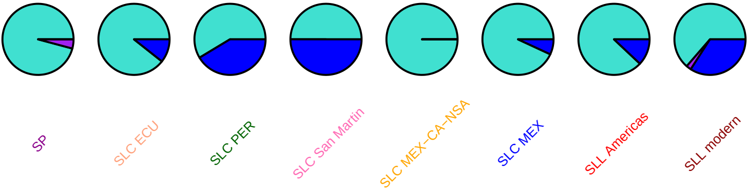

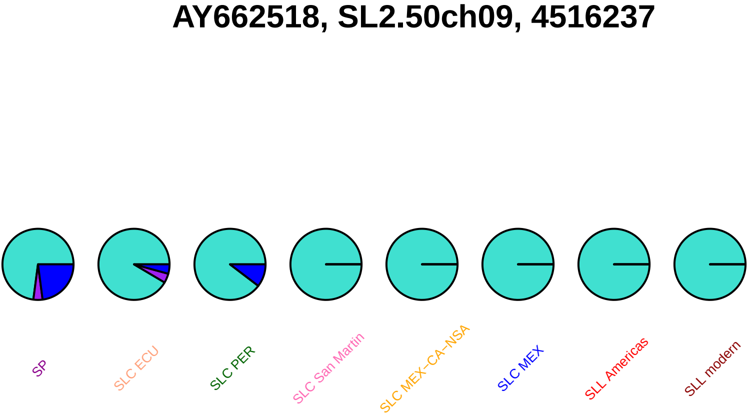
AY662518, SL2.50ch09, 4516237
SP
SLC ECU
SLC PER
SLC San Martin
SLC MEX–CA–NSA
SLC MEX
SLL Americas
SLL modern

### SlMYB1, SL2.50ch09, 5071287

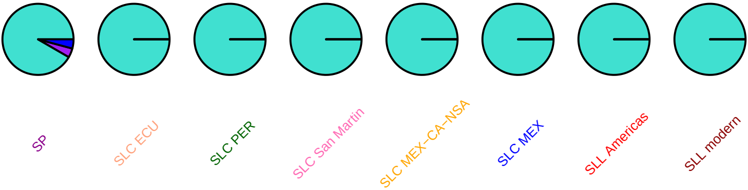

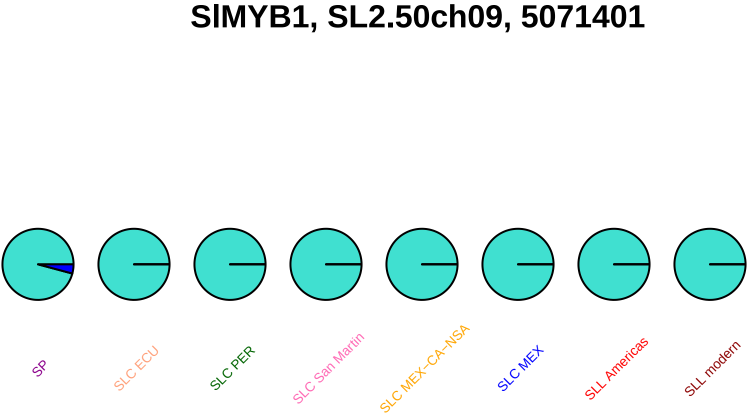
SlMYB1, SL2.50ch09, 5071401
SP
SLC ECU
SLC PER
SLC San Martin
SLC MEX–CA–NSA
SLC MEX
SLL Americas
SLL modern

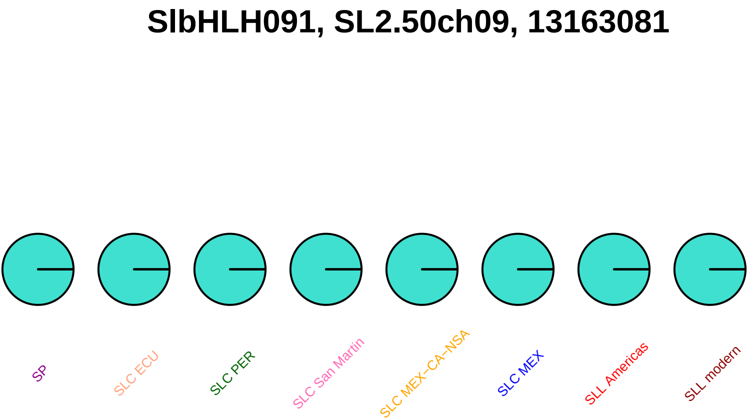
SlbHLH091, SL2.50ch09, 13163081
SP
SLC ECU
SLC PER
SLC San Martin
SLC MEX–CA–NSA
SLC MEX
SLL Americas
SLL modern

#### SlbHLH149, SL2.50ch09, 13309241

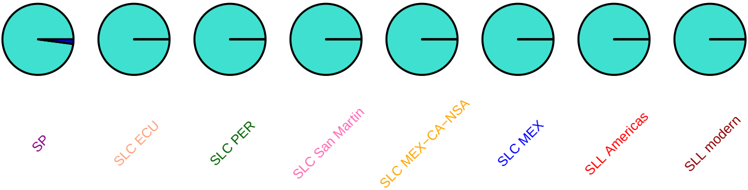

#### SlMAPKKK70, SL2.50ch09, 13406564

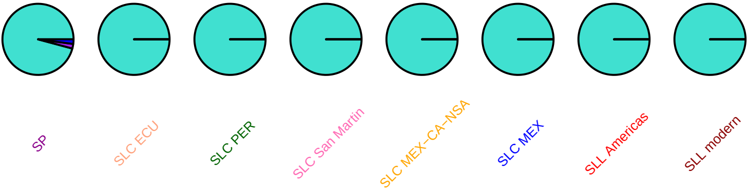

#### SlOFP17, SL2.50ch09, 13537267

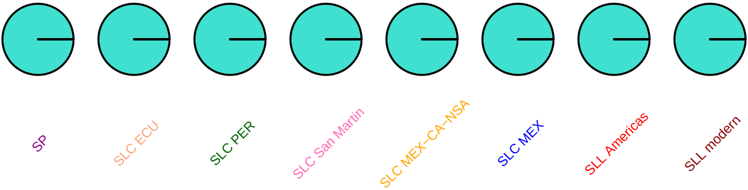

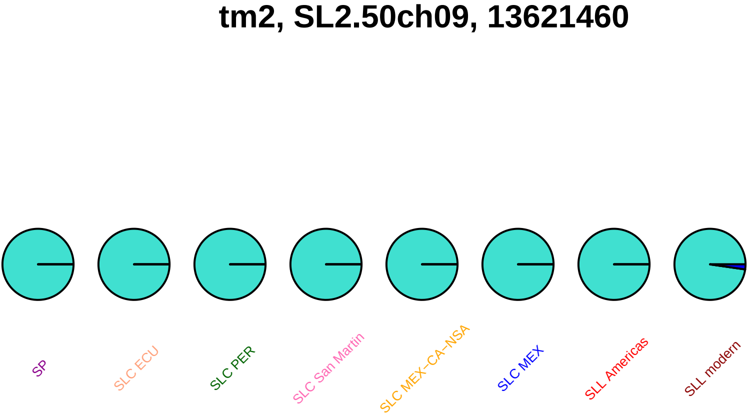

tm2, SL2.50ch09, 13621460
SP
SLC ECU
SLC PER
SLC San Martin
SLC MEX–CA–NSA
SLC MEX
SLL Americas
SLL modern

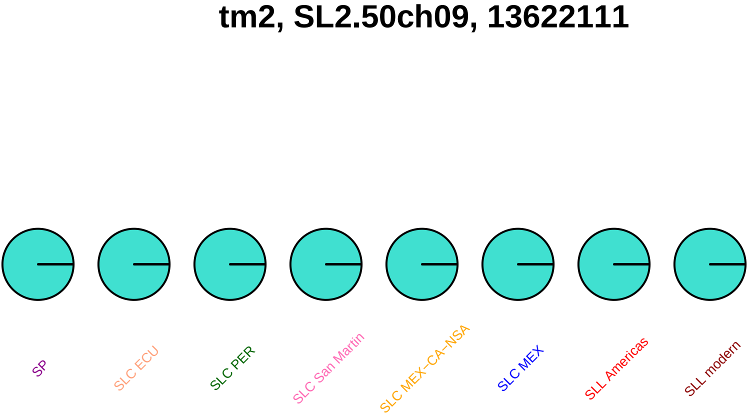

tm2, SL2.50ch09, 13622111
SP
SLC ECU
SLC PER
SLC San Martin
SLC MEX−CA−NSA
SLC MEX
SLL Americas
SLL modern

## tm2, SL2.50ch09, 13622534

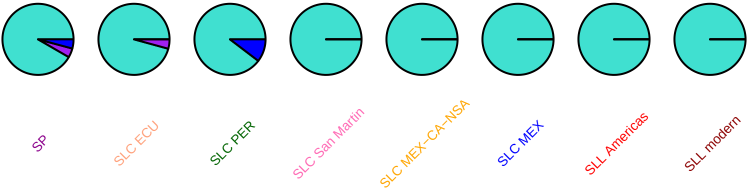

#### SolycHsfA8, SL2.50ch09, 54775599

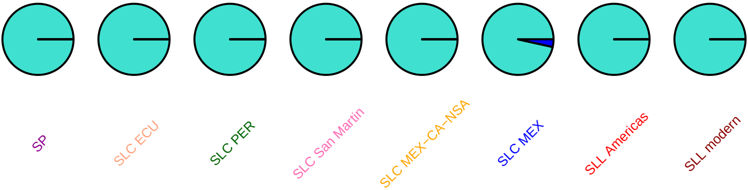

## AF106660, SL2.50ch09, 63110397

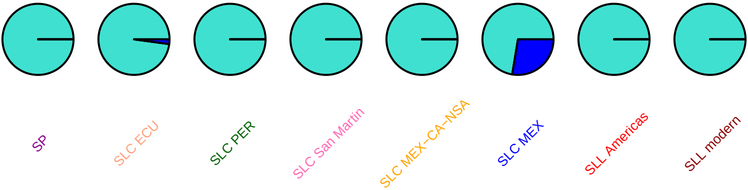

#### SlOFP18, SL2.50ch09, 63433071

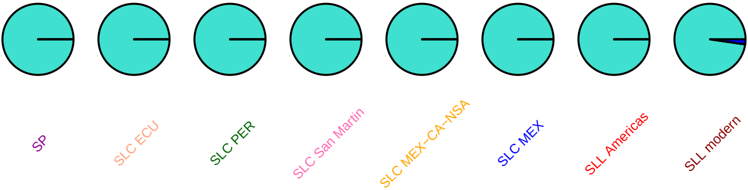

#### SLU20590, SL2.50ch09, 67088452

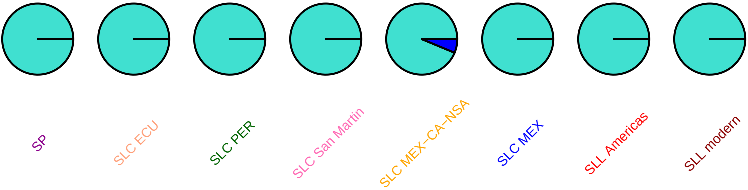

#### SlIAA14, SL2.50ch09, 68967925

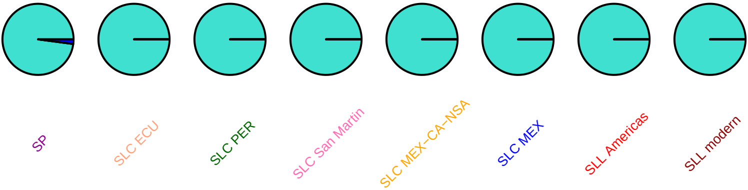

### SlbHLH061, SL2.50ch09, 69491218

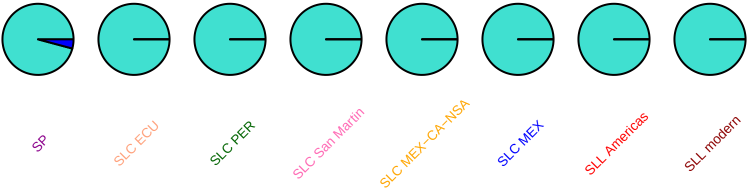

### fer1, SL2.50ch09, 69686671

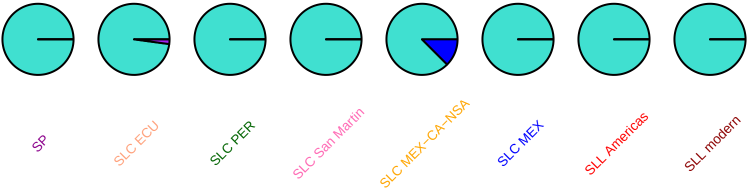

#### SlMYB79, SL2.50ch09, 70204898

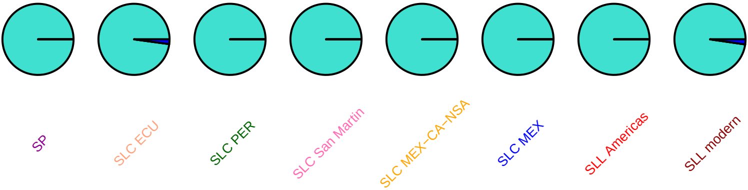

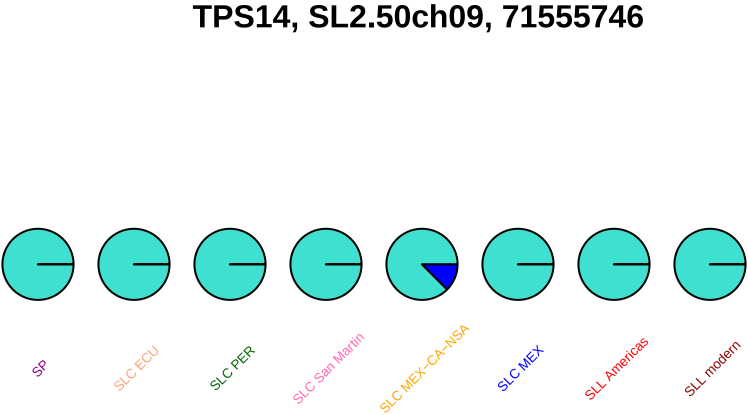

TPS14, SL2.50ch09, 71555746
SP
SLC ECU
SLC PER
SLC San Martin
SLC MEX−CA−NSA
SLC MEX
SLL Americas
SLL modern

### TPS14, SL2.50ch09, 71557185

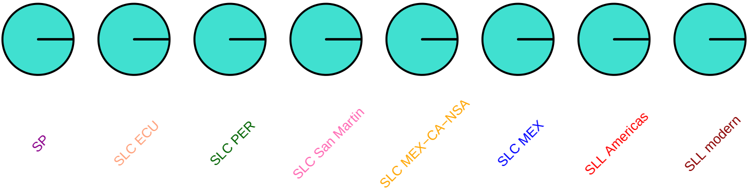

#### TPS14, SL2.50ch09, 71557724

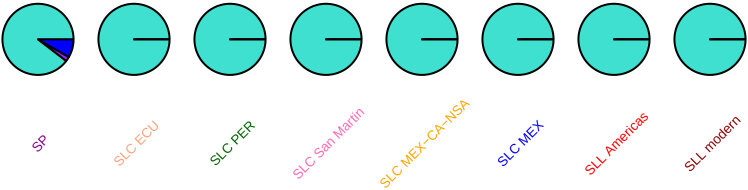

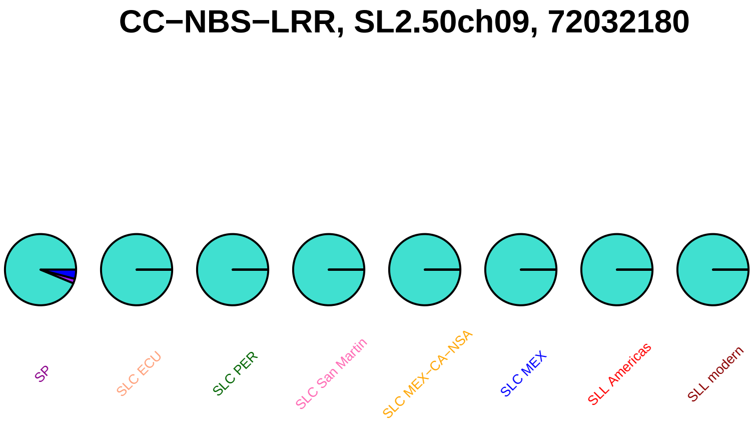
CC−NBS−LRR, SL2.50ch09, 72032180
SP
SLC ECU
SLC PER
SLC San Martin
SLC MEX−CA−NSA
SLC MEX
SLL Americas
SLL modern

### CC−NBS−LRR, SL2.50ch09, 72032583

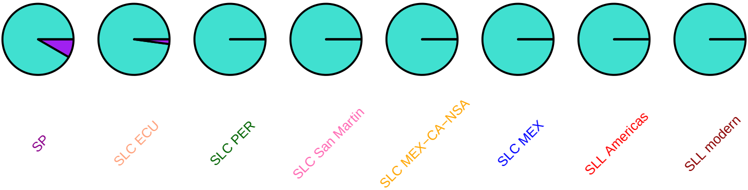

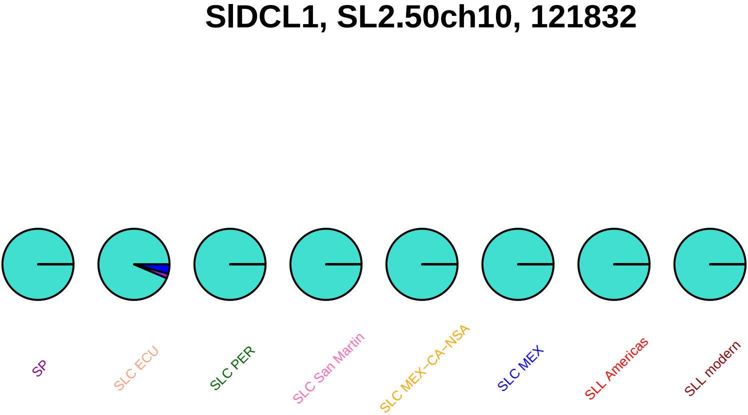

SlDCL1, SL2.50ch10, 121832
SP
SLC ECU
SLC PER
SLC San Martin
SLC MEX−CA−NSA
SLC MEX
SLL Americas
SLL modern

### SlbHLH153, SL2.50ch10, 1146092

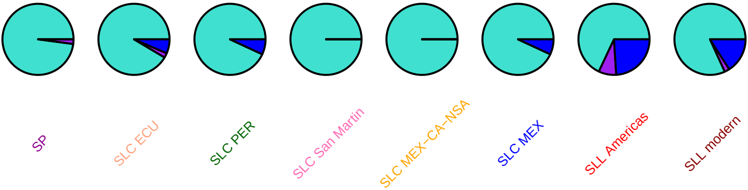

#### SlSUN28, SL2.50ch10, 2862101

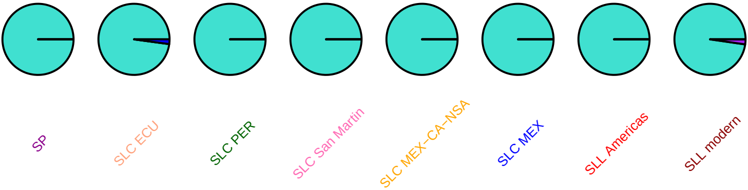

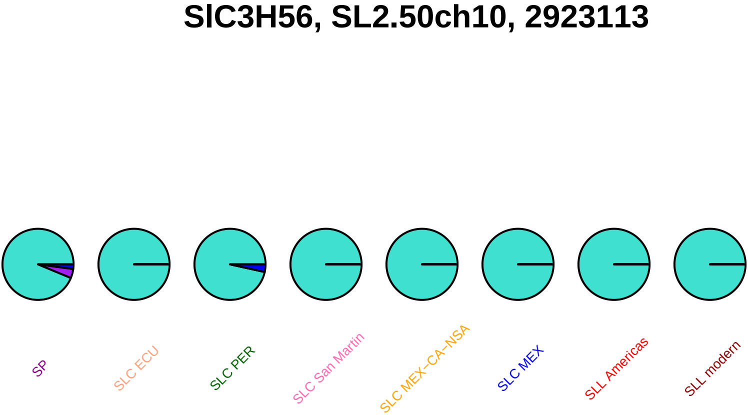
SlC3H56, SL2.50ch10, 2923113
SP
SLC ECU
SLC PER
SLC San Martin
SLC MEX−CA−NSA
SLC MEX
SLL Americas
SLL modern

### SlMAPKKK72, SL2.50ch10, 3092306

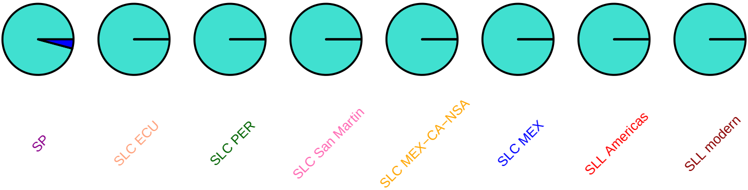

### SlbHLH154, SL2.50ch10, 3260442

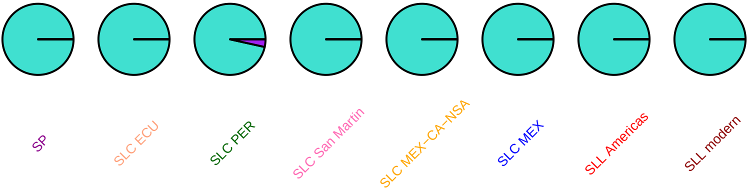

### SlMAPKKK73, SL2.50ch10, 3415982

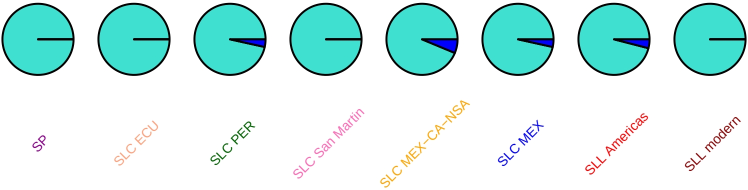

#### SlMAPKKK73, SL2.50ch10, 3418036

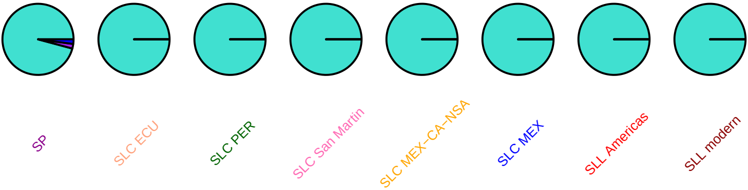

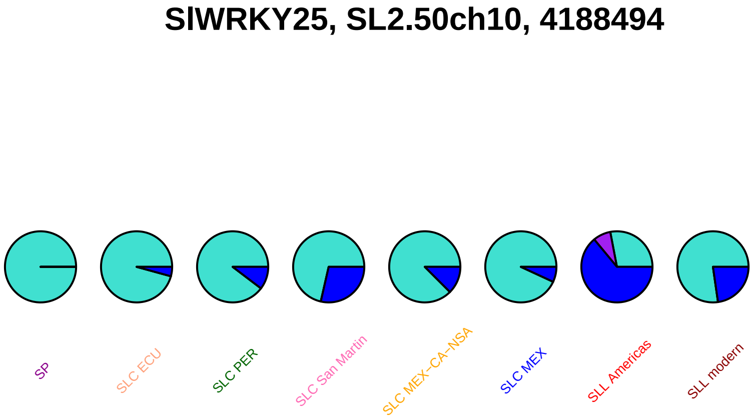
SlWRKY25, SL2.50ch10, 4188494
SP
SLC ECU
SLC PER
SLC San Martin
SLC MEX−CA−NSA
SLC MEX
SLL Americas
SLL modern

### SlbHLH157, SL2.50ch10, 46827509

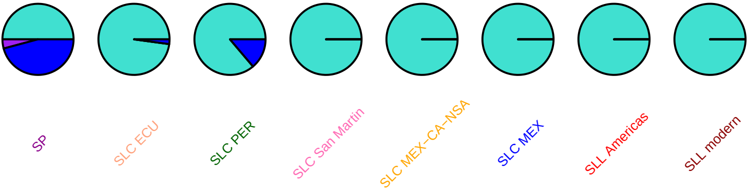

# AY506544, SL2.50ch10, 50976671

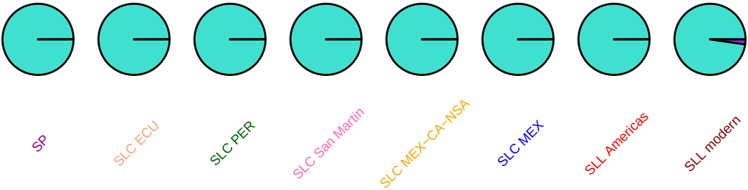

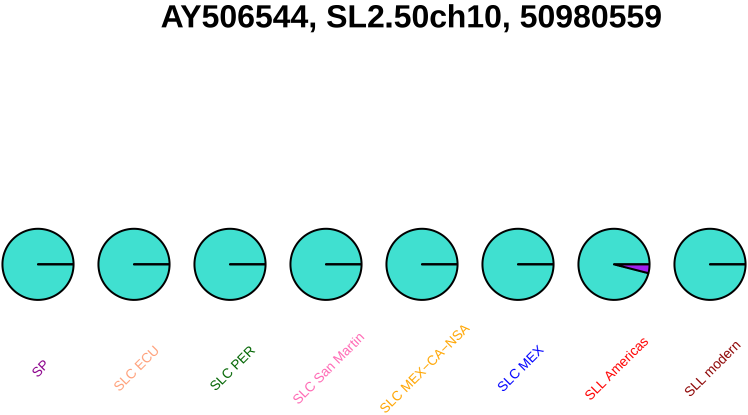
AY506544, SL2.50ch10, 50980559
SP
SLC ECU
SLC PER
SLC San Martin
SLC MEX−CA−NSA
SLC MEX
SLL Americas
SLL modern

### nii2, SL2.50ch10, 51135300

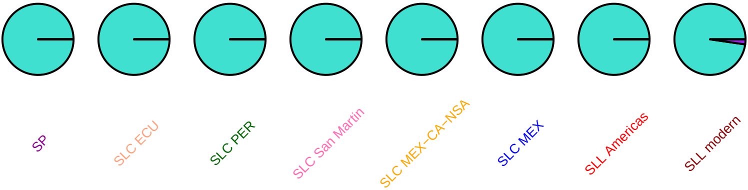

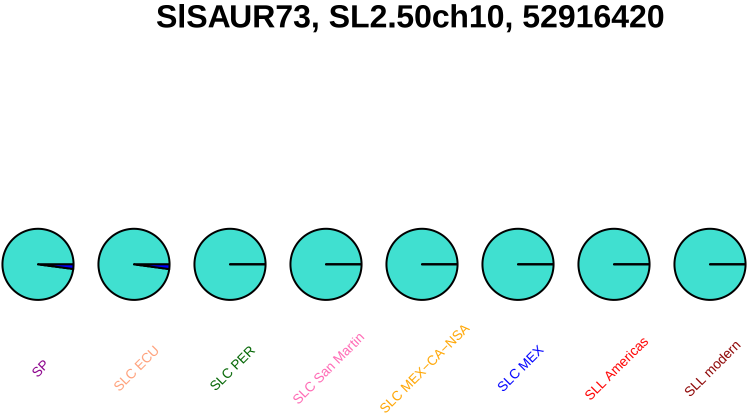
SlSAUR73, SL2.50ch10, 52916420
SP
SLC ECU
SLC PER
SLC San Martin
SLC MEX–CA–NSA
SLC MEX
SLL Americas
SLL modern

### SlSAUR74, SL2.50ch10, 52917832

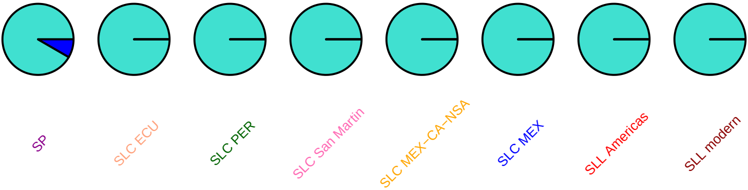

### SlSAUR74, SL2.50ch10, 52917966

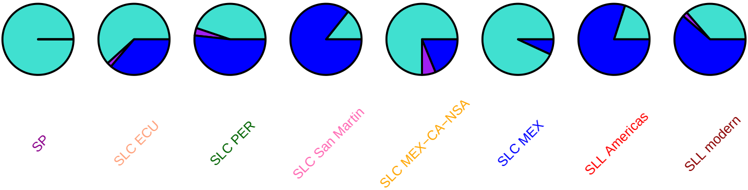

#### SlSAUR75, SL2.50ch10, 52920073

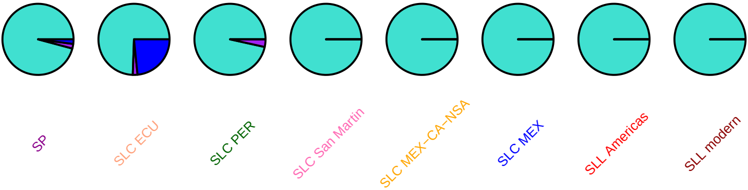

### SlSAUR76, SL2.50ch10, 52935941

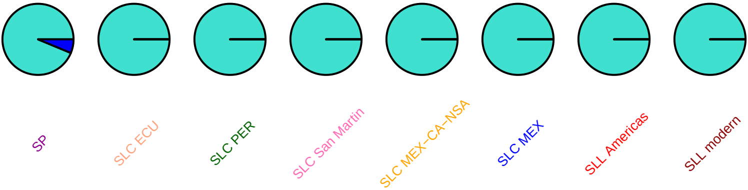

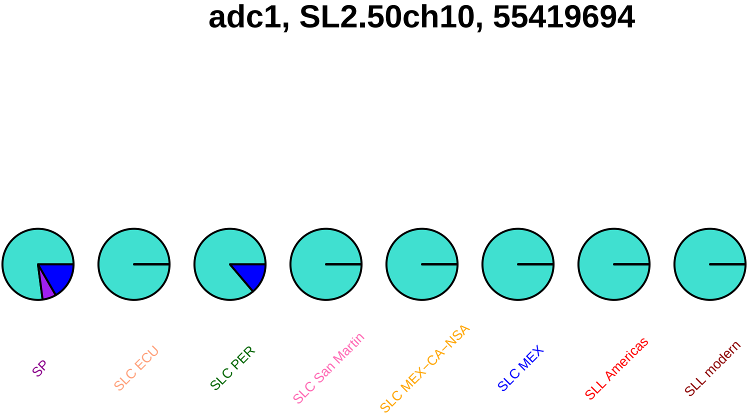
adc1, SL2.50ch10, 55419694
SP
SLC ECU
SLC PER
SLC San Martin
SLC MEX–CA–NSA
SLC MEX
SLL Americas
SLL modern

### SlSAUR81, SL2.50ch10, 55908364

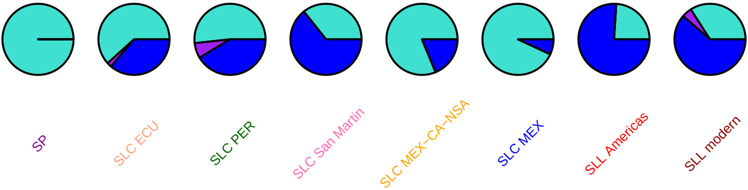

#### SlSAUR81, SL2.50ch10, 55908401

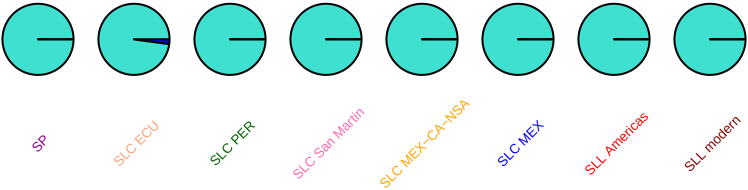

### SlXIP1.3, SL2.50ch10, 55955966

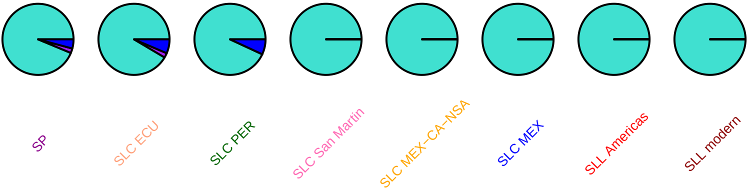

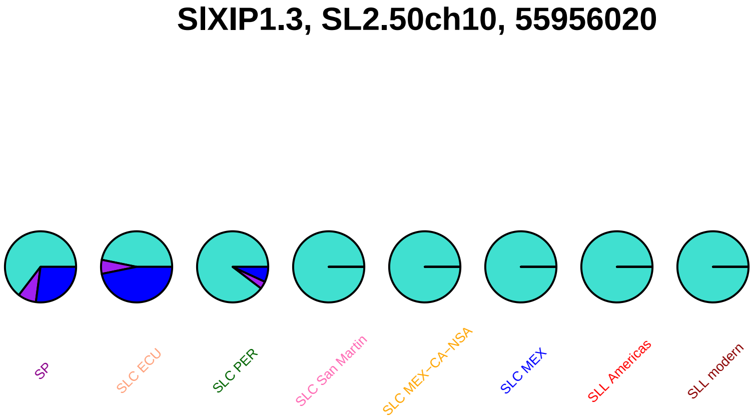
SlXIP1.3, SL2.50ch10, 55956020
SP
SLC ECU
SLC PER
SLC San Martin
SLC MEX−CA−NSA
SLC MEX
SLL Americas
SLL modern

### SlXIP1.3, SL2.50ch10, 55956038

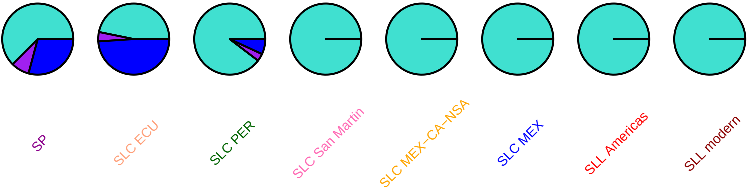

### SlXIP1.3, SL2.50ch10, 55956088

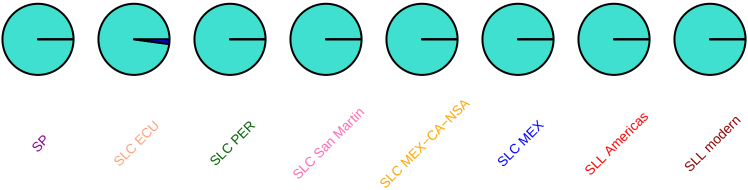

# Z15140, SL2.50ch10, 57417505

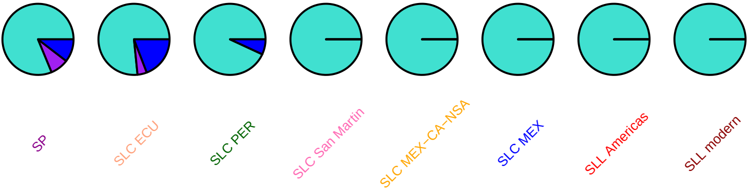

## Z15140, SL2.50ch10, 57417643

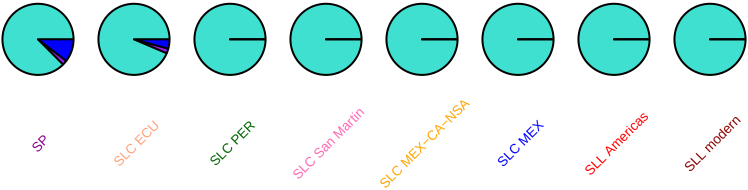

### lysrs, SL2.50ch10, 59568723

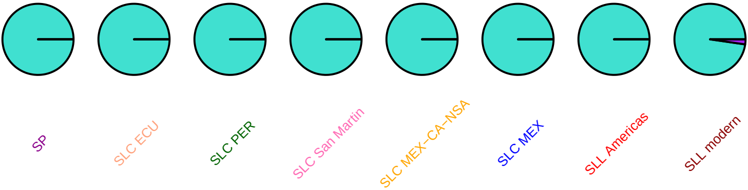

# tf3a, SL2.50ch10, 59994890

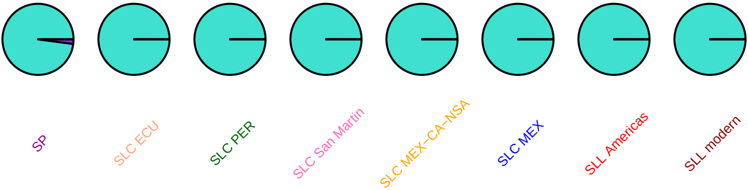

#### cycb1d1, SL2.50ch10, 60154582

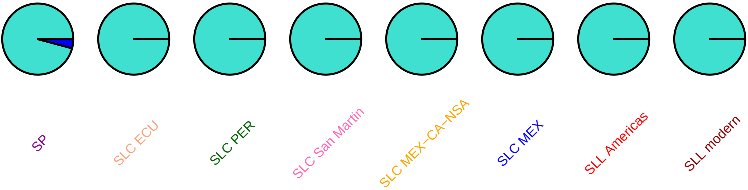

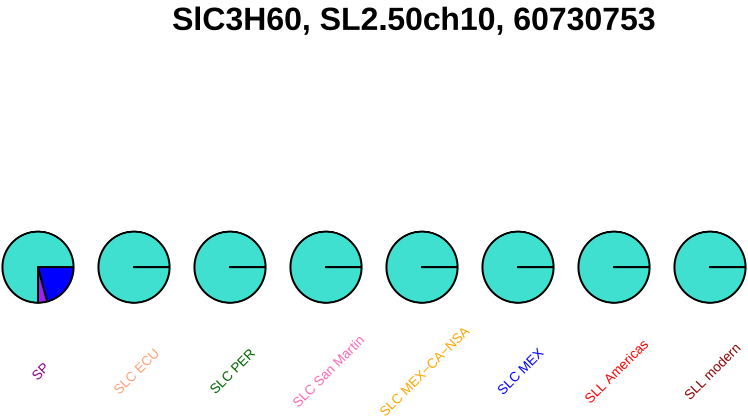

SlC3H60, SL2.50ch10, 60730753
SP
SLC ECU
SLC PER
SLC San Martin
SLC MEX–CA–NSA
SLC MEX
SLL Americas
SLL modern

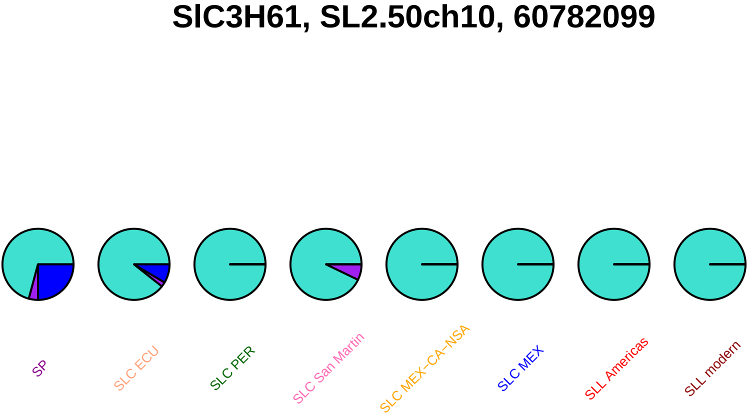
SlC3H61, SL2.50ch10, 60782099
SP
SLC ECU
SLC PER
SLC San Martin
SLC MEX−CA−NSA
SLC MEX
SLL Americas
SLL modern

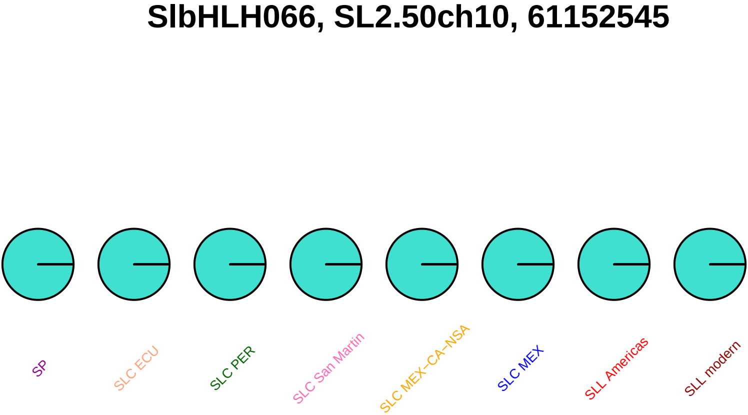
SlbHLH066, SL2.50ch10, 61152545
SP
SLC ECU
SLC PER
SLC San Martin
SLC MEX−CA−NSA
SLC MEX
SLL Americas
SLL modern

### SlbHLH067, SL2.50ch10, 61161564

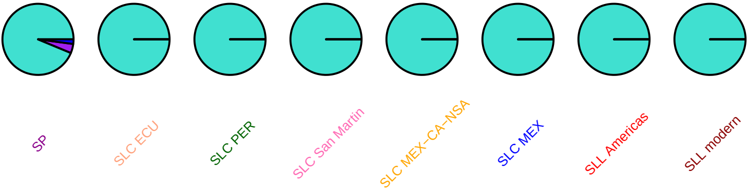

### SlbHLH068, SL2.50ch10, 61177515

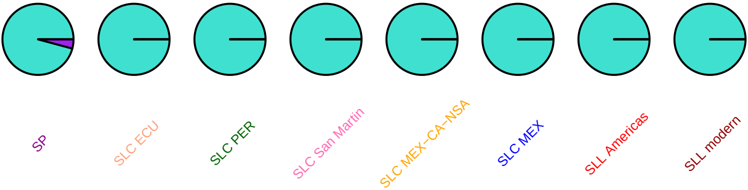

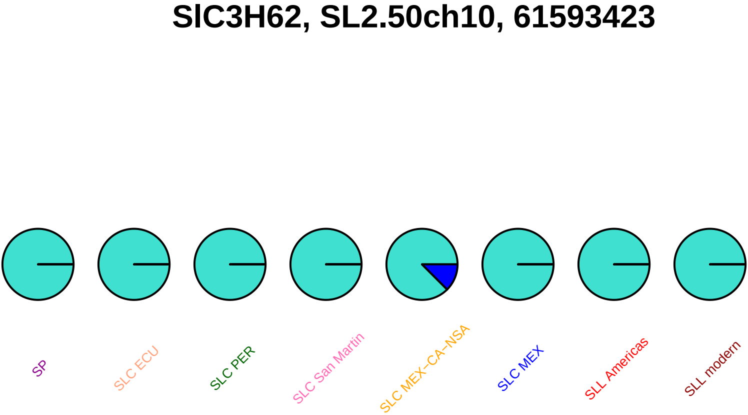

SlC3H62, SL2.50ch10, 61593423
SP
SLC ECU
SLC PER
SLC San Martin
SLC MEX−CA−NSA
SLC MEX
SLL Americas
SLL modern

#### SlC3H62, SL2.50ch10, 61593783

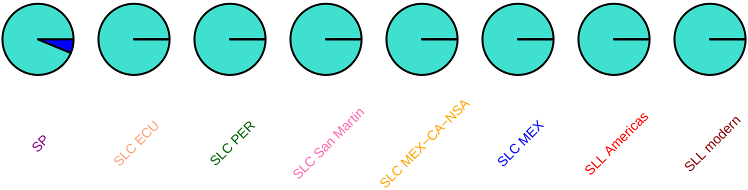

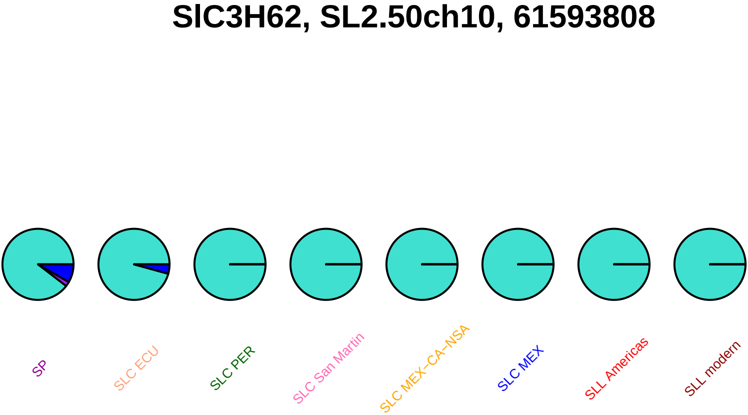
SlC3H62, SL2.50ch10, 61593808
SP
SLC ECU
SLC PER
SLC San Martin
SLC MEX−CA−NSA
SLC MEX
SLL Americas
SLL modern

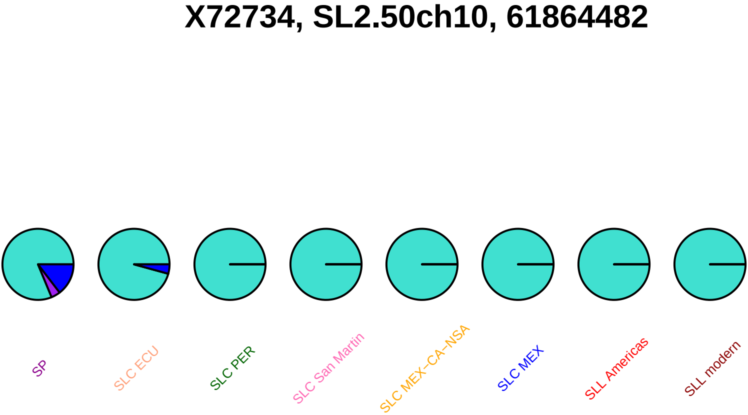
X72734, SL2.50ch10, 61864482
SP
SLC ECU
SLC PER
SLC San Martin
SLC MEX–CA–NSA
SLC MEX
SLL Americas
SLL modern

# X72730, SL2.50ch10, 62072177

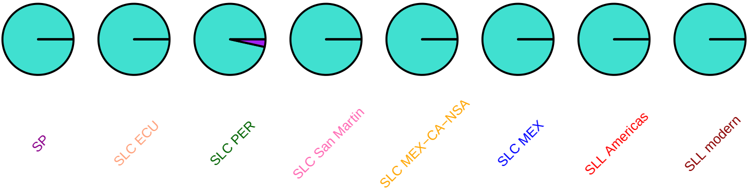

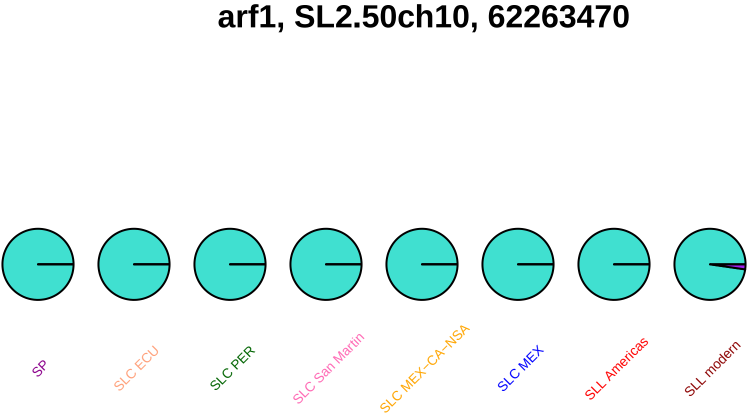
arf1, SL2.50ch10, 62263470
SP
SLC ECU
SLC PER
SLC San Martin
SLC MEX−CA−NSA
SLC MEX
SLL Americas
SLL modern

### crtiso, SL2.50ch10, 62683433

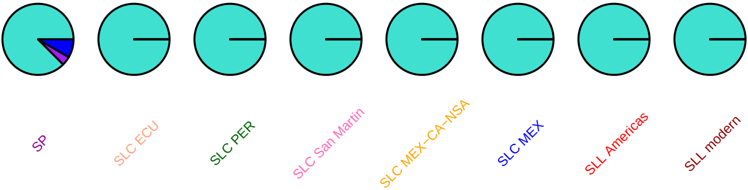

#### crtiso, SL2.50ch10, 62684196

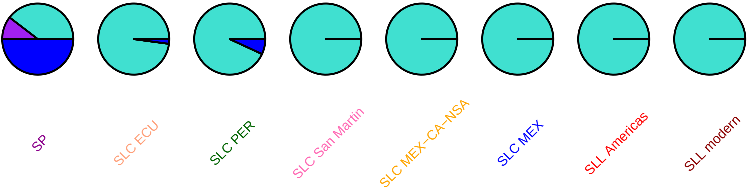

### SlOFP22, SL2.50ch10, 62950440

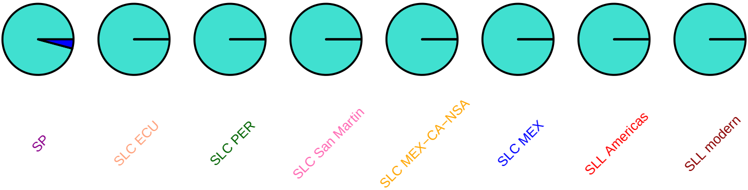

### SlOFP27, SL2.50ch10, 62974806

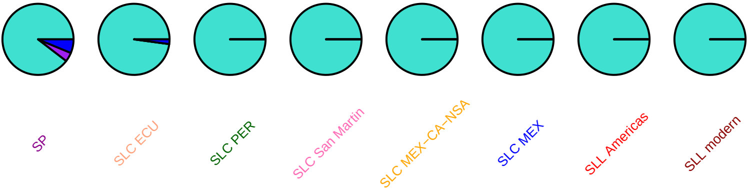

### SlOFP28, SL2.50ch10, 62980418

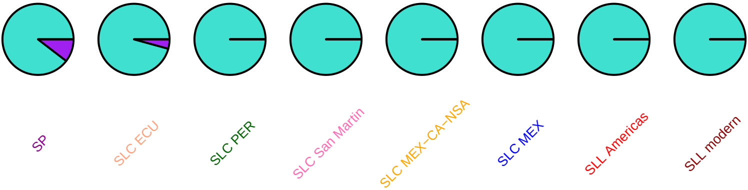

**lin8, SL2.50ch10, 63124370**

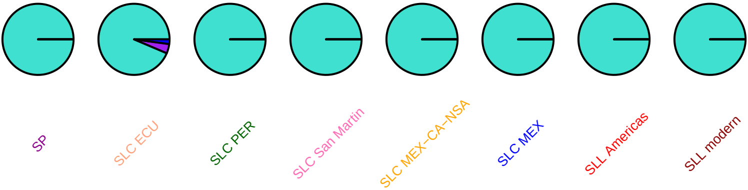

### SlMAPKKK77, SL2.50ch10, 63392348

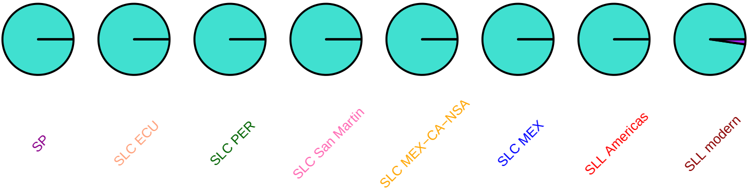

### SlMAPKKK77, SL2.50ch10, 63395038

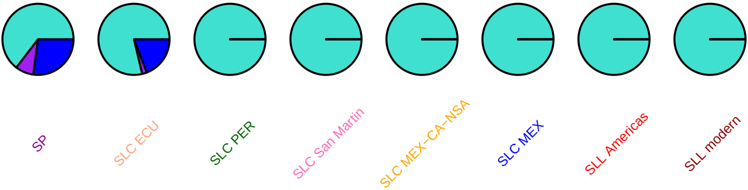

### SlMAPKKK77, SL2.50ch10, 63396602

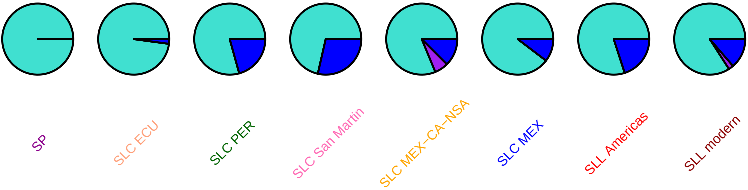

### SlSUN29, SL2.50ch10, 63901631

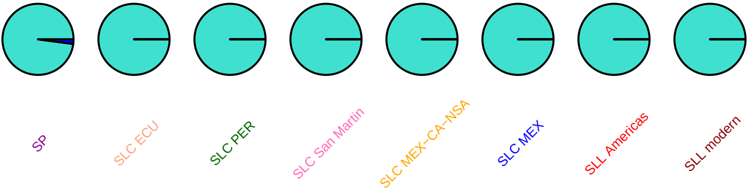

#### ert1b, SL2.50ch10, 64501531

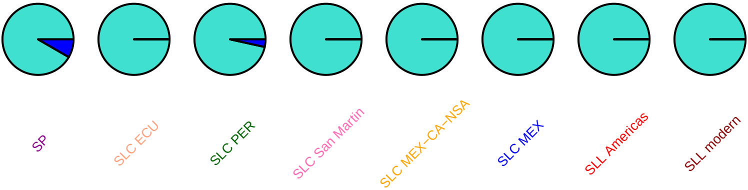

### ctr4, SL2.50ch10, 64684405

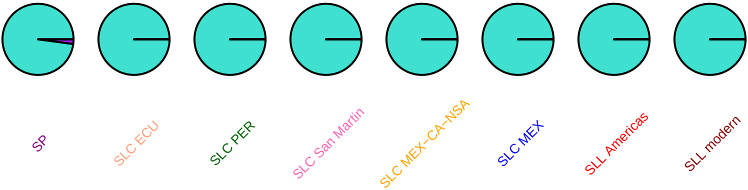

### ctr4, SL2.50ch10, 64693642

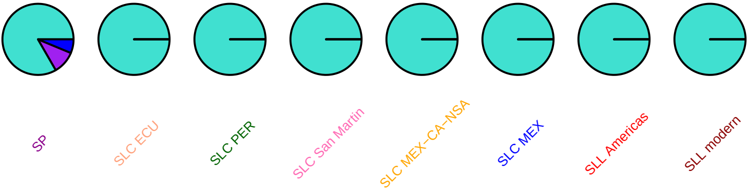

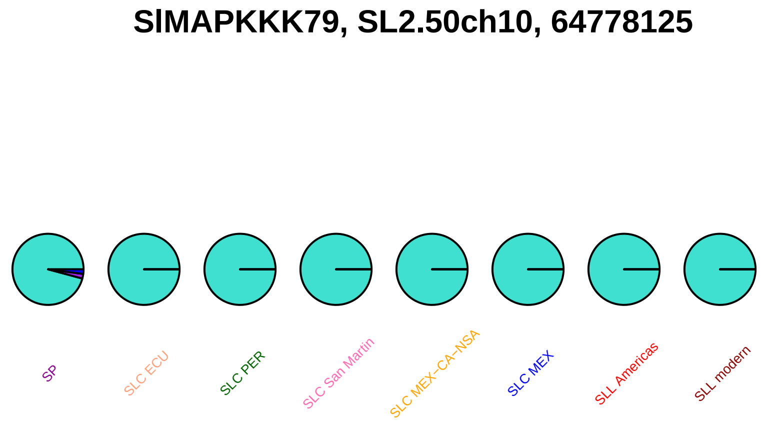
SlMAPKKK79, SL2.50ch10, 64778125
SP
SLC ECU
SLC PER
SLC San Martin
SLC MEX–CA–NSA
SLC MEX
SLL Americas
SLL modern

#### SlSUN30, SL2.50ch10, 65025639

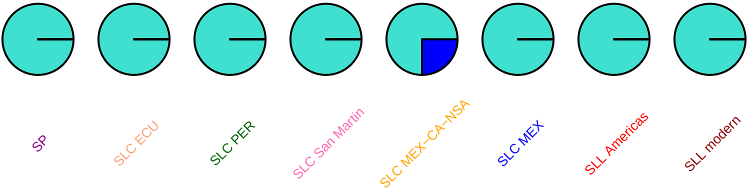

### opr, SL2.50ch10, 65121491

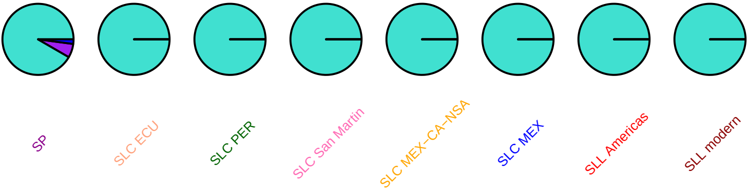

### opr, SL2.50ch10, 65121863

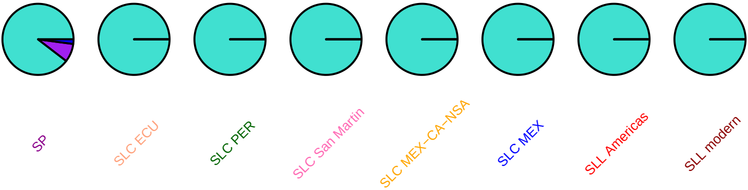

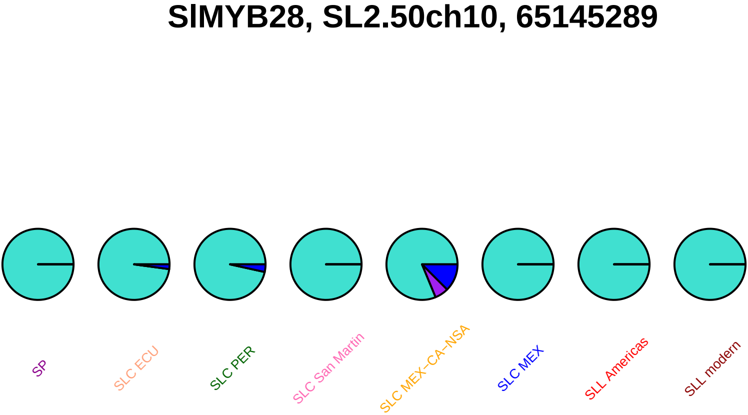
SlMYB28, SL2.50ch10, 65145289
SP
SLC ECU
SLC PER
SLC San Martin
SLC MEX−CA−NSA
SLC MEX
SLL Americas
SLL modern

#### SlMYB28, SL2.50ch10, 65145392

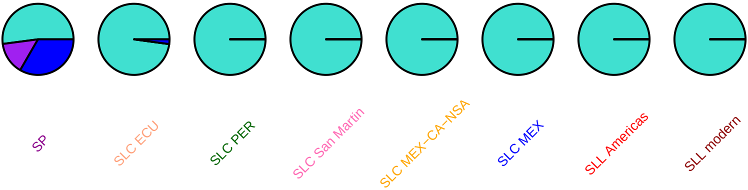

#### SlMYB28, SL2.50ch10, 65145550

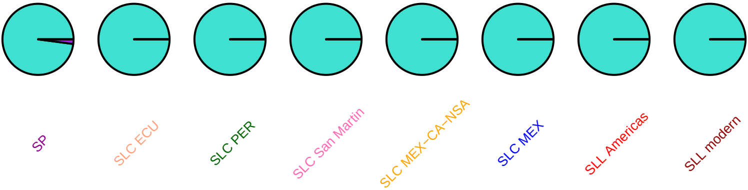

### SlMYB28, SL2.50ch10, 65145571

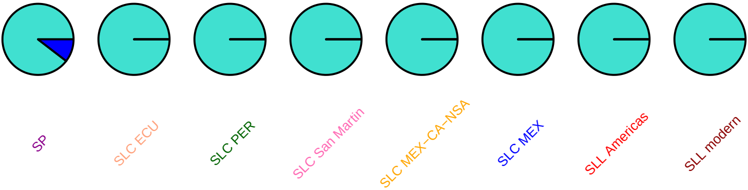

### SlMYB114, SL2.50ch10, 65168172

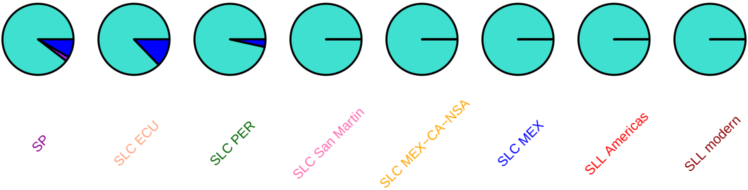

#### SlMYB114, SL2.50ch10, 65168183

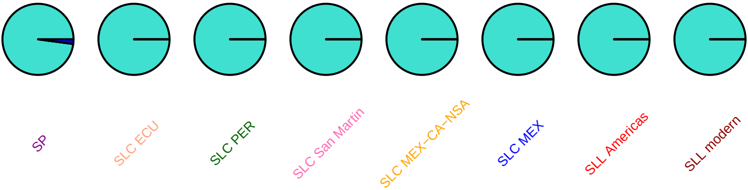

### cyca1, SL2.50ch11, 70078

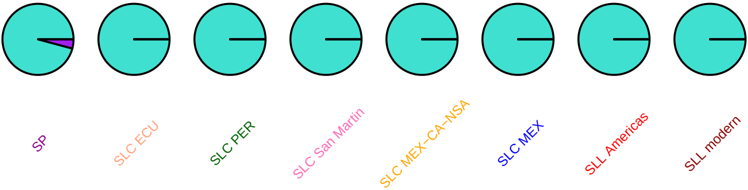

### skip5, SL2.50ch11, 448072

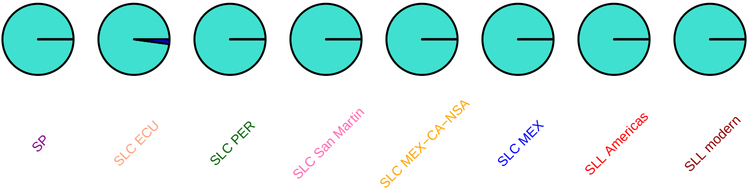

### MAPKKKa, SL2.50ch11, 812116

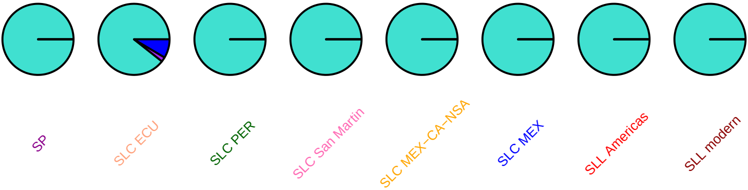

# DQ365932, SL2.50ch11, 1197487

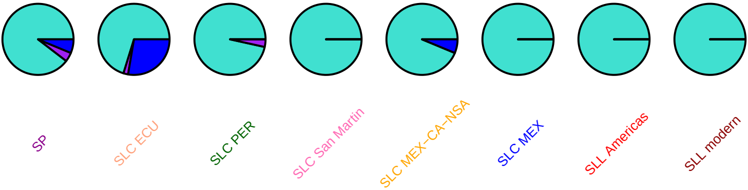

#### mdc, SL2.50ch11, 1514080

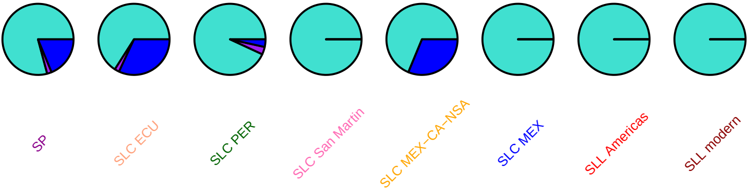

### ptpkis1, SL2.50ch11, 2052247

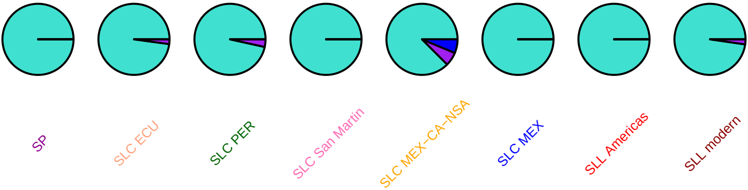

### SolycHsfl3, SL2.50ch11, 2604652

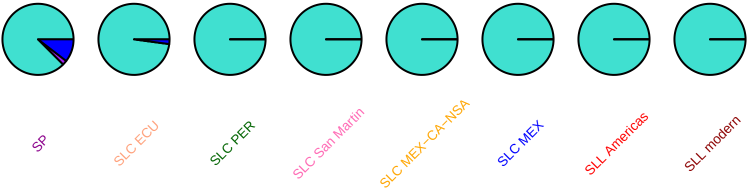

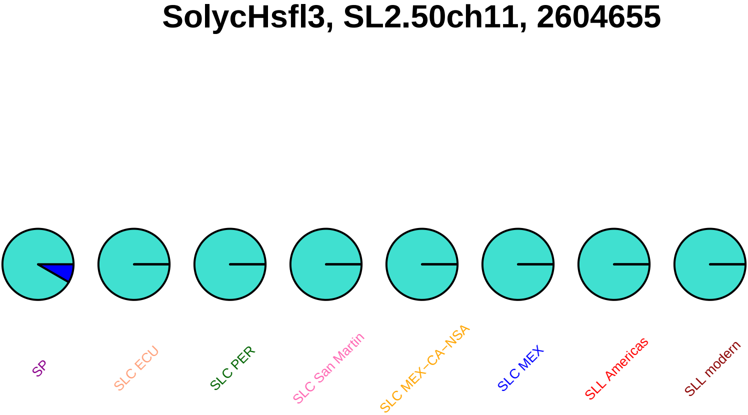

SolycHsfl3, SL2.50ch11, 2604655
SP
SLC ECU
SLC PER
SLC San Martin
SLC MEX–CA–NSA
SLC MEX
SLL Americas
SLL modern

#### SolycHsfl3, SL2.50ch11, 2604695

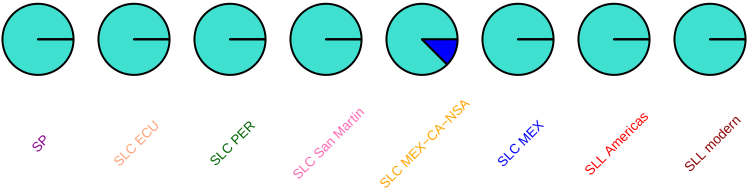

#### SolycHsfl3, SL2.50ch11, 2604856

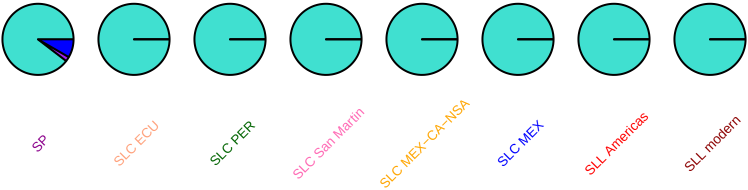

#### SlDCL2c, SL2.50ch11, 2694016

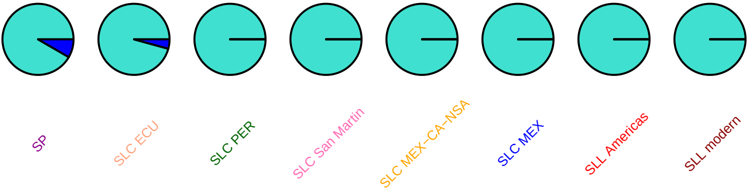

#### SlDCL2c, SL2.50ch11, 2694740

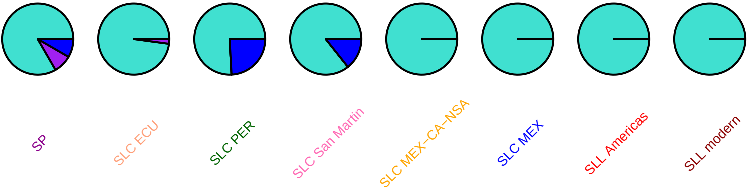

#### SlDCL2c, SL2.50ch11, 2696779

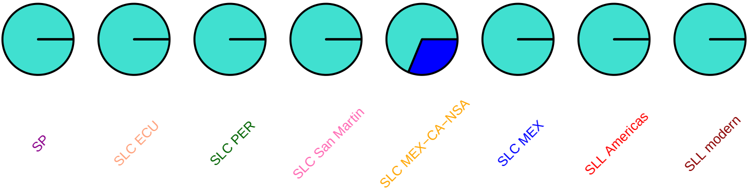

#### SlDCL2d, SL2.50ch11, 2701580

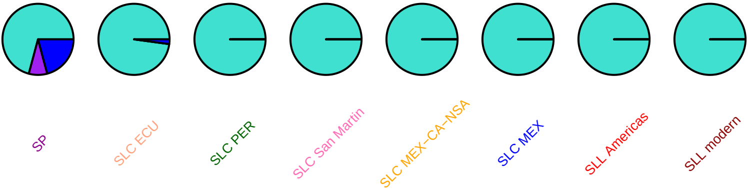

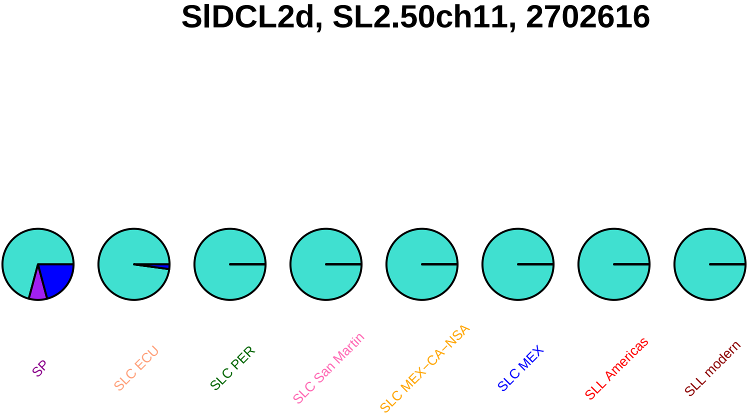
SlDCL2d, SL2.50ch11, 2702616
SP
SLC ECU
SLC PER
SLC San Martin
SLC MEX−CA−NSA
SLC MEX
SLL Americas
SLL modern

### SlDCL2d, SL2.50ch11, 2703552

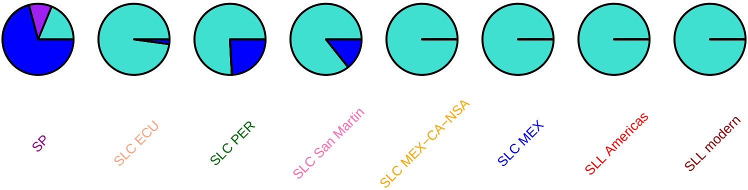

#### SlDCL2d, SL2.50ch11, 2705212

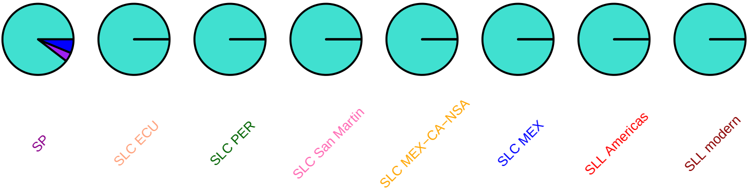

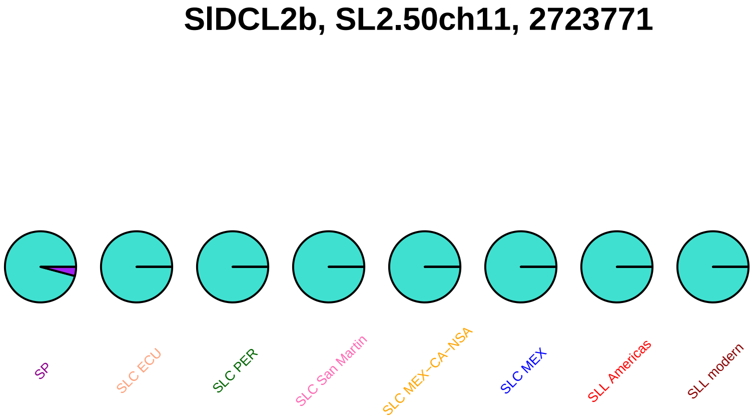

### SlC3H65, SL2.50ch11, 3074739

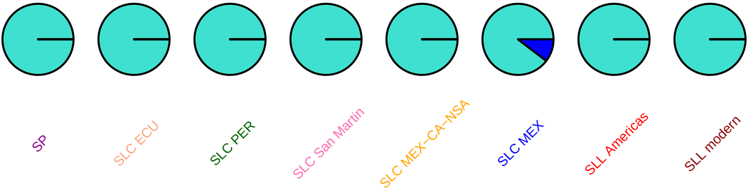

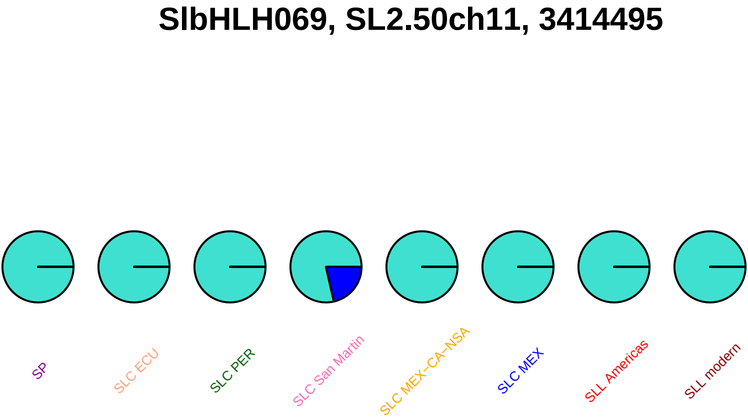
SlbHLH069, SL2.50ch11, 3414495
SP
SLC ECU
SLC PER
SLC San Martin
SLC MEX–CA–NSA
SLC MEX
SLL Americas
SLL modern

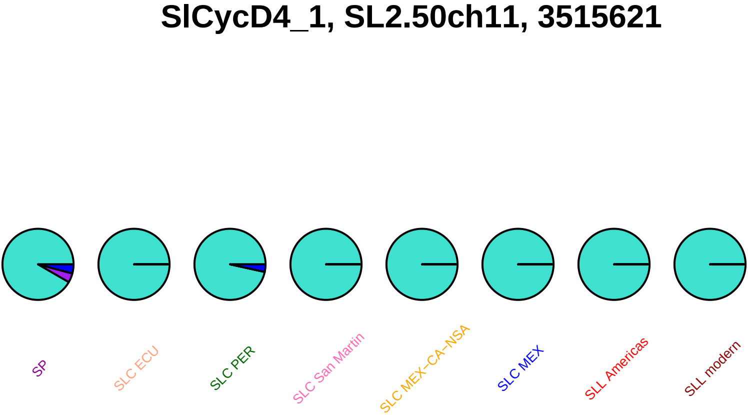
SlCycD4_1, SL2.50ch11, 3515621
SP
SLC ECU
SLC PER
SLC San Martin
SLC MEX−CA−NSA
SLC MEX
SLL Americas
SLL modern

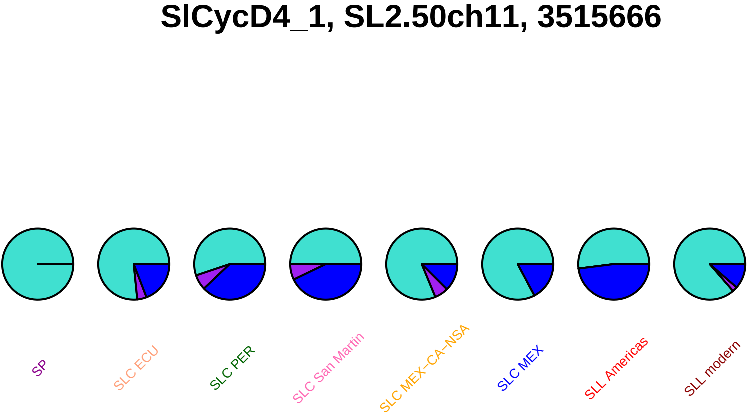
SlCycD4_1, SL2.50ch11, 3515666
SP
SLC ECU
SLC PER
SLC San Martin
SLC MEX−CA−NSA
SLC MEX
SLL Americas
SLL modern

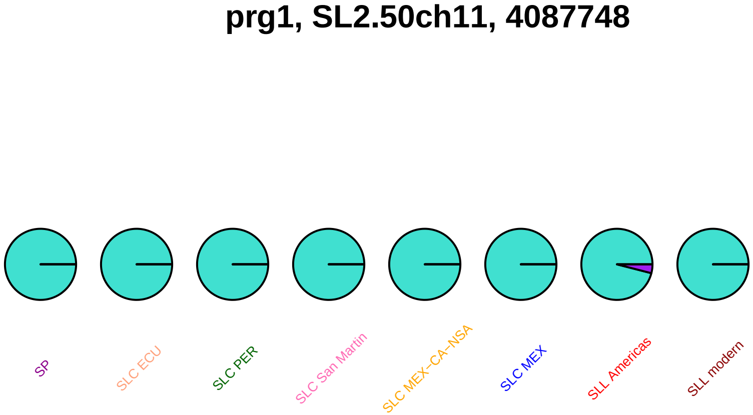
prg1, SL2.50ch11, 4087748
SP
SLC ECU
SLC PER
SLC San Martin
SLC MEX–CA–NSA
SLC MEX
SLL Americas
SLL modern

#### SlSAUR95, SL2.50ch11, 4709062

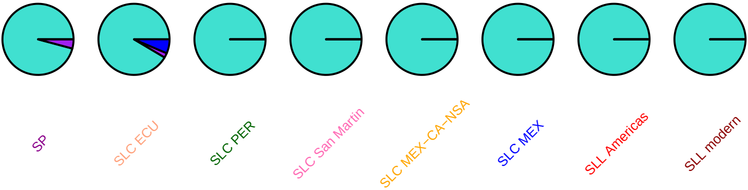

#### SlMAPKKK81, SL2.50ch11, 4994740

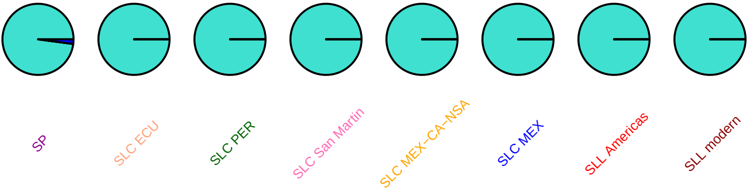

### mntH, SL2.50ch11, 8642460

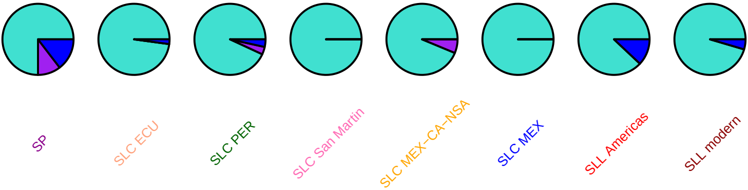

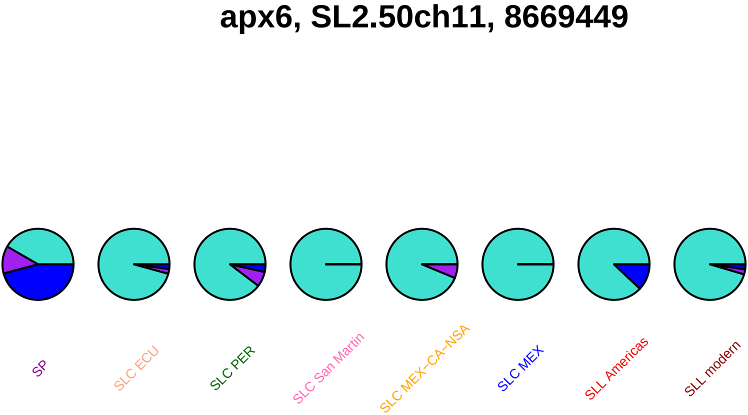
apx6, SL2.50ch11, 8669449
SP
SLC ECU
SLC PER
SLC San Martin
SLC MEX−CA−NSA
SLC MEX
SLL Americas
SLL modern

### hsp70, SL2.50ch11, 10016080

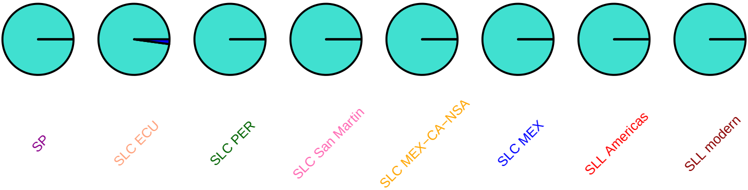

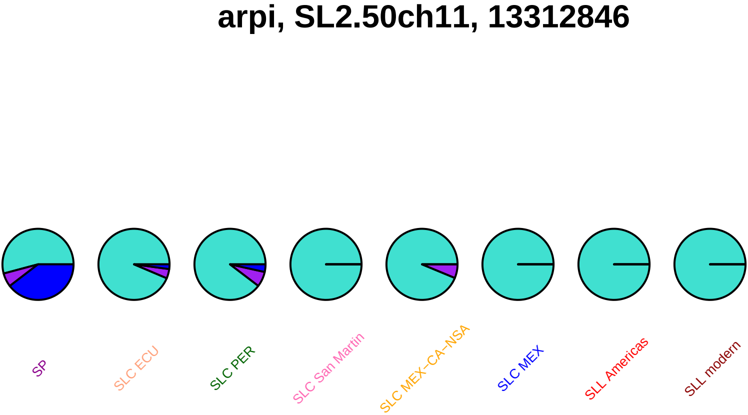
arpi, SL2.50ch11, 13312846
SP
SLC ECU
SLC PER
SLC San Martin
SLC MEX−CA−NSA
SLC MEX
SLL Americas
SLL modern

#### smet, SL2.50ch11, 22206692

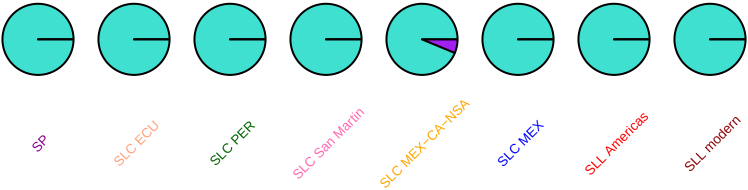

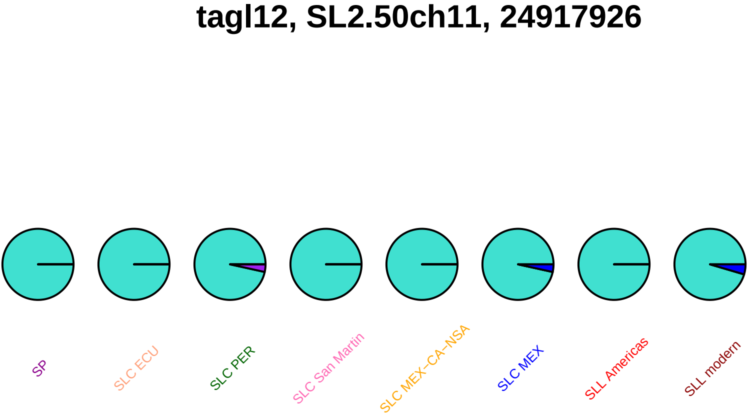
tagl12, SL2.50ch11, 24917926
SP
SLC ECU
SLC PER
SLC San Martin
SLC MEX–CA–NSA
SLC MEX
SLL Americas
SLL modern

### SlMAPKKK82, SL2.50ch11, 26372626

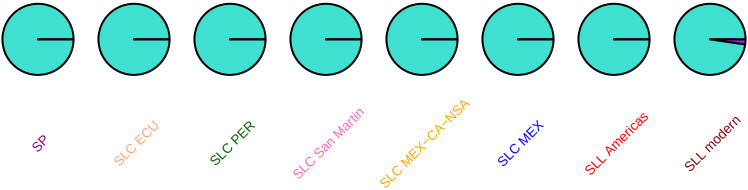

#### cel7, SL2.50ch11, 37864782

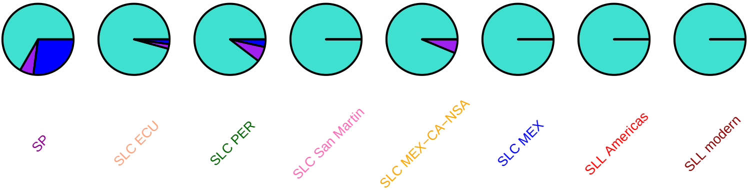

### SlOFP30, SL2.50ch11, 53412751

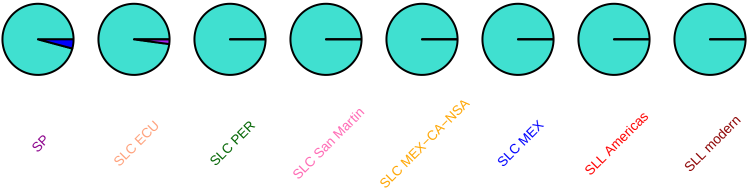

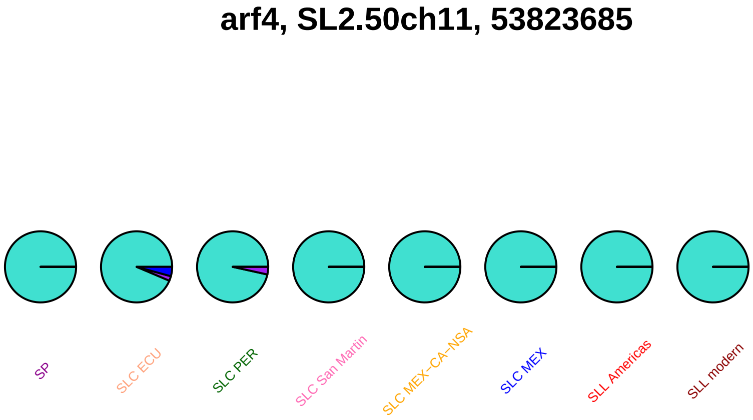
arf4, SL2.50ch11, 53823685
SP
SLC ECU
SLC PER
SLC San Martin
SLC MEX–CA–NSA
SLC MEX
SLL Americas
SLL modern

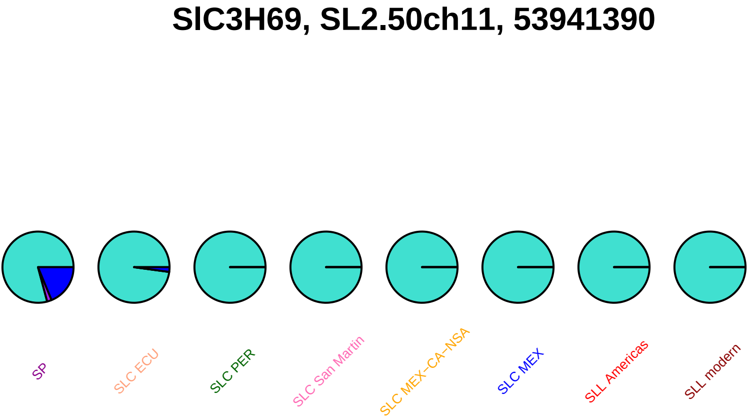
SlC3H69, SL2.50ch11, 53941390
SP
SLC ECU
SLC PER
SLC San Martin
SLC MEX−CA−NSA
SLC MEX
SLL Americas
SLL modern

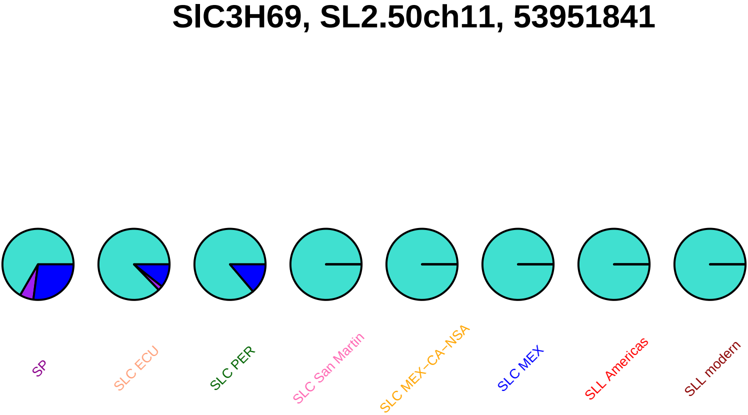
SlC3H69, SL2.50ch11, 53951841
SP
SLC ECU
SLC PER
SLC San Martin
SLC MEX–CA–NSA
SLC MEX
SLL Americas
SLL modern

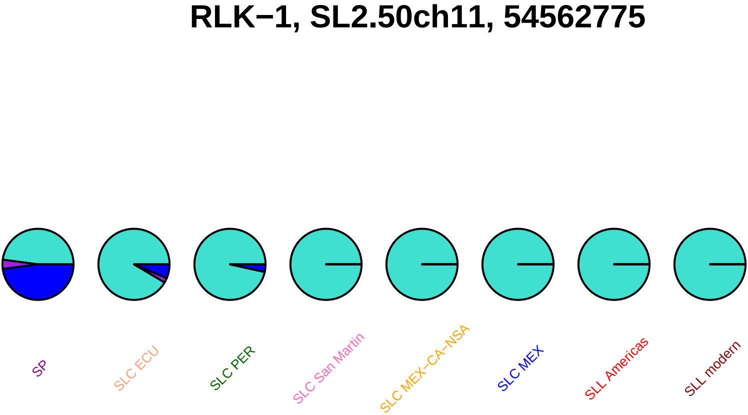
RLK−1, SL2.50ch11, 54562775
SP
SLC ECU
SLC PER
SLC San Martin
SLC MEX−CA−NSA
SLC MEX
SLL Americas
SLL modern

## I2C5, SL2.50ch11, 54578286

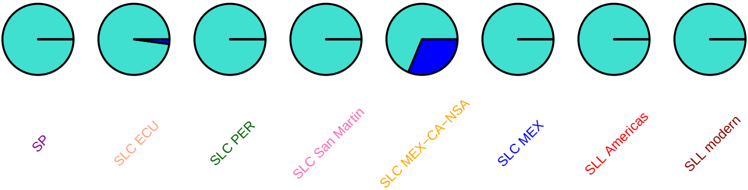

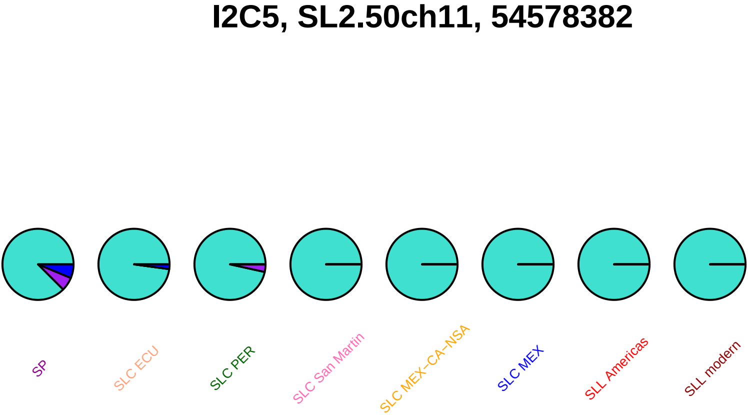
I2C5, SL2.50ch11, 54578382
SP
SLC ECU
SLC PER
SLC San Martin
SLC MEX−CA−NSA
SLC MEX
SLL Americas
SLL modern

# I2C5, SL2.50ch11, 54578516

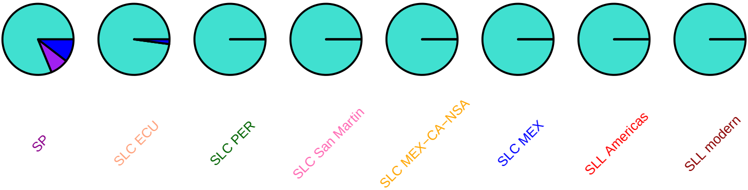

# I2C5, SL2.50ch11, 54580466

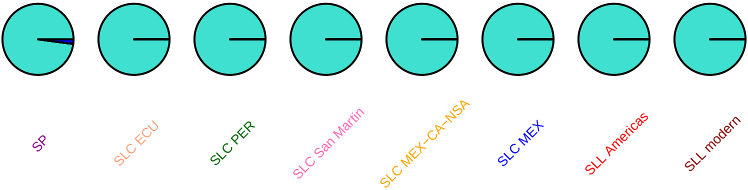

# I2C7, SL2.50ch11, 54882183

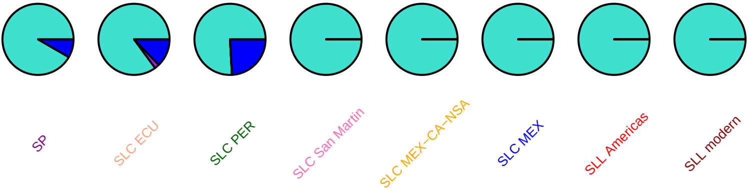

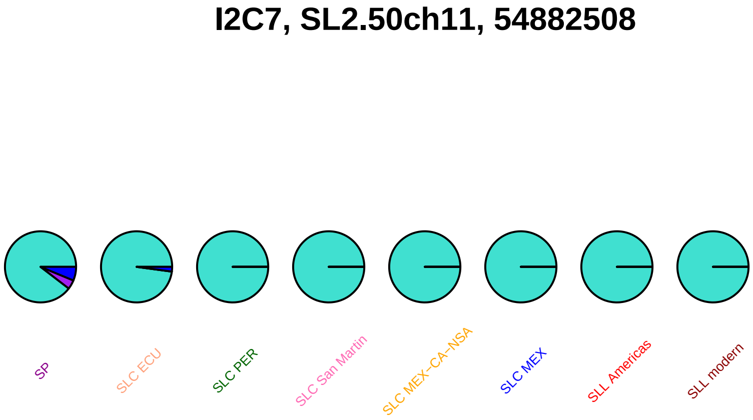
I2C7, SL2.50ch11, 54882508
SP
SLC ECU
SLC PER
SLC San Martin
SLC MEX−CA−NSA
SLC MEX
SLL Americas
SLL modern

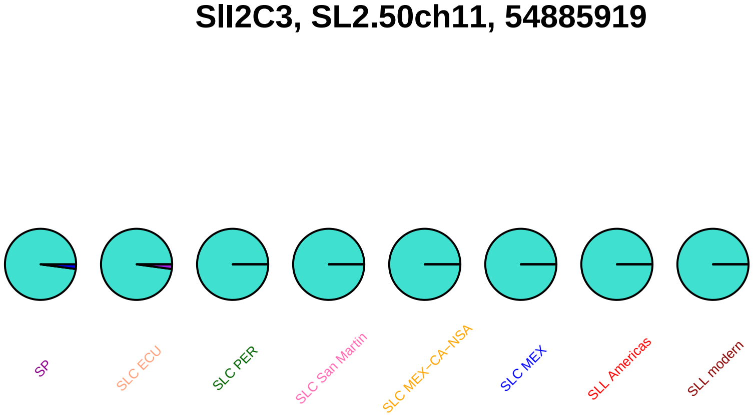
Sll2C3, SL2.50ch11, 54885919
SP
SLC ECU
SLC PER
SLC San Martin
SLC MEX−CA−NSA
SLC MEX
SLL Americas
SLL modern

### Sll2C3, SL2.50ch11, 54886008

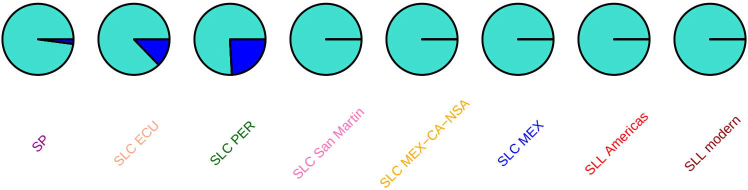

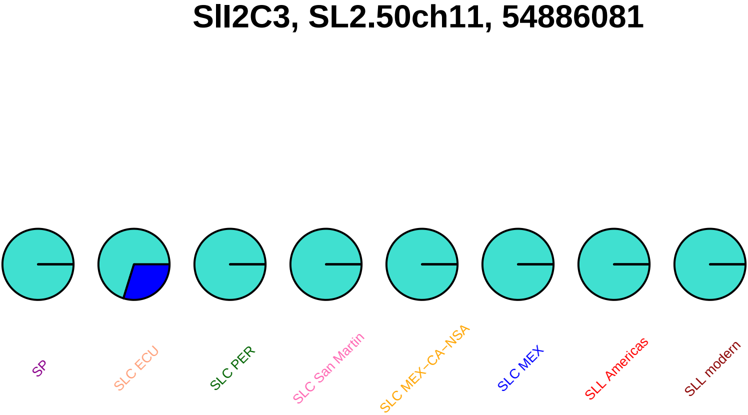
SlI2C3, SL2.50ch11, 54886081
SP
SLC ECU
SLC PER
SLC San Martin
SLC MEX–CA–NSA
SLC MEX
SLL Americas
SLL modern

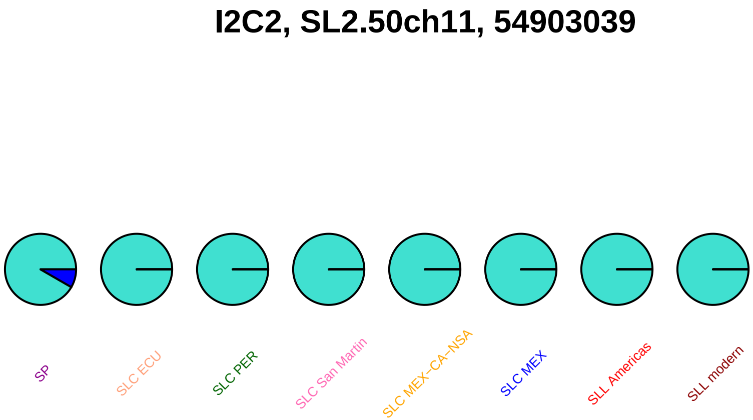

I2C2, SL2.50ch11, 54903039
SP
SLC ECU
SLC PER
SLC San Martin
SLC MEX−CA−NSA
SLC MEX
SLL Americas
SLL modern

## I2C2, SL2.50ch11, 54903142

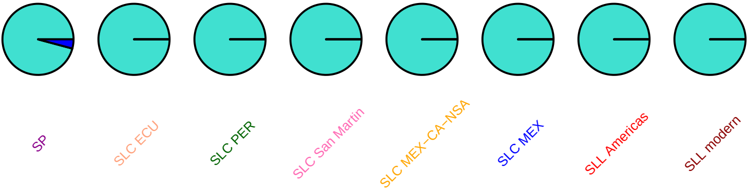

I2C2, SL2.50ch11, 54903300

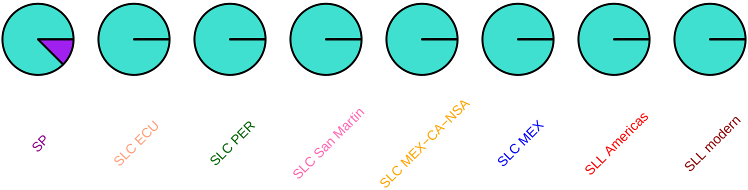

# I2C2, SL2.50ch11, 54903520

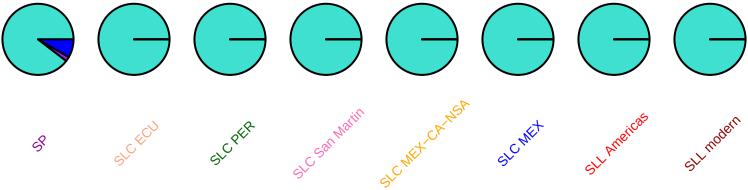

## I2C2, SL2.50ch11, 54905116

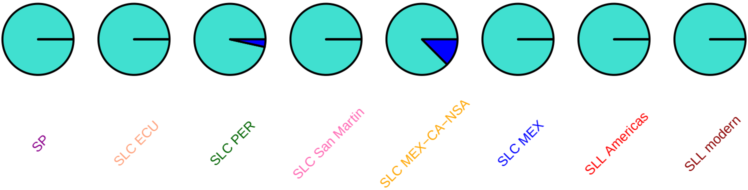

# ao5, SL2.50ch11, 55006969

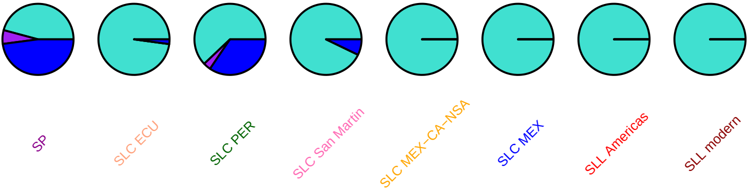

## ao5, SL2.50ch11, 55008375

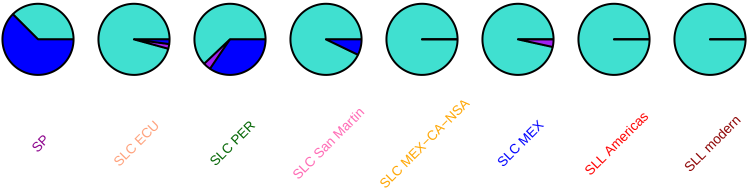

# ao5, SL2.50ch11, 55008416

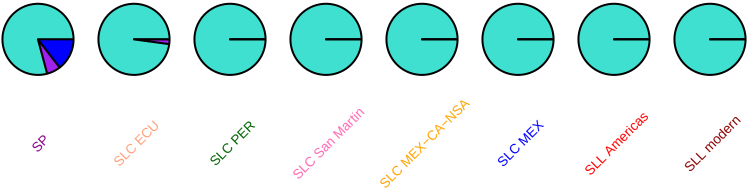

# ao5, SL2.50ch11, 55009253

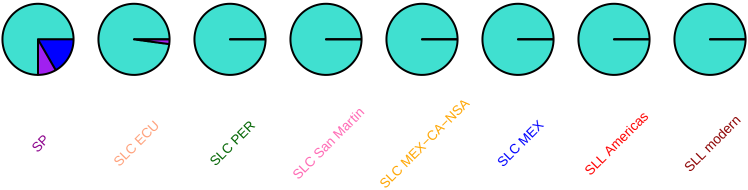

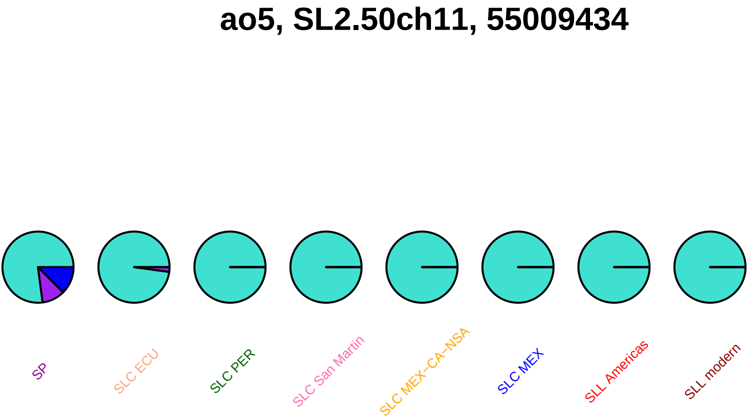
ao5, SL2.50ch11, 55009434
SP
SLC ECU
SLC PER
SLC San Martin
SLC MEX–CA–NSA
SLC MEX
SLL Americas
SLL modern

# ao5, SL2.50ch11, 55011192

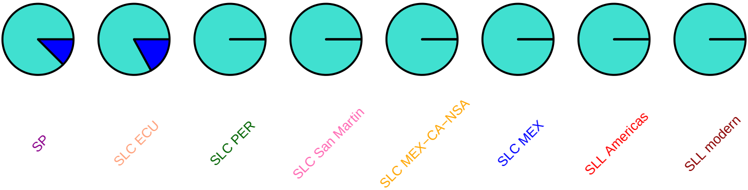

**ao5, SL2.50ch11, 55011400**

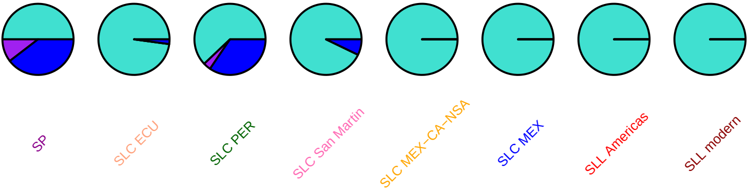

# ao5, SL2.50ch11, 55011927

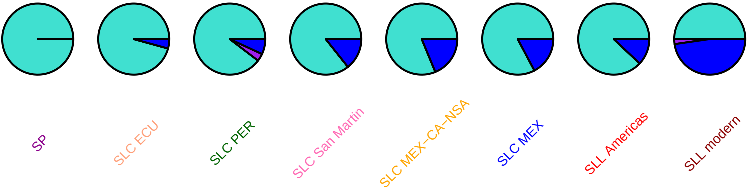

## ao5, SL2.50ch11, 55013010

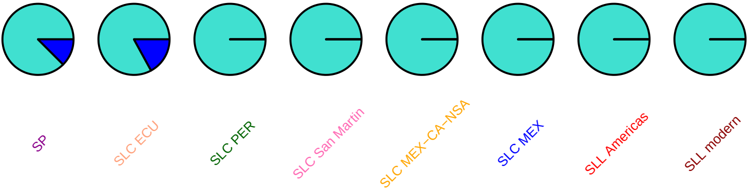

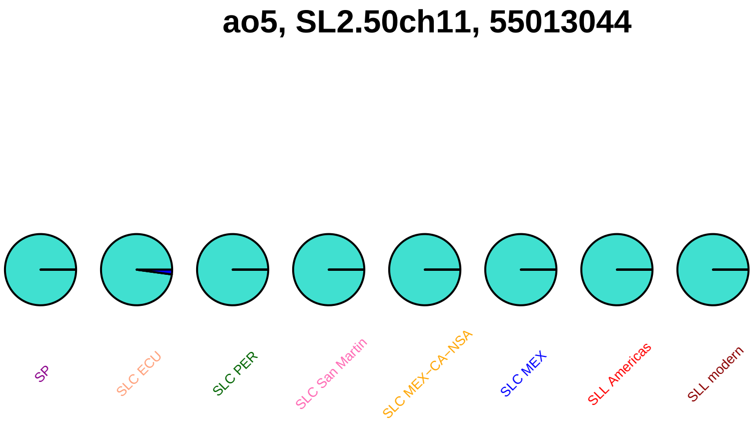

ao5, SL2.50ch11, 55013044
SP
SLC ECU
SLC PER
SLC San Martin
SLC MEX−CA−NSA
SLC MEX
SLL Americas
SLL modern

# ao3, SL2.50ch11, 55016945

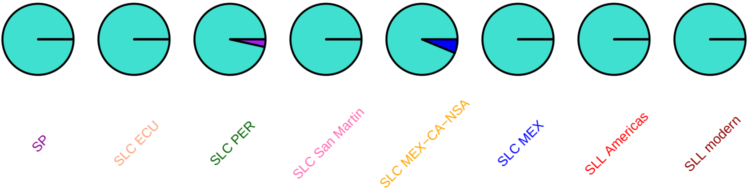

# ao3, SL2.50ch11, 55016948

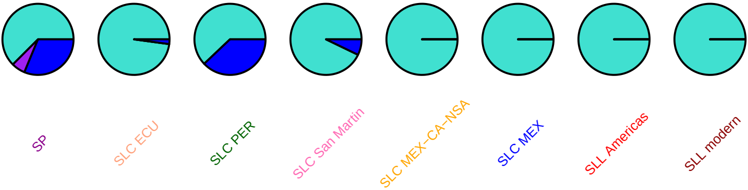

# ao3, SL2.50ch11, 55018275

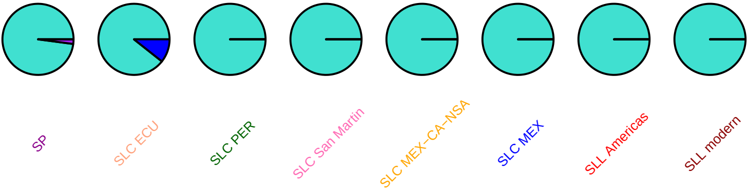

# ao3, SL2.50ch11, 55018518

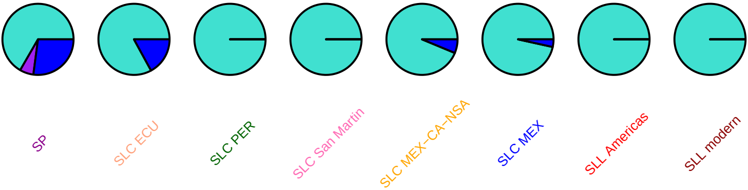

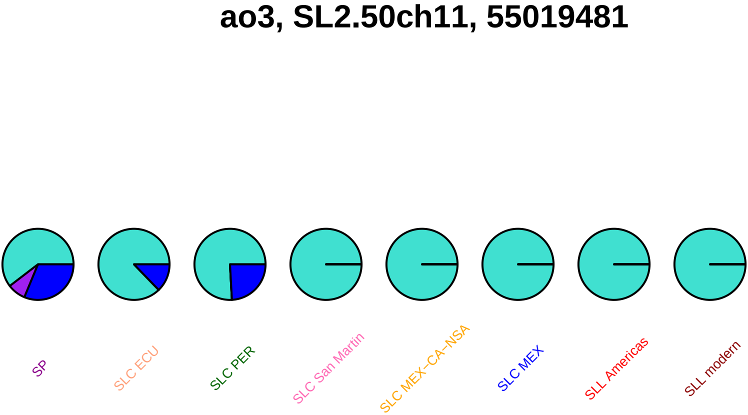

ao3, SL2.50ch11, 55019481
SP
SLC ECU
SLC PER
SLC San Martin
SLC MEX−CA−NSA
SLC MEX
SLL Americas
SLL modern

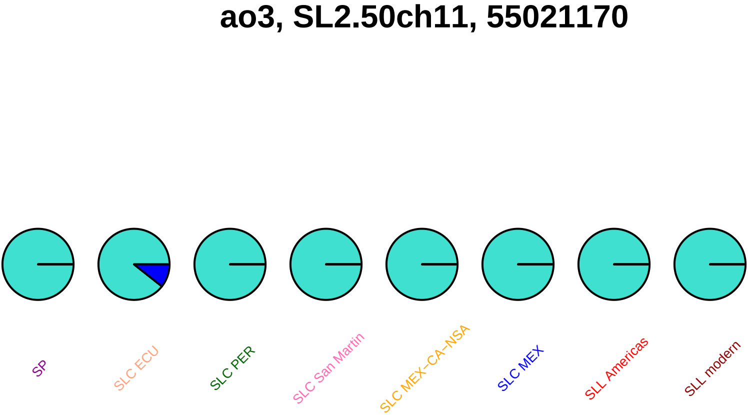
ao3, SL2.50ch11, 55021170
SP
SLC ECU
SLC PER
SLC San Martin
SLC MEX−CA−NSA
SLC MEX
SLL Americas
SLL modern

## ao1, SL2.50ch11, 55048732

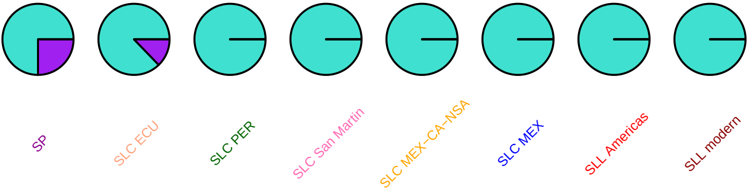

# ao1, SL2.50ch11, 55048856

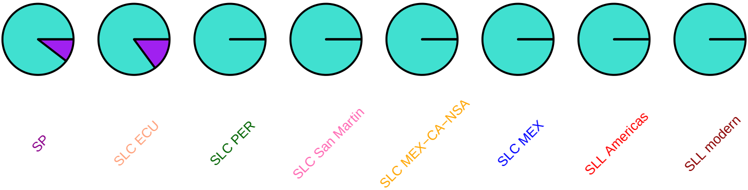

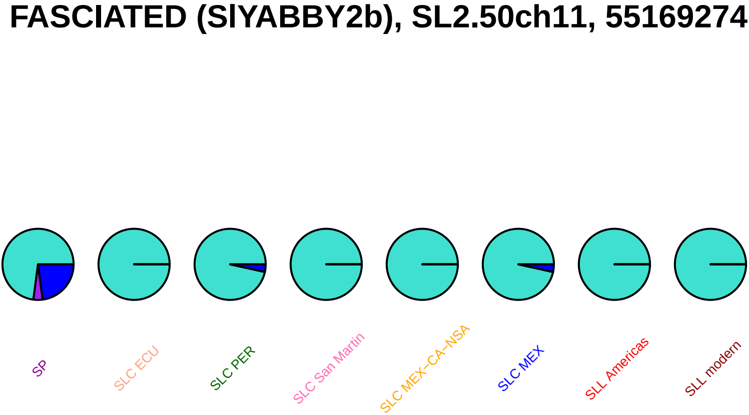
FASCIATED (SlYABBY2b), SL2.50ch11, 55169274
SP
SLC ECU
SLC PER
SLC San Martin
SLC MEX–CA–NSA
SLC MEX
SLL Americas
SLL modern

# AF049900, SL2.50ch11, 55510248

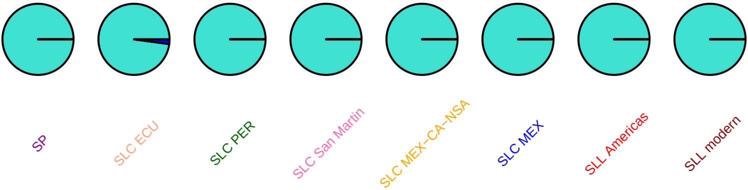

### SlMYB68, SL2.50ch12, 339694

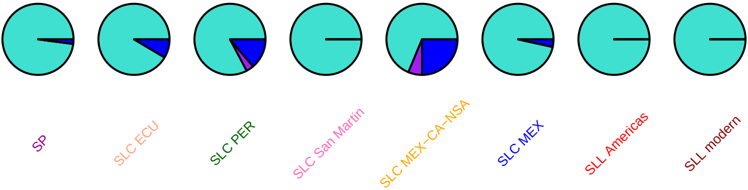

#### HAK5, SL2.50ch12, 361820

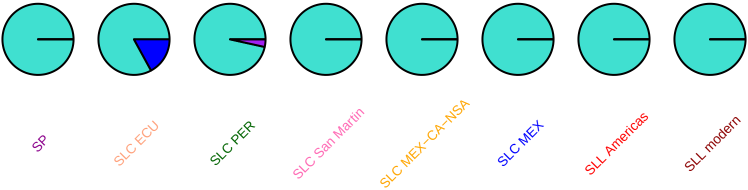

### cab5a, SL2.50ch12, 693285

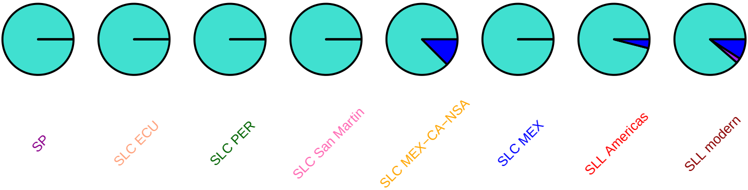

#### SlDEAD35, SL2.50ch12, 843054

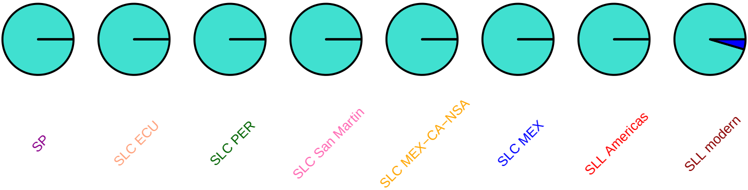

### SlC3H72, SL2.50ch12, 1092658

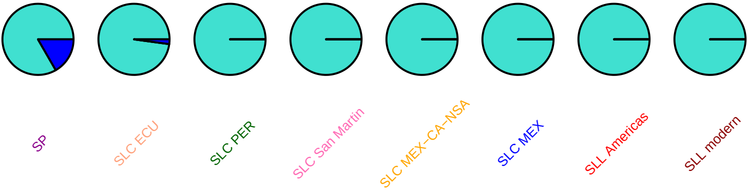

### CYP734A8, SL2.50ch12, 1300693

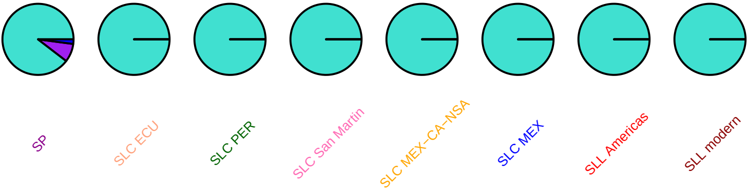

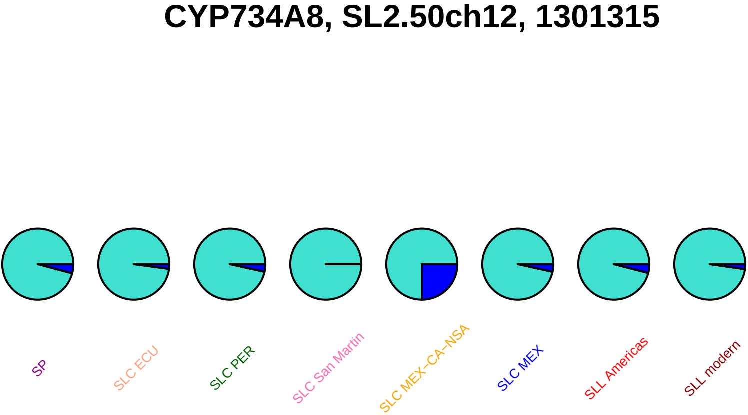

CYP734A8, SL2.50ch12, 1301315
SP
SLC ECU
SLC PER
SLC San Martin
SLC MEX−CA−NSA
SLC MEX
SLL Americas
SLL modern

### TBG4, SL2.50ch12, 2186084

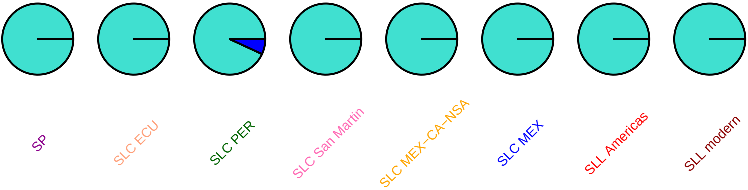

### TBG4, SL2.50ch12, 2186095

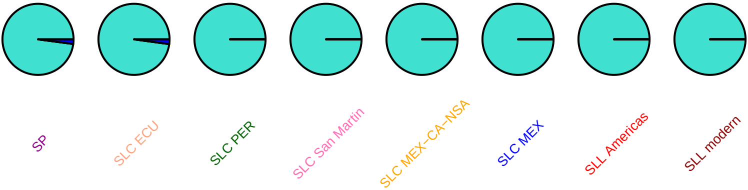

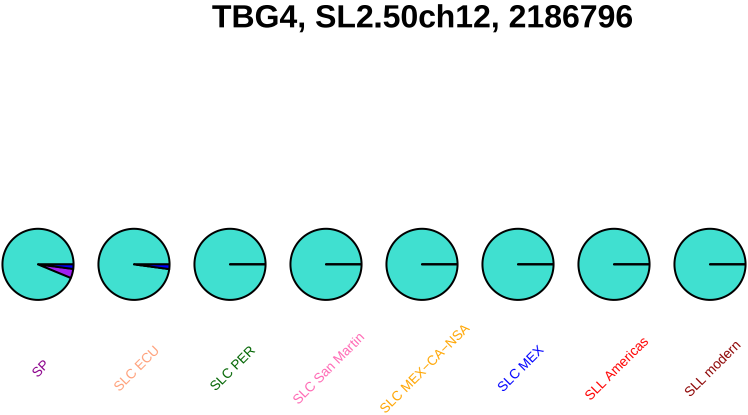
TBG4, SL2.50ch12, 2186796
SP
SLC ECU
SLC PER
SLC San Martin
SLC MEX−CA−NSA
SLC MEX
SLL Americas
SLL modern

### TBG4, SL2.50ch12, 2187169

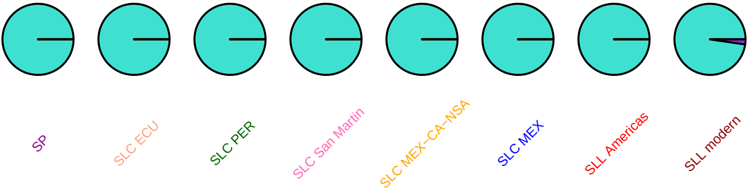

#### TBG4, SL2.50ch12, 2187233

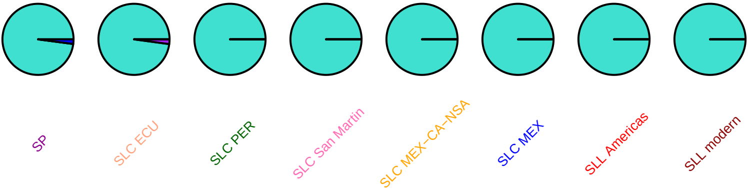

#### SlCKX6, SL2.50ch12, 2210550

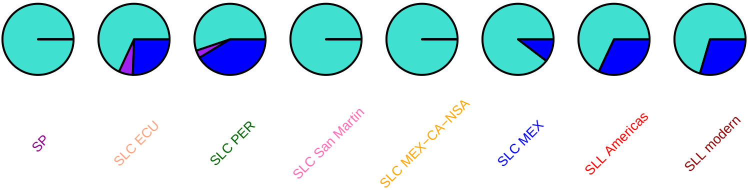

#### SlCKX6, SL2.50ch12, 2210668

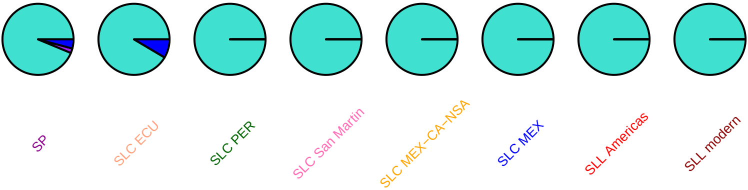

### SlCKX6, SL2.50ch12, 2211574

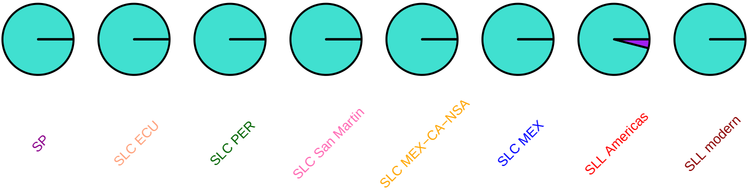

### SlCKX6, SL2.50ch12, 2212632

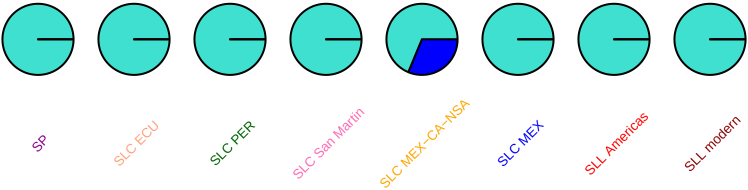

## AF308937, SL2.50ch12, 2922921

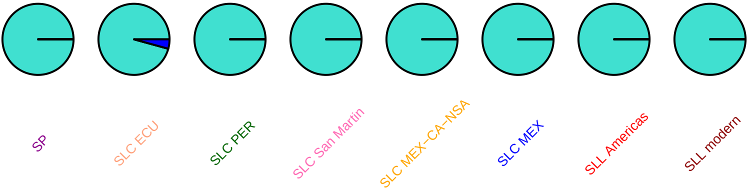

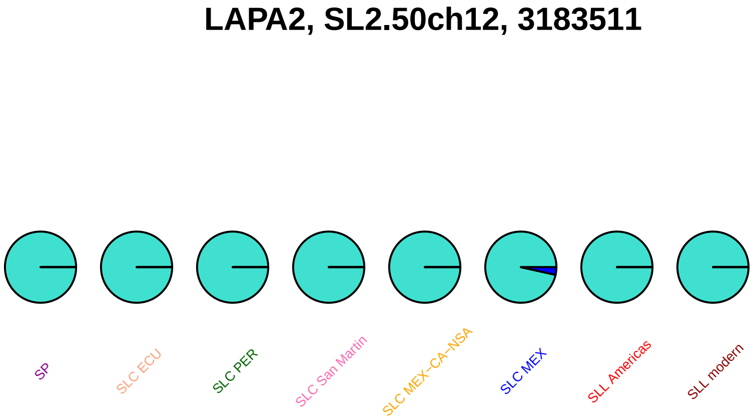
LAPA2, SL2.50ch12, 3183511
SP
SLC ECU
SLC PER
SLC San Martin
SLC MEX−CA−NSA
SLC MEX
SLL Americas
SLL modern

### txetb1, SL2.50ch12, 6426992

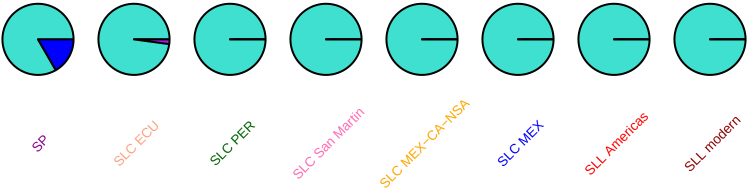

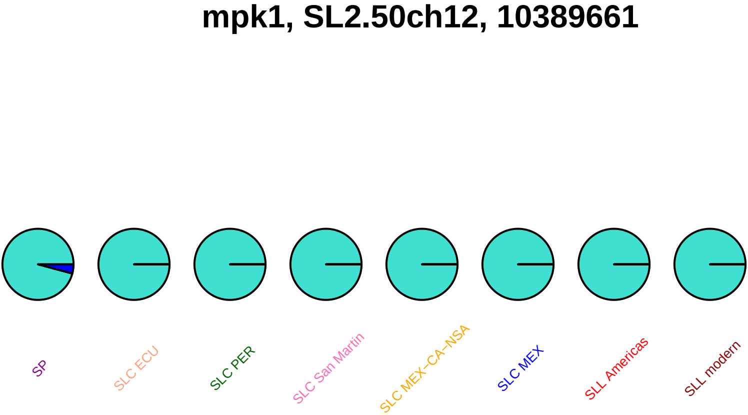

mpk1, SL2.50ch12, 10389661
SP
SLC ECU
SLC PER
SLC San Martin
SLC MEX−CA−NSA
SLC MEX
SLL Americas
SLL modern

# DQ117941, SL2.50ch12, 13969523

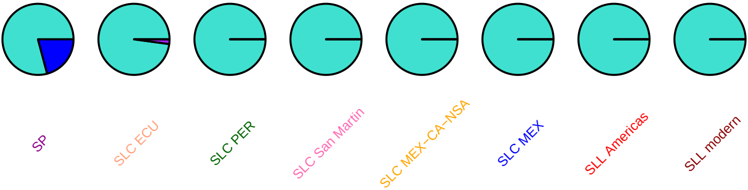

#### SlMYB11, SL2.50ch12, 37018817

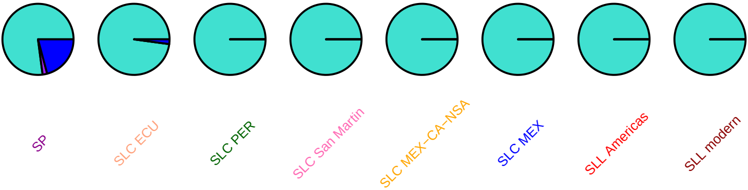

#### teg1b, SL2.50ch12, 37636204

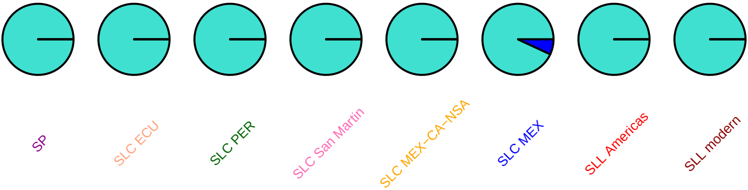

#### SlTIP2.1, SL2.50ch12, 38616084

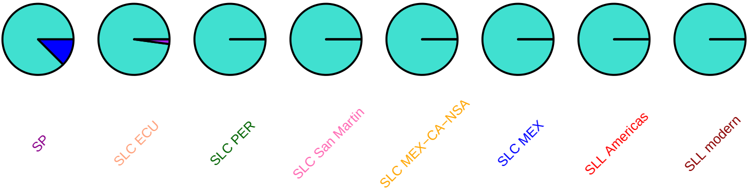

### SlWRKY43, SL2.50ch12, 39936412

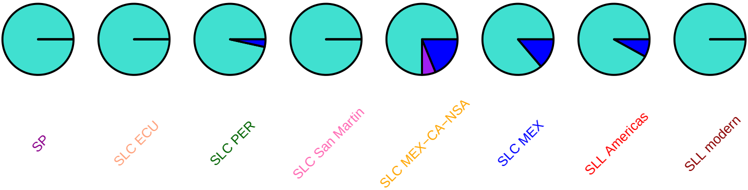

#### SlySBP6c, SL2.50ch12, 47392508

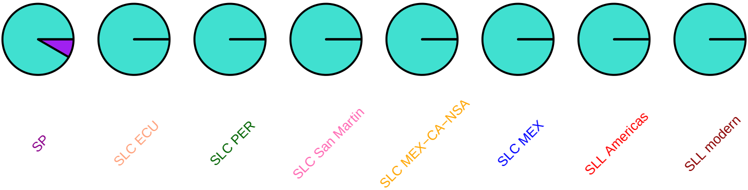

### SlC3H77, SL2.50ch12, 59227463

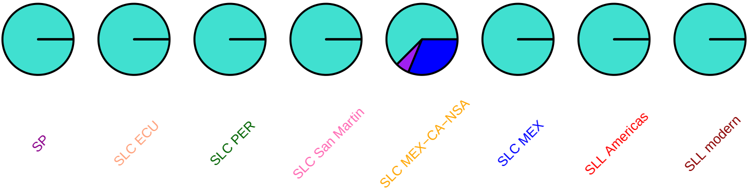

### SlPIP1.3, SL2.50ch12, 62177835

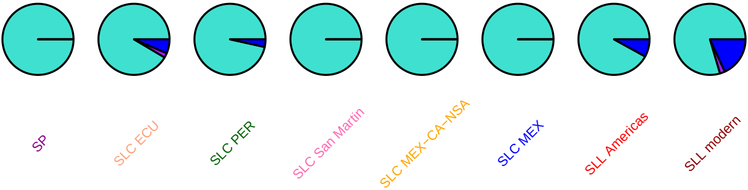

#### SlDEAD38, SL2.50ch12, 62301180

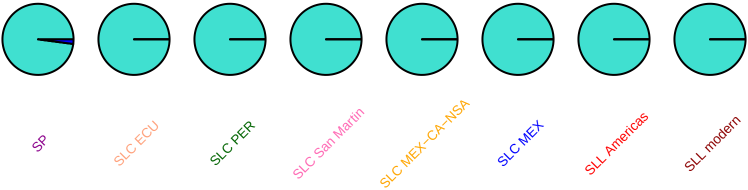

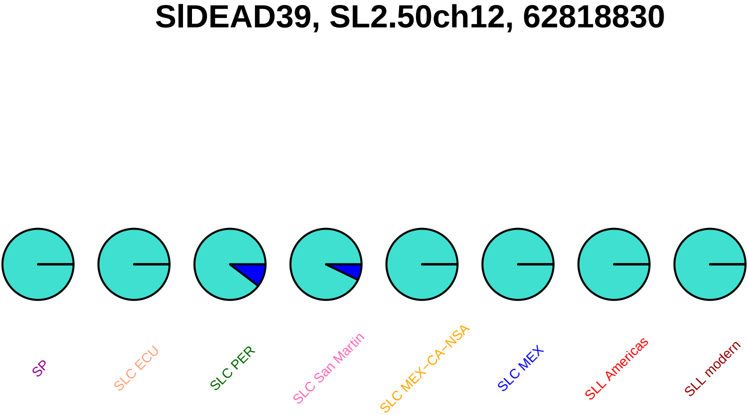

SlDEAD39, SL2.50ch12, 62818830
SP
SLC ECU
SLC PER
SLC San Martin
SLC MEX−CA−NSA
SLC MEX
SLL Americas
SLL modern

### SlWRKY61, SL2.50ch12, 62840579

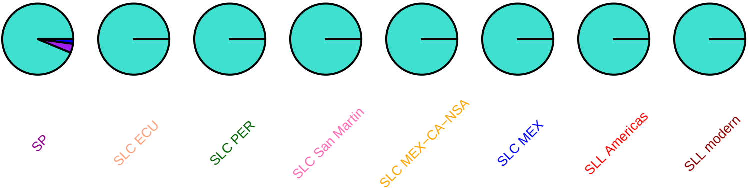

### SlBCAT1, SL2.50ch12, 63665930

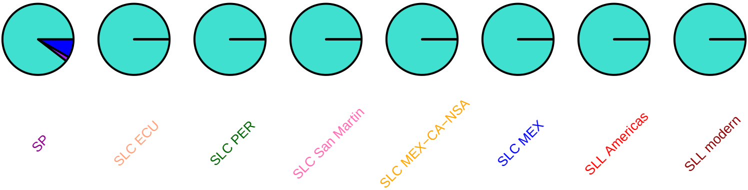

### SlBCAT1, SL2.50ch12, 63667758

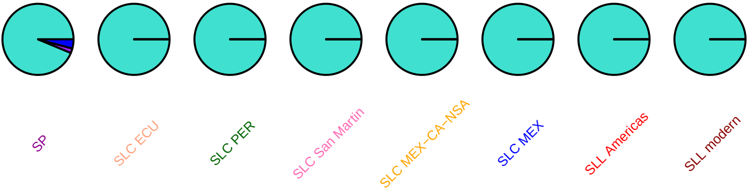

### SlBCAT1, SL2.50ch12, 63668581

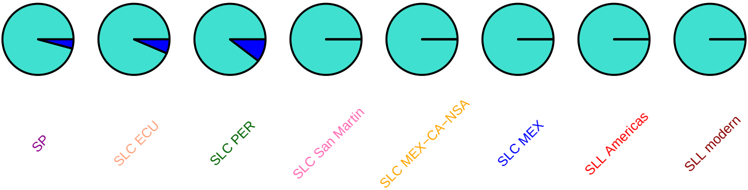

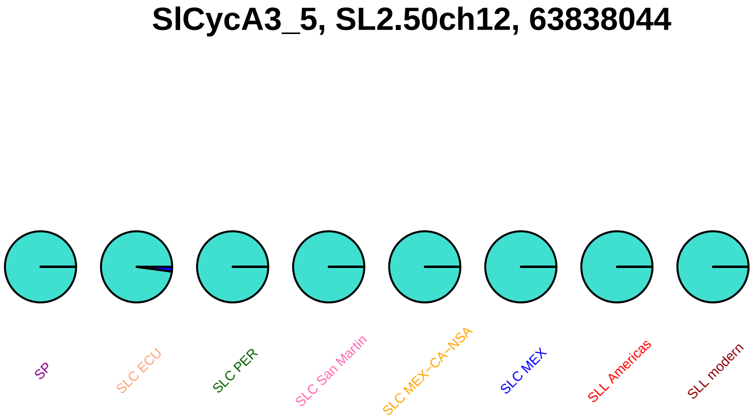
SlCycA3_5, SL2.50ch12, 63838044
SP
SLC ECU
SLC PER
SLC San Martin
SLC MEX–CA–NSA
SLC MEX
SLL Americas
SLL modern

### SlCycA3_5, SL2.50ch12, 63839669

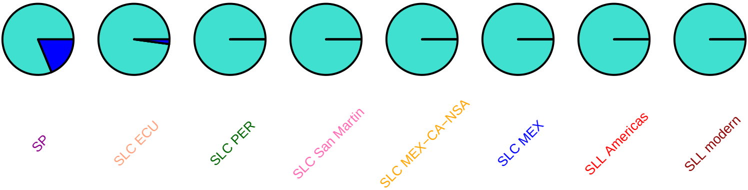

### SlCycA3_1, SL2.50ch12, 63854274

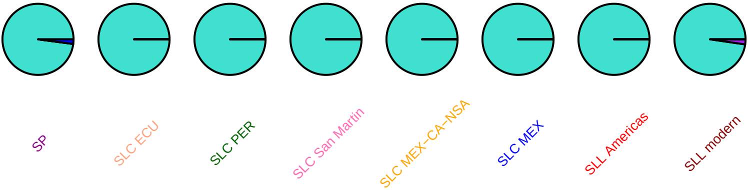

### SlSFP6, SL2.50ch12, 64297553

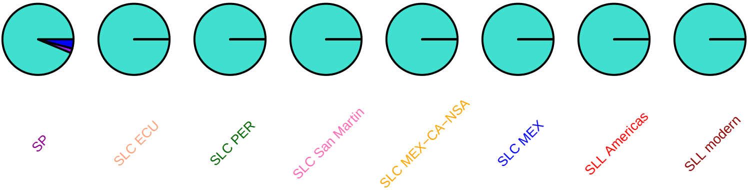

#### BRX, SL2.50ch12, 65168721

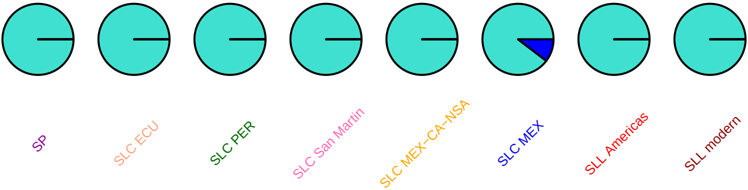

### IAA11, SL2.50ch12, 65673982

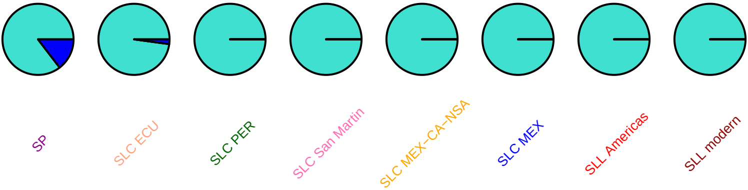

#### AIM1, SL2.50ch12, 66391855

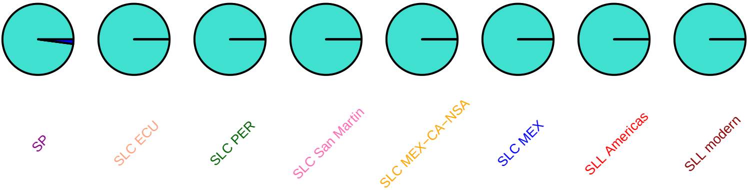

### SlMYB108, SL2.50ch12, 66395658

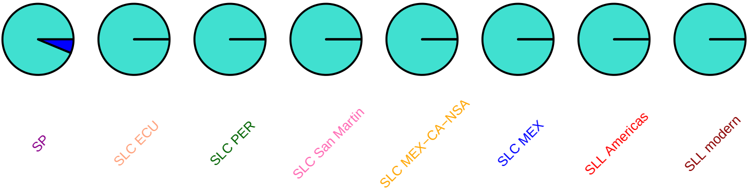

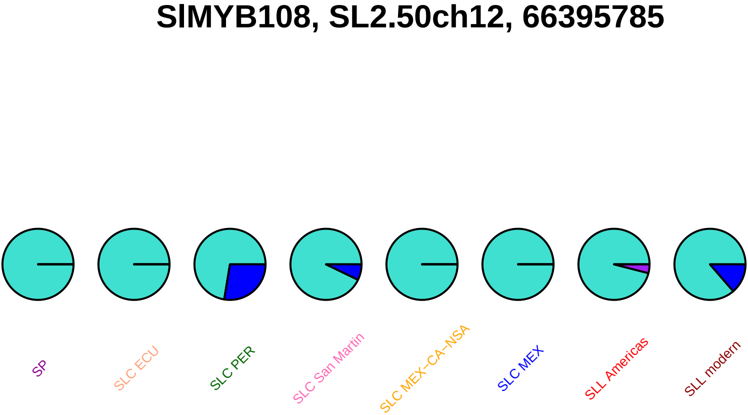
SlMYB108, SL2.50ch12, 66395785
SP
SLC ECU
SLC PER
SLC San Martin
SLC MEX−CA−NSA
SLC MEX
SLL Americas
SLL modern

#### SlMYB108, SL2.50ch12, 66396024

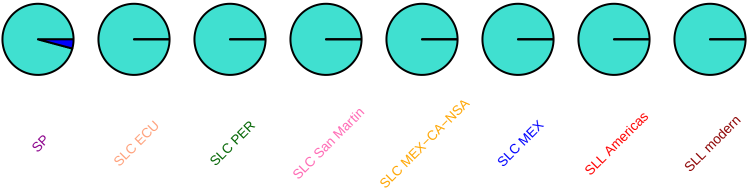

#### SlMAPKKK89, SL2.50ch12, 66453467

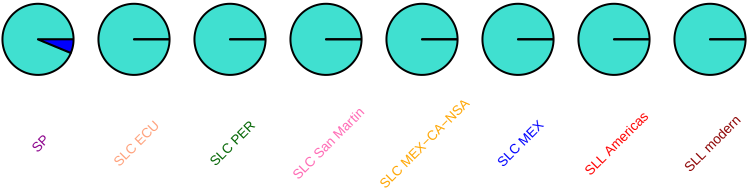

### gal83, SL2.50ch12, 66903828

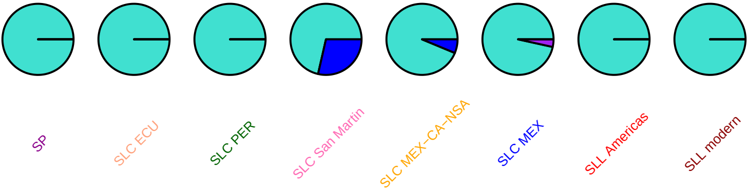

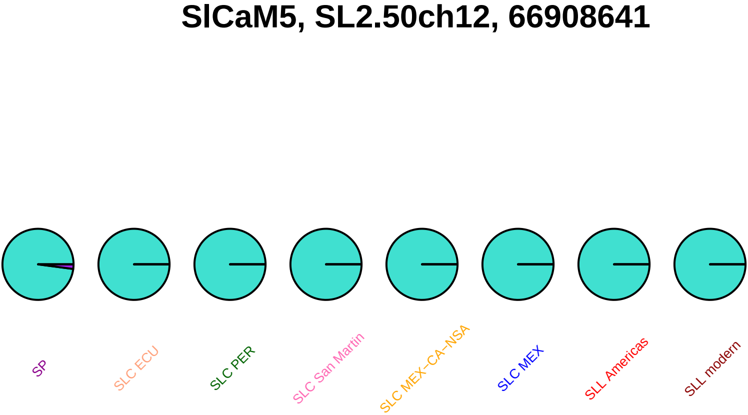
SlCaM5, SL2.50ch12, 66908641
SP
SLC ECU
SLC PER
SLC San Martin
SLC MEX−CA−NSA
SLC MEX
SLL Americas
SLL modern
